# Chemoenzymatically Synthesized *O*-Acetylated GD3 Gangliosides to Examine Viral Receptor Specificities in a Cellular Context

**DOI:** 10.1101/2025.08.15.670503

**Authors:** Zhiyong Zhang, Ruonan Liang, Kevin C. Hooijschuur, Robert P. de Vries, Zeshi Li, Geert-Jan Boons

## Abstract

Gangliosides are a class of sialic acid-containing glycosphingolipids involved in a wide range of biological processes. The terminal sialic acid of gangliosides can be *O*-acetylated at C7 and/or C9 hydroxyl, contributing to ganglioside structural complexity and function. It has been difficult to obtain panels of structurally well-defined *O*-acetylated gangliosides for binding and functional studies. We describe here a chemoenzymatic strategy that can provide, for the first time, 7-*O*-, 9-*O*-, and 7,9-di-*O*-acetylated GD3 gangliosides. It is based on the chemical assembly of a common tetrasaccharide precursor as α-glycosyl fluoride that is coupled to sphingosine by a glycosynthase, followed by *O*-acetyl editing by coronaviral hemagglutinin-esterases. The resulting synthetic glycosphingolipids have been employed for cell surface remodeling of erythrocytes. Analysis by liquid chromatography and ion mobility mass spectrometry (LC-IM-MS) demonstrated successful integration of the glycosphingolipids into the plasma membrane with preservation of acetyl ester patterns. Using human coronavirus HKU1 spike-functionalized virus-like particles, we demonstrate that the resulting glycan-remodeled erythrocytes can be utilized in hemagglutination studies as a label free method to investigate viral protein binding to individual glycoforms in a cellular environment.

**Entry for the Table of Contents:** Table of Contents
7-*O*-, 9-*O*-, and 7,9-di-*O*-acetylated GD3 gangliosides can be prepared by chemical assembly of a tetrasaccharide as α-glycosyl fluoride that is coupled to sphingosine using a glycosynthase, followed by *O*-acetyl editing by coronaviral hemagglutinin-esterases. Cell surface remodeling of erythrocytes using the GD3 derivatives provide a tool to examine viral receptor specificities.

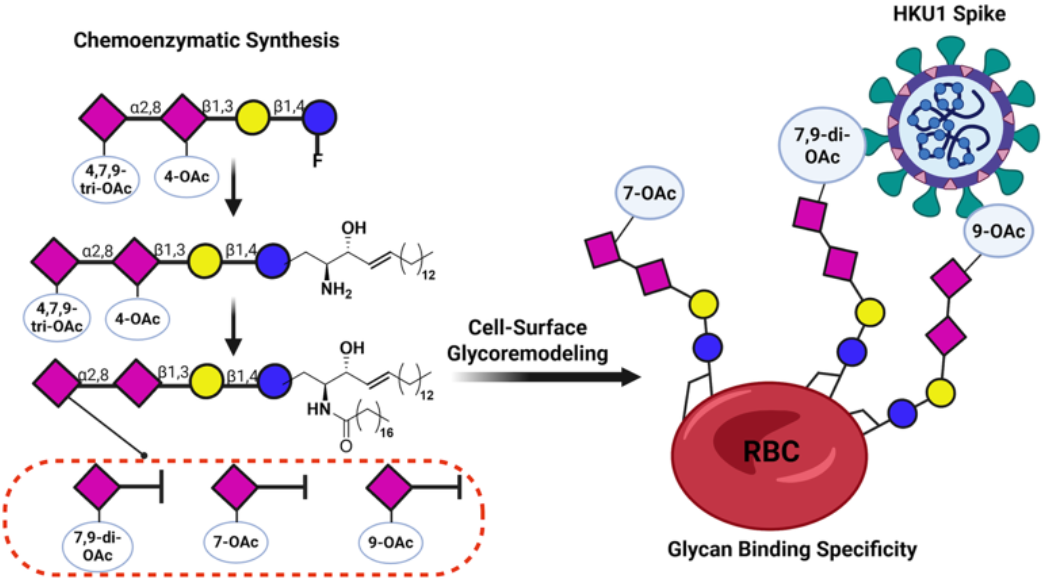

## Introduction

Gangliosides are a class of glycosphingolipids that have one or more sialic acid residues in their carbohydrate portion. They are integral components of the plasma membrane outer leaflet, and mediate membrane nanodomain formation.^[1]^ Gangliosides have been implicated in a broad range of (patho)physiological processes, such as cell growth,^[2]^ differentiation,^[3]^ and embryogenesis,^[4]^ neurobiology^[5]^ as well as malignancies^[6]^ and metastasis.^[7]^ At a molecular level, gangliosides dynamically interact with cellular components,^[8]^ such as transmembrane proteins,^[9]^ phospholipids,^[10]^ and cholesterols.^[11]^ For example, GM3, a mono-sialylated ganglioside, modulates epidermal growth factor receptor signaling *via* carbohydrate-carbohydrate interactions^[12–14]^ and electrostatic interaction with the protein transmembrane domain. Furthermore, gangliosides serve as ligands of immune modulatory proteins,^[15]^ exemplified by the engagement of GD3 with Siglec-7 on natural killer (NK) cells,^[16]^ thereby dampening NK cytotoxicity. Given their central roles in cellular membrane biology, gangliosides are used as entry receptors for toxins and microbes including AB toxins,^[17]^ and influenza C^[18]^ and betacoronaviruses.^[19–20]^

The terminal sialic acid residue of gangliosides can be acetylated at the C-7 and/or C-9 hydroxyl in a tissue/organ specific manner. Such a modification can have profound effects on ganglioside function. A notable example is GD3, which is a pro-apoptotic compound disrupting mitochondrial transmembrane potential.^[21]^ However, upon *O*-acetylation at the C-9 position, the pro-apoptotic effect is suppressed.^[22]^ 9-*O*-acetylated GD3 is an oncofetal marker for several malignancies including neuroblastoma, melanoma, and breast cancer, which implies a possible link with pro-survival function.^[23]^ The molecular mechanism of this functional inversion upon acetylation remains poorly understood. 7-*O*-acetylated GD3, which can non-enzymatically be converted to the 9-*O*-acetylated form, is even more elusive. The biosynthesis of *O*-acetylated GD3 is still controversial. It has been indicated that the acetyl ester is added to the C7 hydroxyl of sialic acid of intact gangliosides,^[24]^ and over time can migrate to C9. A more recent study indicated that the acetyl ester is attached at the stage of cytidine monophosphate linked sialic acid,^[25]^ which can function as a donor for relevant sialyltransferases. The elusive biosynthetic pathway and chemical lability of *O*-acetylated GD3 and other acetylated gangliosides make these compounds difficult to investigate, as they are not easily amenable to isolation or genetic manipulation. Well-defined *O*-acetylated glycosphingolipids are, however, needed to examine the biology of these compounds.

We report here a chemoenzymatic strategy for the concise synthesis of 7-*O*-(**1**), 9-*O*-(**2**), and 7,9-di-*O*-acetylated (**3**) GD3 gangliosides (Figure 1). Our approach involves the chemical assembly of a common tetrasaccharide precursor as α-glycosyl fluoride that is coupled to sphingosine by a glycosynthase,^[26]^ followed by *O*-acetyl editing by coronaviral hemagglutinin-esterases. We demonstrate that the chemically well-defined glycosphingolipids can be employed for cell surface glycosphingolipid remodeling of human erythrocytes. Liquid chromatography and ion mobility mass spectrometry (LC-IM-MS)-based analysis confirmed the successful incorporation of the glycosphingolipids into plasma membrane and retention of the acetyl ester patterns. Using human coronavirus HKU1 spike-functionalized virus-like particles, we demonstrate that the resulting glycan-remodeled erythrocytes can be utilized in hemagglutination studies as a label free method to investigate viral protein binding to individual glycoforms in a cellular environment.

**Figure 1.**
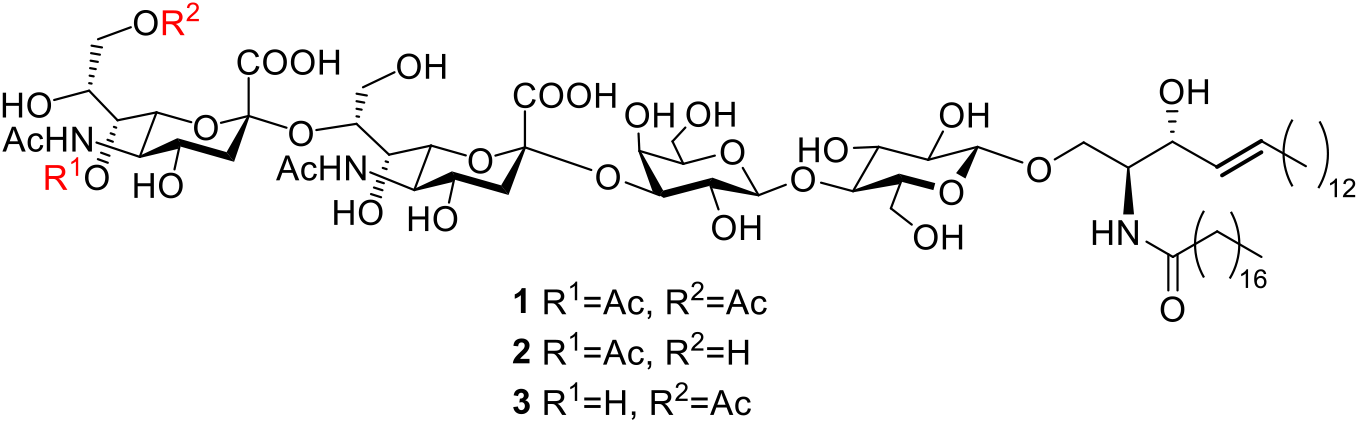
Structure of *O*-acetylated GD3 gangliosides.

## Results and Discussion

The synthesis of intact *O*-acetylated GD3 glycosphingolipids **1**-**3** poses several challenges. The acetyl esters are labile, and prone to migration (pH > 7) or hydrolysis (pH > 9) that hinders the use of basic reaction condition at a late stage of synthesis. Furthermore, removal of protecting groups by hydrogenation must be performed prior to sphingosine installation, as it would otherwise reduce the alkene of the lipid. We envisioned that glycosphingolipid **4** containing 4,7,9-tri-*O*-acetyl esters on the terminal sialic acid and a 4’-*O*-acetyl on the penultimate sialic acid would be an appropriate precursor for the preparation of the target compounds **1**-**3** (Scheme 1). Based on our previous studies,^[19]^ it was expected that murine coronavirus (MHV) hemagglutinin-esterases (HEs) can selectively cleave the acetyl esters at C-4 of the terminal and penultimate sialoside. Subsequent treatment with the HE of bovine coronavirus (BCoV) would result in the removal of the C-9 acetyl ester. The remaining acetyl ester at C-7 can then be migrated to C-9 by treatment with 50 mM NH_4_HCO_3_ (pH 8.0). Acylation of the amine of the sphingosine moiety of the resulting compounds would then provide various acetylated forms of GD3. It was expected that key intermediate **4** could be prepared by the condensation of α-glycosyl fluoride **6** with sphingosine using a glycosynthase mutant of endoglycoceramidase-II (EGC-II, D314Y/E351S) reported by the Withers and co-workers.^[26–27]^ It was envisaged that sialyl donors **9** and **10**, and lactosyl fluoride **8** could readily be assembled to give tetrasaccharide **7**, which upon deprotections would provide fluoride **6**.

**Scheme S1.**
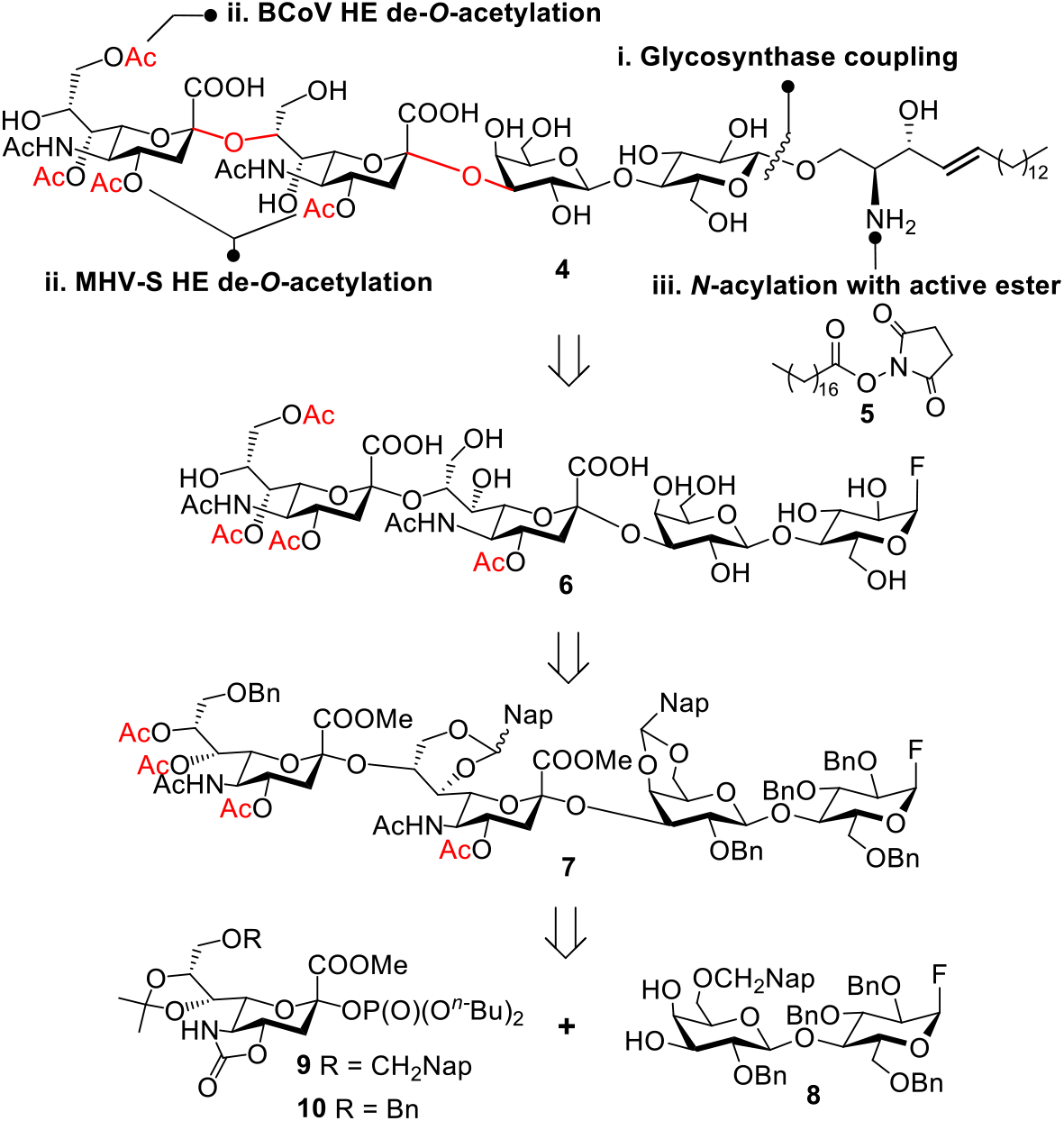
Synthetic strategy for *O*-acetylated GD3 targets as intact glycosphingolipids. Abbreviations: Bn, benzyl; Nap, 2-naphthyl; Bu, butyl.

The preparation of the lactosyl fluoride **8** began with the coupling of thioglucoside acceptor **S10** with galactosyl donor **S6** to give a disaccharide that was subjected to a four-step protecting group interconversion procedure without any intermediate purification to provide **11** in overall yield of 51% (see SI). Lactosyl thiophenol **11** was converted into glycosyl fluoride **12** as only the α-anomer by treatment with Barluenga reagent (IPy_2_BF_4_) (Scheme 2, see SI for NMR characterization).^[28–29]^ Compound **12** was treated with wet trifluoroacetic acid to remove the isopropylidene acetal to afford lactosyl 3,4-diol **8** in a yield of 59% over two-steps. Subsequently, α-selective sialylation of acceptor **8** using sialyl donor **9** in the presence of a catalytic amount of TMSOTf gave an immediate trisaccharide that was immediately subjected to wet TFA to remove the isopropylidene acetal providing the GM1 trisaccharide acceptor **13** in an overall yield of 82%. Next, an α2,8-sialylation of **13** was performed using sialyl donor **10** in the presence of TMSOTf to give a tetrasaccharide that was again subjected to TFA-mediated to remove the isoproplylidene acetal to give a diol. The latter compound was subjected to oxidation by 2,3-dichloro-5,6-dicyano-1,4-benzoquinone (DDQ) under anhydrous conditions to oxidize the Nap ether to the corresponding carbocation that underwent ring closure with the C-7 alcohol to give 2-naphthylmethylene acetal containing **15** in an overall yield of 50% as diastereoisomeric mixture of exo- and endo-products. This conversion blocked the remaining alcohol at the penultimate sialoside with a base-stable and hydrogenolytically removable protecting group allowing late-stage *O*-acetylation of the terminal sialoside. Compound **15** was subjected to a four-step deprotection procedure. It started with saponification of the C1 methyl esters and the oxazolidinone moieties of the sialic acid residues followed by selective sialate-5-*N*-acetylation in the same pot. Next, the carboxylic acids of the sialosides were protected as benzyl ester by reaction with benzyl bromide in the presence of K_2_CO_3_, and finally the remaining alcohols were acetylation under standard conditions to afforded **7** (32% over four steps). Fluoride **7** was subjected to hydrogenation over Pd(OH)_2_ to remove the benzyl ethers and ester to give GD3 glycosyl fluoride **6** in an isolated yield of 63%. Gratifyingly, the α-fluoride had remained intact throughout the entire chemical assembly and deprotection steps. The 8-*O*-acetyl ester of the terminal sialic acid had spontaneously migrated to C-9 during hydrogenation step, which was confirmed by ^1^H-NMR and HMBC experiments (Figure S2), and therefore a separate acetyl migration step was not necessary. The presence of *O*-acetyl ester at C9 was confirmed by the chemical shift analysis of the geminal H-9 protons at 4.07 ppm, which is more downfield than when C9 is not acetylated (4.02 ppm and 3.73 ppm), and by a clear HMBC correlation signal with the carbonyl carbon of the acetyl ester.

**Scheme S2.**
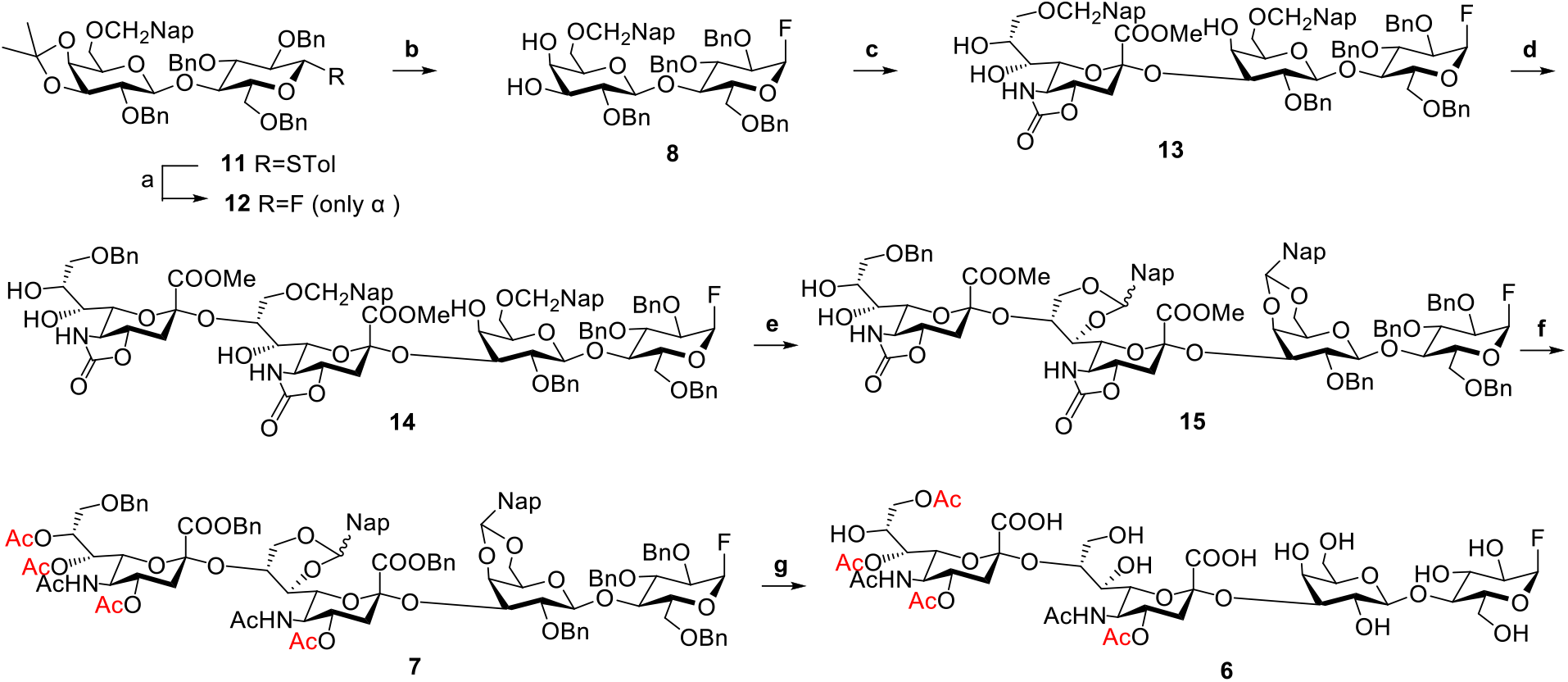
Synthesis of tetrasaccharide fluoride **6**. Reagents and conditions: (a) Barluenga reagent, CH_2_Cl_2_, 0 to 21 °C. (b) wet TFA, CH_2_Cl_2_, 21 °C, 59% over 2 steps. (c) **i. 9**, TMSOTf, −78 °C; **ii**. wet TFA, CH_2_Cl_2_, 21 °C, 82% over 2 steps. (d) **i. 10**, TMSOTf, −78 °C; **ii**. wet TFA, CH_2_Cl_2_, 21 °C, 83% over 2 steps. (e) DDQ, CH_2_Cl_2_, 21 °C, 50%. (f) **i**. KOH, dioxane/H_2_O (1:1), 50 °C; **ii**. Ac_2_O, CH_2_Cl_2_, 21 °C; **iii**. BnBr, K_2_CO_3_, 0 to 21 °C; **iv**. Ac_2_O, pyridine, *N,N*-dimethyl-4-aminopyridine (DMAP), 0 to 21 °C, 32% over 4 steps. (g) Pd(OH)_2_, H_2_, dioxane/H_2_O (1:1), 21 °C, 63%.

Enzymatic glycosylation of sphingosine with glycosyl fluorides using a mutant endoglycoceramidase is a versatile approach to prepare glycosphingolipids.^[26]^ Here, we examined the feasibility of transfer of chemically synthesized anomeric fluorides having an *O*-acetylated sialoside to sphingosine using a mutated endoglycoceramidase (EGC). Previously, it was reported that the double mutant EGC E351S/D314Y has increased transfer efficiency (EGC II)^[27]^ and therefore this enzyme was employed for our studies. Thus, 4,4’,7,9-tetra-*O*-acetylated tetrasaccharide fluoride **6** and sphingosine were incubated with EGC II in NaOAc buffer (pH 5.0) resulting in the facile formation of **4**, which was isolated in a yield of 65% (Scheme 3). The acidic reaction conditions ensured EGC-II operates optimally without acetyl ester migration taking place. Subsequently, exposure of **4** to MHV-S HE resulted in the selective hydrolysis of the C-4 acetyl esters on the terminal and penultimate sialic acid residues, affording 7,9-di-*O*-acetylated sialoside **16** in a yield of 81%. On the other hand, treatment of **4** with a combination of HEs from MHV-S and BCoV to hydrolyze all other acetyl esters, except the 7-*O*-acetyl ester, gave C-7 monoacetylated **17** in a yield of 83%. Compounds **16** and **17** were *N*-acylation using the *N*-hydroxysuccinimide (NHS) ester of stearic acid (18:0, **5**) in DMSO in the presence of cesium fluoride to afford target compounds **1** (83% yield from **16**) and **2** (80% yield from **17**). 9-*O*-acetylated **3** was obtained by treating **2** under mildly basic condition overnight (50 mM NH_4_HCO_3_, pH = 8). Gratifyingly, the *N*-acylation condition was compatible even with the most labile 7-*O*-acetyl ester. The glycolipids were purified using C18 reverse phase chromatography, yielding the final products in high purity. The preservation of the acetyl ester patterns was confirmed by 2D NMR experiments (Figures S3 and S4).

Next, we examined the recognition of the synthetic glycosphingolipids by a viral protein when inserted into the plasma membrane of cells. The receptor specificity of viruses is a critical determinant of host range and tissue/organ tropism.^[30]^ Previously, we demonstrated that the spike proteins the human coronaviruses OC43 and HKU1, which both are of zoonotic origin, have a preference for binding α2,8-linked di-sialoside modified by 9-*O*-acetyl ester.^[19]^ Both virolectins also bound di-sialosides having an additional C-7 acetyl ester, but with a slightly lower responsiveness. Related animal viruses exhibited distinct selectivities for other *O*-acetylation forms and sialosidic linkage types, indicating convergent evolution by adapting to the sialoglycome of the human respiratory tract. 9-*O*-Acetylated α2,8-linked disialosides are common constituents of gangliosides such as GD3 (9-Ac-GD3). At the plasma membrane, gangliosides cluster in microdomains, where they can engage in multivalent interactions resulting in high avidity of binding and endocytosis.^[31]^ Our previous studies had also shown that overexpression α2,8-sialyltransferase 1 (ST8SIA1), which is involved in the biosynthesis of 9-Ac-GD3, results in much strong binding of OC43 and HKU1 S1^A^-Fc fusion proteins,^[19]^ indicating that gangliosides such as 9-Ac-GD3, can function as receptors for these viruses.

Here, we aimed to examine pharmacologically whether the spike protein of HKU1 can engage with GD3 modified by acetyl esters when presented on the plasma membrane. To accomplish this goal, we embarked on a cell-surface glycan remodeling strategy by exposing human erythrocytes to the synthetic GD3 gangliosides **1**-**3** (Figure 2A). Erythrocytes were chosen because these are amenable to hemagglutination, which is widely employed to examine viral receptor specificity owing to a fast and label free visual readout.^[32]^ First, human erythrocytes were incubated with the broad-acting neuraminidase from *Arthrobacter ureafaciens* to remove endogenous sialosides followed by washing to remove the neuraminidase. Next, the cells were incubated with *O*-acetylated GD3 gangliosides **1, 2**, and **3** (2.5 μM) and as a control non-acetylated GD3 (**18**) at pH 6.5 to minimize acetyl ester migration.

**Scheme S3.**
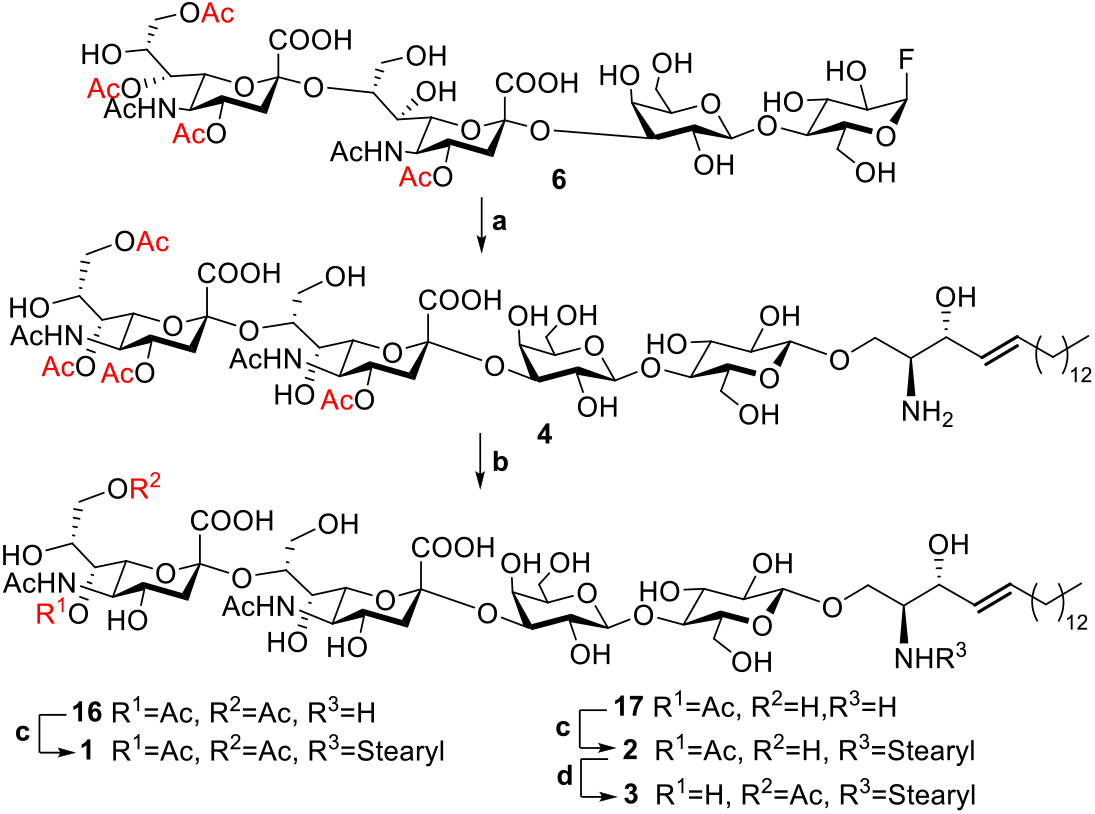
Synthesis of intact *O*-acetylated GD3 gangliosides. Final steps include enzymatic sphingosine coupling, acetyl ester editing, and fatty acid attachment. Reagents and conditions: (a) Sphingosine, EGC-II, Triton X-100, 25 mM NaOAc Buffer (pH 5.0), 37 °C, 65%. (b) MHV-S HE, 50 mM HCOONH_4_, 37 °C for **16**, 81%; MHV-S HE and BCoV HE, 50 mM HCOONH_4_, 37 °C for **17**, 83% (c) stearic acid-NHS ester **5**, CsF, DMSO, 21°C, 83% (**1** from **16**), 80% (**2** from **17**). (d) 50 mM NH_4_HCO_3_ (pH 8.0), 37 °C, quant.

**Figure 2.**
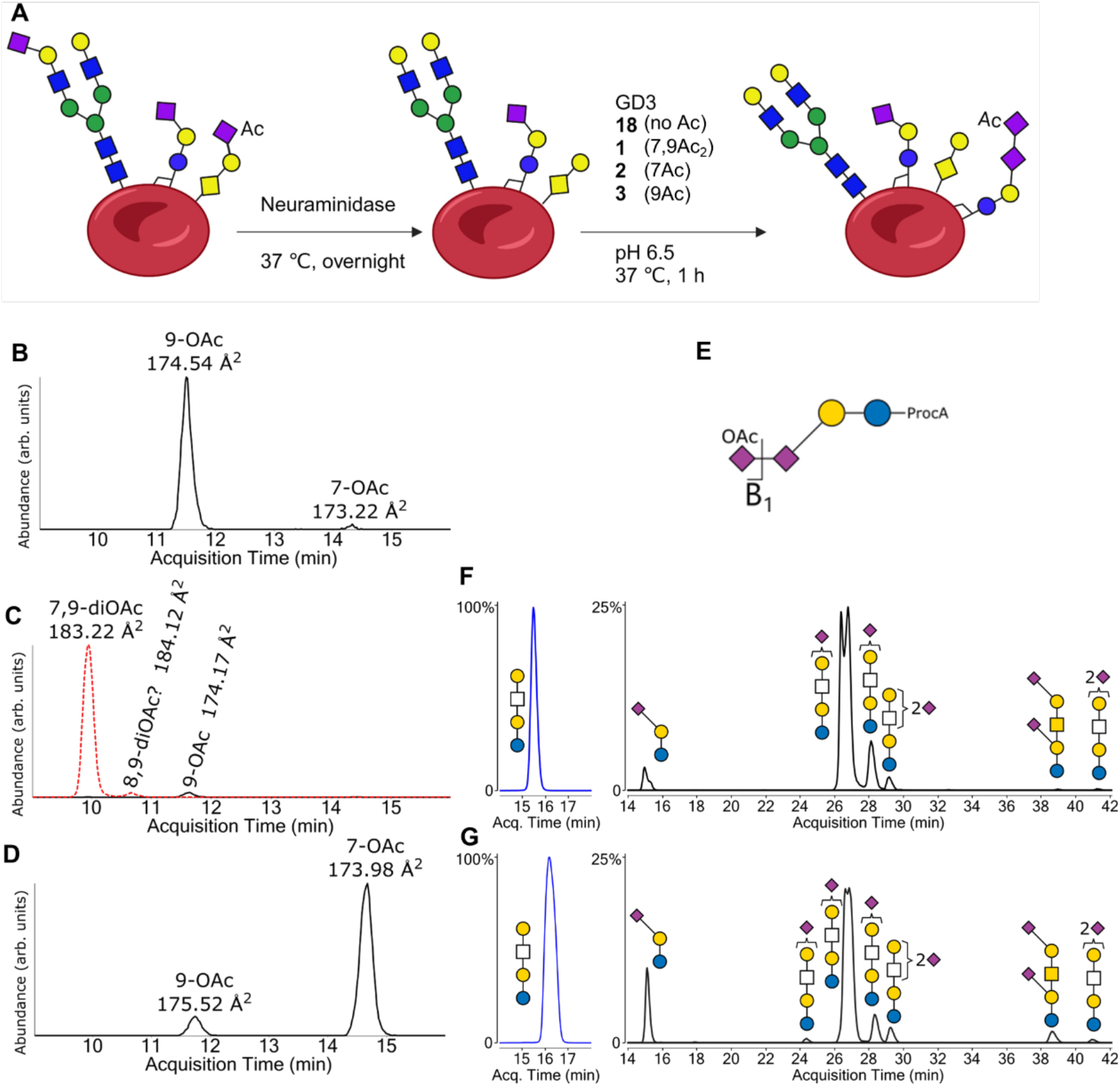
Human erythrocytes remodeling with the synthetic gangliosides and glycolipid profiling for remodeled cells. A) Schematic representation of the procedure for erythrocytes remodeling with the synthetic gangliosides. B) B_1_ fragment used to identify sialic acid *O*-acetylation forms. Abbreviations: ProcA, procainamide. C) – E) Extracted ion chromatograms for mono-*O*-acetylated GD3 (black, solid) and di-*O*-acetylated GD3 (red, dashed) from erythrocytes remodeled with C) 9-*O*-acetylated, D) 7,9-di-*O*-acetylated, and E) 7-*O*-acetylated GD3 with corresponding collision cross section (CCS) values of the *O*-acetylated sialic acid B_1_ fragment displayed at the peak positions. F) and G) Extracted ion chromatograms for representative glycosphingolipids in F) non-treated control and G) neuraminidase-treated erythrocytes. The abundance of the non-sialylated tetrasaccharide was set to 100%. The relative abundances of sialylated structures are shown in a separate chromatogram for clarity. Symbols: magenta diamond, *N*-acetylneuraminic acid; yellow circle, galactose; blue square, *N*-acetylglucosamine; blue circle, glucose; green circle, mannose; empty square, *N*-acetyl hexosamine.

Next, we confirmed the incorporation of the synthetic gangliosides into the plasma membrane of the erythrocytes by employed a liquid chromatography and ion mobility mass spectrometry (LC-IM-MS) approach that can elucidate exact *O*-acetylation patterns of sialosides of glycoconjugates isolated from complex biological samples.^[33]^ It is based on the observation that the B_1_ fragment of isolated sialosides give unique collisional cross section (CCS) values for the various naturally occurring acetylation patterns (9-*O*-acetyl: 174.7 Å^2^, 7-*O*-acetyl: 173.2 Å^2^, and 7,9-di-*O*-acetyl: 183.4 Å^2^, Figure 2E).^[33]^ Thus, glycosphingolipids of the native, neuraminidase treated, and glycan-remodeled erythrocytes were extracted using methanol/chloroform.^[34]^ The glycans were released by treatment with EGCase I and then labeled with procainamide and cleaned up by C18 and porous graphitized carbon solid-phase extraction followed by LC-IM-MS analysis (Figure 2B-D). By a targeted search for *m/z* values of specific glycans, we found that native and neuraminidase-treated erythrocytes did not contain any *O*-acetylated GD3, which was further confirmed by the absence of sialic acid B_1_ fragments for *O*-acetylated sialic acids (Figure S5). On the other hand, *O*-acetylated GD3-remodeled erythrocytes displayed abundantly the exogenously administered glycosphingolipids, which was unambiguously confirmed by the *m/z* values of both unfragmented molecular ion peaks and the B_1_ fragments with matching CCS values. It showed that the majority of *O*-acetylated GD3 had remained in their original forms (Table S1). The 9-*O*-acetylated sialoside had remained fully unchanged. Interestingly, a very small fraction of 7,9-di-*O*-acetylated GD3 had been converted into a different *O*-acetylated regio-isomer (peak at ~11.8 min). It is likely that the acetyl ester on C7 had migrated to the C8 hydroxyl during the incubation yielding 8,9-di-*O*-acetylated GD3.^[35]^ This result is in line with a previous observation that 7,9-di-*O*-acetylated sialic acid can convert into the 8,9-di*-O*-acetylated form. We also observed that ~10% of 7-*O*-acetylated GD3 had undergone acetyl migration giving rise to the 9-*O*-acetylated form, despite the buffer pH was kept at 6.5 during glycosphingolipid incorporation.

Interestingly, the LC-IM-MS analysis of the native and neuraminidase-treated cells showed that the neuraminidase had not removed the sialoside of the glycosphingolipids (Figure 2F,G). As an internal standard, the relative abundance of a non-sialylated tetrasaccharide (Gal-HexNAc-Gal-Glc) was set to 100% for both samples. The relative abundance of various sialylated structures, including GM3, GD1a, and mono- and disialylated Gal-HexNAc-Gal-Glc tetrasaccharides were similar before and after neuraminidase treatment. It indicates that gangliosides, when present on the erythrocyte cell surface, are not readily accessible for enzymatic degradation. In this study, the focus is on receptor binding specificities of acetylated sialic acids and thus a lack of digestion does not affect the interaction of the glycan-binding protein and will not cause ambiguities.

Having confirmed the successful incorporation of GD3 glycosphingolipids into the cell surface of erythrocytes, we performed hemagglutination assays using the spike protein of the human coronavirus HKU1 (Figure 3A). The receptor-binding N-terminus domain (S1^A^) of HKU1 spike, fused with human Fc tag, was recombinantly expressed in HEK293T GnTI-deficient cells. The HKU1 S1^A^-Fc chimera was then presented on a protein A-fused lumazine synthase nanoparticle (pA-LS), which is strategy to mimic viral particles.^[36–37]^ The HKU1 S1^A^-Fc-functionalized nanoparticles hemagglutinated native human erythrocytes at moderate titers (Figure 3B). This indicates that native human erythrocytes endogenously express 9-*O*-acetylated sialosides on glycoconjugates other than glycosphingolipids. Neuraminidase treatment completely abolished hemagglutination. The addition of 9-*O*-acetylated and 7,9-di-*O*-acetylated GD3 at 2.5 μM restored hemagglutination at a significantly higher titers than for native erythrocytes. The addition of non-acetylated GD3 did not result in any hemagglutination, which is in agreement with the binding mode of HKU1 spike, which has an absolute requirement for 9-*O*-acetylation.^[38]^ The addition of 7-*O*-acetylated GD3 resulted in hemagglutination at moderate titers, which is likely caused by the minor presence of 9-*O*-acetylated GD3 (~10% of total GD3 added) as acetyl migration product. As expected, the administration of a lower concentration 9-*O*-acetylated GD3 (0.25 μM) resulted in lower hemagglutination titers without an observable change in specificity (Figure S6).

**Figure 3.**
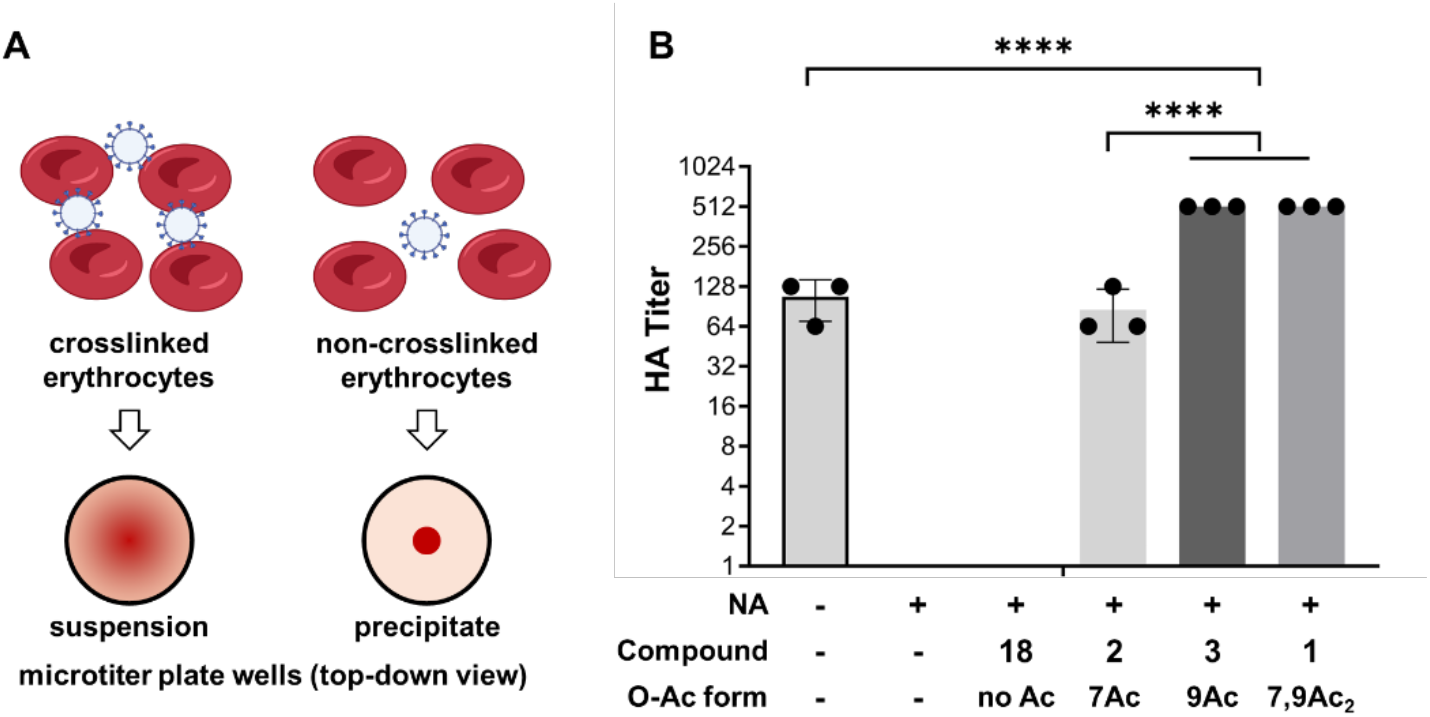
Hemagglutination assays of GD3-remodeled human erythrocytes. A) Schematic representation of visual readouts for hemagglutination. B) Hemagglutination (HA) titers for differently remodeled erythrocytes. HA titer is defined as the largest dilution factor of spike-funcitonalized nanoparticles at which erythrocytes remain in suspension. Bars indicate the mean +/-standard deviation of 3 independent assays. None of the wells for NA-treated or non-acetylated GD3-added samples gave cell suspensions and therefore did not give an HA titer. Statistical significance (two-tailed) calculated by Welch’s unequal variances *t*-test are indicated in the graphs with asterisks. Abbreviations: NA, neuraminidase A treatment.

## Conclusions

We have developed a facile chemoenzymatic strategy that can provide intact GD3 gangliosides having all possible naturally occurring *O*-acetylation patterns, including 7-*O*-, 9-*O*-, and 7,9-di-*O*-acetyl esters. It is the expectation that the method can be applied to the preparation of any naturally occurring *O*-acetylated ganglioside. Earlier reported synthetic strategies for GD3 generally involve the assembly of a tetrasaccharide that is coupled to sphingosine, which is then acylated at the amine of sphingosine, followed by global deprotection. The latter procedures usually involve treatment with strong base which is not compatible with the presence of acetyl esters.^[39–46]^ To access 9-*O*-acetylated GD3, Wong and co-workers reported a post-synthetic modification strategy in which the C-9 hydroxyl of GD3 is acetylated using a lyase in combination with an acetyl donor.^[47]^ Other *O*-acetylated forms of GD3 and other gangliosides have remained synthetically inaccessible. Here, we addressed this deficiency by taking advantage of substrate specificities of HEs to edit *O*-acetylation patterns of a tetra-*O*-acetylated GD3 oligosaccharide linked to sphingosine. The latter compound could be prepared by exploiting the finding that the anomeric fluoride of tetra-*O*-acetylated GD3 is a substrate for a mutant form of an endoglycoceramidase^[27]^ and can be condensed with sphingosine. Acylation of the amine of sphingosine gave entry into all possible acetyl ester forms of GD3. We demonstrated that the synthetic gangliosides could readily be incorporated into the membranes of human erythrocytes. The resulting cell-surface engineered cells were analyzed by LC-IM-MS, which confirmed the presence of exogenously introduced GD3 and preservation of *O*-acetylation patterns of the sialosides. Even the most base-sensitive 7-*O*-acetylated GD3 had remained largely intact providing opportunities to investigate the biological functions of these elusive molecules. We employed the GD3-engineered erythrocytes to examine receptor specificities of the spike of a human coronavirus by hemagglutination. This binding approach is attractive because it is label-free and provides an easy to implement tool for receptor specificity studies. Here, we employed nanoparticles displaying multiple copies of the spike of the human corona virus HKU1. The hemagglutination results agree with our previous observations that HKU1 can bind 9-mono- and 7,9-di-*O*-acetylated GD3 oligosaccharides. A surprising finding was that wildtype human erythrocytes could also be agglutinated by the nanoparticles displaying HKU spikes, albeit with a much low titer. The IM-MS analysis of gangliosides isolated from human red blood showed an absence of structures modified by acetyl esters, and thus another class of *O*-acetylated sialoglycan must be responsible for hemagglutination. Recently, we found that when HKU1 NTD-Fc chimeras is presented on nanoparticles, it can in addition to 9-*O*-acetylated α2,8-linked disialoside also bind to 9-*O*-acetylated α2,3-linked sialylated LacNAc structures albeit with much lower responses.^[37]^ α2,3-Linked *O*-acetylated sialosides have also been observed on *O*-glycans,^[48–49]^ and thus it is like that the wildtype human erythrocytes are agglutinated by the presence of such structures. Respiratory viruses need to penetrate a thick mucus layer before arriving at the cell surface where they need to engage with a receptor for cell entry. Probably, the low affinity interaction of the spike of HKU1 with *O*-glycans of mucus contribute to virion attachment and mucus penetration. The higher affinity binding to 9-*O*-acetylated α2,8-linked disialoside typical of gangliosides, causes a conformational change of the spike^[50]^ which exposes subdomain S1^B^ to bind with the proteinaceous cell surface receptor transmembrane serine protease 2 (TMPRSS2).^[51–52]^ Thus, it is likely that a combination of low affinity and high affinity binding contributes to infectivity. It is the expectation that synthetic *O*-acetylated gangliosides will facility monitoring the evolution of sialoglycan binding specificity and host tropism of emerging pathogens. It will also make it possible to examine other biological properties of gangliosides having distinct *O*-acetylated sialosides.

## Experimental Section

Experimental details can be found in the Supporting Information. The Supporting Information (PDF) contains Figures S1-S6, Table S1, Materials and General Methods, Synthetic Methods and Compound Characterization, and NMR spectra.

## Acknowledgements

This research was supported by the European Commission (grants nr. 101020769), Health~Holland (grant nr. TKI-LSHM21030) and the Chinese Scholarship Council (to Z.Z and to R.L).

## Data Availability Statement

The data that support the findings of this study are available in the article and its Supporting Information.

## References

[1] L. Cantu, M. Corti, P. Brocca, E. Del Favero. Structural aspects of ganglioside-containing membranes. Biochim. Biophys. Acta 2009, 1788, 202–208.

[2] E. G. Bremer, J. Schlessinger, S. Hakomori. Ganglioside-mediated modulation of cell growth. Specific effects of GM3 on tyrosine phosphorylation of the epidermal growth factor receptor. J. Biol. Chem. 1986, 261, 2434–2440.

[3] W. Shen, R. Falahati, R. Stark, D. Leitenberg, S. Ladisch. Modulation of CD4 Th cell differentiation by ganglioside GD1a in vitro. J. Immunol. 2005, 175, 4927–4934.

[4] D. H. Kwak, B. B. Seo, K. T. Chang, Y. K. Choo. Roles of gangliosides in mouse embryogenesis and embryonic stem cell differentiation. Exp. Mol. Med. 2011, 43, 379–388.

[5] Z. Guo. Ganglioside GM1 and the central nervous system. Int. J. Mol. Sci. 2023, 24, 9558.

[6] N. Sasaki, M. Toyoda, T. Ishiwata. Gangliosides as signaling regulators in cancer. Int. J. Mol. Sci. 2021, 22, 5076.

[7] C. Cumin, Y. L. Huang, C. Rossdam, F. Ruoff, S. P. Cespedes, C. Y. Liang, F. C. Lombardo, R. Coelho, N. Rimmer, M. Konantz, M. N. Lopez, S. Alam, A. Schmidt, D. Calabrese, A. Fedier, T. Vlajnic, M. von Itzstein, M. Templin, F. F. R. Buettner, A. Everest-Dass, V. Heinzelmann-Schwarz, F. Jacob. Glycosphingolipids are mediators of cancer plasticity through independent signaling pathways. Cell Rep. 2022, 40, 111181.

[8] C. A. Lingwood. Glycosphingolipid functions. Cold Spring Harb Perspect Biol 2011, 3.

[9] S. Hakomori, K. Handa. Interaction of glycosphingolipids with signal transducers and membrane proteins in glycosphingolipid-enriched microdomains. Methods Enzymol. 2003, 363, 191–207.

[10] B. Maggio, F. A. Cumar, R. Caputto. Interactions of gangliosides with phospholipids and glycosphingolipids in mixed monolayers. Biochem. J. 1978, 175, 1113–1118.

[11] M. R. Morrow, D. Singh, D. Lu, C. W. Grant. Glycosphingolipid fatty acid arrangement in phospholipid bilayers: cholesterol effects. Biophys. J. 1995, 68, 179–186.

[12] S. J. Yoon, K. Nakayama, T. Hikita, K. Handa, S. I. Hakomori. Epidermal growth factor receptor tyrosine kinase is modulated by GM3 interaction with N-linked GlcNAc termini of the receptor. Proc Natl Acad Sci U S A 2006, 103, 18987–18991.

[13] S. J. Yoon, K. Nakayama, N. Takahashi, H. Yagi, N. Utkina, H. Y. Wang, K. Kato, M. Sadilek, S. I. Hakomori. Interaction of N-linked glycans, having multivalent GlcNAc termini, with GM3 ganglioside. Glycoconj. J. 2006, 23, 639–649.

[14] N. Kawashima, S. J. Yoon, K. Itoh, K. Nakayama. Tyrosine kinase activity of epidermal growth factor receptor is regulated by GM3 binding through carbohydrate to carbohydrate interactions. J. Biol. Chem. 2009, 284, 6147–6155.

[15] I. van der Haar Àvila, B. Windhouwer, S. J. van Vliet. Current state-of-the-art on ganglioside-mediated immune modulation in the tumor microenvironment. Cancer Metastasis Rev. 2023, 42, 941–958.

[16] G. Nicoll, T. Avril, K. Lock, K. Furukawa, N. Bovin, P. R. Crocker. Ganglioside GD3 expression on target cells can modulate NK cell cytotoxicity via siglec-7-dependent and -independent mechanisms. Eur. J. Immunol. 2003, 33, 1642–1628.

[17] M. Zuverink, J. T. Barbieri. Protein Toxins That Utilize Gangliosides as Host Receptors. Prog. Mol. Biol. Transl. Sci. 2018, 156, 325–354.

[18] Y. Suzuki. Gangliosides as influenza virus receptors. Variation of influenza viruses and their recognition of the receptor sialo-sugar chains. Prog. Lipid Res. 1994, 33, 429–457.

[19] Z. Li, Y. Lang, L. Liu, M. I. Bunyatov, A. I. Sarmiento, R. J. de Groot, G. J. Boons. Synthetic O-acetylated sialosides facilitate functional receptor identification for human respiratory viruses. Nat. Chem. 2021, 13, 496–503.

[20] L. Nguyen, K. A. McCord, D. T. Bui, K. M. Bouwman, E. N. Kitova, M. Elaish, D. Kumawat, G. C. Daskhan, I. Tomris, L. Han, P. Chopra, T. J. Yang, S. D. Willows, A. L. Mason, L. K. Mahal, T. L. Lowary, L. J. West, S. D. Hsu, T. Hobman, S. M. Tompkins, G. J. Boons, R. P. de Vries, M. S. Macauley, J. S. Klassen. Sialic acid-containing glycolipids mediate binding and viral entry of SARS-CoV-2. Nat. Chem. Biol. 2022, 18, 81–90.

[21] R. De Maria, L. Lenti, F. Malisan, F. d’Agostino, B. Tomassini, A. Zeuner, M. R. Rippo, R. Testi. Requirement for GD3 ganglioside in CD95- and ceramide-induced apoptosis. Science 1997, 277, 1652–1655.

[22] F. Malisan, L. Franchi, B. Tomassini, N. Ventura, I. Condò, M. R. Rippo, A. Rufini, L. Liberati, C. Nachtigall, B. Kniep, R. Testi. Acetylation suppresses the proapoptotic activity of GD3 ganglioside. J. Exp. Med. 2002, 196, 1535–1541.

[23] S. Cavdarli, P. Delannoy, S. Groux-Degroote. O-acetylated gangliosides as targets for cancer immunotherapy. Cells 2020, 9, 741.

[24] H. Y. Chen, A. K. Challa, A. Varki. 9-O-acetylation of exogenously added ganglioside GD3. The GD3 molecule induces its own O-acetylation machinery. J. Biol. Chem. 2006, 281, 7825–7833.

[25] A. M. Baumann, M. J. Bakkers, F. F. Buettner, M. Hartmann, M. Grove, M. A. Langereis, R. J. de Groot, M. Muhlenhoff. 9-O-Acetylation of sialic acids is catalysed by CASD1 via a covalent acetyl-enzyme intermediate. Nat. Commun. 2015, 6, 7673.

[26] M. D. Vaughan, K. Johnson, S. DeFrees, X. Tang, R. A. Warren, S. G. Withers. Glycosynthase-mediated synthesis of glycosphingolipids. J. Am. Chem. Soc. 2006, 128, 6300–6301.

[27] S. M. Hancock, J. R. Rich, M. E. Caines, N. C. Strynadka, S. G. Withers. Designer enzymes for glycosphingolipid synthesis by directed evolution. Nat. Chem. Biol. 2009, 5, 508–514.

[28] K. T. Huang, N. Winssinger. IPy2BF4-mediated glycosylation and glycosyl fluoride formation. Eur. J. Org. Chem. 2007, 2007, 1887–1890.

[29] J. C. Lopez, P. Bernal-Albert, C. Uriel, S. Valverde, A. M. Gomez. IPy2BF4/HF-pyridine: a new combination of reagents for the transformation of partially unprotected thioglycosides and n-pentenyl glycosides to glycosyl fluorides. J. Org. Chem. 2007, 72, 10268–10271.

[30] M. S. Maginnis. Virus–receptor interactions: The key to cellular invasion. J. Mol. Biol. 2018, 430, 2590–2611.

[31] A. Regina Todeschini, S. I. Hakomori. Functional role of glycosphingolipids and gangliosides in control of cell adhesion, motility, and growth, through glycosynaptic microdomains. Biochim. Biophys. Acta 2008, 1780, 421–433.

[32] F. Broszeit, R. J. van Beek, L. Unione, T. M. Bestebroer, D. Chapla, J. Y. Yang, K. W. Moremen, S. Herfst, R. A. M. Fouchier, R. P. de Vries, G. J. Boons. Glycan remodeled erythrocytes facilitate antigenic characterization of recent A/H3N2 influenza viruses. Nat. Commun. 2021, 12, 5449.

[33] G. M. Vos, K. C. Hooijschuur, Z. Li, J. Fjeldsted, C. Klein, R. P. de Vries, J. Toraño Sastre, G. J. Boons. Sialic acid O-acetylation patterns and glycosidic linkage type determination by ion mobility-mass spectrometry. Nat. Commun. 2023, 14, 6795.

[34] D. F. Smith, P. A. Prieto. Special considerations for glycolipids and their purification. Curr. Protoc. Mol. Biol. 2001, 22, 17.13.11–17.13.13.

[35] Y. Ji, A. Sasmal, W. Li, L. Oh, S. Srivastava, A. A. Hargett, B. R. Wasik, H. Yu, S. Diaz, B. Choudhury, C. R. Parrish, D. I. Freedberg, L. P. Wang, A. Varki, X. Chen. Reversible O-acetyl migration within the sialic acid side chain and its influence on protein recognition. ACS Chem. Biol. 2021, 16, 1951–1960.

[36] W. Li, R. J. G. Hulswit, I. Widjaja, V. S. Raj, R. McBride, W. Peng, W. Widagdo, M. A. Tortorici, B. van Dieren, Y. Lang, J. W. M. van Lent, J. C. Paulson, C. A. M. de Haan, R. J. de Groot, F. J. M. van Kuppeveld, B. L. Haagmans, B. J. Bosch. Identification of sialic acid-binding function for the Middle East respiratory syndrome coronavirus spike glycoprotein. Proc. Natl. Acad. Sci. U. S. A. 2017, 114, E8508–E8517.

[37] I. Tomris, A. L. M. Kimpel, R. Liang, R. van der Woude, G. P. H. Boons, Z. Li, R. P. de Vries. The HCoV-HKU1 N-terminal domain binds a wide range of 9-O-acetylated sialic acids presented on different glycan cores. ACS Infect. Dis. 2024, 10, 3880–3890.

[38] R. J. G. Hulswit, Y. Lang, M. J. G. Bakkers, W. Li, Z. Li, A. Schouten, B. Ophorst, F. J. M. van Kuppeveld, G. J. Boons, B. J. Bosch, E. G. Huizinga, R. J. de Groot. Human coronaviruses OC43 and HKU1 bind to 9-O-acetylated sialic acids via a conserved receptor-binding site in spike protein domain A. Proc. Natl. Acad. Sci. U. S. A. 2019, 116, 2681–2690.

[39] H. Ishida, Y. Ohta, Y. Tsukada, M. Kiso, A. Hasegawa. A synthetic approach to polysialogangliosides containing alpha-sialyl-(2-->8)-sialic acid: total synthesis of ganglioside GD3. Carbohydr. Res. 1993, 246, 75–88.

[40] Y. Ito, J. C. Paulson. A novel strategy for synthesis of ganglioside GM3 using an enzymically produced sialoside glycosyl donor. J. Am. Chem. Soc. 1993, 115, 1603–1605.

[41] A. Hasegawa, M. Kato, T. Ando, H. Ishida, M. Kiso. Synthesis of sialyl Lewis X ganglioside analogues containing modified L-fucose residues. Carbohydr. Res. 1995, 274, 165–181.

[42] J. C. Castro-Palomino, B. Simon, O. Speer, M. Leist, R. R. Schmidt. Synthesis of ganglioside GD3 and its comparison with bovine GD3 with regard to oligodendrocyte apoptosis mitochondrial damage. Chem. Eur. J. 2001, 7, 2178–2184.

[43] J. R. Rich, D. R. Bundle. S-linked ganglioside analogues for use in conjugate vaccines. Org. Lett. 2004, 6, 897–900.

[44] T. Kondo, T. Tomoo, H. Abe, M. Isobe, T. Goto. Total synthesis of GD3, a ganglioside. Chem. Lett. 2006, 25, 337–338.

[45] H. Yu, J. Cheng, L. Ding, Z. Khedri, Y. Chen, S. Chin, K. Lau, V. K. Tiwari, X. Chen. Chemoenzymatic synthesis of GD3 oligosaccharides and other disialyl glycans containing natural and non-natural sialic acids. J. Am. Chem. Soc. 2009, 131, 18467–18477.

[46] H. Y. Chuang, C. T. Ren, C. A. Chao, C. Y. Wu, S. S. Shivatare, T. J. Cheng, C. Y. Wu, C. H. Wong. Synthesis and vaccine evaluation of the tumor-associated carbohydrate antigen RM2 from prostate cancer. J. Am. Chem. Soc. 2013, 135, 11140–11150.

[47] S. Takayama, P. O. Livingston, C. H. Wong. Synthesis of the melanoma-associated ganglioside 9-O-acetyl GD3 through regioselective enzymatic acetylation of GD3 using subtilisin. Tetrahedr. Lett. 1996, 37, 9271–9274.

[48] M. A. Langereis, M. J. Bakkers, L. Deng, V. Padler-Karavani, S. J. Vervoort, R. J. Hulswit, A. L. van Vliet, G. J. Gerwig, S. A. de Poot, W. Boot, A. M. van Ederen, B. A. Heesters, C. M. van der Loos, F. J. van Kuppeveld, H. Yu, E. G. Huizinga, X. Chen, A. Varki, J. P. Kamerling, R. J. de Groot. Complexity and diversity of the mammalian sialome revealed by nidovirus virolectins. Cell Rep. 2015, 11, 1966–1978.

[49] C. L. Wardzala, A. M. Wood, D. M. Belnap, J. R. Kramer. Mucins inhibit coronavirus infection in a glycan-dependent manner. ACS Cent. Sci. 2022, 8, 351–360.

[50] M. F. Pronker, R. Creutznacher, I. Drulyte, R. J. G. Hulswit, Z. Li, F. J. M. van Kuppeveld, J. Snijder, Y. Lang, B. J. Bosch, G. J. Boons, M. Frank, R. J. de Groot, D. L. Hurdiss. Sialoglycan binding triggers spike opening in a human coronavirus. Nature 2023, 624, 201–206.

[51] N. Saunders, I. Fernandez, C. Planchais, V. Michel, M. M. Rajah, E. Baquero Salazar, J. Postal, F. Porrot, F. Guivel-Benhassine, C. Blanc, G. Chauveau-Le Friec, A. Martin, L. Grzelak, R. M. Oktavia, A. Meola, O. Ahouzi, H. Hoover-Watson, M. Prot, D. Delaune, M. Cornelissen, M. Deijs, V. Meriaux, H. Mouquet, E. Simon-Loriere, L. van der Hoek, P. Lafaye, F. Rey, J. Buchrieser, O. Schwartz. TMPRSS2 is a functional receptor for human coronavirus HKU1. Nature 2023, 624, 207–214.

[52] I. Fernandez, N. Saunders, S. Duquerroy, W. H. Bolland, A. Arbabian, E. Baquero, C. Blanc, P. Lafaye, A. Haouz, J. Buchrieser, O. Schwartz, F. A. Rey. Structural basis of TMPRSS2 zymogen activation and recognition by the HKU1 seasonal coronavirus. Cell 2024, 187, 4246–4260 e4216.

