## Supplementary material for "Chemoenzymatically Synthesized *O*-Acetylated GD3 Gangliosides to Examine Viral Receptor Specificities in a Cellular Context": SI

**Table of Contents** page

1. Supplementary Figures and Table S2
***Figure S1.*** α-Configuration confirmed by $^{1}H$, $^{19}F$, and HSQC NMR of disaccharide **8** S2
***Figure S2.*** 2D NMR analyses for fully deprotected tetrasaccharide glycosyl fluoride **6** S3
***Figure S3.*** HSQC and HMBC spectra of tetrasaccharides **4**, **16**, and **17** S4-5
***Figure S4.*** HSQC spectra of tetrasaccharides **1**, **2**, and **3** S6-7
***Figure S5.*** IM-MS analysis of glycans released for gangliosides S8-9
***Table S1.*** *O*-acetyl form analysis for sialic acid moieties on GD3 S10
***Figure S6.*** Hemagglutination assay with (*O*-acetylated) GD3 gangliosides S11
2. Materials and General Methods S12
1.1 Chemicals S12
1.2 General Analytical Methods S12
1.3 NMR Assignments S13
1.4 Expression and Purification of Recombinant Hemagglutinin-Esterase S13
1.5 Expression and Purification of EGCaseII E351S, D314Y S14
1.6 Glycosphingolipid Extraction and Release S14
1.7 Glycan Analysis with LC-IM-MS S15
1.8 Expression and Purification of Hku1-S1$^{A}$-FC and pA-LS S16
1.9 *O*-acetylated GD3 Gangliosides Modified Erythrocytes S16
1.10 Hemagglutination Assay S16
1.11 Miscellaneous Information S17
3. Synthetic Methods and Compound Characterization S17
***Scheme S1***. Monosaccharide building block **S6** S17
***Scheme S2.*** Monosaccharide building block **S10** S21
***Scheme S3.*** Monosaccharide building blocks **8** and **9** S23
***Scheme S4.*** Synthetic route of intermediate tetrasaccharides fluoride **11** S32
***Scheme S5.*** Synthetic route of intermediate tetrasaccharides fluoride **6** S33
4. References S56
5. NMR spectra S57

### 1. Supplementary Figures and Table

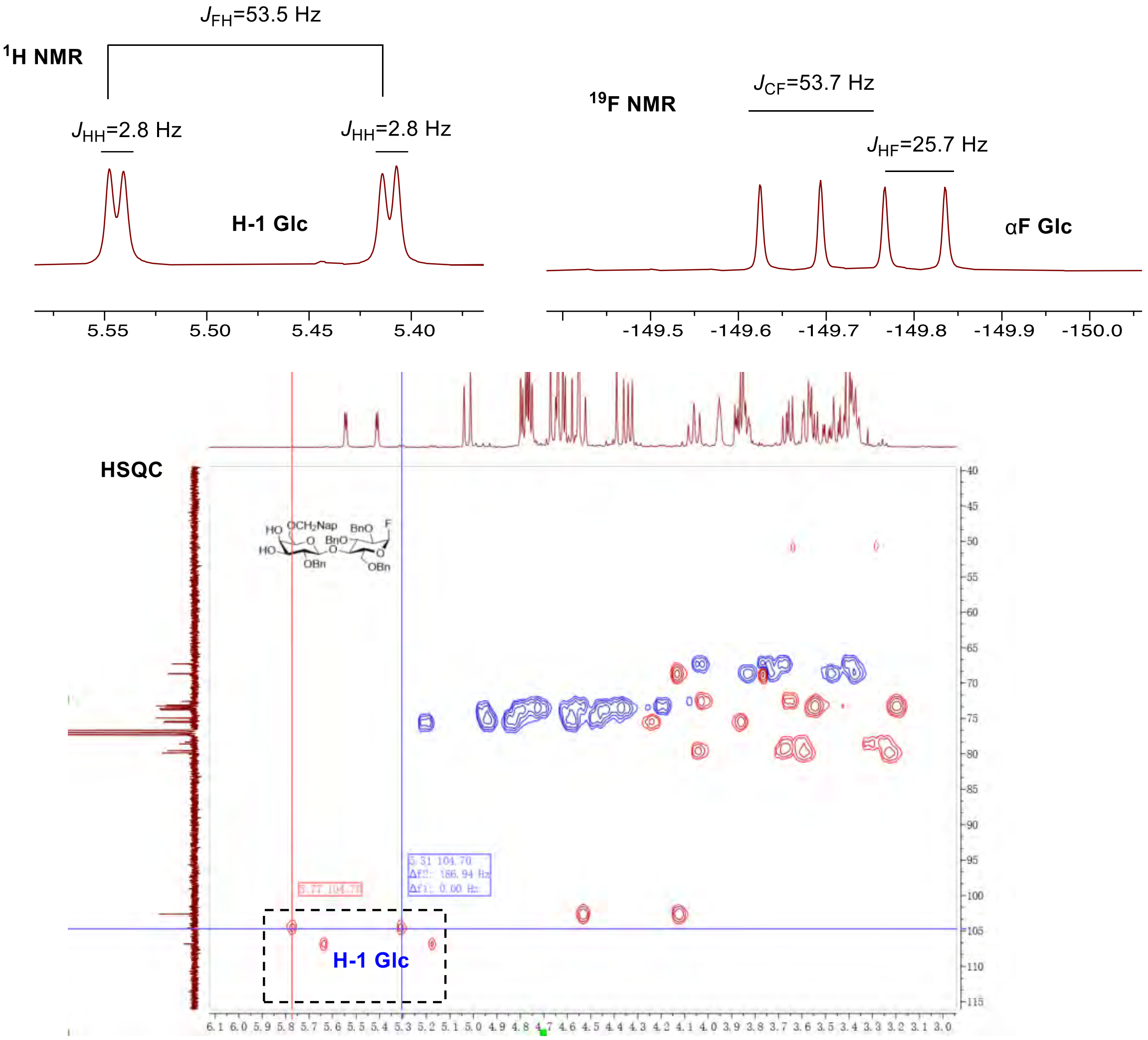


***Figure S1.*** α-Configuration confirmed by $^{1}$H, $^{19}$F, and HSQC (no decoupling) NMR of disaccharide **8**. The H-1$_{Glc}$ signal appears as doublet of doublets in $^{1}$H-NMR with a characteristic chemical shift and coupling constants (*J* values) consistent with the formation of an anomeric α-fluoride. $^{19}$F NMR spectrum of the product matches with the reported chemical shifts and coupling constants of anomeric α-fluorides. In the HSQC spectrum, the coupling constant $^{1}J_{C1, H1}$ is larger than 170 Hz, confirming the formation of desired α-glycosyl fluoride.

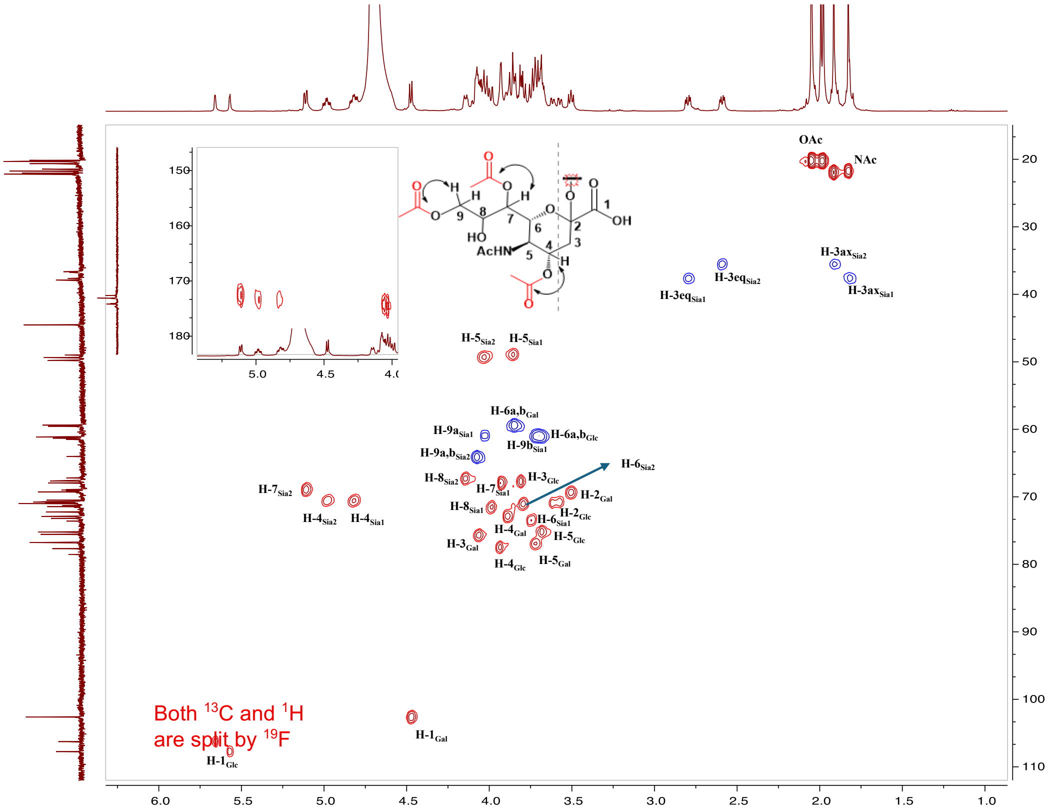


***Figure S2.*** 2D NMR analyses of fully deprotected tetrasaccharide glycosyl fluoride **6**. HSQC spectrum of compound **6** shows the preservation of the anomeric fluoride. The $^{1}$H and $^{13}$C peak splitting by $^{19}$F is present. The zoomed-in partial spectrum is the HMBC signals between protons of C4, C5, C7, and C9 positions and the carbonyl carbons of acetyl esters.

**A.** 4,7,9-tri-OAc-GD3βSph (**4**)

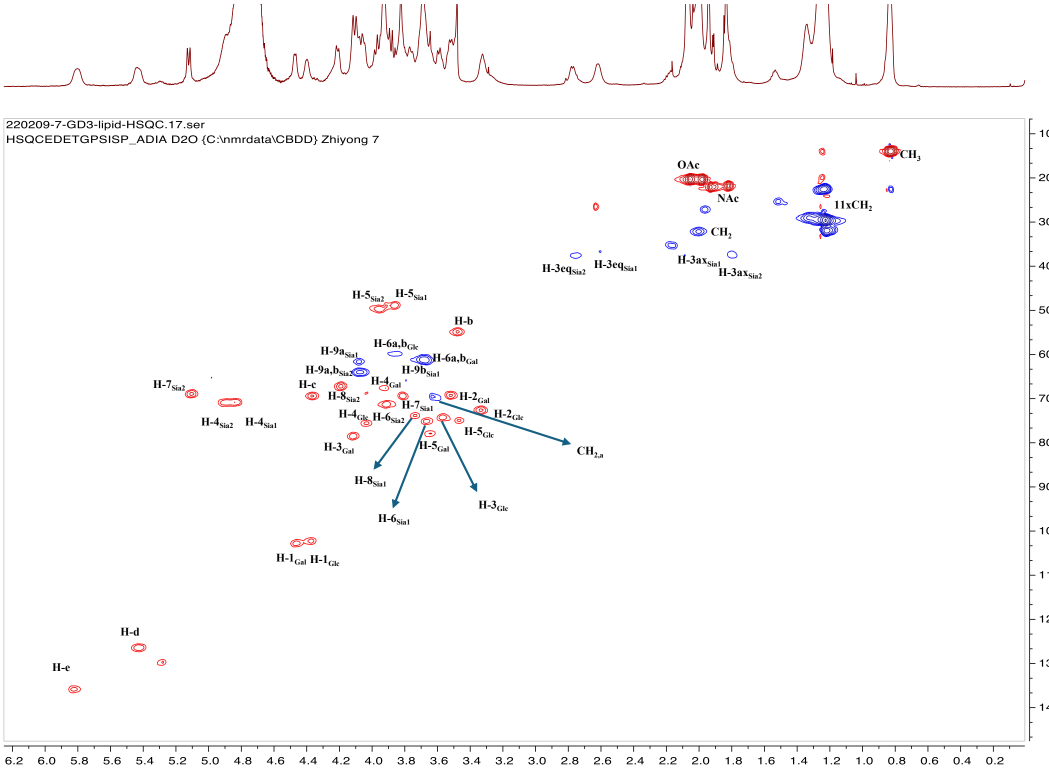


**B.** 7,9-di-OAc-GD3βSph (**16**)

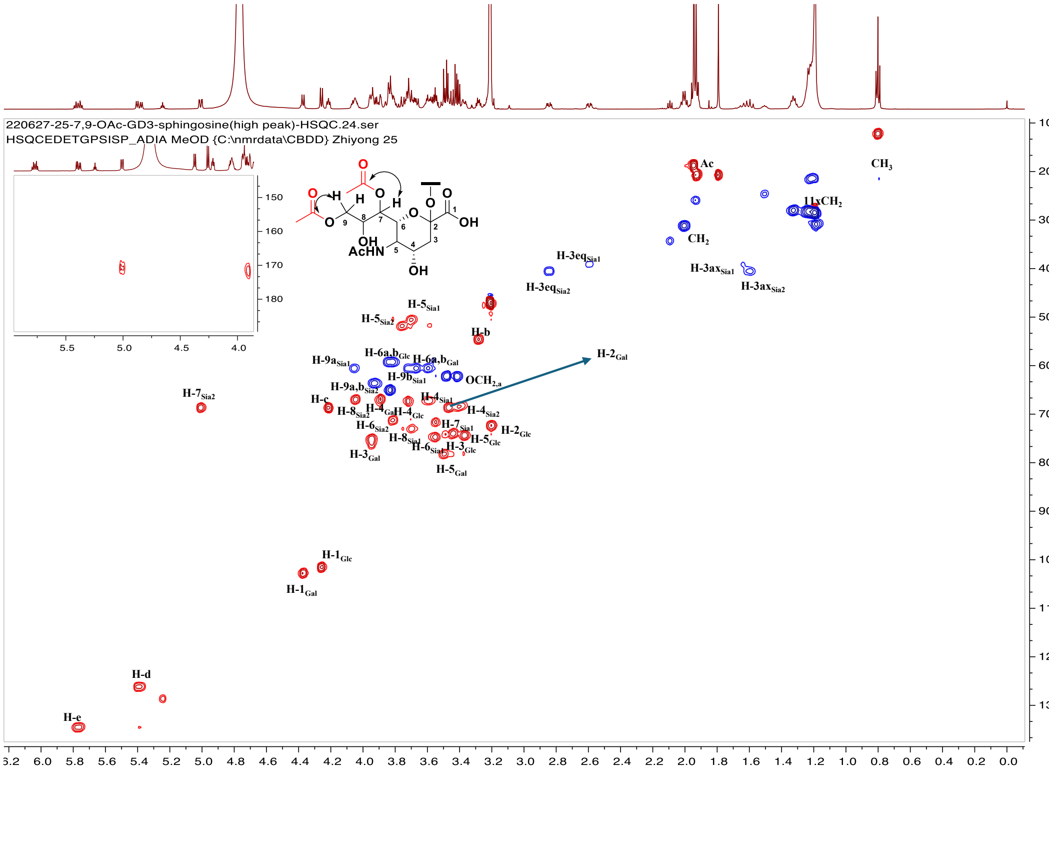

**C.** 7-OAc-GD3βSph (**17**)

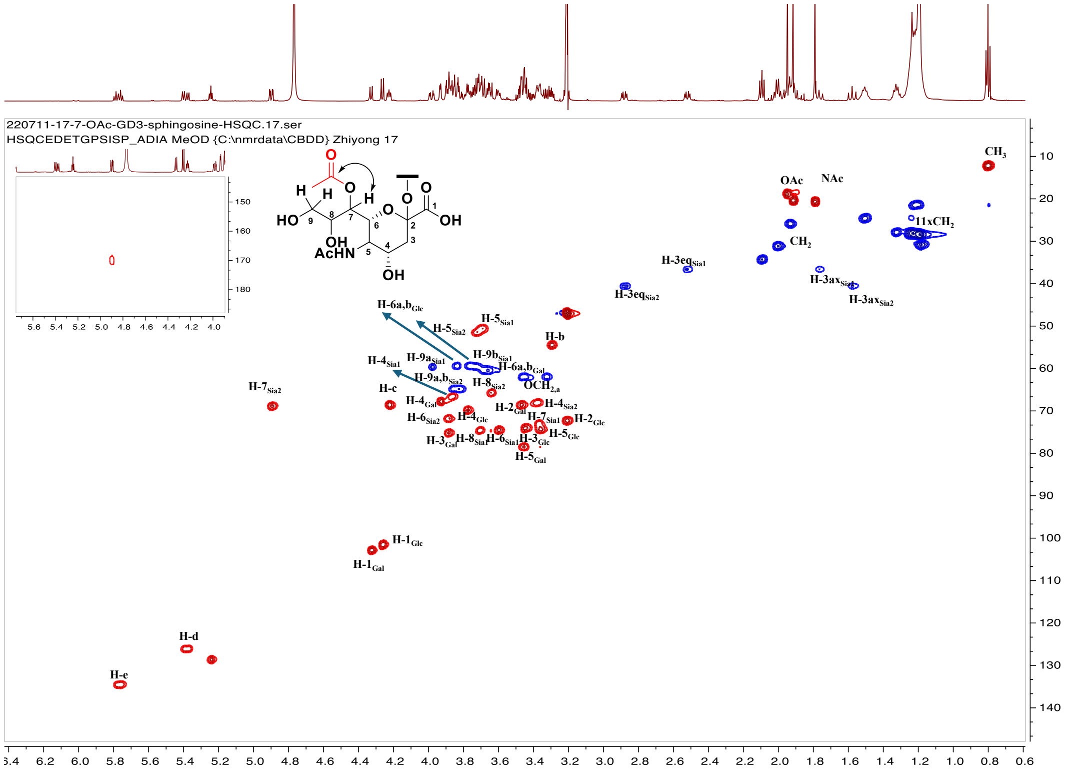


***Figure S3.*** HSQC and HMBC spectra of tetrasaccharides (A) **4**; (B) **16**; and (C) **17**. All acetyl esters remained at the expected positions after EGC glycosynthase-mediated sphingosine attachment and HE-mediated *O*-acetyl editing. See Experimental Section for signal assignment.

**A.** 7,9-di-OAc-GD3 (**1**)

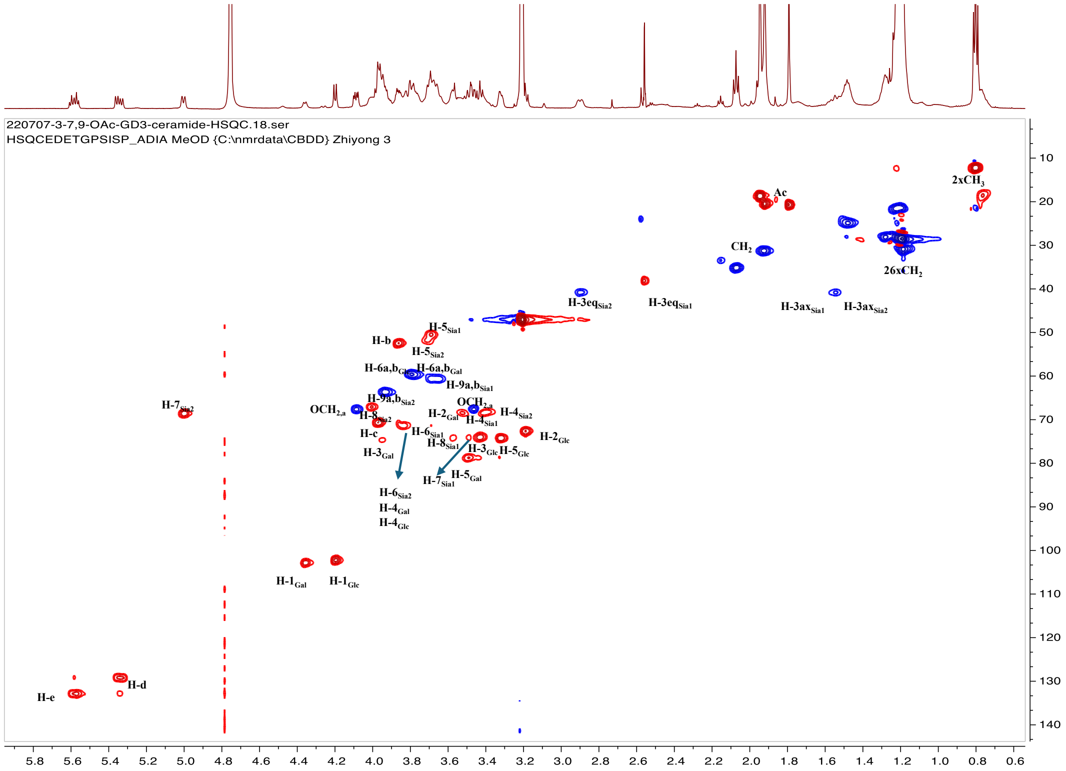


**B.** 7-OAc-GD3 (**2**)

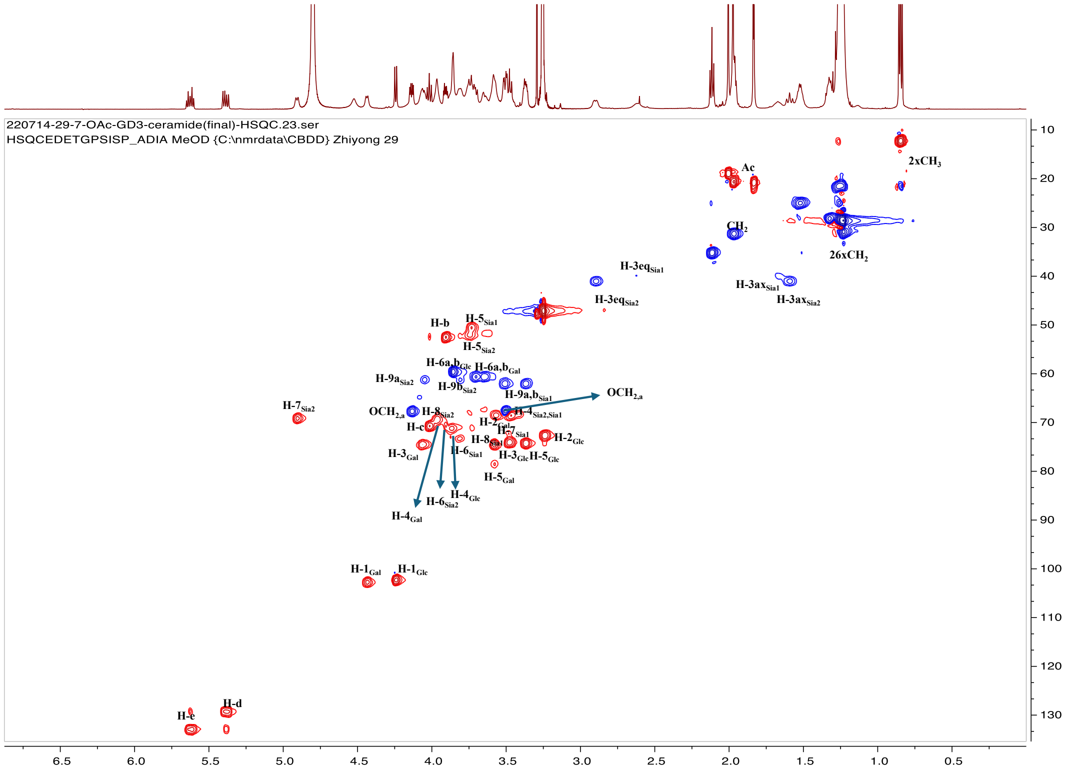

**C.** 9-OAc-GD3 (**3**)

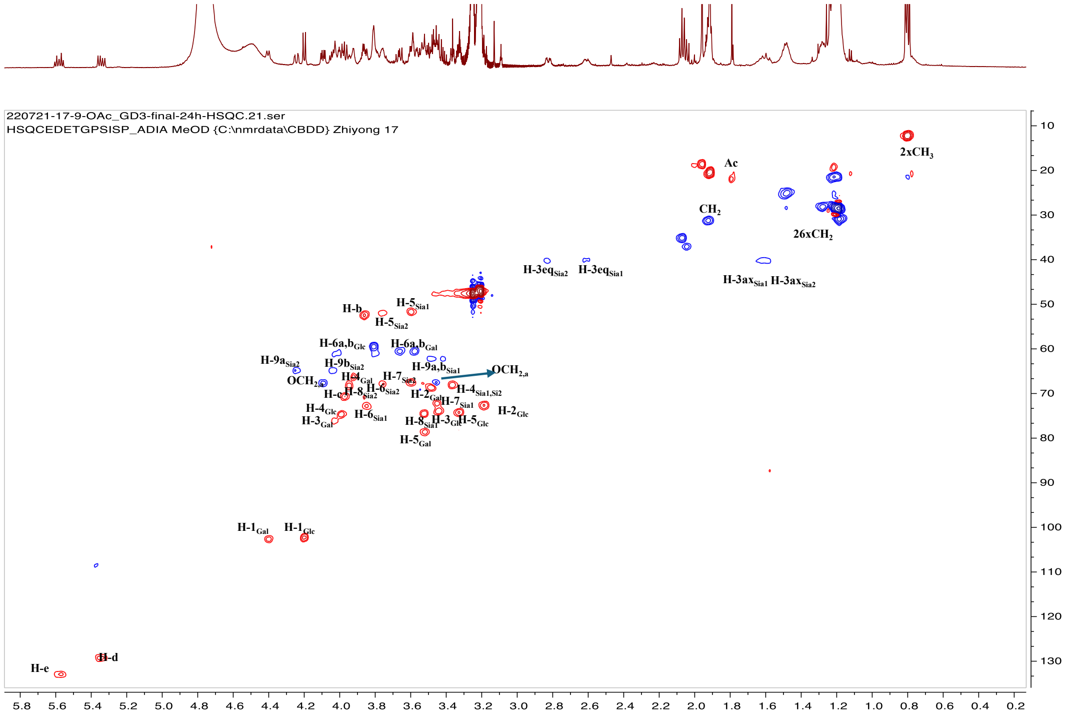

***Figure S4.*** HSQC spectra for tetrasaccharides **1**, **2**, and **3**. Preservation of *O*-acetyl moieties in final products (A) **1**; (B) **2**; and (C) **3** after acylation with stearic acid NHS ester. See Experimental Section for signal assignment.

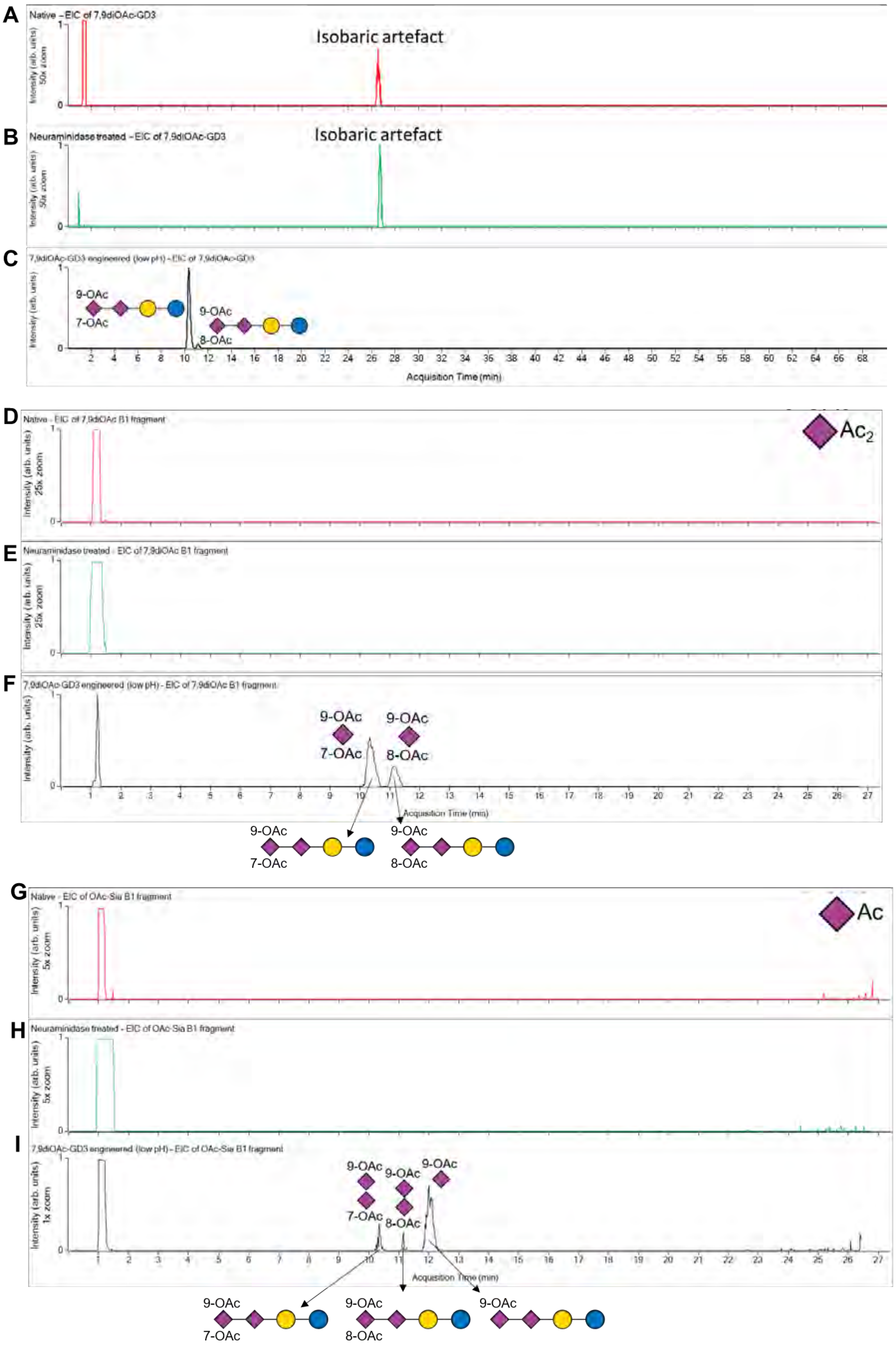
A
Native – EIC of 7,9diOAc-GD3
Isobaric artefact
Intensity (arb. units) 50x zoom
B
Neuraminidase treated – EIC of 7,9diOAc-GD3
Isobaric artefact
Intensity (arb. units) 50x zoom
C
7,9diOAc-GD3 engineered (low pH) – EIC of 7,9diOAc-GD3
9-OAc
7-OAc
9-OAc
8-OAc
Intensity (arb. units)
Acquisition Time (min)
D
Native - EIC of 7,9diOAc B1 fragment
Ac2
Intensity (arb. units) 25x zoom
E
Neuraminidase treated - EIC of 7,9diOAc B1 fragment
Intensity (arb. units) 25x zoom
F
7,9diOAc-GD3 engineered (low pH) - EIC of 7,9diOAc B1 fragment
9-OAc
7-OAc
9-OAc
8-OAc
Intensity (arb. units)
Acquisition Time (min)
9-OAc
7-OAc
9-OAc
8-OAc
G
Native - EIC of OAc-Sia B1 fragment
Ac
Intensity (arb. units) 5x zoom
H
Neuraminidase treated - EIC of OAc-Sia B1 fragment
Intensity (arb. units) 5x zoom
I
7,9diOAc-GD3 engineered (low pH) - EIC of OAc-Sia B1 fragment
9-OAc
7-OAc
9-OAc
8-OAc
9-OAc
Intensity (arb. units) 1x zoom
Acquisition Time (min)
9-OAc
7-OAc
9-OAc
8-OAc
9-OAc

***Figure S5.*** IM-MS analysis of glycans released from gangliosides. Extracted ion chromatograms showing absence of *O*-acetylated gangliosides in native and neuraminidase-treated samples. (A-C) The molecular ion of di-*O*-acetylated GD3 (with procainamide label) was extracted for (A) native; (B) neuraminidase-treated; and (C) 7,9-di-*O*-acetylated GD3-remodeled erythrocyte samples. (D-F) $B_1$ fragment of di-*O*-acetylated sialic acid was extracted for (D) native; (E) neuraminidase-treated; and (F) 7,9-di-*O*-acetylated GD3-remodeled erythrocyte samples. (G-I) $B_1$ fragment of mono-*O*-acetylated sialic acid was extracted for (G) native; (H) neuraminidase-treated; and (I) 7,9-di-*O*-acetylated GD3-remodeled erythrocyte samples. The mono-acetylated $B_1$ fragments arose from the loss of one Ac during in-source fragmentation (left and middle peaks), as well as a minor, hydrolyzed and subsequently Ac migrated product (right peak) prior to MS analysis.

***Table S1.*** Acetyl ester analysis of sialic acid moieties on GD3 from remodeled erythrocytes. Quantification of *O*-acetyl sialic acid forms on GD3 extracted from remodeled erythrocytes was based on the area-under-curve values of the extracted ion chromatogram.

| **Structure** | **7-OAc-GD3 added** | **9-OAc-GD3 added** | **7,9-di-OAc-GD3 added** |
|---|---:|---:|---:|
| 7-OAc-GD3 | 89.4% | 1.8% | n.d. |
| 9-OAc-GD3 | 10.6% | 98.2% | 3.0% |
| 7,9-di-OAc-GD3 | - | - | 94.2% |
| 7,8-di-OAc-GD3 | - | - | 2.8% |

n.d. indicates not detected

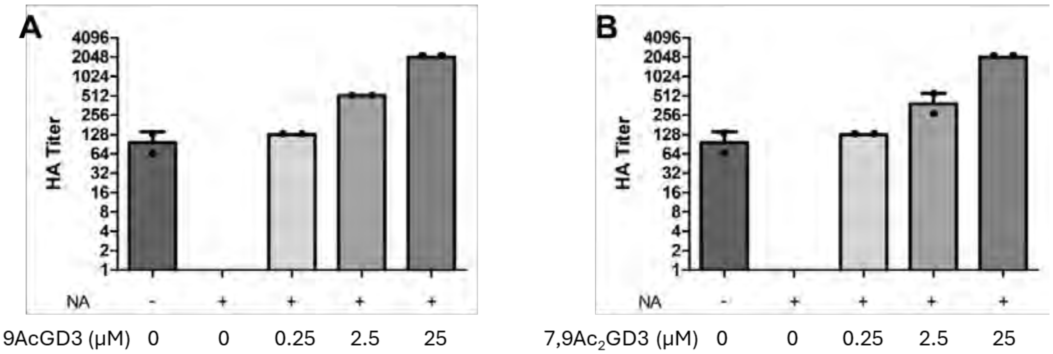


***Figure S6.*** Hemagglutination assay with varying concentrations of (*O*-acetylated) GD3 gangliosides. (A) HKU1 $S1^{A}$-domain functionalized pA-LS nanoparticle with 9-OAc-GD3-modified human erythrocytes, which was pre-treated with neuraminidase to remove endogenous sialic acids; (B) HKU1 $S1^{A}$-domain functionalized pA-LS nanoparticle with 7,9-di-OAc-GD3-modified human erythrocytes, which was pre-treated with neuraminidase to remove endogenous sialic acids.

#### 2. Material and General Methods

##### 1.1 Chemicals

Unless otherwise stated, all chemical reagents were acquired from commercial sources and utilized without additional purification. Acetonitrile, dichloromethane, toluene, tetrahydrofuran and *N*,*N*-dimethylformamide used for synthesis were anhydrous grade from an MB SPS 5 solvent purification system. Other organic solvents used for reactions were obtained from Biosolve Chemie. Technical grade organic solvents for work-up procedures were supplied from VWR Chemicals. Deuterated solvents for NMR experiments were purchased from Cambridge Isotope Laboratories. Molecular sieves (MS) were flame dried in a round bottom flask with a Bunsen burner and were placed under high vacuum to allow the activated molecular sieves to cool prior to use.

##### 1.2 General Analytical Methods

TLC analysis was performed using precoated silica gel 60 F-254 plates (Merck). The plates were either visualized with UV lamp or directly stained with ceric ammonium molybdate (5 g $Ce(SO4)_2$, 25 g $(NH_4)_6Mo_7O_{24}{\cdot}4H_2O$, 50 mL conc. $H_2SO_4$, 450 mL $H_2O$). Column chromatography was performed using flash silica gel (60 Å, from Silicycle, Canada). Water soluble compounds were purified using P-2 biogel (Biorad) and collected with BioFrac fraction collector (Biorad). C18 column were purchased from Dr. Maisch GmbH (ReproSil-Pur 120 C18-AQ, 10 µm) for reverse phase HPLC chromatography. $^{1}H$ and $^{13}C$ NMR spectroscopy was conducted on an Agilent 400-MR, VARIAN INOVA-500 or Bruker 600 UltraShield. Chemical shifts are reported in parts per million (ppm) relative to $CDCl_3$ as the internal standard. NMR data are presented as follows: Chemical shift, multiplicity (s = singlet, d = doublet, t = triplet, dd = doublet of doublets, ddd = doublet of doublet of doublets, td = triplet of doublets, m = multiplet); coupling constants are reported in Hertz (Hz). All NMR signals were assigned on the basis of $^{1}H$ NMR, COSY, HSQC, HMBC and TOCSY experiments. MS-based reaction monitoring was performed on Shimadzu Kratos Axima-CFR MALDI-ToF or Brucker micrOTOF-Q II ESI mass spectrometer. LC-MS traces were collected on Shimadzu LC-ESI-IT-TOF. HRMS data was collected on Agilent 6560 ion mobility Q-ToF MS.

#### 1.3 NMR Assignments

Atom numbering and residue labels for NMR assignment.

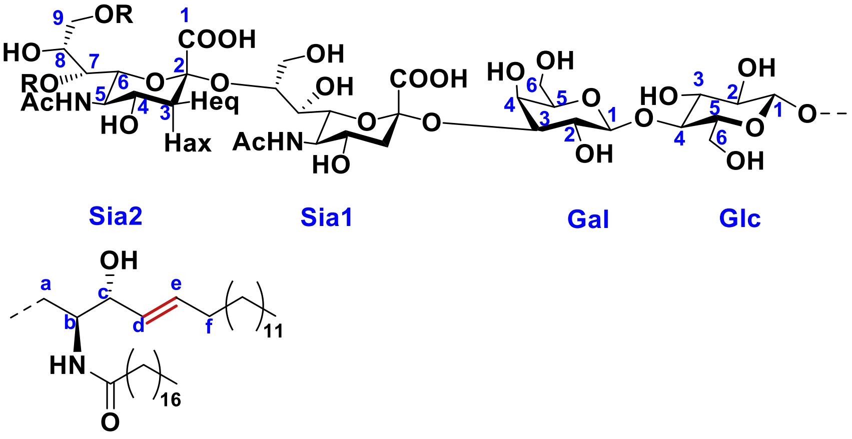


For NMR assignments, the protons on the carbohydrate portion are denoted H-n (n = number of position) with a subscript specifying the sugar unit. For example, H-$1_{Glc}$ stands for the anomeric proton of glucose. For the primary alcohol positions, including C6 of Glc and Gal and C9 of Sia, a and b are used to distinguish the geminal protons. For example, H-$9a_{Sia1}$ and H-$9b_{Sia1}$. In particular, the C3 geminal protons of both Sia residues are denoted H-3eq and H-3ax, for equatorial and axial protons, respectively. For coupling constant ($J$ values) between protons, they are specified with a subscript of sugar unit followed by the two positions. For example, $J_{Glc1,2}$ stands for the coupling constant between protons at C1 and C2. For monosaccharides, the sugar units are not specified. The carbon numbering follows the same rules. For the sphingosine portion, the protons are denoted H-x (x = positions a – f). For the coupling constant, $J_{vic}$ denotes the value between two vicinal protons, $J_{gem}$ between two geminal protons, and $J_{trans}$ between two protons across a trans double bond. The carbon numbering follows the same rules. The chemical shifts of thiotolyl, benzyl, and naphthyl groups, and aliphatic $CH_2$ and $CH_3$ of the sphingosine and the fatty acid are not individually assigned.

#### 1.4 Expression and Purification of Recombinant Hemagglutinin-Esterase (HE)

All HEs used in this study were by a reported procedure.[1] In brief, codon-optimized sequences encoding the ectodomains of HEs fused to the Fc domain of human IgG1 *via* a thrombin-cleavable linker were cloned into pCD5 or pCAGGS vectors. HEK293T cells (~2 × $10^7$) were transfected with 20 μg plasmid DNA complexed with 2 μg polyethyleneimine (Polysciences). Sixteen hours post-transfection, the medium was replaced with 293 SFM II (Invitrogen) supplemented with 44.1 mM sodium bicarbonate, 11.1 mM glucose, 3.0 g/L Primatone RL-UF, 100 IU/mL penicillin, 100 μg/mL streptomycin, 1% GlctaMAX (Gibco), and 1.5% DMSO. Supernatants were collected 6–7 days later and clarified by sequential centrifugation (1200 rpm for 5 min, followed by 4000 rpm for 10 min at 4 °C). The proteins were captured by overnight incubation with Protein A-Sepharose (150 μL per 50 mL supernatant) at 4 °C, and purified using Poly-Prep columns (Bio-Rad). Elution was performed with 0.1 M citric acid, and eluates were immediately neutralized with one-third volume of 1 M Tris-HCl (pH 8.8). Purified HE-Fc proteins were dialyzed against PBS and stored at −80 °C until use.

**1.5 Expression and Purification of EGCaseII E351S, D314Y[2]**

The gene encoding endo-glycoceramidase II (EGCII) from *Rhodococcus* sp. strain M-777, optimized for *Escherichia coli* codon usage and lacking the N-terminal 30-residue secretion signal, was synthesized and subcloned into the pET28a vector via *NdeI* and *XhoI* restriction sites. Site-directed mutagenesis was performed to introduce E351S and D314Y substitutions, resulting in a double mutant construct (EGCII E351S, D314Y) that was transformed into *E. coli* BL21(Tuner) cells. Cells were cultured in Lysogeny Broth (LB) medium containing 50 μg/mL kanamycin at 37 °C to stationary phase. Protein expression was induced by adding 0.1 mM IPTG following a temperature shift to room temperature. The $His_6$-tagged EGCII E351S, D314Y (residues 31−490) was purified by Ni(II)-affinity chromatography to >95% purity as determined by SDS-PAGE. The protein was dialyzed against 25 mM sodium acetate buffer (pH 5.0) and stored at −80 °C until use.

**1.6 Glycosphingolipid Extraction and Release**

To erythrocytes in an eppendorf vial was added 750 μL of 2:1 $CHCl_3$/MeOH followed by sonication on ice for 30 min. 750 μL 1:1 MeOH/$H_2O$ was added to the mixture, which was vortexed and centrifuged at 15 xg, and the supernatant was collected. The pellet was extracted again with 4:8:3 $CHCl_3$/MeOH/$H_2O$ and with 2:1 $CHCl_3$/MeOH. The supernatants were combined, dried under a flow of nitrogen, and resuspended in 5% MeOH in $H_2O$. The lipids were isolated with C18 SPE (Avantor Bakerbond, Amsterdam): a cartridge was washed with MeOH, $CHCl_3$, MeOH, and 5% MeOH in $H_2O$, followed by sample loading and washing with 5% MeOH in $H_2O$. Lipids were eluted consecutively with MeOH, 4:8:3 $CHCl_3$/MeOH/$H_2O$, and 2:1 $CHCl_3$/MeOH and the elution was dried under a flow of nitrogen. Glycans were released from the extracted lipid with endoglycoceramidase I (expressed in house): lipids were dissolved in 45 μL of 50 mM sodium acetate buffer (pH 5.2) with 0.1% Triton X-100, 5 μL EGCase I was added, and the mixture was incubated at 37 °C overnight. The mixture was diluted with 5% MeOH in $H_2O$ and loaded on a C18 SPE (washed with MeOH, $CHCl_3$, MeOH, 5% MeOH in $H_2O$). The glycans were released with 5% MeOH in $H_2O$, while the lipids remained on the column. Glycans were dissolved in 120 μL $H_2O$, and 40 μL procainamide labeling mixture (60 mg/mL of procainamide-HCl and $NaBH_3CN$) and 20 μL acetic acid were added, followed by incubation at RT for 4 h. Labeled glycans were cleaned up with C18 SPE by washing the cartridge with ACN and $H_2O$, loading the glycans in $H_2O$, and eluting the glycans in 10% ACN in $H_2O$. The ACN was evaporated from the elute, which was loaded on a porous graphitized carbon SPE that was washed with ACN and $H_2O$. The cartridge was washed with $H_2O$ and glycans were eluted with 1 mL 50% ACN in $H_2O$, dried under nitrogen, and resuspended in 70% acetonitrile.

##### 1.7 Glycan Analysis with LC-IM-MS

Glycans were analyzed using an Agilent Technologies 1290 LC system coupled to an Agilent Technologies 6560B drift tube ion mobility-quadrupole-time of flight mass spectrometer via a dual-spray AJS electrospray source. Glycans were separated with HILIC using a ZIC-HILIC (150 x 4.6 mm, 3.5 μm) column outfitted with a ZIC-HILIC (20x2.1 mm) guard column (Merck, Darmstadt, Germany). The gradient used 0.1% aqueous formic acid as A and acetonitrile as B and was programmed as follows: 70% B at 0 min, 70% B at 2 min, 50% at 20 min, and 50% at 22 min, followed by 15 min equilibration at 70% B. The 6560 IM-QTOF was operated with a capillary voltage of 3500 V, nozzle voltage of 1000 V, nebulizer pressure of 40 psi, 8 l/min nitrogen drying gas at 300 °C, and 11 l/min sheath gas at 350 °C. The drift tube used a trap fill time of 3900 μs, a release time of 250 μs, a drift tube entrance voltage of 1400 V, and a 4-bit multiplexed pulse sequence. The transfer capillary was modified with an exit lens that creates a potential difference with the first funnel of the IM, which was set to 600 V to promote glycan fragmentation.[3] Data files were processed with the PNNL preprocessor v4.1 (Pacific Northwest National Laboratory, Richland, WA) with 3 drift bins interpolation and 5-point moving average smoothing and with

Agilent's HRdm v2.0 software. CCS values were calculated from arrival time by single field calibration with the ESI TOF-MS calibration standard with known *m/z* and CCS values.

##### 1.8 Expression and Purification of Hku1-S1$^{A}$-FC and pA-LS

HKU1-S1$^{A}$ was cloned in the pCDNA5 expression vector, with a C-terminal human IgG1 Fc, and a Strep-tag. pA-LS was cloned in the pCD5 vector with a strep-tag at the C-terminus. HKU1-S1$^{A}$-Fc and pA-LS were expressed in HEK293S GnTI(-) cells. The transfection was performed using plasmids and polyethyleneimine I. After 6 h of transfection, the medium was replaced with 293 SFM II expression medium (Gibco) supplemented with 1 % glutaMAX (Gibco), and 1.5% DMSO. 3.0 g/L Primatone (Kerry), 3.6 g/L bicarbonate, 2.0 g/L glucose, 0.4 g/L valproic acid. After 5-6 days, the cell culture supernatants were harvested, and the proteins were purified by using Strep-Tactin sepharose beads (IBA, Germany).

##### 1.9 *O*-acetylated GD3 Gangliosides Modified Erythrocytes

Fresh human blood was centrifuged (400 rcf, 10 min) and the supernatant was removed. Next, the erythrocytes were washed 3-times with PBS and stored in 50% PBS. 250 µL of 50% erythrocytes were treated with α2-3,6,8,9 Neuraminidase A (24 U) (NEB, P0722L) in 750 µL reaction solution at 37 °C overnight. The reaction solution contains 2 µM $MnCl_2$, 12 µg/mL BSA, 6 U alkaline phosphatase (Thermo scientific, EF0651), and 7 µM Amikacin in PBS. Next, the erythrocytes were washed 3-times with 900 µL PBS. The resulting erythrocytes were kept in 50% PBS (pH 6.5). 50 µL of 50% neuraminidase-treated erythrocytes were incubated with 2.5 µM *O*-acylated GD3 in 450 µL PBS (pH 6.5) at 37 °C for 1 h. The modified human blood cells were washed with 3-times with PBS and stored in 50% PBS for further use.

##### 1.10 Hemagglutination Assay

HKU1-S1$^{A}$-Fc and pA-LS were pre-complexed in a 1:1 molar ratio on ice for 30 min. The hemagglutination assay with a starting concentration of 20 µg/mL was performed with human erythrocytes at pH 6.5 according to the standard methods.[4] Briefly, in a conical-bottom 96-well plate, a 2-fold dilution of pre-complexed HKU1-NTD-FC and pA-LS were incubated with 1% human erythrocytes at 4 °C for approximately 4 h. The precipitation of cells is visually distinguishable from those that are hemagglutinated.

##### 1.11 Miscellaneous Information

Enzymatic reactions were performed in MaxQ 4450 incubator (Thermo Scientific) at 37 °C with shaking. RT was controlled at 21 (±1) °C. Chemical structures were made using ChemDraw Professional 16.0. NMR spectra were processed and analyzed using MestReNova.

#### 3. Synthetic Methods and Compound Characterization

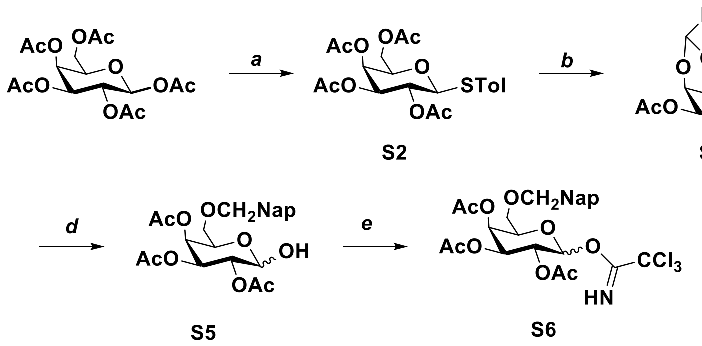


***Scheme S1.*** Monosaccharide building block **S6**. Reagents and conditions: *a*) PhSTol, $BF_3$•$OEt_2$, $CH_2Cl_2$, 99%. *b*) i MeoNa, MeOH; ii NapCH$(OMe)_2$, camphorsulfonic acid, $CH_3CN$; iii $Ac_2O$, Pyiride, 88% over 3 steps. *c*) i $Et_3SiH$, trifluoroacetic acid (TFA), $CH_2Cl_2$; ii $Ac_2O$, Pyiride, 64% over 2 steps. *d*) NBS, acetone/$H_2O$, 69%. *e*) trichloroacetonitrile, DBU, $CH_2Cl_2$, 76%.

**Tolyl 2,3,4,6-tetra-*O*-acetyl-1-thio-β-D-galactopyranoside (S2)**

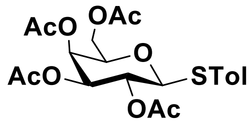


To a solution of D-galactose pentaacetate **S1** (20.0 g, 51.3 mmol) and 4-tolyl mercaptan (TolSH) (7.0 g, 56.4 mmol) in $CH_2Cl_2$ (200 mL) was added $BF_3$·$OEt_2$ (16.1 mL, 128.0 mmol) at 0 °C. The resulting reaction mixture was stirred at RT under an argon atmosphere. After 6 h, the reaction was quenched at 0 °C by the addition of saturated aq. $NaHCO_3$. The organic layer was separated, dried over $Na_2SO_4$, filtered, and the filtrate concentrated under reduced pressure. The residue was purified by silica gel column chromatography (petroleum ether: ethyl acetate = 1: 1) to provide **S2** as a colorless liquid (23.0 g, 99%). $^1$H NMR (600 MHz, $CDCl_3$) $\delta$ 7.41 (d, $J$ = 8.1 Hz, 2H, Ar), 7.13 (d, $J$ = 7.9 Hz, 2H, Ar), 5.41 (dd, $J_{3,4}$ = 3.3 Hz, $J_{4,5}$ = 1.1 Hz, 1H, H-4), 5.21 (t, $J_{1,2}$=$J_{2,3}$ = 10.0 Hz, 1H, H-2), 5.05 (dd, $J_{2,3}$ = 10.0 Hz, $J_{3,4}$=3.3 Hz, 1H, H-3), 4.66 (d, $J_{1,2}$ = 10.0 Hz, 1H, H-1), 4.22 – 4.10 (m, 2H, H-6), 3.95

– 3.90 (m, 1H, H-5), 2.34 (s, 3H, C<u>H</u>$_3$Ph), 2.12 (s, 3H, Ac), 2.11 (s, 3H, Ac), 2.06 (s, 3H, Ac); $^{13}$C NMR (150 MHz, $CDCl_3$) $\delta$ 170.83, 170.67, 169.91, 138.53, 133.17, 129.67, 128.51, 86.86 (C-1), 74.28 (C-4), 72.17 (C-3), 67.44 (C-5), 67.40 (C-2), 61.86 (C-6), 21.16 (<u>C</u>H$_3$Ph), 20.86 (CO<u>C</u>H$_3$), 20.67 (CO<u>C</u>H$_3$), 20.62 (CO<u>C</u>H$_3$), 20.58 (CO<u>C</u>H$_3$); LCMS(ESI-TOF): *m/z* calculated for $C_{21}H_{26}OSNa$ ([M+Na]$^+$) 447.1190, found 447.1058.

**Tolyl 2,3-di-*O*-acetyl-4,6-*O*-(2-naphthylmethylidene)-1-thio-β-D-galactopyranoside (S3)**

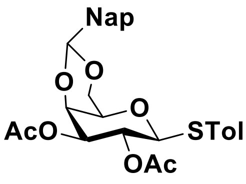


To a solution of **S2** (39.2 g, 86.2 mmol) in $CH_3OH$ (300 mL) was added $NaOCH_3$ (0.5 g, 8.6 mmol). The mixture was stirred for 1 h at RT and then neutralized with DOWEX 50W (H$^+$-form). The resin was removed by filtration, and the filtrate was concentration under reduced pressure to provide the corresponding 2,3,4,6-tetraol, which was used in the next step without further purification. 2-Naphthaldehyde dimethyl acetal (15.5 g, 76.9 mmol) and camphorsulfonic acid (0.6 g, 2.5 mmol) were added to a solution of the residue in $CH_3CN$ (300 mL) at RT. After stirring overnight, the reaction mixture was neutralized by the addition of $NEt_3$ and then concentrated under reduced pressure. The resulting residue was dissolved in $CH_2Cl_2$, washed with saturated aq. $NaHCO_3$, the organic layer was dried over $Na_2SO_4$. After filtration, the solution was concentrated under reduced pressure. Petroleum ether was added to the mixture, which was then stored at −20 °C for 30 min. The solids were filtered off, washed with petroleum ether, and used in the next step. To a solution of the residue in pyridine (60 mL) was added $Ac_2O$ (30 mL) and DMAP (0.2 g, 1.6 mmol) at 0 °C. After stirring at RT for 24 h, the solvents were evaporated under reduced pressure. The residue was dissolved in $CH_2Cl_2$ and washed with 1 N aq. HCl, saturated aq. $NaHCO_3$ and brine. The organic layer was dried over $Na_2SO_4$. After filtration, the solution was concentrated to afford **S3** (23.0 g, 88%) as white foam. $^1$H NMR (600 MHz, $CDCl_3$) $\delta$ 7.92 – 7.05 (m, 7H, Ar), 5.62 (s, 1H, -C<u>H</u>Nap), 5.33 (t, $J_{1,2}$ = $J_{2,3}$ = 9.8 Hz, 1H, H-2), 5.02 (dd, $J_{2,3}$ = 9.9 Hz, $J_{3,4}$ = 3.4 Hz, 1H, H-3), 4.67 (d, $J_{1,2}$ = 9.7 Hz, 1H, H-1), 4.42 (d, $J_{3,4}$ = 3.5 Hz, 1H, H-4), 4.41 (dd, $J_{6a,6b}$ = 12.4 Hz, $J_{5,6a}$ = 1.9 Hz, 1H, H-6a), 4.07 (dd, $J_{6a,6b}$ = 12.4 Hz, $J_{5,6b}$ = 1.7 Hz, 1H, H-6b), 3.61 (d, $J_{5,6b}$ = 1.8 Hz, 1H, H-5), 2.27 (s, 3H, C<u>H</u>$_3$Ph), 2.10 (s, 3H, Ac), 2.02 (s, 3H, Ac); $^{13}$C NMR (150 MHz, $CDCl_3$) $\delta$ 170.76, 169.09, 138.35, 134.90, 134.16, 133.80, 132.83, 129.58, 128.44, 128.02, 127.73, 127.33, 126.40, 126.05, 125.91, 124.20, 101.32 (-<u>C</u>HNap), 85.36 (C-1), 73.59 (C-4), 73.27 (C-3),

69.74 (C-5), 69.21 (C-6), 66.89 (C-2), 21.20 (C̲H$_3$Ph), 20.94 (COC̲H$_3$), 20.91(COC̲H$_3$); LCMS(ESI-TOF): *m/z* calculated for $C_{28}H_{28}O_7SNa$ ([M+Na]$^+$) 531.1448, found 531.1309.

**Tolyl 2,3,4-tri-*O*-acetyl-6-*O*-(2-naphthylmethyl)-1-thio-β-D-galactopyranoside (S4)**

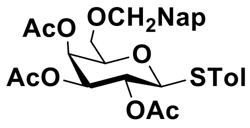


To a solution of **S3** (3.8 g, 7.0 mmol) in $CH_2Cl_2$ (50 mL) was added sequentially $Et_3SiH$ (5.6 mL, 35.2 mmol) and trifluoroacetic acid (2.7 mL, 35.2 mmol) at 0 °C. After complete consumption of starting material (monitored by TLC), $NEt_3$ was added to quench the reaction. Solvents were removed under reduced pressure and the resulting crude product was purified by silica gel column chromatography (petroleum ether: ethyl acetate = 4: 1) to give **S4** (16.0 g, 64%). $^1$H NMR (400 MHz, $CDCl_3$) $\delta$ 7.85 – 7.77 (m, 3H, Ar), 7.74 – 7.69 (m, 1H, Ar), 7.49 – 7.42 (m, 2H, Ar), 7.42 – 7.35 (m, 3H, Ar), 7.06 – 6.99 (m, 2H, Ar), 5.49 (dd, $J_{3,4}$ = 3.3 Hz, $J_{4,5}$ = 1.1 Hz, 1H, H-4), 5.20 (t, $J_{1,2}$ = $J_{2,3}$ = 9.9 Hz, 1H, H-2), 5.04 (dd, $J_{2,3}$ = 9.9 Hz, $J_{3,4}$ = 3.4 Hz, 1H, H-3), 4.68 (d, $J$= 12.0 Hz, 1H,CH̲$_2$Nap), 4.66 (d, $J_{1,2}$ = 10.0 Hz, 1H, H-1), 4.57 (d, $J$= 12.0 Hz, 1H, CH̲$_2$Nap), 3.89 (t, $J_{5,6a}$ = $J_{5,6b}$ = 6.3 Hz, 1H, H-5), 3.62 (dd, $J_{6a,6b}$ = 9.7, $J_{5,6a}$ = 6.2 Hz, 1H, H-6a), 3.52 (dd, $J_{6a,6b}$ = 9.7 Hz, $J_{5,6b}$ = 6.4 Hz, 1H, H-6b), 2.26 (s, 3H, CH̲$_3$Ph), 2.07 (s, 3H, Ac), 1.96 (s, 3H, Ac), 1.95 (s, 3H, Ac); $^{13}$C NMR (100 MHz, $CDCl_3$) $\delta$ 170.17, 170.01, 169.48, 138.13, 135.04, 133.19, 133.03, 132.65, 129.64, 129.62, 129.05, 128.23, 127.91, 127.65, 126.71, 126.13, 125.97, 125.81, 87.05 (C-1), 76.03 (C-5), 73.61(C̲H$_2$Nap), 72.22 (C-3), 67.76 (C-6), 67.74 (C-4), 67.53 (C-2), 21.07 (C̲H$_3$Ph), 20.85(COC̲H$_3$), 20.60 (COC̲H$_3$), 20.57 (COC̲H$_3$); LCMS(ESI-TOF): *m/z* calculated for $C_{30}H_{32}O_8SNa$ ([M+Na]$^+$) 575.1710, found 575.1735.

**6-*O*-(2-naphthylmethyl)-2,3,4-tri-*O*-acetyl galactopyranose hemiacetal (S5)**

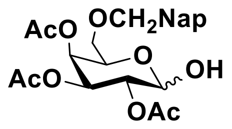


To a solution of **S4** (16.0 g, 29.0 mmol) in acetone/ water (250 mL, 9:1) was added *N*-bromosuccinimide (15.5 g, 87.1 mmol) at RT under an argon atmosphere. After 5 h, the mixture was neutralized by the addition of $NEt_3$ and then concentrated under reduced pressure. The resulting residue was dissolved in $CH_2Cl_2$, washed with 1N aq. HCl and saturated brine, dried over $Na_2SO_4$, filtered, and concentrated under reduced pressure. The crude product was purified by silica gel column chromatography (petroleum ether: ethyl acetate = 5: 1 to 2: 1) to give **S5** (9 g, 69%, α/β = 3:1) as an amorphous solid. The α/β ratio was determined by $^1$H NMR analysis. α-configuration: $^1$H NMR (600 MHz, $CDCl_3$) $\delta$ 7.82 –7.38

(m, 7H, Ar), 5.50 (d, $J_{1,2}$ = 3.9 Hz, 1H, H-1), 5.46 (d, $J_{3,4}$ = 3.6 Hz, 1H, H-4), 5.41 (dd, $J_{2,3}$ = 10.9 Hz, $J_{3,4}$ = 3.6 Hz, 1H, H-3), 5.14 (dd, $J_{2,3}$ = 10.8 Hz, $J_{1,2}$ = 3.8 Hz, 1H, H-2), 4.67 (d, $J$= 12.1 Hz, 1H, C<u>H</u>$_2$Nap), 4.55 (d, $J$= 12.2 Hz, 1H, C<u>H</u>$_2$Nap), 4.46 (t, $J_{5,6a}$ = $J_{5,6b}$ = 6.3 Hz, 1H, H-5), 3.51 (dd, $J_{6a,6b}$ = 9.7 Hz, $J_{5,6b}$ = 6.3 Hz, 1H, H-6a), 3.42 (dd, $J_{6a,6b}$ = 9.8 Hz, $J_{5,6a}$ = 6.0 Hz, 1H, H-6b), 2.06 (s, 3H, Ac), 1.99 (s, 3H, Ac), 1.98 (s, 3H, Ac); $^{13}$C NMR (150 MHz, $CDCl_3$) $\delta$ 170.56, 170.38, 170.15, 134.74, 133.17, 133.07, 128.40, 127.93, 127.75, 126.87, 126.27, 126.09, 125.82, 90.61 (C-1), 73.53 (<u>C</u>H$_2$Nap), 68.92 (C-4), 68.54 (C-2), 68.31 (C-6), 67.63 (C-3), 67.25 (C-5), 20.83 (CO<u>C</u>H$_3$), 20.71 (CO<u>C</u>H$_3$), 20.62 (CO<u>C</u>H$_3$); LCMS(ESI-TOF): *m/z* calculated for $C_{23}H_{26}O_9Na$ ([M+Na]$^+$) 469.1469, found 469.1478.

**2,3,4-Tri-*O*-acety-6-*O*-(2-napthylmethyl)-α-D-galactopyranosyl trichloroacetimidate (S6)**

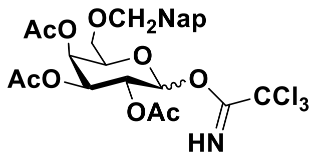


A solution of **S5** (9.0 g, 20.0 mmol) in $CH_2Cl_2$ (200 mL) under an argon atmosphere was cooled to 0 °C. $Cl_3CCN$ (20.0 mL, 200.0 mmol) and 1,8-diazabicyclo[5.4.0]undec-7-ene (0.6 g, 4.0 mmol) were subsequently added. The reaction mixture was stirred at RT. After 12 h, the dark brown reaction mixture was passed through a glass fritted vacuum filter funnel equipped with a plug of Celite. The filter cake and reaction flask were rinsed with $CH_2Cl_2$. The filtrate was concentrated to an oil. The residue was purified by silica gel column chromatography (petroleum ether: ethyl acetate = 4: 1 with 1% $NEt_3$) to give **S6** (8.9 g, 75%). $^1$H NMR (400 MHz, $CDCl_3$) $\delta$ 8.62 (s, 1H, NH), 7.83 – 7.75 (m, 3H, Ar), 7.69 (t, $J$ = 1.6 Hz, 1H, Ar), 7.48 – 7.41 (m, 2H, Ar), 7.36 (dd, $J$ = 8.4, 1.6 Hz, 1H, Ar), 6.57 (d, $J_{1,2}$ = 3.6 Hz, 1H, H-1), 5.65 (dd, $J_{3,4}$ = 3.2 Hz, $J_{4,5}$ = 1.3 Hz, 1H, H-4), 5.43 (dd, $J_{2,3}$ = 10.8 Hz, $J_{3,4}$ = 3.2 Hz, 1H, H-3), 5.32 (dd, $J_{2,3}$ = 10.8 Hz, $J_{1,2}$ = 3.6 Hz, 1H, H-2), 4.68 (d, $J$ = 12.2 Hz, 1H, C<u>H</u>$_2$Nap), 4.54 (d, $J$ = 12.2 Hz, 1H, C<u>H</u>$_2$Nap), 4.42 (ddd, $J_{5,6b}$ = 7.1 Hz, $J_{5,6a}$ = 5.5 Hz, $J_{4,5}$ = 1.3 Hz, 1H, H-5), 3.57 (dd, $J_{6a,6b}$ = 9.7, 5.8 Hz, 1H, H-6a), 3.49 (dd, $J_{6a,6b}$ = 9.7 Hz, $J_{5,6a}$ = 7.1 Hz, 1H, H-6b), 2.02 – 1.94 (m, 9H, Ac); $^{13}$C NMR (100 MHz, $CDCl_3$) $\delta$ 170.11, 170.01, 169.91, 161.00, 134.88, 133.16, 133.02, 128.24, 127.90, 127.64, 126.72, 126.13, 125.97, 125.75, 93.70 (C-1), 73.47 (<u>C</u>H$_2$Nap), 70.13 (C-5), 67.82 (C-4), 67.71(C-3), 67.13 (C-2), 67.11 (C-6), 20.68 (CO<u>C</u>H$_3$), 20.53 (CO<u>C</u>H$_3$), 20.51 (CO<u>C</u>H$_3$); LCMS(ESI-TOF): *m/z* calculated for $C_{25}H_{26}Cl_3NO_9Na$ ([M+Na]$^+$) 612.0565, found 612.0521.

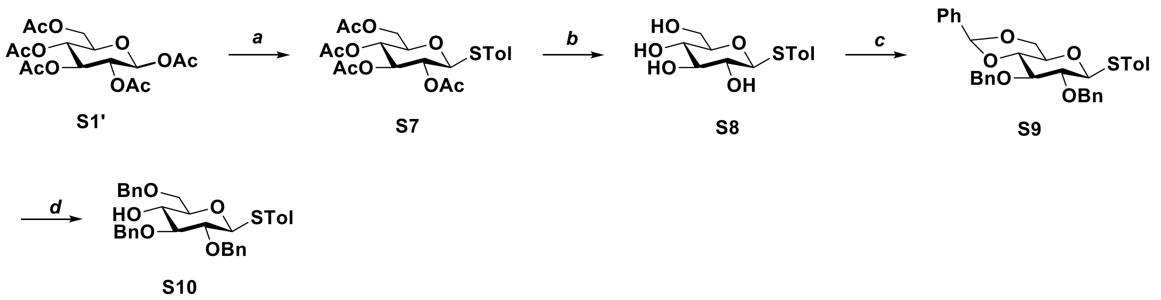

***Scheme S2.*** Monosaccharide building block **S10**. Reagents and conditions: *a*) PhSTol, $BF_3.OEt_2$, $CH_2Cl_2$, 85%. *b*) MeONa, MeOH, 95%. *c*) i $PhCH(OMe)_2$, camphorsulfonic acid, $CH_3CN$; ii BnBr, NaH, DMF, 54% over 2 steps. *d*) $Et_3SiH$, TFA, $CH_2Cl_2$, 89%.

**Tolyl 2,3,4,6-tetra-*O*-acetyl-β-D-thioglucopyranoside (S7)**

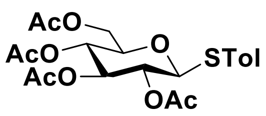

Peracetylated glucose **S1'** (20.0 g, 51.3 mmol) was dissolved in dichloromethane (200 mL) to which was added by p-thiocresol (7.0 g, 56.4 mmol) and boron trifluoride–diethyl etherate (16.1 mL, 128.0 mmol) at 0 °C. The resulting reaction mixture was stirred at RT under argon atmosphere. After 6 h, the reaction was quenched at 0 °C by addition of saturated aq. $NaHCO_3$. The organic layer was separated, dried over $Na_2SO_4$, filtered, and the filtrate concentrated under reduced pressure. The residue was purified by silica gel column chromatography (petroleum ether:ethyl acetate = 1:1) to afford **S7** as an amorphous solid (19.0 g, 85%). The NMR data are in agreement with reported data.[5]

**Tolyl 1-thio-β-D-glucopyranoside (S8)**

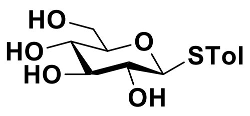

Compound **S7** (8.0 g, 17.6 mmol) was dissolved in methanol (200 mL), and a catalytic amount of sodium methoxide was added dropwise until the pH of the solution reached approximately 9. The reaction mixture was stirred at RT and monitored by TLC until complete deacetylation was observed. The reaction was quenched by the addition of Amberlite IRA-120 ($H^+$) resin to neutralize the solution. The suspension was filtered, and the filtrate was concentrated under reduced pressure to afford compound **S8** as an amorphous solid (4.8 g, 95%). The NMR data are in agreement with reported data.[5]

**Tolyl 2,3-di-*O*-benzyl-4,6-*O*-[(R)-phenylmethylene]-1-thio-β-D-glucopyranoside (S9)**

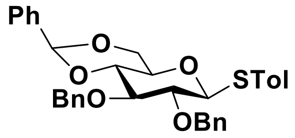


To a solution of compound **S8** (7.3 g, 25.6 mmol) was added 2-benzaldehyde dimethyl acetal (5.8 mL, 38.5 mmol) and camphorsulfonic acid (0.3 g, 1.3 mmol) in $CH_3CN$ (300 mL) at RT. After stirring overnight, the mixture was neutralized by addition of $NEt_3$ and then concentrated under reduced pressure. The resulting residue was dissolved in $CH_2Cl_2$, washed with saturated aq. $NaHCO_3$, dried over $Na_2SO_4$, filtered, and the filtrate concentrated under reduced pressure. Petroleum ether was added to the mixture, which was then stored at −20 °C for 30 min. The solid was filtered off, washed with petroleum ether, and used in the next reaction step. The residue was dissolved in DMF (100 mL). To the solution was added sodium hydride (60% dispersed in mineral oil, 2.9 g) at 0 °C. After 30 min, benzyl bromide (7.2 mL) was added dropwise. The reaction mixture was allowed to warm to RT slowly and stirred for 5 h. The reaction was quenched by the addition of methanol and the solvents were removed under reduced pressure. The residue was diluted with $CH_2Cl_2$, and the insoluble solids were removed by filtration. The filtrate was concentrated and the resulting residue purified by silica gel column chromatography (petroleum ether:ethyl acetate = 8:1) to give compound **S9** (7.5 g, 54% over 2 steps) as a white amorphous solid. The NMR data are in agreement with reported data.[6]

**Tolyl 2,3,6-tri-*O*-benzyl-1-thio-β-D-glucopyranoside (S10)**

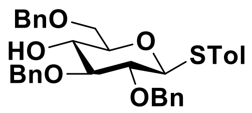


Trifluoroacetic acid (5.2 ml, 68.0 mmol) was slowly added to a solution of **S9** (7.5 g, 13.5 mmol) and triethylsilane (10.8 ml, 68.0 mmol) in $CH_2Cl_2$ (100 ml) at 0 ºC under argon atmosphere. After 2 h, the reaction mixture was diluted with $CH_2Cl_2$ and quenched with ice and then washed with sat. aq. $NaHCO_3$ and brine. The organic layer was separated, dried over $Na_2SO_4$, filtered, and the filtrate concentrated under reduced pressure. The resulting residue was purified by silica gel column chromatography (petroleum ether:ethyl acetate = 4:1) to afford **S10** (6.7 g, 89%) as a colourless oil. $^1$H NMR (400 MHz, $CDCl_3$) $\delta$ 7.53 – 7.41 (m, 4H, Ar), 7.40 – 7.28 (m, 13H, Ar), 7.07 (d, $J$ = 7.9 Hz, 2H, Ar), 4.95 (d, $J$ = 10.3 Hz, 1H, C$\underline{H_2}$Ph), 4.93 (d, $J$ = 11.3 Hz, 1H, C$\underline{H_2}$Ph), 4.80 (d, $J$ = 11.3 Hz, 1H, C$\underline{H_2}$Ph), 4.76 (d, $J$ = 10.3 Hz, 1H, , C$\underline{H_2}$Ph), 4.65 (d, $J_{1,2}$ = 8.8 Hz, 1H, H-1), 4.58 (d, $J$ = 4.3 Hz, 2H, C$\underline{H_2}$Ph), 3.85 – 3.73 (m, 2H, H-6a, H-6b), 3.66 (td, $J_{3,4}$ = 9.2 Hz, $J_{4,5}$ =2.3 Hz, 1H, H-4), 3.55 (t, $J_{1,2}$ = $J_{2,3}$ = 8.7 Hz, 1H, H-3), 3.52 – 3.46 (m, 2H, H-2, H-5), 2.33 (s, 3H, C$\underline{H_3}$Ph); $^{13}$C NMR

(100 MHz, $CDCl_3$) $\delta$ 138.50, 138.06, 137.99, 137.77, 132.63, 129.81, 129.70, 128.62, 128.46, 128.43, 128.28, 127.96, 127.93, 127.91, 127.73, 127.72, 87.96 (C-1), 86.22 (C-3), 80.48 (C-5), 78.11 (C-2), 75.49 ($\underline{C}H_2Ph$), 75.35 ($\underline{C}H_2Ph$), 73.66 ($\underline{C}H_2Ph$), 71.71 (C-4), 70.41 (C-6), 21.14 ($\underline{C}H_3Ph$); LCMS(ESI-TOF): *m/z* calculated for $C_{34}H_{36}O_5Na$ ($[M+Na]^+$) 579.2176, found 579.2158.

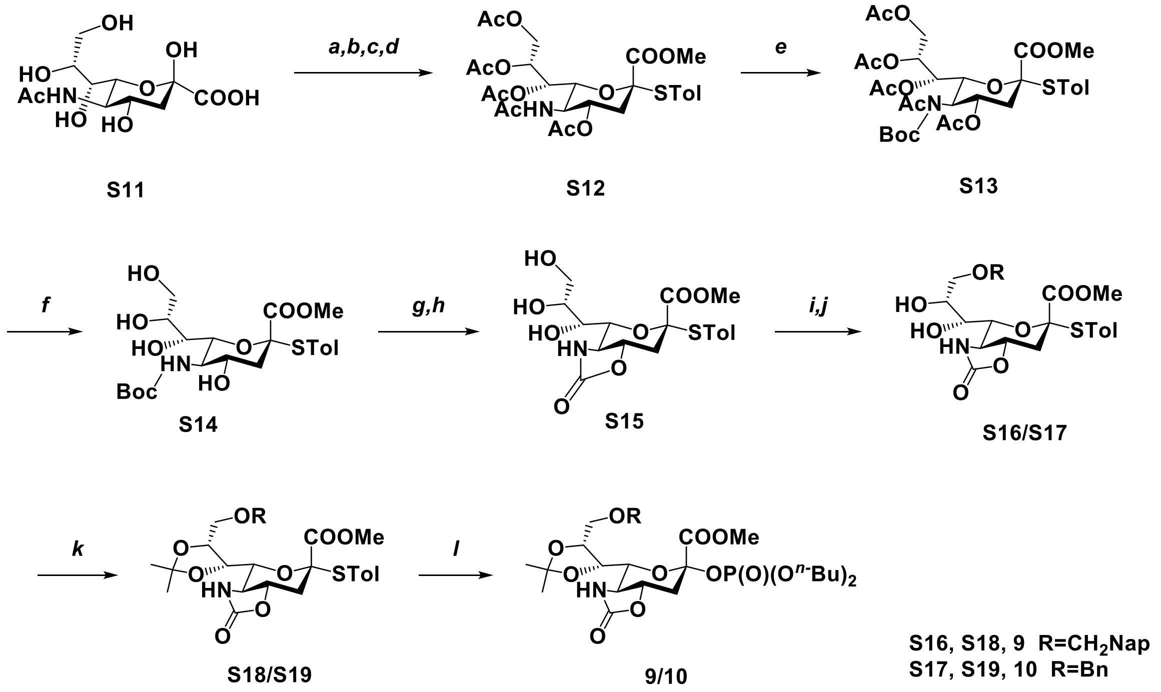


***Scheme S3.*** Monosaccharide building blocks **9** and **10**. Reagents and conditions: *a*) $H^+$ Resin, MeOH. *b*) $Ac_2O$, pyridine. *c*) AcCl, MeOH. *d*) p-toluenethiol, NaH, DMF, 40% over 4 steps. *e*) $Boc_2O$, DMAP, THF, 60°C, 89%. *f*) NaOMe, MeOH, 66%. *g*) AcCl, MeOH, $CH_2Cl_2$. *h*) $4\text{-}NO_2C_6H_4OCOCl$, $NaHCO_3$, $CH_3CN/H_2O$, 72% over 2 steps. *i*) $NapCH(OMe)_2$ for **S16**; $PhCH(OMe)_2$ for **S17**, Camphorsulfonic acid, $CH_3CN$. *j*) $BH_3NMe_3$, $AlCl_3$, THF,73% (**S16**), 76% (**S17**) over 2 steps. *k*) 2,2-Dimethoxypropane, Camphorsulfonic acid, $CH_3CN$, 83% (**S18**), 85% (**S19**). *l*) Dibutyl phosphate, NIS, TfOH, $CH_2Cl_2$, 52% (**9**), 67% (**10**).

#### Methyl 4-methylphenyl 5-acetamido-4,7,8,9-tetra-*O*-acetyl-3,5-dideoxy-2-thio-D-glycero-α-D-galacto-2-nonulopyranosonate (S12)

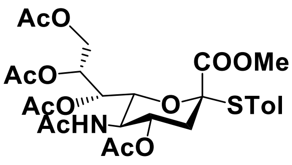

A mixture of *N*-acetylneuraminic acid monohydrate **S11** (20.0 g, 64.7 mmol) and Amberlite IR-120 $H^+$ resin (9.0 g) in methanol (600 mL) was stirred at 60 °C for 6 h. The reaction mixture was filtered and the filtrate concentrated under reduced pressure to yield methyl 5-acetamido-3,5-dideoxy-D-glycero-β-D-galacto-non-2-ulopyranosonate as an off white solid (20.0 g, quant.). The residue was used without additional purification. To a solution of the residue in pyridine (200 mL) was added $Ac_2O$ (100 mL) and DMAP (1.0 g, 8.0 mmol) at 0 °C. After stirring at RT for 24 h, the solvents were evaporated under reduced pressure. The residue was dissolved in $CH_2Cl_2$ and washed with 1N aq. HCl, saturated aq. $NaHCO_3$ and brine. The organic layer was separated, dried over $Na_2SO_4$, filtered, and the filtrate concentrated under reduced pressure to afford per-acetylated product as a yellow foam (35.0 g, quant.). The residue was used without additional purification. Methanol (70 mL) was added dropwise to AcCl (250 mL) with cooling in an ice-salt bath. The resulting solution was added to a cold solution of glycosyl acetate (35.0 g, 64.7 mmol) in a mixture of $CH_2Cl_2$ (1 L) and the reaction mixture was kept at RT for 72 h (TLC control: $R_f = 0.6$). Volatile components were evaporated, $CH_2Cl_2$ (100 mL) was added, and the solution was concentrated again. Addition of $CH_2Cl_2$ and concentration was repeated until no smell of HCl could be detected. The residue was dried *in vacuo* to give the glycosyl chloride as a slight yellow amorphous solid (32 g; $R_f = 0.6$; ethyl acetate). The residue was used without additional purification. To a solution of 4-methylthiophenol (4.5 g, 35.8 mmol) in DMF (50 mL) was slowly added NaH (60 % dispersion in mineral oil, 1.3 g, 23.8 mmol) at 0 °C, stirred for 10 min at this temperature. The mixture was added dropwise to a solution of crude sia-chloride (15.2 g, 29.8 mmol) in DMF (150 mL). After stirring the reaction mixture at RT for 1 h, it was extracted with $CH_2Cl_2$ (200 mL) and washed with saturated brine. The organic layer was dried over $Na_2SO_4$, filtered, and the filtrate concentrated under reduced pressure. Petroleum ether (1 L) was added to the residue and the resulting suspension was stirred overnight at RT, then filtrated and the residue was crystallized from $CH_2Cl_2$ – hexanes to give to **S12** as a yellow amorphous solid (15.7 g, 40 % over 4 steps). $^1$H NMR (400 MHz, $CDCl_3$) $\delta$ 7.35 (d, $J$ = 8.1 Hz, 2H, Ar), 7.10 (d, $J$ = 7.9 Hz, 1H, Ar), 5.36 (d, $J$ = 9.8 Hz, 1H, N<u>H</u>), 5.26 (dd, $J_{7,8}$ = 6.5 Hz, $J_{6,7}$ = 1.9 Hz, 1H), 5.22 (td, $J_{7,8}$ = 6.1Hz, $J_{8,9a}$ = 2.7 Hz, 1H), 4.37 (d, $J_{8,9a}$ = 2.7 Hz, 1H), 4.18 (dd, $J_{9a,9ab}$ = 12.4 Hz, $J_{8,9b}$ = 5.9 Hz, 1H), 4.80 (ddd, $J_{3a,4}$ = 11.8 Hz, $J_{4,5}$ = 10.1 Hz, $J_{3e,4}$ = 4.7 Hz, 1H, H-4), 3.93 (t, $J_{4,5} = J_{5,6}$ = 10.2 Hz, 1H, H-5), 3.85 (dd, $J_{5,6}$ = 10.7 Hz, $J_{6,7}$ = 2.0 Hz, 1H, H-6), 3.57 (s, 3H, OC<u>H</u>3), 2.74 (dd, $J_{3e,3a}$ = 12.9 Hz, $J_{3e,4}$ = 4.7 Hz, 1H), 2.33 (s, 3H, C<u>H</u>3Ph), 2.10 (s, 3H, Ac), 2.02 (s, 3H, Ac), 2.01 (s, 3H, Ac), 2.00 (s, 3H, Ac), 1.97 (s, 3H, Ac), 1.94 (m, 1H, H-3ax); $^{13}$C NMR (100 MHz, $CDCl_3$) $\delta$ 170.89, 170.60, 170.20, 170.17, 170.02, 168.04 (C-1), 140.23, 136.49, 129.60, 124.95, 87.46 (C-2), 74.82 (C-6), 70.28 (C-4), 69.72 (C-8), 67.79 (C-7), 62.02 (C-9), 52.70 (O<u>C</u>H3), 49.14 (C-5), 37.99 (C-3), 23.10 (NHCO<u>C</u>H3), 21.29

(CO$\underline{C}$H$_3$), 20.95 (CO$\underline{C}$H$_3$), 20.81 (CO$\underline{C}$H$_3$), 20.78 (CO$\underline{C}$H$_3$), 20.72 (CO$\underline{C}$H$_3$); HRMS (ESI): *m/z* calculated for $C_{27}H_{35}NO_{12}SNa$ ([M+Na]$^+$) 620.1772, found 620.1782.

**Methyl 4-methylphenyl 5-acetamido-4,7,8,9-tetra-*O*-acetyl-3,5-dideoxy-5-*tert*-butoxycarbonyl-2-thio-D-glycero-α-D-galacto-2-nonulopyranosonate (S13)**

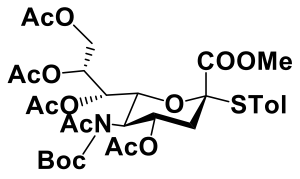


To a solution of **S12** (15.7 g, 26.3 mmol) in anhydrous THF (135 mL) were added di-tertbutyl dicarbonate (11.5 g, 52.6 mmol) and DMAP (1.0 g, 7.9 mmol) at RT. The mixture was stirred for 2 h at 60 °C. Then it was cooled to RT and concentrated under reduced pressure. The residue was purified by silica gel column chromatography (petroleum ether:ethyl acetate = 15:1 to 5:1) to give **S13** as yellow oil (16.3 g, 89%). $^1$H NMR (400 MHz, CDCl$_3$) $\delta$ 7.32 (d, $J$ = 8.1 Hz, 2H, Ar), 7.08 (d, $J$ = 8.0 Hz, 3H, 2H, Ar), 5.30 (td, $J_{3a,4}$ = 10.9 Hz, $J_{3e,4}$ = 5.0 Hz, 1H, H-4), 5.16 (m, 1H, H-8), 5.08 (d, $J_{6,7}$ = 1.7 Hz, 1H, H-7), 4.75 (t, $J_{4,5}$ = $J_{5,6}$ = 10.5 Hz, 1H, H-5), 4.48 (dd, $J_{5,6}$ = 10.3, $J_{6,7}$ = 1.7 Hz, 1H, H-6), 4.35 (dd, $J_{9a,9b}$ = 12.4 Hz, $J_{8,9a}$ = 2.7 Hz, 1H, H-9a), 4.12 (dd, $J_{9a,9b}$ = 12.5 Hz, $J_{8,9b}$ = 5.6 Hz, 1H, H-9b), 3.52 (s, 3H, OC$\underline{H}_3$), 2.86 (dd, $J_{3a,3e}$ = 12.8 Hz, $J_{3e,4}$ = 5.0 Hz, 1H, H-3eq), 2.30 (s, 6H, Ac, C$\underline{H}_3$Ph), 2.04 (s, 3H, Ac), 2.01 – 1.94 (m, 7H, Ac, H-3ax), 1.89 (s, 3H, Ac), 1.45 (s, 9H, C$\underline{H}_3^{Boc}$); $^{13}$C NMR (100 MHz, CDCl$_3$) $\delta$ 173.85, 170.51, 170.08, 169.81, 169.65, 168.05 (C-1), 151.63, 140.15, 136.37, 129.56, 125.13, 87.52 (C-2), 84.62, 73.87 (C-6), 70.09 (C-8), 67.18 (C-4, C-7), 61.66 (C-9), 52.54 (O$\underline{C}$H$_3$), 52.32 (C-5), 39.16 (C-3), 27.70 ($\underline{C}$H$_3^{Boc}$), 26.65 (NCO$\underline{C}$H$_3$), 21.28 ($\underline{C}$H$_3$Ph), 20.91(CO$\underline{C}$H$_3$), 20.78 (CO$\underline{C}$H$_3$), 20.69 (CO$\underline{C}$H$_3$), 20.62 (CO$\underline{C}$H$_3$); HRMS (ESI): *m/z* calculated for $C_{32}H_{43}NO_{14}SNa$ ([M+Na]$^+$) 720.2296, found 720.2312.

**Methyl 5-(tert-butoxycarbamado)-2,3,5-dideoxy-2-para-methylthiophenol-D-glycero-α-galacto-non-2-ulopyranosonate (S14)**

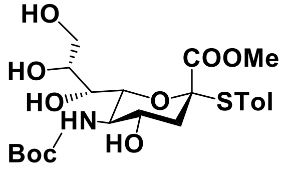


To a solution of **S13** (16.3 g, 23.4 mmol) in methanol (150 mL) were added dropwise 30 wt% aq. NaOMe (40 drops). After stirring the reaction mixture at RT for 2 h, Amberlite IR-120 H$^+$ resin was added to make the solution pH ~ 6. After filtration, the solution was concentrated under reduced pressure and the residue purified by silica gel column chromatography (petroleum ether: ethyl acetate = 5: 1 to 1:1) to give **S14** as white foam (9.9 g, 87%). $^1$H NMR (400 MHz, CDCl$_3$) δ 7.43 (d, $J$ = 8.0 Hz, 2H, Ar), 7.18 (d, $J$ = 8.0 Hz, 2H, Ar), 4.83 (d, $J$ = 8.4 Hz, 1H, NH), 3.92 (m,

1H, H-9a), 3.87 (m, 1H, H-8), 3.73 (m, 1H, H-9b), 3.68 (m, 1H, H-5), 3.66 (s, 3H, OC<u>H</u>3), 3.63 (m, 1H, H-7), 3.54 (m, 1H, H-4), 3.33 (d, $J_{5,6}$ = 10.3 Hz, 1H, H-6), 2.93 (dd, $J_{3a,3e}$ = 13.0 Hz, $J_{3e,4}$ = 4.6 Hz, 1H, H-3eq), 2.50 (t, $J$ = 5.6 Hz, 1H), 2.39 (s, 3H, C<u>H</u>3Ph), 2.04 – 1.95 (m, 1H, H-3ax), 1.44 (s, 9H, $C\underline{H}_3^{Boc}$); $^{13}$C NMR (100 MHz, $CDCl_3$) δ 169.95 (C-1), 157.79, 140.61, 136.49, 129.63, 124.85, 86.69 (C-2), 80.95, 76.90 (C-6), 71.40 (C-8), 69.47 (C-7), 68.87 (C-4), 64.19 (C-9), 53.04 ($O\underline{C}H_3$, C-5), 40.18 (C-3), 28.21($\underline{C}H_3^{Boc}$), 21.33 ($\underline{C}H_3Ph$); HRMS (ESI): *m/z* calculated for $C_{22}H_{33}NO_9SNa$ ($[M+Na]^+$) 510.1768, found 510.1776.

**Methyl (4-methylphenyl 5-amino-5-N,4-*O*-carbonyl-3,5-dideoxy-2-thio-D-glycero-α-D-galacto-non-2-ulopyranoside) onate (S15)**

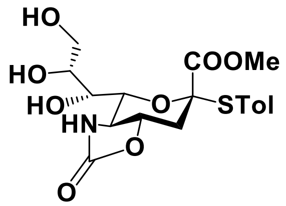


Acetyl chloride (11.0 mL, 156.0 mmol) was added to cooled (0 °C) methanol (32 mL) and stirred at this temperature for 0.5 h. The mixture was added to a solution of **S14** (7.6 g, 15.6 mmol) in $CH_2Cl_2$ (200 mL). After stirring the reaction mixture for 0.5 h at RT, it was concentrated under reduced pressure to give crude product as white solid. The residue was used without additional purification. The residue was dissolved in $CH_3CN/H_2O$ (1: 2, 120 mL) in the presence of $NaHCO_3$ (6.5 g, 77.0 mmol). The mixture was cooled to 0 °C and 4-nitrophenyl chloroformate (7.8 g, 38.5 mmol) in MeCN (40 mL) was added under vigorously stirred mixture over 20 min through an addition funnel. The resulting reaction mixture was stirred for an additional 12 h at 0 °C. After being warmed to RT, the reaction mixture was quenched with a 10% aq. HCl solution, and the suspended solid was dissolved with ethyl acetate. The aqueous layer was extracted with ethyl acetate, washed with saturated brine, dried over $Na_2SO_4$, and concentrated under reduced pressure. The residue was purified by silica gel column chromatography, eluting first with ethyl acetate and then with ethyl acetate: methanol (10: 1 to 5: 1) to give compound **S15** as white foam (6.1 g, 72% over two steps). $^1$H NMR (400 MHz, $CDCl_3$) $\delta$ 7.32 (d, $J$ = 8.0 Hz, 2H, Ar), 7.12 (d, $J$ = 7.8 Hz, 2H, Ar), 6.59 (b, 1H, N<u>H</u>), 3.95 (m, 1H, H-4), 3.85 (dd, $J_{9a,9b}$ = 12.0 Hz, $J_{8,9a}$ = 2.8 Hz, 1H, H-9a), 3.80 – 3.75 (m, 1H, H-9b), 3.76 – 3.67 (m, 6H, $OC\underline{H}_3$, H-6, H-7, H-8), 3.56 (t, $J_{4,5}$ = $J_{5,6}$ = 10.3 Hz, 1H, H-5), 3.13 (dd, $J_{3a,3e}$ = 11.9 Hz, $J_{3e,4}$ = 3.8 Hz, 1H, H-3eq), 2.34 (s, 3H, $C\underline{H}_3Ph$), 2.21 (t, $J_{3a,3e}$ = $J_{3a,4}$ = 12.0 Hz, 1H, H-3ax); $^{13}$C NMR (100 MHz, $CDCl_3$) $\delta$ 168.97 (C-1), 159.49, 140.99, 136.48, 129.74, 124.50, 87.55 (C-2), 78.47 (C-6), 78.08 (C-4), 71.34 (C-7), 69.87 (C-8),

62.51 (C-9), 57.16, 53.54 (O<u>C</u>$H_3$), 36.94 (C-3), 21.34 (<u>C</u>$H_3$Ph); HRMS (ESI): *m/z* calculated for $C_{18}H_{24}NO_8S$ ($[M+H]^+$) 414.1217, found 414.1403.

**Methyl (4-methylphenyl 5-amino-9-*O*-naphthylmethyl-5-*N*,4-*O*-carbonyl-3,5-dideoxy-2-thio-D-glycero-α-D-galacto-2-nonulopyranosid) onate (S16)**

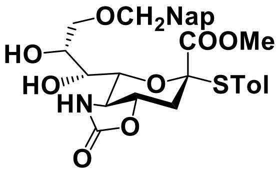


To a stirred solution of the compound **S15** (1.5 g, 3.5 mmol) in $CH_3CN$ (20 mL) were added 2-naphthaldehyde dimethyl acetal (0.8 g, 4.2 mmol) and camphorsulfonic acid (41.0 mg, 0.2 mmol) at RT under argon atmosphere. After being stirred at RT for 0.5 h, the reaction mixture was neutralized with triethylamine and poured into ice-cooled 1N HCl. The aqueous phase was extracted with two portions of ethyl acetate. The organic layer was washed with saturated aq. $NaHCO_3$ and brine, dried over $Na_2SO_4$, filtered, and the filtrate concentrated *in vacuo* to give a crude product as white foam. The residue was used without additional purification. To a solution of crude mixture and pulverizaed activated MS-4Å in THF (40 mL) was added borane-trimethylamine complex (1.6 g, 22.5 mmol) and aluminum chloride (2.9 g, 21.8 mmol) at 0 °C. After being stirred at RT for 3 h, the reaction mixture was poured into ice-cooled solution of 1 N HCl and $CH_2Cl_2$ for 20 min at 0 °C. The combined extracts were washed with saturated aq. $NaHCO_3$ and brine, dried over $Na_2SO_4$, filtered, and the filtrate concentrated *in vacuo*. The residue was purified by silica gel column chromatography (toluene:ethyl acetate = 10:1 to 1:1) to give pure **S16** (1.41 g, 73% over 2 steps) as yellow foam. $^1$H NMR (400 MHz, $CDCl_3$) $\delta$ 7.87 – 7. 07 (m, 11H, Ar), 5.84 (s, 1H, N<u>H</u>), 4.73 (d, $J$ = 2.1 Hz, 2H, C<u>H</u>$_2$Nap), 3.85 (m, 1H, H-4), 3.83(m, 1H, H-8), 3.70 (m, 2H, H-9a, H-9b), 3.67 (m, 1H, H-6), 3.64 (m, 1H, H-7), 3.58 (s, 3H, OC<u>H</u>$_3$), 3.50 (t, $J_{4,5} = J_{5,6}$ = 10.4 Hz, 1H, H-5), 3.14 (dd, $J_{3a,3e}$ = 11.9 Hz, $J_{3e,4}$ = 3.7 Hz, 1H, H-3eq), 2.31 (s, 3H, C<u>H</u>$_3$Ph), 2.15 (t, $J_{3a,3e} = J_{3a,4}$ = 12.0 Hz, 1H, H-3ax); $^{13}$C NMR (100 MHz, $CDCl_3$) $\delta$ 168.69 (C-1), 159.46, 140.94, 136.48, 135.03, 133.20, 133.04, 129.79, 129.01, 128.35, 128.19, 127.87, 127.71, 126.73, 126.25, 126.07, 125.67, 125.27, 124.60, 87.57(C-2), 79.61 (C-6), 78.10 (C-4), 73.66 (<u>C</u>H$_2$Nap), 72.45 (C-7), 71.30 (C-9), 69.86 (C-8), 57.58 (C-5), 53.13 (O<u>C</u>H$_3$), 36.80 (C-3), 21.33 (<u>C</u>H$_3$Ph); HRMS (ESI): *m/z* calculated for $C_{29}H_{31}NO_8SNa$ ($[M+Na]^+$) 576.1663, found 576.1672.

**Methyl (4-methylphenyl 5-amino-9-*O*-benzyl-5-*N*,4-*O*-carbonyl-3,5-dideoxy-2-thio-D-glycero-α-Dgalacto-2-nonulopyranosid) onate (S17)**

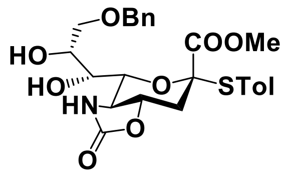

To a stirred solution of the compound **S15** (6.1 g, 14.8 mmol) in $CH_3CN$ (75 mL) was added benzaldehyde dimethyl acetal (2.7 g, 17.7 mmol) and camphorsulfonic acid (0.2 g, 0.7 mmol) at RT under argon. After being stirred at RT for 0.5 h, the reaction mixture was neutralized with triethylamine and poured into ice-cooled 1 M HCl. The aqueous phase was extracted with two portions of ethyl acetate. The extract was washed with saturated aq. $NaHCO_3$ and brine, dried over $Na_2SO_4$, filtered, and the filtrate concentrated *in vacuo* to give a crude product as white foam. The residue was used without additional purification. To a solution of crude product and pulverizaed activated MS-4Å in THF (150 mL) was added borane-trimethylamine complex (6.7 g, 91.6 mmol) and aluminum chloride (11.8 g, 88.6 mmol) at 0 °C. After being stirred at RT for 3 h, the reaction mixture was poured into ice-cooled solution of 1 N HCl and $CH_2Cl_2$ for 20 min at 0 °C. The extract was washed with saturated aq. $NaHCO_3$ and brine, dried over $Na_2SO_4$, filtered, and the filtrate concentrated *in vacuo*. The residue was purified by silica gel column chromatography (toluene:ethyl acetate = 10:1 to 1:1) to give pure **S17** (5.7 g, 76% over 2 steps) as yellowish foam. $^1H$ NMR (400 MHz, $CDCl_3$) $\delta$ 7.44 – 7.08 (m, 9H, Ar), 6.48 (s, 1H, N$\underline{H}$), 4.58 (s, 2H, C$\underline{H}_2$Ph), 3.91 (m, 1H, H-8), 3.87 (dd, $J_{3a,4}$ = 12.3 Hz, $J_{3e,4}$ = 3.8 Hz, 1H, H-4), 3.76 – 3.69 (m, 3H, H9a, H9b, H-6), 3.68 (m, 1H, H-7), 3.59 (s, 3H, OC$\underline{H}_3$), 3.57 (m, 1H, H-5), 3.11 (dd, $J_{3a,3e}$ = 11.9 Hz, $J_{3e,4}$ = 3.7 Hz, 1H, H-3eq), 2.33 (s, 3H, C$\underline{H}_3$Ph), 2.17 (t, $J_{3a,3e}$ = $J_{3a,4}$ = 12.3 Hz, 1H, H-3ax); $^{13}C$ NMR (100 MHz, $CDCl_3$) $\delta$ 168.90 (C-1), 160.26, 140.87, 137.99, 137.82, 136.53, 129.79, 129.04, 128.46, 128.23, 127.77, 127.75, 126.28, 125.31, 124.70, 87.72 (C-2), 78.97 (C-6), 78.32 (C-4), 73.42 ($\underline{C}H_2$Ph), 71.35 (C-9), 71.15 (C-7), 70.46 (C-8), 57.28 (C-5), 53.20 (O$\underline{C}H_3$), 36.83 (C-3), 21.37 ($\underline{C}H_3$Ph); HRMS (ESI): *m/z* calculated for $C_{25}H_{29}NO_8SNa$ ($[M+Na]^+$) 526.1506, found 526.1528.

**Methyl (4-methylphenyl 5-amino-9-*O*-naphthylmethyl-5-*N*,4-*O*-carbonyl-3,5-dideoxy-7, 8-*O*-isopropylidene-2-thio-D-glycero-D-galacto-2-nonulopyranosid) onate (S18)**

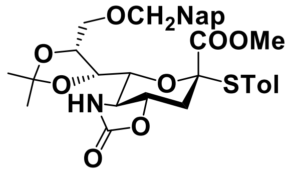

To a solution of compound **S16** (0.9 g, 1.6 mmol) in acetone dimethyl acetal (10.0 mL) was added camphorsulfonic acid (76.0 mg, 0.3 mmol) at RT. After being stirred at RT for 20 min, the reaction mixture was neutralized with triethylamine and concentrated *in vacuo*. The residue was purified by silica gel column chromatography (toluene:ethyl acetate = 5:1) to give **S18** (0.8 g, 83%) as white foam. $^1$H NMR (400 MHz, $CDCl_3$) $\delta$ 7.85 – 7.13 (m, 11H, Ar), 5.76 (s, 1H, N<u>H</u>), 4.65 (d, $J$ = 12.1 Hz, 1H, C<u>H</u>$_2$Ph), 4.51 (m, 1H, H-8), 4.45 (d, 1H, $J$ = 12.1 Hz, C<u>H</u>$_2$Ph), 4.03 (d, $J_{7,8}$ = 6.7 Hz, 1H, H-7), 3.87 (dd, $J_{3a,4}$ = 12.0 Hz, $J_{3e,4}$ = 3.6 Hz, 1H, H-4), 3.81 (d, $J_{8,9a}$ = 7.0 Hz, 1H, H-9a), 3.76 (dd, $J_{9a,9b}$ = 10.3 Hz, $J_{8,9b}$ = 5.0 Hz, 1H, H-9b), 3.67 – 3.58 (m, 2H, H-5, H-6), 3.53 (s, 3H, OC<u>H</u>$_3$), 3.15 (dd, $J_{3a,3e}$ = 12.1 Hz, $J_{3e,4}$ = 3.6 Hz, 1H, H-3eq), 2.30 (s, 3H, C<u>H</u>$_3$Ph), 2.16 (t, $J_{3a,3e}$ = $J_{3a,4}$ = 12.1 Hz, 1H, H-3ax), 1.73 (s, 3H, CC<u>H</u>$_3$), 1.39 (s, 3H, CC<u>H</u>$_3$); $^{13}$C NMR (100 MHz, $CDCl_3$) $\delta$ 168.20 (C-1), 159.70, 140.36, 136.85, 135.51, 133.18, 132.95, 129.56, 128.07, 127.80, 127.66, 126.65, 126.05, 125.85, 125.20, 110.22, 88.87 (C-2), 77.61 (C-4), 76.91 (C-6), 76.18 (C-8), 75.61 (C-7), 73.12 (<u>C</u>H$_2$Nap), 67.81 (C-9), 57.82 (C-5), 52.60 (O<u>C</u>H$_3$), 37.36 (C-3), 26.49 (C<u>C</u>H$_3$), 25.64 (C<u>C</u>H$_3$), 21.33 (<u>C</u>H$_3$Ph); HRMS (ESI): *m/z* calculated for $C_{32}H_{35}NO_8SNa$ ([M+Na]$^+$) 616.1976, found 616.1984.

**Methyl (4-methylphenyl 5-amino-9-*O*-benzyl-5-N,4-*O*-carbonyl-3,5-dideoxy-7, 8-*O*-isopropylidene-2-thio-D-glycero-D-galacto-2-nonulopyranosid) onate (S19)**

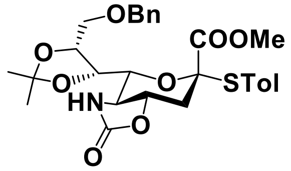

To a solution of compound **S17** (5.7 g, 11.3 mmol) in acetone dimethyl acetal (60.0 ml) was added camphorsulfonic acid (0.5 g, 10.2 mmol) at RT. After being stirred at RT for 20 min, the reaction mixture was neutralized with triethylamine and concentrated *in vacuo*. The residue was purified by silica gel column chromatography (toluene:ethyl acetate = 5:1) to give **S19** (5.4 g, 88%) as white foam. $^1$H NMR (400 MHz, $CDCl_3$) $\delta$ 7.58– 7.15 (m, 9H, Ar), 5.72 (s, 1H, N<u>H</u>), 4.47 (m, 1H, H-8), 4.46 (d, $J$ = 12.0 Hz, 1H, C<u>H</u>$_2$Ph), 4.32 (d, $J$ = 11.9 Hz, 1H, C<u>H</u>$_2$Ph), 4.03 (d, $J_{7,8}$ = 6.7 Hz, 1H, H-7), 3.87 (m, 1H, H-4), 3.75 (m, 2H, H-9a, H-9b), 3.68 – 3.60 (m, 2H, H-5, H-6), 3.53 (s, 3H, OC<u>H</u>$_3$), 3.17 (dd, $J_{3a,3e}$ = 12.0 Hz, $J_{3e,4}$ = 3.6 Hz, 1H, H-3eq), 2.33 (s, 3H, C<u>H</u>$_3$Ph), 2.16 (t, $J_{3a,3e}$ = $J_{3a,4}$ = 12.2 Hz, 1H, H-3ax), 1.72 (s, 3H, CC<u>H</u>$_3$), 1.40 (s, 3H, CC<u>H</u>$_3$); $^{13}$C NMR (100 MHz,

CDCl3) $\delta$ 168.24 (C-1), 159.72, 140.32, 138.06, 136.82, 129.53, 129.00, 128.30, 128.19, 127.81, 127.63, 125.18, 110.20, 88.89 (C-2), 77.62 (C-4), 76.92 (C-6), 76.11 (C-8), 75.64 (C-7), 72.99 (<u>C</u>H$_2$Ph), 67.71 (C-9), 57.82 (C-5), 52.58 (O<u>C</u>H$_3$), 37.37 (C-3), 26.47 (C<u>C</u>H$_3$), 25.62 (C<u>C</u>H$_3$), 21.32 (<u>C</u>H$_3$Ph); HRMS (ESI): *m/z* calculated for $C_{28}H_{33}NO_8SNa$ ([M+Na]$^+$) 566.1819, found 566.1832.

**(3aR,4R,6S,7aS)-Methyl 4-(4S,5R)-5-(naphthylmethyl)-2,2-dimethyl-1,3-dioxolan-4-yl)-6-(dibutoxyphosphoryloxy)-2-oxohexahydro-2H-pyrano[3,4-d] oxazole-6-carboxylate (9)**

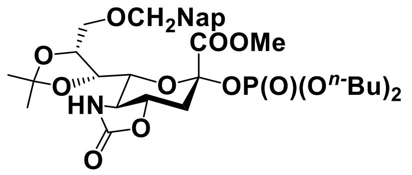


A mixture of thiosialoside donor **S18** (1 g, 1.68 mmol), dibutyl phosphate (1.06 g, 5.06 mmol), and pulverized activated 4Å MS in $CH_2Cl_2$ (20 mL) was stirred under argon at RT for 1 h. The mixture was then cooled to 0 °C, followed by the addition of *N*-iodosuccinimide (0.8 g, 3.4 mmol) and 0.1M TfOH solution in THF (0.8 mL, 0.08 mmol). After stirring for 6 h, the reaction mixture was neutralized with triethylamine and diluted with dichloromethane and filtered through a pad of celite. The filtrate was poured into a mixture of 5% aq. $Na_2S_2O_3$ and saturated aq. $NaHCO_3$. The aqueous layer was extracted with $CH_2Cl_2$. The collected organic phases were washed with saturated brine, dried over $Na_2SO_4$, filtered, and the filtrate concentrated *in vacuo*. The residue was purified by silica gel column chromatography (toluene:ethyl acetate = 10:1 to 1:1) to give **9** as yellow oil (0.8 g, 68%, α/β = 3: 1). The α/β ratio was determined by $^1$H NMR analysis. Only the α-anomer was used for glycosylation. α-anomer: $^1$H NMR (600 MHz, CDCl3) $\delta$ 7.86 – 7.42 (m, 7H, Ar), 5.38 (s, 1H, N<u>H</u>), 4.78 (m, 2H, C<u>H</u>$_2$Nap), 4.53 (m, 1H, H-8), 4.35 (dd, $J_{5,6}$ = 10.2 Hz, $J_{6,7}$ = 2.0 Hz, 1H, H-6), 4.12 (m, 1H, H-9a), 4.11 (m, 1H, H-7), 4.06 (m, 5H, H-9b, 2xOC<u>H</u>$_2$$^{Bu}$), 4.02 (dd, $J_{4,5}$ = 10.4 Hz, $J_{3e,4}$ = 3.6 Hz, 1H, H-4), 3.80 (s, 3H, OC<u>H</u>$_3$), 3.68 (t, $J_{4,5}$ = $J_{5,6}$ = 10.5 Hz, 1H, H-5), 2.97 (dd, $J_{3a,3e}$ = 11.9 Hz, $J_{3e,4}$ = 3.6 Hz, 1H, H-3eq), 2.34 (t, $J_{3a,3e}$ = $J_{3a,4}$ = 12.0 Hz, 1H, H-3ax), 1.74 –1.60 (m, 4H, -C<u>H</u>$_2$CH$_2$$^{Bu}$), 1.50 (s, 3H, CC<u>H</u>$_3$), 1.45 – 1.36 (m, 4H, C<u>H</u>$_2$CH$_3$$^{Bu}$), 1.36 (s, 3H, CC<u>H</u>$_3$), 0.94 (t, $J$ = 7.4 Hz, 3H, C<u>H</u>$_3$$^{Bu}$), 0.91 (t, $J$ = 7.4 Hz, 3H, CH$_3$$^{Bu}$); $^{13}$C NMR (150 MHz, CDCl3) $\delta$ 167.90 (C-1), 135.63, 133.27, 133.00, 128.06, 127.92, 127.66, 126.97, 126.16, 125.99, 125.82, 109.49, 99.32 (C-2), 76.20(C-6, C-8), 76.14 (C-4), 75.31 (C-7), 73.50 (<u>C</u>H$_2$Nap), 68.62 (C-9), 68.29 (O<u>C</u>H$_2$$^{Bu}$), 67.81 (O<u>C</u>H$_2$$^{Bu}$), 57.79 (C-5), 53.22 (O<u>C</u>H$_3$), 38.24 (C-3), 32.13 (-<u>C</u>H$_2$CH$_2$$^{Bu}$), 32.06 (-<u>C</u>H$_2$CH$_2$$^{Bu}$), 26.38 (C<u>C</u>H$_3$), 25.07

($C\underline{C}H_3$), 18.62($\underline{C}H_2CH_3^{Bu}$), 18.60 ($\underline{C}H_2CH_3^{Bu}$), 13.59 ($\underline{C}H_3^{Bu}$), 13.57 ($\underline{C}H_3^{Bu}$); HRMS (ESI): *m/z* calculated for $C_{33}H_{47}NO_{12}P$ ($[M+H]^+$) 680.2830, found 680.2844.

**(3aR,4R,6S,7aS)-Methyl 4-((4S,5R)-5-(benzyloxymethyl)-2,2-dimethyl-1,3-dioxolan-4-yl)-6-(dibutoxyphosphoryloxy)-2-oxohexahydro-2H-pyrano[3,4-d] oxazole-6-carboxylate (10)**

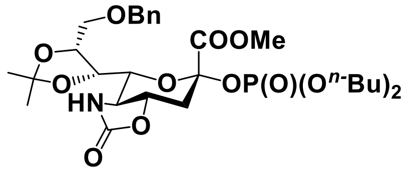


A mixture of thiosialoside donor **S19** (1.0 g, 1.8 mmol), dibutyl phosphate (1.2 g, 5.5 mmol), and pulverized activated 4Å MS in $CH_2Cl_2$ (20 mL) was stirred under argon at RT for 1 h. The mixture was cooled to 0 °C, followed by the addition of *N*-Iodosuccinimide (0.8 g, 3.7 mmol) and 0.1 M TfOH solution in THF (0.9 mL, 0.09 mmol). After stirring for 4 h, the reaction mixture was neutralized with triethylamine and diluted with dichloromethane and filtered through a pad of celite. The filtrate was poured into a mixture of 5% aq. $Na_2S_2O_3$ and saturated aq. $NaHCO_3$. The aqueous layer was extracted with $CH_2Cl_2$. The collected organic phases were washed with saturated brine, dried over $Na_2SO_4$, filtered, and concentrated *in vacuo*. The residue was purified by silica gel column chromatography (toluene:ethyl acetate = 10:1 to 4:1) to give **10** as an oil (0.9 g, 78%, α/β = 4: 1). The α/β ratio was determined by $^1$H NMR analysis. Only the α-anomer was used for glycosylation. α-anomer: $^1$H NMR (400 MHz, $CDCl_3$) $\delta$ 7.41 – 7.26 (m, 5H, Ar), 5.27 (s, 1H, N$\underline{H}$), 4.62 (s, 2H, $PhC\underline{H}_2$), 4.50 (m, 1H, H-8), 4.36 (dd, $J_{5,6}$ = 10.1 Hz, $J_{6,7}$ = 2.0 Hz, 1H, H-6), 4.09 (m, 4H, 2x$OC\underline{H}_2^{Bu}$), 4.11 (m, 1H, H-7), 4.02 (m, 1H, H-4), 4.01 (m, 2H, H-9a, H-9b), 3.83 (s, 3H, $OC\underline{H}_3$), 3.68 (t, $J_{4,5} = J_{5,6}$ = 10.5 Hz, 1H, H-5), 2.98 (dd, $J_{3a,3e}$ = 11.9 Hz, $J_{3e,4}$ = 3.7 Hz, 1H, H-3eq), 2.35 (t, $J_{3a,3e} = J_{3a,4}$ = 12.0 Hz, 1H, H-3ax), 1.71 – 1.59 (m, 4H, -$C\underline{H}_2CH_2^{Bu}$), 1.50 (s, 3H, $CC\underline{H}_3$), 1.46 – 1.38 (m, 4H, $C\underline{H}_2CH_3^{Bu}$), 1.35 (s, 3H, $CC\underline{H}_3$), 0.94 (m, 6H, $C\underline{H}_3^{Bu}$); $^{13}$C NMR (100 MHz, $CDCl_3$) $\delta$ 167.88 (C-1), 159.42, 138.11, 128.55, 128.44, 128.30, 128.09, 128.01, 127.62, 109.45, 99.28(C-2), 76.21 (C-4), 76.09 (C-6, C-8), 75.33 (C-7), 73.38 ($\underline{C}H_2Ph$), 68.48 (C-9), 68.24 ($O\underline{C}H_2^{Bu}$), 67.78 ($O\underline{C}H_2^{Bu}$), 57.76 (C-5), 53.18 ($O\underline{C}H_3$), 38.20 (C-3), 32.09 (-$\underline{C}H_2CH_2^{Bu}$), 32.02 (-$\underline{C}H_2CH_2^{Bu}$), 26.32 ($C\underline{C}H_3$), 25.04 ($C\underline{C}H_3$), 18.57 ($\underline{C}H_2CH_3^{Bu}$), 18.57 ($\underline{C}H_2CH_3^{Bu}$), 13.55 ($\underline{C}H_3^{Bu}$), 13.52 ($\underline{C}H_3^{Bu}$); HRMS (ESI): *m/z* calculated for $C_{29}H_{45}NO_{12}P$ ($[M+H]^+$) 630.2674, found 630.2687.

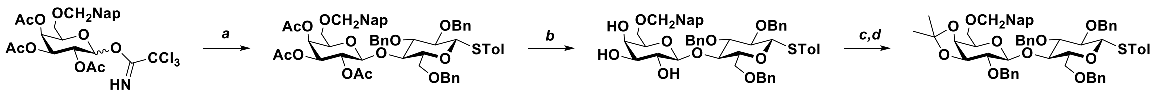


***Scheme S4.*** Synthetic route of intermediate tetrasaccharides fluoride **11**. Reagents and conditions: *a*) **S10**, TMSOTf, $CH_2Cl_2$. *b*) NaOMe, MeOH. *c*) 2,2-dimethoxypropane, camphorsulfonic acid, $CH_3CN$. *d*) BnBr, NaH, DMF, 51% over 4 steps.

**Tolyl [2-*O*-benzyl-3,4-*O*-isopropylidene-6-*O*-(2-naphthylmethyl)-β-D-galactopyranosyl-(1→4)]-2,3,6-tri-*O*-benzyl-β-D-glucopyranoside (11)**

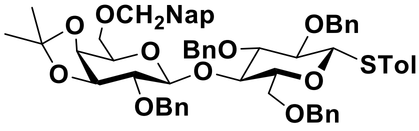


A mixture of glycosyl donor **S6** (6.2 g, 10.6 mmol), acceptor **S10** (4.9 g, 8.8 mmol), and pulverized activated 4Å MS in $CH_2Cl_2$ (100 mL) was stirred at RT under argon for 1 h. The mixture was then cooled to −45 °C, at which temperature TMSOTf (1.9 mL, 10.6 mmol) was added slowly. The reaction was allowed to stir at this temperature for 1 h, followed by quenching with triethylamine. The mixture was filtered through Celite, and the filtrate was diluted with $CH_2Cl_2$, washed with saturated aq. $NaHCO_3$ and dried over $Na_2SO_4$. After filtration, the solvent was removed *in vacuo*. The crude material **S20** was used for the next step without purification. To a solution of the disaccharide **S20** in methanol (50 mL) was added a catalytic amount of sodium methoxide (30% w/v in methanol). The reaction was stirred at RT for 2 h. When TLC indicated completion, Amberlite IR 120 $H^+$ resin was added with vigorous stirring. The resin was then removed by filtration, and the solution was concentrated *in vacuo* to give the de-*O*-acetylated product **S21**. The de-*O*-acetylated disaccharide **S21** was dissolved in acetone, 2,2-dimethoxypropane (50 mL), to which camphorsulfonic acid (0.5 g, 2.0 mmol) were added. The reaction was stirred at RT for 1 h and was then quenched with triethylamine. The solution was concentrated *in vacuo*. The residue was taken up in $CH_2Cl_2$ and washed with sat. $NaHCO_3$. The organic phase was dried over $Na_2SO_4$, followed by filtration and evaporation of $CH_2Cl_2$ to give a crude material for the next step. The isopropylidene protected compound was dissolved in DMF (20 mL), followed by addition of benzyl bromide (3.4g, 20.0 mmol), and was cooled to 0 °C, at which temperature 60% sodium hydride (0.6 g, 16.0 mmol) was added. The mixture was then allowed to warm to RT and stirred for 2 h. Upon completion, excess NaH was quenched with acetic acid. The mixture was concentrated *in vacuo*, the residue taken up in $CH_2Cl_2$ and washed with saturated aq. $NaHCO_3$. After filtration, the solvent was removed *in vacuo*. The crude material was purified by silica gel column chromatography (petroleum: ethyl acetate = 4: 1) to give **11** (4.4 g, 51% over 4 steps). $^1$H NMR (400 MHz, $CDCl_3$) $\delta$ 7.77 – 6.97 (m, 31H, Ar), 4.98 (d, $J$ = 10.3 Hz, 1H, C$\underline{H}_2$Nap), 4.91 – 4.71 (m, 6H, C$\underline{H}_2$Nap, C$\underline{H}_2$Ph), 4.58 (d, $J_{Glc1,2}$ = 9.8 Hz, 1H, H-1$_{Glc}$), 4.57 (d, $J$ = 12.0 Hz, 1H, C$\underline{H}_2$Ph), 4.45 (d, $J_{Gal1,2}$ = 3.0 Hz, 1H, H-1$_{Gal}$), 4.44 (m, 2H, C$\underline{H}_2$Ph), 4.14 (dd, $J_{Gal3,4}$ = 5.6 Hz, $J_{Gal4,5}$ = 1.6 Hz, 1H, H-4$_{Gal}$), 4.07 (dd, $J_{Gal2,3}$ = 6.7 Hz, $J_{Gal3,4}$ = 5.6 Hz,1H, H-3$_{Gal}$), 4.01 (t, $J_{Glc2,3}$ = $J_{Glc3,4}$ = 9.4 Hz, 1H, H-3$_{Glc}$), 3.85 (dd, $J_{Glc6a,6b}$ = 11.1 Hz, $J_{Glc5,6a}$ =4.1 Hz, 1H, H-6a$_{Glc}$), 3.77 (dd, $J_{Glc6a,6b}$ = 11.2 Hz, $J_{Glc5,6b}$ = 1.9 Hz, 1H, H-6b$_{Glc}$), 3.74 – 3.68 (m, 2H, H-6a$_{Gal}$, H-5$_{Gal}$), 3.62 (d, $J_{Glc3,4}$ = 8.9 Hz, 1H, H-4$_{Glc}$), 3.58 (m, 1H, H-6b$_{Gal}$), 3.41 (m, 1H, H-2$_{Glc}$), 3.36 (d, $J_{Gal2,3}$ = 6.7 Hz, 1H, H-2$_{Gal}$), 3.29 (m, 3H, H-2$_{Glc}$, H-5$_{Glc}$, H-2$_{Gal}$), 2.25 (s, 3H, C$\underline{H}_3$Ph), 1.38 (s, 3H, C$\underline{CH}_3$), 1.36 (s, 3H, CC$\underline{H}_3$); $^{13}$C NMR (100 MHz, $CDCl_3$) $\delta$ 138.80, 138.50, 138.47, 138.41, 138.39, 137.70, 136.11, 133.34, 133.01, 132.83, 132.07, 129.77, 129.75, 129.68, 128.57, 128.49, 128.42, 128.39, 128.38, 128.29, 128.25, 128.22, 128.15, 128.07, 128.05, 128.02, 127.98, 127.96, 127.91, 127.80, 127.75, 127.67, 127.58, 127.55, 127.47, 126.16, 126.08, 125.80, 125.74, 109.86 , 101.96 (C-1$_{Gal}$), 87.77 (C-1$_{Glc}$), 85.12 (C-4$_{Glc}$), 80.65 (C-2$_{Gal}$), 80.28(C-5$_{Glc}$), 79.45(C-2$_{Glc}$), 79.40 (C-3$_{Gal}$), 77.36, 76.08 (C-3$_{Glc}$), 75.77 ($\underline{C}$H$_2$Nap), 75.50 ($\underline{C}$H$_2$Ph), 73.70 (C-4$_{Gal}$), 73.56 ($\underline{C}$H$_2$Ph), 73.38 ($\underline{C}$H$_2$Ph), 73.22 ($\underline{C}$H$_2$Ph), 72.14 (C-5$_{Gal}$), 68.94 (C-6$_{Gal}$), 68.38 (C-6$_{Glc}$), 28.06 (C$\underline{C}$H$_3$), 26.54 (C$\underline{C}$H$_3$), 21.16 ($\underline{C}$H$_3$Ph); HRMS (ESI): $m/z$ calculated for $C_{61}H_{64}O_{10}SNa$ ([M+NH$_4$]$^+$) 1011.4112, found 1011.4120.

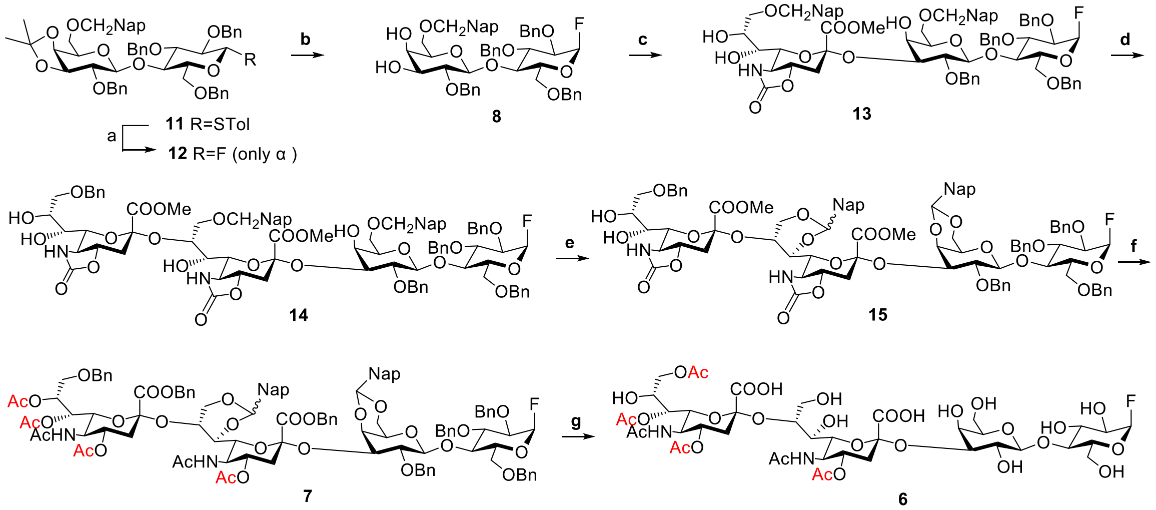

***Scheme S5.*** Synthetic route of intermediate tetrasaccharides fluoride **6**. See also in main text for details. Reagents and conditions: (a) Barluenga reagent, $CH_2Cl_2$, 0 to 21 °C. (b) wet TFA, $CH_2Cl_2$, 21 °C, 59% over 2 steps. (c) i **9**, TMSOTf, −78 °C; ii wet TFA, $CH_2Cl_2$, 21 °C, 82% over 2 steps. (d) i **10**, TMSOTf, −78 °C;

ii wet TFA, $CH_2Cl_2$, 21 °C, 83% over 2 steps. (e) DDQ, $CH_2Cl_2$, 21 °C, 50% (f) i KOH, Dioxane/$H_2O$ (1: 1), 50 °C; ii $Ac_2O$, $CH_2Cl_2$, 21 °C; iii BnBr, $K_2CO_3$, 0 to 21 °C; iv $Ac_2O$, pyridine, DMAP, 0 to 21 °C, 32% over 4 steps (g) $Pd(OH)_2$, $H_2$, Dioxane/$H_2O$, 21 °C, 63%.

**[2-*O*-Benzyl-3,4-*O*-isopropylidene-6-*O*-(2-naphthylmethyl)-β-D-galactopyranosyl-(1→4)]-2,3,6-tri-*O*-benzyl-β-D-glucopyranosyl fluoride (12)**

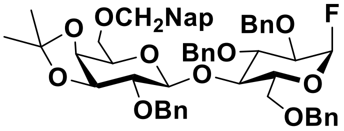


To a solution of **11** (2.0 g, 2.0 mmol) in $CH_2Cl_2$ (30 mL) was added bis(pyridine)iodonium(I) tetrafluoroborate ($IPy_2BF_4$) (2.3 g, 6.2 mmol) under argon and cooled at 0 ℃. After 2 h, the reaction mixture was diluted with $CH_2Cl_2$ and washed with 10% aq. $Na_2S_2O_3$, saturated aq. $NaHCO_3$ and brine. The organic layer was dried ($Na_2SO_4$), filtered and the filtrate concentrated *in vac*uo. The residue was purified by silica gel column chromatography (petroleum: ethyl acetate 4: 1) to give **12** (1.3 g, 70%). $^1$H NMR (400 MHz, $CDCl_3$) $\delta$ 7.85 – 7.12 (m, 27H, Ar), 5.52 (d, $J_{Glc1,2}$ = 2.8 Hz, 0.5H, H-1$_{Glc}$), 5.39 (d, $J_{Glc1,2}$ = 2.8 Hz, 0.5H, H-1$_{Glc}$), 4.98 (d, $J$ = 10.4 Hz, 1H, $C\underline{H}_2$Nap), 4.78 (d, $J$ = 11.7 Hz, 1H, $C\underline{H}_2$Ph), 4.77 (d, $J$ = 11.8 Hz, 1H, $C\underline{H}_2$Ph), 4.72 (d, $J$ = 10.0 Hz, 1H, $C\underline{H}_2$Nap), 4.66 (d, $J$ = 11.6 Hz, 2H, $C\underline{H}_2$Ph), 4.58 (dd, $J$ = 11.9 Hz, 1H, $C\underline{H}_2$Ph), 4.56 (d, $J$ = 12.1 Hz, 1H, $C\underline{H}_2$Ph), 4.45 (d, $J$ = 12.3 Hz, 1H, $C\underline{H}_2$Ph), 4.36 (d, $J$ = 12.1 Hz, 1H, $C\underline{H}_2$Ph), 4.30 (d, $J_{Gal1,2}$ = 8.1 Hz, 1H, H-1$_{Gal}$), 4.10 (dd, $J_{Gal3,4}$ = 5.6 Hz, $J_{Gal4,5}$ = 1.8 Hz, 1H, H-4$_{Gal}$), 4.04 (d, $J_{Glc3,4}$ = 9.6 Hz, 1H, H-4$_{Glc}$), 3.98 (dd, $J_{Gal2,3}$ = 6.8 Hz, $J_{Gal3,4}$ = 5.6 Hz, 1H, H-3$_{Gal}$), 3.89 (m, 1H, H-6a$_{Glc}$), 3.86 (m, 1H, H-5$_{Glc}$), 3.84 (t, $J_{Glc2,3}$ = $J_{Glc3,4}$ = 9.4 Hz, 1H, H-3$_{Glc}$), 3.72 (m, 1H, H-6a$_{Gal}$), 3.69 (m, 1H, H-5$_{Gal}$), 3.58 (m, 1H, H-6b$_{Gal}$), 3.56 (m, 1H, H-6b$_{Glc}$), 3.42 (dd, $J_{Glc2,3}$ = 9.5 Hz, $J_{Glc1,2}$ = 2.9 Hz, 0.5H, H-2$_{Glc}$), 3.36 (m, 0.5H, H-2$_{Glc}$), 3.35 (m, 1H, H-2$_{Gal}$), 1.40 (s, 3H, $C\underline{H}_3$), 1.36 (s, 3H, $CC\underline{H}_3$); $^{13}$C NMR (100 MHz, $CDCl_3$) $\delta$ 138.81, 138.29, 137.87, 137.72, 136.14, 133.26, 132.90, 132.72, 129.58, 128.53, 128.47, 128.41, 128.40, 128.33, 128.27, 128.15, 128.03, 128.00, 127.96, 127.94, 127.92, 127.88, 127.85, 127.82, 127.72, 127.64, 127.58, 127.52, 127.38, 126.01, 125.97, 125.71, 125.64, 109.84, 106.90 (C-1$_{Glc}$), 104.66 (C-1$_{Glc}$), 101.87 (C-1$_{Gal}$), 80.48 (C-2$_{Gal}$), 79.71 (C-3$_{Glc}$), 79.36 (C-3$_{Gal}$), 78.58 (C-2$_{Glc}$), 78.33 (C-2$_{Glc}$), 77.30, 76.99, 76.67, 75.73 ($\underline{C}H_2$Nap), 75.19 (C-4$_{Glc}$), 73.82 ($\underline{C}H_2$Ph), 73.67 (C-4$_{Gal}$), 73.57 ($\underline{C}H_2$Ph), 73.38 ($\underline{C}H_2$Ph), 73.22 ($\underline{C}H_2$Ph), 72.65 (C-5$_{Glc}$), 72.33 (C-5$_{Gal}$), 69.00 (C-6$_{Gal}$), 67.21 (C-6$_{Glc}$), 27.96 ($C\underline{C}H_3$), 26.44 ($C\underline{C}H_3$); $^{19}$F NMR (376 MHz, $CDCl_3$) $\delta$ -149.62; HRMS (ESI): *m/z* calculated for $C_{54}H_{61}FNO_{10}$ ($[M+NH_4]^+$) 902.4274, found 902.4272.

**[2-*O*-Benzyl-3,4-*O*-isopropylidene-6-*O*-(2-naphthylmethyl)-β-D-galactopyranosyl-(1→4)]-2,3,6-tri-*O*-benzyl-β-D-glucopyranosyl fluoride (8)**

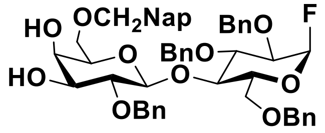


To a solution of compound **12** (1.2 g, 1.4 mmol) in $CH_2Cl_2$ and water (20 mL, 19:1) was added trifluoroacetic acid (2 mL, 10% v/v). The biphasic mixture was vigorously stirred at RT for 1 h and then poured into cold saturated aq. $NaHCO_3$. The mixture was gently shaken in a separatory funnel until no gas was produced. The organic phase was collected and dried over $Na_2SO_4$. The solvent was removed *in vacuo* after filtration. The residue was purified by silica gel column chromatography (petroleum ether:ethyl acetate 3:1 to 1:1) to give the **8** (1.0 g, 84%). $^1$H NMR (400 MHz, $CDCl_3$) $\delta$ 7.86 – 7.11 (m, 27H, Ar), 5.54 (d, $J_{Glc1,2}$ = 2.8 Hz, 0.5H, H-1$_{Glc}$), 5.41 (d, $J_{Glc1,2}$ = 2.8 Hz, 0.5H, H-1$_{Glc}$), 5.03 (d, $J$ = 10.9 Hz, 1H, C<u>H</u>$_2$Nap), 4.78 (d, $J$ = 11.9 Hz, 1H, C<u>H</u>$_2$Ph), 4.77 (d, $J$ = 11.9 Hz, 1H, C<u>H</u>$_2$Ph), 4.76 (d, $J$ = 10.8 Hz, 1H, C<u>H</u>$_2$Nap),4.81 – 4.74 (m, 2H, C<u>H</u>$_2$Ph), 4.68 – 4.51 (m, 4H, C<u>H</u>$_2$Ph), 4.56 (d, $J$ = 12.1 Hz, 1H, C<u>H</u>$_2$Ph), 4.37 (d, $J$ = 12.1 Hz, 1H, C<u>H</u>$_2$Ph), 4.33 (d, $J_{Gal1,2}$ = 7.1 Hz, 1H, H-1$_{Gal}$), 4.06 (t, $J_{Glc3,4}$ = $J_{Glc4,5}$ = 9.7 Hz, 1H, H-4$_{Glc}$), 3.95 (d, $J_{Gal3,4}$ = 3.3 Hz, 1H, , H-3$_{Gal}$), 3.87 (m, 1H, H-3$_{Glc}$), 3.87 – 3.82 (m, 2H, H-6a$_{Glc}$, H-5$_{Glc}$), 3.66 (dd, $J_{Gal6a,6b}$ = 10.0 Hz, $J_{Gal5,6a}$ = 6.5 Hz, 1H, H-6a$_{Gal}$), 3.61 – 3.51 (m, 2H, H-6b$_{Glc}$, H-6b$_{Gal}$), 3.49 (dd, $J_{Glc2,3}$ = 9.5 Hz, $J_{Glc1,2}$ = 2.8 Hz, 0.5H, H-2$_{Glc}$), 3.43 (m, 0.5H, H-2$_{Glc}$), 3.40 (m, 1H, H-2$_{Gal}$), 3.40 – 3.33 (m, 2H, H-4$_{Gal}$, H-5$_{Gal}$); $^{13}$C NMR (100 MHz, $CDCl_3$) $\delta$ 138.96, 138.20, 137.84, 137.66, 135.50, 133.22, 132.97, 128.50, 128.43, 128.39, 128.19, 128.07, 128.05, 127.91, 127.90, 127.86, 127.83, 127.73, 127.67, 127.32, 126.34, 126.10, 125.88, 125.55, 106.83 (C-1$_{Glc}$), 104.58 (C-1$_{Glc}$), 102.61 (C-1$_{Gal}$), 79.87 (C-2$_{Gal}$), 79.57 (C-3$_{Glc}$), 78.58 (C-2$_{Glc}$), 78.33 (C-2$_{Glc}$), 75.53 (<u>C</u>H$_2$Nap), 75.41 (C-4$_{Glc}$), 74.95 (<u>C</u>H$_2$Ph), 73.79 (<u>C</u>H$_2$Ph), 73.63 (<u>C</u>H$_2$Ph), 73.43 (<u>C</u>H$_2$Ph), 73.24 (C-4$_{Gal}$, C-5$_{Gal}$), 73.12, 72.59 (C-5$_{Glc}$), 68.71(C-3$_{Gal}$), 68.66 (C-6$_{Gal}$), 67.28 (C-6$_{Glc}$); $^{19}$F NMR (376 MHz, $CDCl_3$) $\delta$ -149.84; HRMS (ESI): *m/z* calculated for $C_{61}H_{68}NO_{10}S$ ($[M+NH_4]^+$) 862.3961, found 862.3960.

**(Methyl 5-acetamido-9-*O*-(2-naphthylmethyl)-5-*N*,4-*O*-carbonyl-3,5-dideoxy-D-glycero-α-D-galacto-non-2-ulopyranosylonate)-(2→3)-[2-O-benzyl-6-*O*-(2-naphthylmethyl)-β-D-galactopyranosyl-(1→4)]-2,3,6-tri-*O*-benzyl-α-D-glucopyranosyl fluoride (13)**

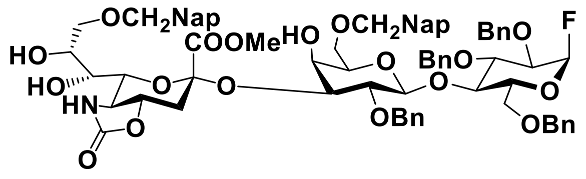


A mixture of diol acceptor **8** (0.9 g, 1.1 mmol), sialyl donor **9** (2.0 g, 2.1 mmol), and pulverized activated 4Å MS in $CH_2Cl_2$ (50 mL) was stirred at RT under argon for 1 h. The mixture was then cooled to −78 °C, at which temperature TMSOTf (350 µL, 2.1 mmol) was added dropwise. The reaction was stirred at this temperature for 10 min and then quenched with triethylamine. The mixture was filtered through Celite, diluted with $CH_2Cl_2$ and washed with saturated aq. $NaHCO_3$. The organic phase was dried over $Na_2SO_4$, filtered and concentrated *in vacuo*. The residue was dissolved in $CH_2Cl_2/H_2O$ (19/1, 20 mL), trifluoroacetic acid (2 mL, 10%V) was added dropwise into the solution at 0 °C. After stirring for 1 h at RT, the reaction mixture was diluted with $CH_2Cl_2$ and washed with saturated aq. $NaHCO_3$ and brine, dried over $MgSO_4$, filtered, and the solvent evaporated *in vacuo*. The residue was purified by silica gel column chromatography (toluene:ethyl acetate 4:1) to give the alpha isomer **13** (1.1 g, 82%). $^1$H NMR (600 MHz, $CDCl_3$) $\delta$ 7.83 – 7.10 (m, 34H, Ar), 5.65 (s, 1H, N$\underline{H}$), 5.51 (d, $J_{Glc1,2}$ = 2.8 Hz, 0.5H, H-1$_{Glc}$), 5.42 (d, $J_{Glc1,2}$ = 2.8 Hz, 0.5H, H-1$_{Glc}$), 4.98 (d, $J$ = 10.6 Hz, 1H, C$\underline{H}_2$Nap), 4.79 (d, $J$ = 11.8 Hz, 1H, C$\underline{H}_2$Ph), 4.73 (d, $J$ = 10.6 Hz, 1H, C$\underline{H}_2$Nap), 4.66 (m, 2H, C$\underline{H}_2$Nap), 4.62 (d, $J$ = 11.7 Hz, 1H, C$\underline{H}_2$Ph), 4.60 (m, 3H, C$\underline{H}_2$Ph), 4.52 (d, $J$ = 12.1 Hz, 1H, C$\underline{H}_2$Ph), 4.50 (d, $J$ = 12.2 Hz, 1H, C$\underline{H}_2$Ph), 4.38 (d, $J$ = 12.1 Hz, 1H, C$\underline{H}_2$Ph), 4.32 (d, $J_{Gal1,2}$ = 7.8 Hz, 1H, H-1$_{Gal}$), 4.03 (t, $J_{Glc3,4}$ = $J_{Glc4,5}$ = 9.5 Hz, 1H, H-4$_{Glc}$), 3.90 (dd, $J_{Sia4,5}$ = 10.0 Hz, $J_{Sia3e,4}$ = 4.3 Hz, 1H, H-4$_{Sia}$), 3.88 (m, 1H, H-3$_{Glc}$), 3.85 – 3.76 (m, 5H, H-6$_{Sia}$, H-3$_{Gal}$, H-6a$_{Glc}$, H-5$_{Glc}$, H-8$_{Sia}$), 3.71 (m, 1H, H-7$_{Sia}$), 3.70 (s, 3H, OC$\underline{H}_3$), 3.67 (dd, $J_{Sia9a,9b}$ = 9.6 Hz, $J_{Sia8,9a}$ = 4.8 Hz, 1H, H-9a$_{Sia}$), 3.64 (dd, $J_{Gal6a,6b}$ = 9.9 Hz, $J_{Gal5,6a}$ = 6.6 Hz, 1H, H-6a$_{Gal}$), 3.60 (d, $J_{Glc6a,6b}$ = 10.4 Hz, 1H, H-6b$_{Glc}$), 3.57 (m, 1H, H-9b$_{Sia}$), 3.54 (d, $J_{Gal1,2}$ = 7.7 Hz, 1H, H-2$_{Gal}$), 3.51 (dd, $J_{Gal6a,6b}$ = 9.9 Hz, $J_{Gal5,6b}$ = 6.0 Hz, 1H, H-6b$_{Gal}$), 3.46 – 3.40 (m, 2H, H-2$_{Glc}$, H-4$_{Gal}$), 3.37 (t, $J_{Sia4,5}$ = $J_{Sia5,6}$ = 10.5 Hz, 1H, H-5$_{Sia}$), 3.24 (t, $J_{Gal5,6a}$ = $J_{Gal5,6b}$ = 6.3 Hz, 1H, H-5$_{Gal}$), 2.94 (dd, $J_{Sia3a,3e}$ = 12.1 Hz, $J_{Sia3e,4}$ = 3.7 Hz, 1H, H-3eq$_{Sia}$), 2.10 (t, $J_{Sia3a,3e}$ = $J_{Sia3a,4}$ = 12.7 Hz, 1H, H-3ax$_{Sia}$); $^{13}$C NMR (150 MHz, $CDCl_3$) $\delta$ 168.70 (C-1$_{Sia}$), 159.41, 138.91, 137.89, 135.70, 134.93, 133.28, 133.21, 133.08, 133.06, 132.98, 132.96, 129.07, 129.06, 128.50, 128.48, 128.44, 128.42, 128.30, 128.29, 128.26, 128.24, 128.15, 128.12, 128.10, 128.05, 127.96, 127.94, 127.88, 127.86,

127.82, 127.81, 127.78, 127.76, 127.74, 127.72, 127.42, 127.40, 126.77, 126.75, 126.34, 126.32, 126.20, 126.17, 126.14, 125.95, 125.92, 125.64, 125.54, 125.52, 125.33, 125.31, 106.47 (C-1$_{Glc}$), 104.97 (C-1$_{Glc}$), 102.39 (C-1$_{Gal}$), 99.50 (C-2$_{Sia}$), 79.53 (C-6$_{Sia}$), 78.57 (C-2$_{Glc}$), 78.41 (C-2$_{Glc}$), 78.12 (C-2$_{Gal}$), 77.69 (C-3$_{Glc}$), 77.66 (C-4$_{Sia}$), 76.55 (C-3$_{Gal}$), 75.84 (<u>C</u>H$_2$Nap), 75.68 (<u>C</u>H$_2$Nap), 75.18 (C-4$_{Glc}$), 73.85 (<u>C</u>H$_2$Ph), 73.62 (<u>C</u>H$_2$Ph), 73.56 (<u>C</u>H$_2$Ph), 73.18 (<u>C</u>H$_2$Ph), 72.58 (C-5$_{Glc}$), 72.47 (C-5$_{Gal}$), 72.21(C-4$_{Gal}$), 71.23 (C-9$_{Sia}$), 69.75 (C-8$_{Sia}$), 68.24 (C-6$_{Gal}$), 68.02 (C-7$_{Sia}$), 67.43 (C-6$_{Glc}$), 57.91 (C-5$_{Sia}$), 53.45 (O<u>C</u>H$_3$), 36.19 (C-3$_{Sia}$); $^{19}$F NMR (376 MHz, CDCl$_3$) $\delta$ -149.88; HRMS (ESI): *m/z* calculated for $C_{73}H_{80}FN_2O_{18}$ ([M+NH$_4$]$^+$) 1291.5385, found 1291.5381.

**(Methyl 5-acetamido-9-*O*-benzyl-5-*N*,4-*O*-carbonyl-3,5-dideoxy-D-glycero-α-D-galacto-non-2-ulopyranosylonate)-(2→8)-(Methyl 5-acetamido-9-*O*-(2-naphthylmethyl)-5-*N*,4-*O*-carbonyl-3,5-dideoxy-D-glycero-α-D-galacto-non-2-ulopyranosylonate)-(2→3)-[2-*O*-benzyl-6-*O*-(2-naphthylmethyl)-β-D-galactopyranosyl-(1→4)]-2,3,6-tri-*O*-benzyl-α-D-glucopyranosyl fluoride (14)**

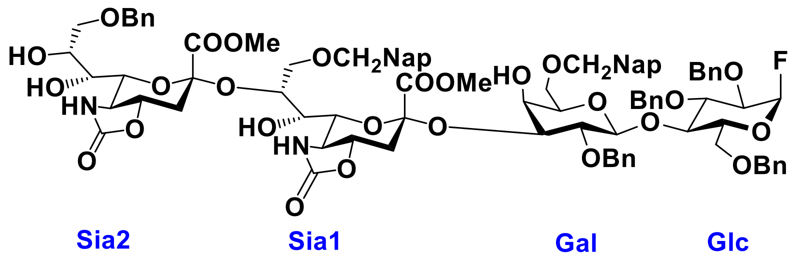


A suspension of acceptor **13** (1.1 g, 0.8 mmol), sialyl donor **10** (1.0 g, 1.6 mmol) and pulverized activated 4Å MS in $CH_2Cl_2$ (25 mL) was stirred at RT under an atmosphere of argon for 1 h. It was then cooled to −78 °C, at which temperature TMSOTf (310 μL, 1.7 mmol) was added dropwise. The reaction mixture was stirred at this temperature for 10 min and was then quenched with triethylamine. The mixture was filtered through Celite, diluted in $CH_2Cl_2$, and washed with saturated aq. $NaHCO_3$. The organic phase was dried over $Na_2SO_4$, filtered and the filtrate and concentrated *in vacuo*. The residue was dissolved in $CH_2Cl_2/H_2O$ (19/1, 20 mL), trifluoroacetic acid (2 mL) was added dropwise into the solution at 0 °C. After stirring for 1 h at RT, the reaction mixture was diluted with $CH_2Cl_2$ and washed with saturated aq. $NaHCO_3$ and brine, dried over $MgSO_4$, filtered, and the filtrate concentrated *in vacuo*. The residue was purified by silica gel column chromatography (toluene:ethyl acetate 4:1) to give the alpha isomer **14** (1.2 g, 83%). $^1$H NMR (600 MHz, CDCl$_3$) $\delta$ 7.86 – 7.09 (m, 39H, aromatic), 6.30 (s, 1H, NH$_{Sia2}$), 5.91 (s, 1H, NH$_{Sia1}$), 5.51 (d, $J_{Glc1,2}$ = 2.8 Hz, 0.5H, H-1$_{Glc}$), 5.42 (d, $J_{Glc1,2}$ = 2.8 Hz, 0.5H, H-1$_{Glc}$), 4.98 (d, *J*

= 10.7 Hz, 1H, C$\underline{H}_2$Nap), 4.78 (d, $J$ = 11.8 Hz, 1H, C$\underline{H}_2$Ph), 4.71 (d, $J$ = 10.6 Hz, 1H, C$\underline{H}_2$Nap), 4.63 (m, 2H, C$\underline{H}_2$Nap), 4.61 (d, $J$ = 11.7 Hz, 1H, C$\underline{H}_2$Ph), 4.59 – 4.48 (m, 6H, C$\underline{H}_2$Ph), 4.47 (d, $J$ = 11.5 Hz, 1H, C$\underline{H}_2$Ph), 4.31 (d, $J$ = 11.9 Hz, 1H, C$\underline{H}_2$Ph), 4.32 (m, 1H, H-1$_{Gal}$), 4.11 (m, 1H, H-8$_{Sia1}$), 4.02 (t, $J_{Glc3,4}$ = $J_{Glc4,5}$ = 9.6 Hz, 1H, H-4$_{Glc}$), 3.97 – 3.85 (m, 4H, H-4$_{Sia1}$, H-4$_{Sia2}$, H-6$_{Sia1}$, H-3$_{Glc}$), 3.81 (m, 2H, H-8$_{Sia2}$, H-6$_{Sia2}$), 3.80 (d, $J_{Glc6a,6b}$ = 10.9 Hz, 1H, H-6a$_{Glc}$), 3.78 (m, 1H, H-5$_{Glc}$), 3.75 (m, 2H, H-7$_{Sia2}$, H-3$_{Gal}$), 3.72 (m, 1H, H-9a$_{Sia1}$), 3.69 (s, 3H, OC$\underline{H}_3$), 3.68 – 3.61 (m, 4H, H-6a$_{Gal}$, H-9b$_{Sia1}$, H-9a$_{Sia2}$, H-7$_{Sia1}$), 3.62 (s, 3H, OC$\underline{H}_3$), 3.59 (m, 1H, H-9b$_{Sia2}$), 3.55 (d, $J_{Glc6a,6b}$ = 10.9 Hz, 1H, H-6b$_{Glc}$), 3.53 (m, 1H, H-4$_{Gal}$), 3.52 (m, 2H, H-6b$_{Gal}$, H-2$_{Gal}$), 3.44 (d, $J_{Glc2,3}$ = 9.7 Hz, 0.5H, H-2$_{Glc}$), 3.39 (d, $J_{Glc2,3}$ = 9.6 Hz, 0.5H, H-2$_{Glc}$), 3.38 (m, 1H, H-5$_{Sia1}$), 3.29 (m, 1H, H-5$_{Sia2}$), 3.25 (m, 1H, H-5$_{Gal}$), 2.92 (dd, $J_{Sia3a,3e}$ = 12.2 Hz, $J_{Sia3e,4}$ = 3.8 Hz, 1H, H-3eq$_{Sia1}$), 2.81 (dd, $J_{Sia3a,3e}$ = 12.2 Hz, $J_{Sia3e,4}$ = 3.7 Hz, 1H, H-3eq$_{Sia2}$), 2.10 (m, 2H, H-3ax$_{Sia2}$, H-3ax$_{Sia1}$); $^{13}$C NMR (150 MHz, CDCl$_3$) $\delta$ 168.54 (C-1$_{Sia2}$), 168.26 (C-1$_{Sia1}$), 160.25, 159.93, 138.88, 138.03, 137.91, 137.86, 137.64, 135.75, 134.75, 133.21, 133.11, 132.99, 132.91, 128.48, 128.43, 128.40, 128.36, 128.32, 128.29, 128.08, 128.06, 127.96, 127.89, 127.82, 127.80, 127.70, 127.68, 127.37, 126.74, 126.33, 126.14, 125.88, 125.64, 125.54, 106.42 (C-1$_{Glc}$), 104.92 (C-1$_{Glc}$), 102.19 (C-1$_{Gal}$), 100.32 (C-2$_{Sia2}$), 99.77 (C-2$_{Sia1}$), 79.51 (C-6$_{Sia2}$), 78.48 (C-2$_{Glc}$), 78.32 (C-2$_{Glc}$), 78.08 (C-2$_{Gal}$), 77.14 (C-4$_{Sia1}$, C-4$_{Sia2}$, C-6$_{Sia1}$), 76.72 (C-3$_{Gal}$), 75.97 (C-3$_{Glc}$), 75.62 ($\underline{C}$H$_2$Nap), 75.19 (C-8$_{Sia1}$), 75.01 (C-4$_{Glc}$), 73.80 ($\underline{C}$H$_2$Ph), 73.53 ($\underline{C}$H$_2$Ph), 73.20 ($\underline{C}$H$_2$Ph), 72.63 (C-5$_{Glc,}$ C-5$_{Gal}$), 72.49 (C-7$_{Sia1}$), 71.47 (C-9$_{Sia2}$), 71.32 (C-4$_{Gal}$), 69.87 (C-9$_{Sia1}$), 69.72 (C-8$_{Sia2}$), 68.42 (C-6$_{Gal}$), 68.01 (C-7$_{Sia2}$), 67.38 (C-6$_{Glc}$), 59.00 (C-5$_{Sia2}$), 57.24 (C-5$_{Sia1}$), 53.54 (O$\underline{C}$H$_3$), 53.15 (O$\underline{C}$H$_3$), 35.81 (C-3$_{Sia1}$, C-3$_{Sia2}$); $^{19}$F NMR (376 MHz, CDCl$_3$) $\delta$ -149.56; HRMS (ESI): *m/z* calculated for $C_{91}H_{101}FN_3O_{26}$ ($[M+NH_4]^+$) 1670.6652, found 1670.6620.

**(Methyl 5-acetamido-9-*O*-benzyl-5-*N*,4-*O*-carbonyl-3,5-dideoxy-D-glycero-α-D-galacto-non-2-ulopyranosylonate)-(2→8)-(Methyl 5-acetamido-7,9-*O*-(2-naphthylmethylidene)-5-*N*,4-*O*-carbonyl-3,5-dideoxy-D-glycero-α-D-galacto-non-2-ulopyranosylonate)-(2→3)-[2-*O*-benzyl-5,6-*O*-(2-naphthylmethylidene)-β-D-galactopyranosyl-(1→4)]-2,3,6-tri-*O*-benzyl-α-D-glucopyranosyl fluoride (15)**

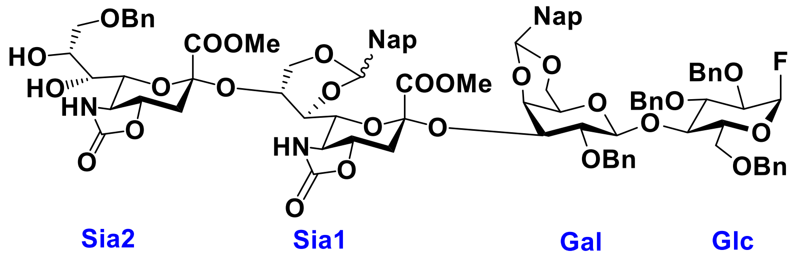


To a suspension of **14** (0.5 g, 0.3 mmol) and pulverized activated 4Å MS in $CH_2Cl_2$ (12 mL) under an atmosphere of argon was added DDQ (0.3 g, 1.2 mmol). The reaction mixture was stirred in the dark for 2 h at RT. Upon completion, the reaction mixture was diluted with $CH_2Cl_2$, and then washed with saturated aq. $NaHCO_3$ until the color of the organic phase turned light yellow. The organic layer was collected, dried over $Na_2SO_4$, filtered and the filtrate concentrated under reduced pressure. The residue was purified by silica gel column chromatography (toluene:ethyl acetate 4:1 to 1:1) to give **15** (248 mg, 50%). HRMS (ESI): *m/z* calculated for $C_{91}H_{97}FN_3O_{26}$ ($[M+NH_4]^+$) 1666.6339, found 1666.6302. *NMR analysis was complicated due to the presence of isomers. For detailed analysis, see compound **7'**.

**(Benzyl 5-acetamido-9-*O*-benzyl-4,7,8-tri-*O*-acetyl-3,5-dideoxy-D-glycero-α-D-galacto-non-2-ulopyranosylonate)-(2→8)-(Benzyl 5-acetamido-4-*O*-acetyl-7,9-*O*-(2-aphthylmethylidene)-3,5-dideoxy-D-glycero-α-D-galacto-non-2-ulopyranosylonate)-(2→3)-[2-*O*-benzyl-7,9-*O*-(2-aphthylmethylidene)-β-D-galactopyranosyl-(1→4)]-2,3,6-tri-*O*-benzyl-α-D-glucopyranosyl fluoride (7)**

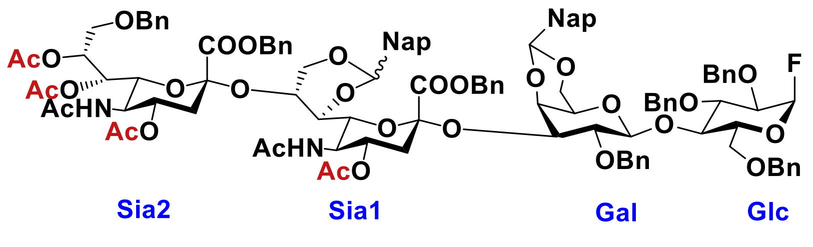


To a stirred solution of **15** (40.0 mg, 0.02 mmol) in 1,4-dioxane (4 mL) and $H_2O$ (4 mL) was added KOH (30.0 mg, 0.5 mmol) at RT. After being stirred at 50 °C for 2 h, the reaction mixture was concentrated *in vacuo*. The residue was dissolved in $CH_2Cl_2$ (2 mL), $Ac_2O$ (10.0 μL, 3.6 mmol) was added dropwise. After stirring the reaction mixture at RT for 2 h, it was concentrated *in vacuo*. The residue was purified by size exclusion chromatography

(LH-20, MeOH:$H_2O$=1:1). The residue was dissolved in $H_2O$ (2 mL) to which was $K_2CO_3$ was added (14.0 mg, 0.1 mmol) to achieve a pH ~ 8. The mixture was then concentrated *in vacuo*, and the residue was dissolved in DMF (2 mL), followed by the addition of BnBr (25.0 µL, 0.2 mmol). After being stirred at the same temperature for 2 h, pyridine (1 mL), $Ac_2O$ (0.5 mL) and a catalytic amount of DMAP (5.0 mg, 0.04 mmol) were added. The reaction mixture was stirred at RT overnight and then was extracted with $CH_2Cl_2$ and washed with 1 N HCl, saturated aq. $NaHCO_3$ and brine. The organic layer was dried over $Na_2SO_4$, filtered and the filtrate concentrated *in vacuo*. The residue was purified by size exclusion chromatography (LH-20, MeOH: $CH_2Cl_2$ = 1: 1) to give **7** (15 mg, 32% over 4 steps). HRMS (ESI): *m/z* calculated for $C_{113}H_{122}FN_3O_{30}$ ($[M+NH_4+H]^{2+}$) 1010.4060, found 1010.4040. *NMR analysis was complicated due to the presence of isomers. For detailed analysis, see compound **7'**.

**(Benzyl 5-acetamido-9-*O*-benzyl-4,7,8-tri-*O*-acetyl-3,5-dideoxy-D-glycero-α-D-galacto-non-2-ulopyranosylonate)-(2→8)-(Benzyl 5-acetamido-4-*O*-acetyl-3,5-dideoxy-D-glycero-α-D-galacto-non-2-ulopyranosylonate)-(2→3)-[2-*O*-benzyl-β-D-galactopyranosyl-(1→4)]-2,3,6-tri-*O*-benzyl-α-D-glucopyranosyl fluoride (7')**

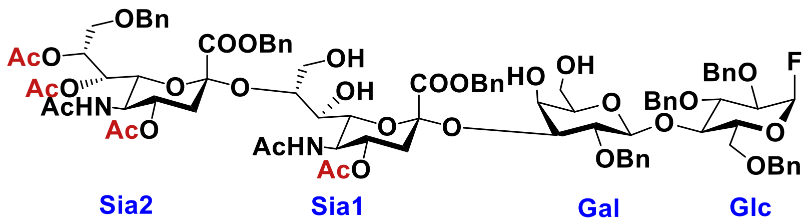


Compound **7** (2.0 mg, 1.0 µmol) was dissolved in $CH_2Cl_2/H_2O$ (19/1, 2 mL), and then trifluoroacetic acid (0.2 mL) was added dropwise at 0 °C. After stirring for 1 h at RT, the reaction mixture was diluted with $CH_2Cl_2$ and washed with saturated aq. $NaHCO_3$ and brine, dried over $Na_2SO_4$, filtered and the filtrate concentrated *in vacuo*. The residue was purified by silica gel column chromatography ($CH_2Cl_2$:MeOH = 50:1 to 30:1) to give **7'** (0.8 mg). $^1H$ NMR (600 MHz, $CDCl_3$) $\delta$ 7.43 – 7.17 (m, 35H, aromatic), 5.53 (d, $J_{Glc1,2}$ = 2.7 Hz, 0.5H, H-$1_{Glc}$), 5.44 (d, $J_{Glc1,2}$ = 2.7 Hz, 0.5H, H-$1_{Glc}$), 5.39 (d, $J_{Sia5,NH}$ = 8.8 Hz, 1H, $NH_{Sia2}$), 5.35 (d, $J_{Sia7,8}$ = 9.3 Hz, 1H, H-$7_{Sia2}$), 5.29 (d, $J$ = 10.4 Hz, 1H, $COOCH_2Ph$), 5.27 – 5.21 (m, 2H, $COOCH_2Ph$, H-$8_{Sia2}$), 5.18 (d, $J$ = 12.2 Hz, 1H, $COOCH_2Ph$), 5.14 (d, $J$ = 12.2 Hz, 1H, $COOCH_2Ph$), 5.05 (d, $J_{Sia5,NH}$ = 9.8 Hz, 1H, $NH_{Sia1}$), 5.02 (dd, $J$ = 10.2 Hz, 1H, $CH_2Ph$), 4.90 (tdd, $J_{Sia3a,4}$ = 11.9 Hz, $J_{Sia4,5}$ =9.4 Hz, $J_{Sia3e,4}$ = 4.6 Hz, 2H, H-$4_{Sia1}$, H-$4_{Sia2}$), 4.82 (d, $J$ = 11.9 Hz, 1H, $CH_2Ph$), 4.72 (d, $J$ = 10.6 Hz, 1H, $CH_2Ph$), 4.66 (d, $J$ = 11.9 Hz, 1H, $CH_2Ph$), 4.61 (d, $J$ = 10.8 Hz,

2H, C$\underline{H}_2$Ph), 4.57 (d, $J$ = 11.9 Hz, 1H, C$\underline{H}_2$Ph), 4.49 (d, $J$ = 11.8 Hz, 1H, C$\underline{H}_2$Ph), 4.38 (d, $J$ = 11.8 Hz, 1H, C$\underline{H}_2$Ph), 4.37 (d, $J$ = 12.0 Hz, 1H, C$\underline{H}_2$Ph), 4.28 (d, $J_{Gal1,2}$ = 7.8 Hz, 1H, H-1$_{Gal}$), 4.13 (dd, $J_{Sia5,6}$ = 10.7 Hz, $J_{Sia6,7}$ = 2.0 Hz, 1H, H-6$_{Sia1}$), 4.09 (m, 1H, H-8$_{Sia1}$), 4.03 (m, 1H, H-5$_{Sia1}$), 4.00 (m, 1H, H-5$_{Sia2}$), 3.95 (d, $J_{Glc4,5}$ = 9.5 Hz, 1H, H-4$_{Glc}$), 3.89 (m, 2H, H-9a$_{Sia1}$, H-9b$_{Sia1}$), 3.84 (m, 1H, H-3$_{Gal}$), 3.82 (m, 1H, H-3$_{Glc}$), 3.80 (m, 1H, H-6$_{Sia2}$), 3.79 (m, 1H, H-6a$_{Glc}$), 3.76 (d, $J_{Glc4,5}$ = 10.6 Hz, 1H, H-5$_{Glc}$), 3.69 (m, 1H, H-7$_{Sia1}$), 3.59 (m, 1H, H-4$_{Gal}$), 3.56 (m, 1H, H-9a$_{Sia2}$), 3.52 (m, 2H, H-6a$_{Gal}$, H-6b$_{Gal}$), 3.49 (m, 1H, H-6b$_{Glc}$), 3.48(m, 2H, H-2$_{Gal}$, H-2$_{Glc}$), 3.40 (dd, $J_{Sia9a,9b}$ = 10.9 Hz, $J_{Sia8,9b}$ = 4.8 Hz, 1H, H-9b$_{Sia2}$), 3.15 (t, $J_{Gal5,6a}$ = $J_{Gal5,6b}$ = 6.0 Hz, 1H, H-5$_{Gal}$), 2.70 (dd, $J_{Sia3a,3e}$ = 12.9 Hz, $J_{Sia3e,4}$ = 4.5 Hz, 1H, H-3eq$_{Sia1}$), 2.39 (dd, $J_{Sia3a,3e}$ = 13.2 Hz, $J_{Sia3e,4}$ = 4.8 Hz, 1H, H-3eq$_{Sia2}$), 2.11 (s, 3H, Ac), 2.06 – 1.94 (m, 14H, 3xAc, H-3ax$_{Sia1}$, H-3ax$_{Sia2}$), 1.86 (s, 3H, Ac), 1.84 (s, 3H, Ac); $^{13}$C NMR (150 MHz, $CDCl_3$) $\delta$ 128.81, 128.49, 128.33, 128.17, 127.84, 106.41 (C-1$_{Glc}$), 104.96 (C-1$_{Glc}$), 102.55 (C-1$_{Gal}$), 79.67 (C-3$_{Glc}$), 78.22 (C-2$_{Glc}$, C-2$_{Gal}$), 76.93 (C-8$_{Sia1}$), 75.80 (C-3$_{Gal}$, $\underline{C}H_2$Ph), 75.64 (C-4$_{Glc}$), 75.32 (C-4$_{Gal}$, $\underline{C}H_2$Ph), 73.87 ($\underline{C}H_2$Ph), 73.71 (C-5$_{Gal}$), 73.38 ($\underline{C}H_2$Ph), 72.74 (C-5$_{Glc}$, C-6$_{Sia1}$), 70.32 (C-7$_{Sia1}$), 68.87 (C-8$_{Sia2}$), 68.75 (C-4$_{Sia1}$, C-4$_{Sia2}$), 68.55 (C-6$_{Sia2}$, C-9$_{Sia2}$), 67.90 (COO$\underline{C}H_2$Ph), 67.74 (C-7$_{Sia2}$, COOC$\underline{H}_2$Ph), 67.10 (C-6$_{Glc}$), 63.07 (C-9$_{Sia1}$), 61.78 (C-6$_{Gal}$), 50.99 (C-5$_{Sia2}$), 49.54 (C-5$_{Sia1}$), 38.26 (C-3$_{Sia1}$), 36.00 (C-3$_{Sia2}$), 23.27 (NCO$\underline{C}H_3$), 23.11(NCO$\underline{C}H_3$), 21.34 (CO$\underline{C}H_3$), 20.86 (CO$\underline{C}H_3$); HRMS (ESI): *m/z* calculated for $C_{91}H_{109}FN_3O_{30}$ ($[M+NH_4]^+$) 1742.7074, found 1742.7088.

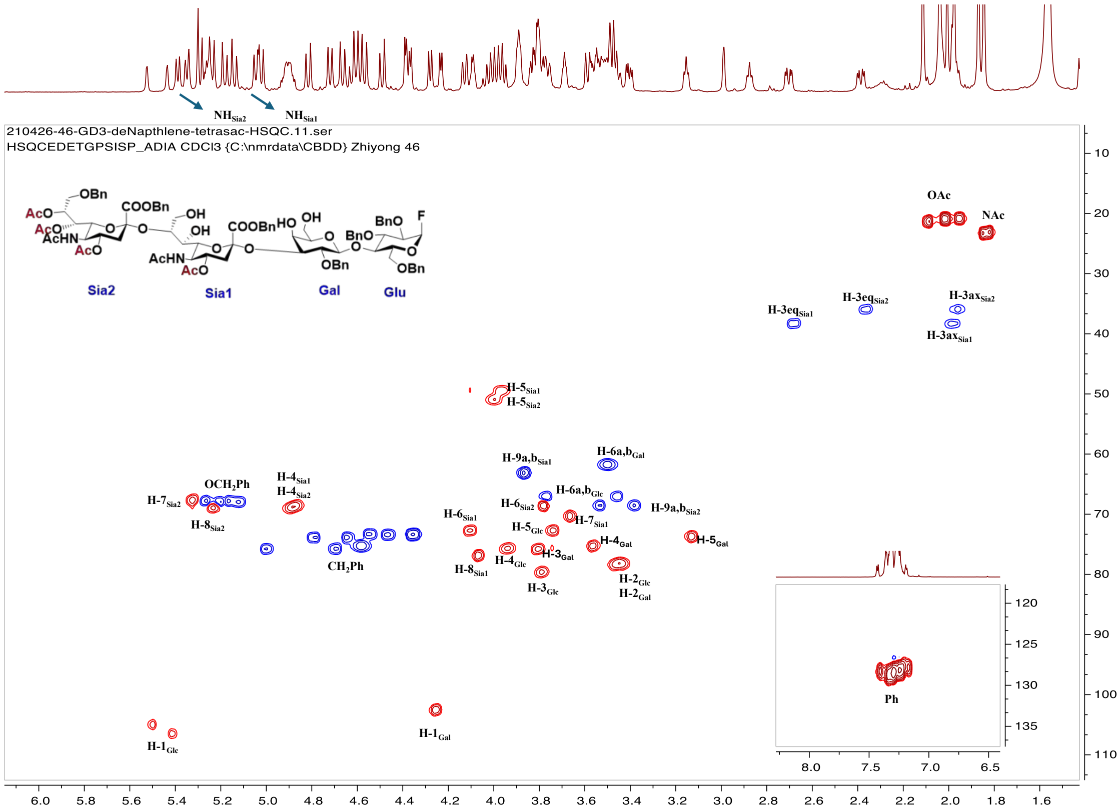


**(5-Acetamido-4,7,9-tri-*O*-acetyl-3,5-dideoxy-D-glycero-α-D-galacto-non-2-ulopyranosylonate)-(2→8)-(5-acetamido-4-*O*-acetyl-3,5-dideoxy-D-glycero-α-D-galacto-non-2-ulopyranosylonate)-(2→3)-[β-D-galactopyranosyl-(1→4)]-α-D-glucopyranosyl fluoride (6)**

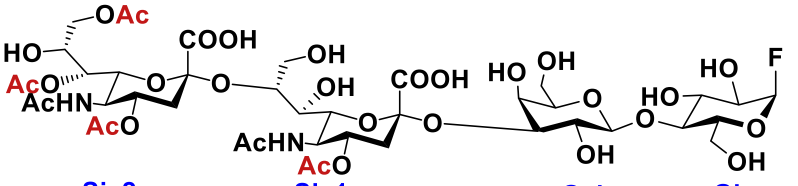


To a stirred solution of compound **7** (5.0 mg, 3.0 μmol) in dioxane/$H_2O$ (1:1, 1 mL) was added Pd $(OH)_2$ (5.0 mg). The mixture was placed under an atmosphere of $H_2$ and stirred overnight. The reaction mixture was filtered, and the filtrate was concentrated *in vacuo*. The residue was dissolved in deionized water and freeze-dried for 24 h. The resulted residue was purified by Biogel P-2 size exclusion column chromatography (5% n-BuOH) to give a product that was further purified by reverse phase (RP)-HPLC (C18, acetonitrile/water 0% –10%, 25 min) to afford **6** (2.1 mg, 63%). $^1$H NMR (600 MHz, $D_2O$) $\delta$ 5.66 (d, $J_{Glc1,2}$ = 2.9 Hz, 0.5H, H-1$_{Glc}$), 5.57 (d, $J_{Glc1,2}$ = 2.9 Hz, 0.5H,

H-$1_{Glc}$), 5.11 (dd, $J_{Sia7,8}$ = 8.7 Hz, $J_{Sia6,7}$ = 2.8 Hz, 1H, H-$7_{Sia2}$), 4.98 (td, $J_{Sia4,5}$ = $J_{Sia3a,4}$ = 11.1 Hz, $J_{Sia3e,4}$ = 4.7 Hz, 1H, H-$4_{Sia2}$), 4.82 (dt, $J_{Sia4,5}$ = $J_{Sia3a,4}$ = 11.0 Hz, $J_{Sia3e,4}$ = 5.6 Hz, 1H, H-$4_{Sia1}$), 4.47 (d, $J_{Gal1,2}$ = 8.0 Hz, 1H, H-$1_{Gal}$), 4.14 (m, 1H, H-$8_{Sia2}$), 4.10 – 4.03 (m, 3H, H-$9a_{Sia2}$, H-$9b_{Sia2}$, H-$3_{Gal}$), 4.02 (m, 2H, H-$5_{Sia2}$, H-$9a_{Sia1}$), 3.98 (m, 1H, H-$8_{Sia1}$), 3.93 (m, 2H, H-$7_{Sia1}$, H-$4_{Glc}$), 3.88 (m, 1H, H-$4_{Gal}$), 3.85 (m, 3H, H-$5_{Sia1}$, H-$6a_{Gal}$, H-$6b_{Gal}$), 3.81 (m, 1H, H-$3_{Glc}$), 3.79 (m, 1H, H-$6_{Sia2}$), 3.74 (m, 1H, H-$6_{Sia1}$), 3.73 – 3.64 (m, 5H, H-$9b_{Sia1}$, H-$6a_{Glc}$, H-$6b_{Glc}$, H-$5_{Gal}$, H-$5_{Glc}$), 3.62 (dd, $J_{Glc2,3}$ = 9.8 Hz, $J_{Glc1,2}$ = 2.9 Hz, 0.5H, H-$2_{Glc}$), 3.57 (dd, $J_{Glc2,3}$ = 9.8 Hz, $J_{Glc1,2}$ = 3.1 Hz, 0.5H, H-$2_{Glc}$), 3.50 (m, 1H, H-$2_{Gal}$), 2.80 (dd, $J_{Sia3a,3e}$ = 12.8 Hz, $J_{Sia3e,4}$ = 4.9 Hz, 1H, H-$3eq_{Sia2}$), 2.59 (dd, $J_{Sia3a,3e}$ = 12.6 Hz, $J_{Sia3e,4}$ = 4.8 Hz, 1H, H-$3eq_{Sia1}$), 2.16 – 1.76 (m, 20H, 6xAc, H-$3ax_{Sia2}$, H-$3ax_{Sia1}$); $^{13}$C NMR (150 MHz, $D_2O$) $\delta$ 174.19 (C-$1_{Sia2}$), 174.08 (C-$1_{Sia1}$), 173.19, 173.14, 173.11, 172.67, 172.61, 107.75 (C-$1_{Glc}$), 106.27 (C-$1_{Glc}$), 102.62 (C-$1_{Gal}$), 78.55, 77.69 (C-$4_{Glc}$), 76.77 (C-$5_{Gal}$), 76.73, 75.55 (C-$3_{Gal}$), 75.22 (C-$5_{Glc}$), 73.61 (C-$6_{Sia1}$), 73.19, 72.90 (C-$4_{Gal}$), 72.23, 71.46 (C-$8_{Sia1}$), 71.27, 71.00 (C-$6_{Sia2}$), 70.87(C-$2_{Glc}$), 70.75 (C-$2_{Glc}$), 70.60 (C-$4_{Sia1}$, C-$4_{Sia2}$), 69.26 (C-$2_{Gal}$), 69.12, 68.95 (C-$7_{Sia2}$), 67.93 (C-$7_{Sia1}$), 67.65 (C-$3_{Glc}$), 67.20 (C-$8_{Sia2}$), 64.02 (C-$9_{Sia2}$), 61.60 (C-$6_{Glc}$, C-$9_{Sia1}$), 59.42 (C-$6_{Gal}$), 49.80, 49.73, 49.37 (C-$5_{Sia2}$), 48.99 (C-$5_{Sia1}$), 44.52, 37.90, 37.65 (C-$3_{Sia1}$), 35.50 (C-$3_{Sia2}$), 22.12 (NCO<u>C</u>$H_3$), 21.65 (NCO<u>C</u>$H_3$), 20.65 (CO<u>C</u>$H_3$), 20.32 (CO<u>C</u>$H_3$), 20.27 (CO<u>C</u>$H_3$), 20.15 (CO<u>C</u>$H_3$); HRMS (ESI): *m/z* calculated for $C_{42}H_{62}FN_2O_{30}$ ($[M-H]^-$) 1093.3377, found 1093.3399.

| **6** | H1 | H2 | H3 | H4 | H5 | H6 | H7 | H8 | H9 |
|---|---|---|---|---|---|---|---|---|---|
| Glc | 5.66 (d, $J_{1,2}$ = 2.9 Hz), 5.57 (d, $J_{1,2}$ = 2.9 Hz) | 3.62 (dd, $J_{2,3}$ = 9.8 Hz, $J_{1,2}$ = 2.9 Hz), 3.57 (dd, $J_{2,3}$ = 9.8 Hz, $J_{1,2}$ = 3.1 Hz) | 3.81 | 3.93 | 3.68 | 3.70-3.64 | - | - | - |
| Gal | 4.47 | 3.51 | 4.06 | 3.89 | 3.72 | 3.88-3.80 | - | - | - |

| | | | | | | | | | |
|---|---|---|---|---|---|---|---|---|---|
| | (d, $J_{1,2}$ = 8.0 Hz) | | | | | | | | |
| Sia1 | - | - | 2.79 (dd, $J_{3a,3e}$ = 12.8 Hz, $J_{3e,4}$ = 4.9 Hz), 1.82 | 4.82 (td, $J_{4,5}$ = $J_{3a,4}$ = 11.0 Hz, $J_{3e,4}$ = 5.6 Hz) | 3.86 | 3.75 | 3.93 | 3.98 | 4.02, 3.73 |
| Sia2 | - | - | 2.59 (dd, $J_{3a,3e}$ = 12.6 Hz, $J_{3e,4}$ = 4.8 Hz), 1.91 | 4.98 (td, $J_{4,5}$ = $J_{3a,4}$ = 11.1 Hz, $J_{3e,4}$ = 4.7 Hz) | 4.03 | 3.79 | 5.11 (dd, $J_{7,8}$ = 8.7 Hz, $J_{6,7}$ = 2.8 Hz) | 4.14 | 4.07 |

| **6** | C1 | C2 | C3 | C4 | C5 | C6 | C7 | C8 | C9 |
|---|---|---|---|---|---|---|---|---|---|
| Glc | 107.7, 106.2 | 70.8 | 67.7 | 77.5 | 75.2 | 61.0 | - | - | - |
| Gal | 102.6 | 69.4 | 75.7 | 72.9 | 76.9 | 59.4 | - | - | - |
| Sia1 | na | na | 37.6 | 70.6 | 48.9 | 73.2 | 68.0 | 71.4 | 60.9 |
| Sia2 | na | na | 35.5 | 70.6 | 49.3 | 71.1 | 68.9 | 67.2 | 64.1 |

**4, 4', 7, 9-Tetra-*O*-acetyl GD3βSph (4)**

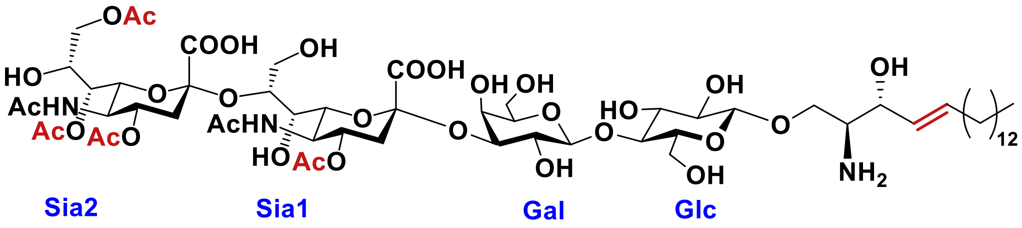


Compound **6** (2.0 mg, 1.8 μmol) was dissolved in 25 mM NaOAc (pH 5.0, 1 mL) containing 0.2% Triton X-100 to which D-erythrosphingosine (1.0 mg, 3.6 μmol) and EGC-II mutant (2 mg/mL, 5 μL) were added. The reaction mixture was incubated at 37 °C for 3 days. The solution was lyophilized, and the resulting mixture was passed through a Biogel P-2 size exclusion column (5% n-BuOH) to give a product that was further purified by reverse phase (RP)-HPLC (C18, acetonitrile/water 0% – 20%, 30 min) affording compound **4** (1.6 mg, 65%). $^{1}$H NMR (600 MHz, $D_2O$) $\delta$ 5.80 (m, 1H, H-$e_{CH=CH}$), 5.43 (m, 1H, H-$d_{CH=CH}$), 5.12 (d, $J_{Sia7,8}$ = 10.2 Hz, 1H, H-$7_{Sia2}$),

4.87 (m, 2H, H-4$_{Sia2}$, H-4$_{Sia1}$), 4.47 (d, $J_{Gal1,2}$ = 7.8 Hz, 1H, H-1$_{Gal}$), 4.40 (m, 1H, H-1$_{Glc}$), 4.37 (m, 1H, H-c), 4.21 (m, 1H, H-8$_{Sia2}$), 4.12 (m, 1H, H-3$_{Gal}$), 4.07 (m, 3H, H-9a$_{Sia1}$, H-9a$_{Sia2}$, H-9b$_{Sia2}$), 4.03 (m, 1H, H-4$_{Glc}$), 3.95 (m, 1H, H-5$_{Sia2}$), 3.91 (m, 2H, H-4$_{Gal}$, H-6$_{Sia2}$), 3.89 – 3.83 (m, 3H, H-6a$_{Glc}$, H-6b$_{Glc}$, H-5$_{Sia1}$), 3.81 (m, 1H, H-7$_{Sia1}$), 3.73 (m, 1H, H-8$_{Sia1}$), 3.68 (m, 3H, H-9b$_{Sia1}$, H-6a$_{Gal}$, H-6b$_{Gal}$), 3.66 (m, 1H, H-6$_{Sia1}$), 3.64 (m, 1H, H-5$_{Gal}$), 3.61 (m, 2H, O$C\underline{H}_{2,a}$), 3.56 (m, 1H, H-3$_{Glc}$), 3.52 (m, 1H, H-2$_{Gal}$), 3.48 (m, 1H, H-b), 3.46 (m, 1H, H-5$_{Glc}$), 3.30 (m, 1H, H-2$_{Glc}$), 2.77 (m, 1H, H-3eq$_{Sia2}$), 2.61 (m, 1H, H-3eq$_{Sia1}$), 2.25 – 1.75 (m, 22H, 6xAc, H-3ax$_{Sia1}$, H-3ax$_{Sia2}$, $C\underline{H}_{2,f}$), 1.23 (m, 22H, 11x$C\underline{H}_2$), 0.83 (m, 3H, $C\underline{H}_3$); $^{13}$C NMR (150 MHz, $D_2O$) $\delta$ 135.83 (=$\underline{C}$H sphingosine), 126.42 (H$\underline{C}$=C sphingosine), 102.84 (C-1$_{Gal}$), 102.26 (C-1$_{Glc}$), 78.54 (C-3$_{Gal}$), 77.90 (C-5$_{Gal}$), 75.58 (C-4$_{Glc}$), 75.05 (C-6$_{Sia1}$), 74.91(C-5$_{Glc}$), 74.27 (C-3$_{Glc}$), 73.84 (C-8$_{Sia1}$), 72.59 ( C-2$_{Glc}$), 71.30 (C-6$_{Sia2}$), 70.91 (C-4$_{Sia1}$, C-4$_{Sia2}$), 69.70 (-O$\underline{C}H_2$- C1-sphingosine), 69.46 (C-7$_{Sia1}$), 69.40 (-$\underline{C}$HOH sphingosine), 69.31 (C-2$_{Gal}$), 68.88 (C-7$_{Sia2}$), 67.61(C-4$_{Gal}$), 67.24 (C-8$_{Sia2}$), 64.05 (C-9$_{Sia2}$), 61.61 (C-9$_{Sia1}$), 61.31 (C-6$_{Gal}$), 59.87 (C-6$_{Glc}$), 54.80 (-$\underline{C}HNH_2$ sphingosine), 49.71 (C-5$_{Sia2}$), 48.89 (C-5$_{Sia1}$), 37.53 (C-3$_{Sia2}$), 36.70 (C-3$_{Sia1}$), 32.05 (-$\underline{C}H_2$-), 31.93 (-$\underline{C}H_2$-), 29.61 (-$\underline{C}H_2$-), 29.02 (-$\underline{C}H_2$-), 22.54 (-$\underline{C}H_2$-), 22.07 (NCO$\underline{C}H_3$), 21.82 (NCO$\underline{C}H_3$), 20.36 (CO$\underline{C}H_3$), 20.35 (CO$\underline{C}H_3$), 13.90 (-$CH_2\underline{C}H_3$); HRMS (ESI): *m/z* calculated for $C_{60}H_{98}N_3O_{32}$ ([M-H]$^-$) 1372.6139, found 1372.6161.

| **4** | H1 | H2 | H3 | H4 | H5 | H6 | H7 | H8 | H9 |
|---|---|---|---|---|---|---|---|---|---|
| Glc | 4.37 | 3.33 | 3.57 | 4.03 | 3.47 | 3.90-3.82 | - | - | - |
| Gal | 4.46(d, $J_{1,2}$ = 7.8 Hz) | 3.52 | 4.12 | 3.92 | 3.64 | 3.72-3.62 | - | - | - |
| Sia1 | - | - | 2.61, 1.82 | 4.84 | 3.86 | 3.67 | 3.81 | 3.74 | 4.08, 3.68 |
| Sia2 | - | - | 2.75, 1.80 | 4.90 | 3.95 | 3.91 | 5.10(d, $J_{7,8}$ = 10.2 Hz) | 4.19 | 4.12 |

| **4** | C1 | C2 | C3 | C4 | C5 | C6 | C7 | C8 | C9 |
|---|---|---|---|---|---|---|---|---|---|
| Glc | 102.3 | 72.6 | 74.3 | 75.6 | 74.9 | 59.8 | - | - | - |
| Gal | 102.8 | 69.3 | 78.5 | 67.6 | 77.9 | 61.2 | - | - | - |
| Sia1 | na | na | 36.7 | 70.9 | 48.9 | 75.1 | 69.4 | 73.8 | 61.6 |
| Sia2 | na | na | 37.5 | 70.9 | 49.7 | 71.3 | 68.9 | 67.2 | 64.0 |

**7,9-di-*O*-Acetyl GD3βSph (16)**

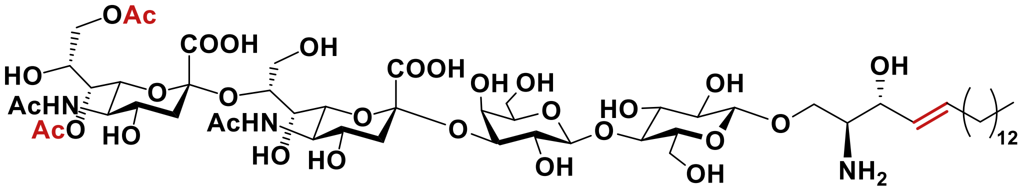


Compound **4** (2.0 mg, 1.5 μmol) was dissolved in 50 mM $HCOONH_4$ (500 μL) to which MHVS HE (1 mg/mL, 2 μL) was added. The reaction mixture was incubated at 37 °C for 2 h. The solution was lyophilized, and the resulting mixture passed through a Biogel P-2 size exclusion column (5% n-BuOH) to give a product that was further purified by reverse phase (RP)-HPLC (C18, acetonitrile/water 0% – 20%, 30 min) to afford **16** (0.8 mg, 81%). $^1$H NMR (600 MHz, $CD_3OD$) $\delta$ 5.77 (dt, $J_{trans}$ = 15.1 Hz, $J_{vic}$ = 7.4 Hz, 1H, H-$e_{CH=CH}$), 5.39 (dd, $J_{trans}$ = 15.4, $J_{vic}$ = 6.7 Hz, 1H, H-$d_{CH=CH}$), 5.01 (dd, $J_{Sia7,8}$ = 9.4 Hz, $J_{Sia6,7}$ = 2.4 Hz, H, H-$7_{Sia2}$), 4.37 (d, $J_{Gal1,2}$ = 7.9 Hz, 1H, H-$1_{Gal}$), 4.26 (d, $J_{Glc1,2}$ = 7.8 Hz, 1H, H-$1_{Glc}$), 4.22 (m, 1H, H-c), 4.10 – 4.01 (m, 2H, H-$8_{Sia2}$, H-$9a_{Sia1}$), 3.95 (m, 1H, H-$3_{Gal}$), 3.93 (m, 2H, H-$9a_{Sia2}$, H-$9b_{Sia2}$), 3.89 (m, 1H, H-$4_{Gal}$), 3.84 – 3.79 (m, 3H, H-$6a_{Glc}$, H-$6b_{Glc}$, H-$6_{Sia2}$), 3.76 (m, 1H, H-$5_{Sia2}$), 3.72 (m, 2H, H-$9b_{Sia1}$, H-$4_{Glc}$), 3.70 (m, 2H, H-$5_{Sia1}$, H-$8_{Sia1}$), 3.67 (m, 1H, H-$6a_{Gal}$), 3.59 (m, 2H, H-$6b_{Gal}$, H-$4_{Sia1}$), 3.55 (m, 2H, H-$7_{Sia1}$, H-$6_{Sia1}$), 3.50 (m, 1H, H-$5_{Gal}$), 3.48 (m, 1H, OC$\underline{H}_{2,a}$), 3.47 (m, 1H, H-$2_{Gal}$), 3.44 (m, 1H, H-$3_{Glc}$), 3.41 (m, 1H, OC$\underline{H}_{2,a}$), 3.40 (m, 1H, H-$4_{Sia2}$), 3.37 (m, 1H, H-$5_{Glc}$), 3.28 (m, 1H, H-b), 3.20 (m, 1H, H-$2_{Glc}$), 2.84 (dd, $J_{Sia3a,3e}$ = 12.6 Hz, $J_{Sia3e,4}$ = 4.8 Hz, 1H, H-$3eq_{Sia2}$), 2.60 (dd, $J_{Sia3a,3e}$ = 12.4 Hz, $J_{Sia3e,4}$ = 4.5 Hz, 1H, H-$3eq_{Sia1}$), 2.04 – 1.90 (m, 14H, 4xAc, C$\underline{H}_{2,f}$), 1.64 (t, $J_{Sia3a,3e}$ = $J_{Sia3a,4}$ = 12.0 Hz, 1H, H-$3ax_{Sia1}$), 1.60 (t, $J_{Sia3a,3e}$ = $J_{Sia3a,4}$ = 12.1 Hz, 1H, H-$3ax_{Sia2}$), 1.33 – 1.15 (m, 22H, 11xC$\underline{H}_2$), 0.80 (t, $J$ = 7.0 Hz, 3H, C$\underline{H}_3$); $^{13}$C NMR (150 MHz, $CD_3OD$) $\delta$ 134.49 (=$\underline{C}$H sphingosine), 126.15 (H$\underline{C}$=C sphingosine), 102.78 (C-$1_{Gal}$), 101.53 (C-$1_{Glc}$), 78.30 (C-$5_{Gal}$), 75.25 (C-$3_{Gal}$), 74.73 (C-$6_{Sia1}$), 74.41 (C-$5_{Glc}$), 73.98 (C-$3_{Glc}$), 72.99 (C-$8_{Sia1}$), 72.32 (C-$2_{Glc}$), 71.61 (C-$7_{Sia1}$), 71.21 (C-$6_{Sia2}$), 68.70 (C-$7_{Sia2}$), 68.64 (C-$2_{Gal}$, -$\underline{C}$HOH sphingosine), 68.53 (C-$4_{Sia2}$), 67.35 (C-$4_{Glc}$), 67.24 (C-$4_{Sia1}$), 66.90 (C-$8_{Sia2}$), 66.82 (C-$4_{Gal}$), 63.71 (C-$9_{Sia2}$), 62.21 (-O$\underline{C}H_2$- C1-sphingosine), 60.60 (C-$9_{Sia1}$), 60.49 (C-$6_{Gal}$), 59.24 (C-$6_{Glc}$), 54.57 (-$\underline{C}HNH_2$ sphingosine), 51.81 (C-$5_{Sia2}$), 50.57 (C-$5_{Sia1}$), 40.58 (C-$3_{Sia2}$), 39.13 (C-$3_{Sia1}$), 31.10 (-$\underline{C}H_2$-), 28.46 (-$\underline{C}H_2$-), 28.31 (-$\underline{C}H_2$-), 27.95 (-$\underline{C}H_2$-), 21.56 (-$\underline{C}H_2$-), 20.72 (NCO$\underline{C}H_3$), 18.77 (CO$\underline{C}H_3$), 12.18 (-$CH_2\underline{C}H_3$); HRMS (ESI): *m/z* calculated for $C_{56}H_{94}N_3O_{30}$ ($[M-H]^-$) 1288.5928, found 1288.5910.

| **16** | H1 | H2 | H3 | H4 | H5 | H6 | H7 | H8 | H9 |
|---|---|---|---|---|---|---|---|---|---|
| Glc | 4.26 ($J_{1,2}$ = 7.8 Hz) | 3.20 | 3.44 | 3.72 | 3.37 | 3.85, 3.82 | - | - | - |
| Gal | 4.37 (d, $J_{1,2}$ = 7.9 Hz) | 3.47 | 3.95 | 3.89 | 3.50 | 3.67, 3.60 | - | - | - |
| Sia1 | - | - | 2.60 (dd, $J_{3a,3e}$ = 12.4 Hz, $J_{3e,4}$ = 4.5 Hz), 1.64 (t, $J_{3a,3e}$ = $J_{3a,4}$ = 12.0 Hz) | 3.59 | 3.70 | 3.55 | 3.55 | 3.70 | 4.05, 3.67 |
| Sia2 | - | - | 2.84 (dd, $J_{3a,3e}$ = 12.6 Hz, $J_{3e,4}$ = 4.8 Hz), 1.95 | 3.40 | 3.76 | 3.81 | 5.01 ($J_{7,8}$ = 9.4 Hz, $J_{6,7}$ = 2.4 Hz) | 4.05 | 3.93 |

| **16** | C1 | C2 | C3 | C4 | C5 | C6 | C7 | C8 | C9 |
|---|---|---|---|---|---|---|---|---|---|
| Glc | 101.5 | 72.3 | 74.0 | 67.3 | 74.4 | 59.2 | - | - | - |
| Gal | 102.8 | 68.6 | 75.2 | 66.8 | 78.2 | 60.5 | - | - | - |
| Sia1 | na | na | 39.1 | 67.2 | 50.6 | 74.7 | 71.6 | 73.0 | 60.6 |
| Sia2 | na | na | 40.5 | 68.5 | 51.8 | 71.2 | 68.7 | 66.9 | 63.6 |

**7-*O*-Acetyl GD3βSph (17)**

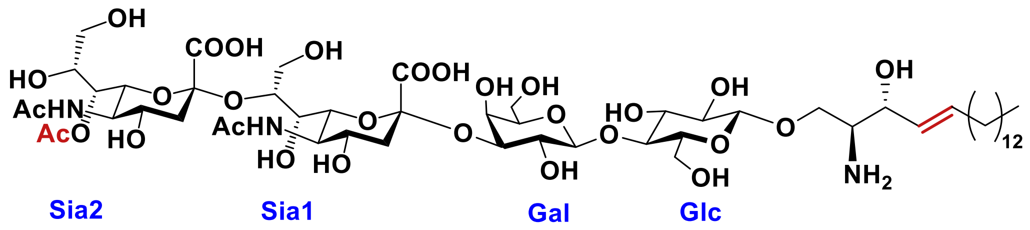


Compound **4** (2.0 mg, 1.5 µmol) was dissolved in 50 mM $HCOONH_4$ (500 µL) to which MHVS HE (1 mg/mL, 2 µL) and BCov HE (1 mg/mL, 2 µL) was added. The reaction mixture was incubated at 37 °C for 2 h. The solution was lyophilized, and the resulting mixture was passed through a Biogel P-2 (5% n-BuOH) column to give a product that was further purified by reverse phase (RP)-HPLC (C18, acetonitrile/water 0% – 20%, 30 min) to afford **17** (0.8 mg, 83%). $^{1}$H NMR (600 MHz, $CD_3OD$) $\delta$ 5.78 (dt, $J_{trans}$ = 15.4 Hz, $J_{vic}$ = 7.4 Hz, 1H, H-$e_{CH=CH}$), 5.39 (dd, $J_{trans}$ = 15.3, $J_{vic}$ = 6.8 Hz, 1H, H-$d_{CH=CH}$), 4.89 (dd, $J_{Sia7,8}$ = 9.3 Hz, $J_{Sia6,7}$ = 2.4 Hz, 1H, H-$7_{Sia2}$), 4.32

(d, $J_{Gal1,2}$ = 7.9 Hz, 1H, H-1$_{Gal}$), 4.26 (d, $J_{Glc1,2}$ = 7.8 Hz, 1H, H-1$_{Glc}$), 4.23 (m, 1H, H-c), 3.98 (dd, $J_{Sia9a,9b}$ = 12.0 Hz, $J_{Sia8,9a}$ = 3.3 Hz, 1H, H-9a$_{Sia1}$), 3.93 (d, $J_{Gal3,4}$ = 3.4 Hz, 1H, H-4$_{Gal}$), 3.89 (m, 2H, H-6$_{Sia2}$, H-3$_{Gal}$), 3.87 (m, 1H, H-4$_{Sia1}$), 3.86 – 3.81 (m, 3H, H-6a$_{Glc}$, H-9a$_{Sia2}$, H-9b$_{Sia2}$), 3.78 (m, 1H, H-4$_{Glc}$), 3.75 (m, 1H, H-6b$_{Glc}$), 3.72 (m, 2H, H-9b$_{Sia1}$, H-5$_{Sia2}$), 3.70 (m, 1H, H-8$_{Sia1}$), 3.69 (m, 1H, H-5$_{Sia1}$), 3.66 (m, 2H, H-6a$_{Gal}$, H-6b$_{Gal}$), 3.64 (m, 1H, H-8$_{Sia2}$), 3.60 (m, 1H, H-6$_{Sia1}$), 3.46 (m, 3H, OC$\underline{H}_{2,a}$, H-2$_{Gal,}$ H-5$_{Gal}$), 3.45 (m, 1H, H-3$_{Glc}$), 3.38 (m, 1H, H-4$_{Sia2}$), 3.36 (m, 2H, H-7$_{Sia1}$, H-5$_{Glc}$), 3.34 (m, 1H, OC$\underline{H}_{2,a}$), 3.29 (m, 1H, H-b), 3.20 (m, 1H, H-2$_{Glc}$), 2.89 (dd, $J_{Sia3a,3e}$ = 12.9 Hz, $J_{Sia3e,4}$ = 4.6 Hz, 1H, H-3eq$_{Sia2}$), 2.52 (dd, $J_{Sia3a,3e}$ = 12.7 Hz, $J_{Sia3e,4}$ = 4.7 Hz, 1H, H-3eq$_{Sia1}$), 2.00 – 1.89 (m, 8H, 2xAc, C$\underline{H}_{2,f}$), 1.79 (s, 3H, Ac), 1.76 (t, $J_{Sia3a,3e}$ = $J_{Sia3a,4}$ = 12.1 Hz, 1H, H-3ax$_{Sia1}$), 1.58 (t, $J_{Sia3a,3e}$ = $J_{Sia3a,4}$ = 12.5 Hz, 1H, H-3ax$_{Sia2}$), 1.36 – 1.12 (m, 22H, 11xC$\underline{H}_2$), 0.80 (t, $J$ = 7.0 Hz, 3H, C$\underline{H}_3$); $^{13}$C NMR (150 MHz, $CD_3OD$) $\delta$ 134.53 (=$\underline{C}$H sphingosine), 126.07 (H$\underline{C}$=C sphingosine), 102.84 (C-1$_{Gal}$), 101.48 (C-1$_{Glc}$), 78.48 (C-5$_{Gal}$), 75.16 (C-3$_{Gal}$), 74.57 (C-8$_{Sia1}$), 74.49 (C-6$_{Sia1}$), 74.35 (C-5$_{Glc}$), 74.14 (C-3$_{Glc}$), 73.11(C-7$_{Sia1}$), 72.27 (C-2$_{Glc}$), 71.81 (C-6$_{Sia2}$), 69.79 (C-4$_{Glc}$), 68.95 (C-7$_{Sia2}$), 68.65 (-$\underline{C}$HOH sphingosine), 68.57 (C-2$_{Gal}$), 68.07 (C-4$_{Sia2}$), 67.83 (C-4$_{Gal}$) , 66.60 (C-4$_{Sia1}$), 65.75 (C-8$_{Sia2}$), 64.86 (C-9$_{Sia2}$), 61.95 (-O$\underline{C}H_2$- C1-sphingosine), 60.45 (C-6$_{Gal}$), 59.66 (C-9$_{Sia1}$), 59.43 (C-6$_{Glc}$), 54.47 (-$\underline{C}HNH_2$ sphingosine), 51.42 (C-5$_{Sia2}$), 50.59 (C-5$_{Sia1}$), 40.64 (C-3$_{Sia2}$), 36.60 (C-3$_{Sia1}$), 34.30 (-$\underline{C}H_2$-), 31.21 (-$\underline{C}H_2$-), 30.81(-$\underline{C}H_2$-), 28.41(-$\underline{C}H_2$-), 28.14(-$\underline{C}H_2$-), 27.92 (-$\underline{C}H_2$-), 25.84 (-$\underline{C}H_2$-), 24.60 (-$\underline{C}H_2$-), 21.41 (-$\underline{C}H_2$-), 20.67 (NCO$\underline{C}H_3$), 20.37 (NCO$\underline{C}H_3$), 18.88 (CO$\underline{C}H_3$), 12.17 (-$CH_2\underline{C}H_3$); HRMS (ESI): $m/z$ calculated for $C_{54}H_{92}N_3O_{29}$ ([M-H]$^-$) 1246.5822, found 1246.5798.

| **17** | H1 | H2 | H3 | H4 | H5 | H6 | H7 | H8 | H9 |
|---|---|---|---|---|---|---|---|---|---|
| Glc | 4.26 (d, $J_{1,2}$ = 7.8 Hz) | 3.21 | 3.45 | 3.77 | 3.36 | 3.83, 3.76 | - | - | - |
| Gal | 4.32 (d, $J_{1,2}$ = 7.9 Hz) | 3.38 | 3.88 | 3.93 (d, $J_{3,4}$ = 3.4 Hz) | 3.45 | 3.69, 3.66 | - | - | - |
| Sia1 | - | - | 2.52 (dd, $J_{3a,3e}$ = 12.7 Hz, $J_{3e,4}$ = | 3.86 | 3.69 | 3.60 | 3.37 | 3.70 | 3.98, 3.71 |

| | | | | | | | | | |
|---|---|---|---|---|---|---|---|---|---|
| | | | 4.7 Hz), 1.76 (t, $J_{3a,3e}$ = $J_{3a,4}$ = 12.1 Hz) | | | | | | |
| Sia2 | - | - | 2.89, (dd, $J_{3a,3e}$ = 12.9 Hz, $J_{3e,4}$ =4.6 Hz), 1.58 (t, $J_{3a,3e}$ = $J_{3a,4}$ = 12.5 Hz) | 3.38 | 3.72 | 3.88 | 4.89 (dd, $J_{7,8}$ = 9.3 Hz, $J_{6,7}$ = 2.4 Hz) | 3.64 | 3.86, 3.82 |

| **17** | C1 | C2 | C3 | C4 | C5 | C6 | C7 | C8 | C9 |
|---|---|---|---|---|---|---|---|---|---|
| Glc | 101.5 | 72.3 | 74.1 | 69.8 | 74.4 | 59.4 | - | - | - |
| Gal | 102.9 | 68.5 | 75.2 | 67.8 | 78.5 | 60.5 | - | - | - |
| Sia1 | na | na | 36.6 | 66.6 | 50.6 | 74.5 | 73.1 | 74.6 | 59.6 |
| Sia2 | na | na | 40.6 | 68.1 | 51.4 | 71.8 | 68.9 | 65.8 | 64.9 |

**7,9-Di-*O*-acetyl GD3 (1)**

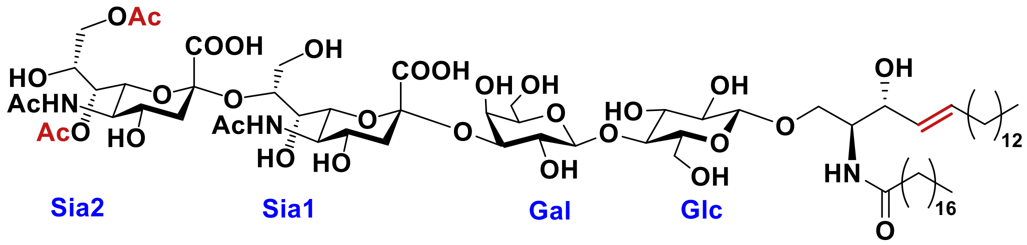


To a solution of 7,9-di-OAc-GD3βSph **16** (1.0 mg, 0.8 μmol) in DMSO (500 μL), CsF (0.2 mg, 1.3 μmol) and stearic acid-NHS ester (0.5 mg, 1.3 μmol) were added. The resulting reaction mixture was vigorously stirred at RT for 2 h. The solution was concentrated, and the residue was purified by silica gel column chromatography (ethyl acetate:methanol:water= 7:2:1) to give compound **1** (1.0 mg, 83%). $^1$H NMR (600 MHz, $CD_3OD$) $\delta$ 5.59 (m, 1H, H-$e_{CH=CH}$), 5.34 (dd, $J_{trans}$ = 15.3 Hz, $J_{vic}$ = 7.8 Hz, 1H), H-$d_{CH=CH}$), 5.00 (dd, $J_{Sia7,8}$ = 9.2 Hz, $J_{Sia6,7}$ = 2.3 Hz, 1H, H-

7$_{Sia2}$), 4.36 (d, $J_{Gal1,2}$ = 7.8 Hz, 1H, H-1$_{Gal}$), 4.20 (d, $J_{Glc1,2}$ = 7.9 Hz, 1H, H-1$_{Glc}$), 4.09 (dd, $J_{gem}$ = 10.0, $J_{vic}$ = 4.3 Hz, 1H, OC$\underline{H}_{2,a}$), 4.04 – 3.90 (m, 5H, H-8$_{Sia2}$, H-c, H-3$_{Gal}$, H-9a$_{Sia2}$, H-9b$_{Sia2}$), 3.89 – 3.75 (m, 6H, H-b, H-6a$_{Glc}$, H-6b$_{Glc}$, H-6$_{Sia2}$, H-4$_{Glc}$, H-4$_{Gal}$), 3.72 – 3.61 (m, 7H, H-5$_{Sia1}$, H-5$_{Sia2}$, H-6a$_{Gal}$, H-6b$_{Gal}$, H-9a$_{Sia1}$, H-9b$_{Sia1}$, H-6$_{Sia1}$), 3.58 (m, 1H, H-8$_{Sia1}$), 3.54 (m, 1H, H-2$_{Gal1}$), 3.50 (m, 1H, H-5$_{Gal}$), 3.49 (m, 1H, H-7$_{Sia1}$), 3.46(m, 1H, OC$\underline{H}_{2,a}$), 3.43 (m, 1H, H-3$_{Glc}$), 3.40(m, 2H, H-4$_{Sia2}$, H-4$_{Sia1}$), 3.32 (m, 1H, H-5$_{Glc}$), 3.19 (m, 1H, H-2$_{Glc}$), 2.90 (d, $J_{Sia3a,3e}$ = 12.6 Hz, 2H, H-3eq$_{Sia2}$, H-3eq$_{Sia1}$), 2.00 – 1.79 (m, 14H, 4xAc, H-3ax$_{Sia1}$, H-3ax$_{Sia2}$, C$\underline{H}_{2,f}$), 1.20 (m, 56H, 26xC$\underline{H}_2$), 0.80 (t, $J$ = 7.0 Hz, 6H, 2xC$\underline{H}_3$); $^{13}$C NMR (150 MHz, CD$_3$OD) $\delta$ 132.85 (=$\underline{C}$H sphingosine), 129.22 (H$\underline{C}$=C sphingosine), 102.81 (C-1$_{Gal}$), 102.27 (C-1$_{Glc}$), 78.78 (C-5$_{Gal}$), 74.64 (C-3$_{Gal}$), 74.30 (C-5$_{Glc}$, C-8$_{Sia1}$), 74.10 (C-7$_{Sia1}$), 74.00 (C-3$_{Glc}$), 72.63 (C-2$_{Glc}$), 71.42 (C-4$_{Glc}$, C-4$_{Gal}$, C-6$_{Sia2}$), 71.29 (C-6$_{Sia1}$), 70.67 (-$\underline{C}$HOH sphingosine), 68.50 (C-4$_{Sia1}$, C-4$_{Sia2}$), 68.39 (C-2$_{Gal}$), 67.61 (-O$\underline{C}$H$_2$- C1-sphingosine), 67.08 (C-8$_{Sia2}$) , 63.66 (C-9$_{Sia2}$), 60.68 (C-9$_{Sia1}$), 60.65 (C-6$_{Gal}$), 59.66 (C-6$_{Glc}$), 52.45 (-$\underline{C}$HNH$_2$ sphingosine), 51.66 (C-5$_{Sia2}$), 50.47 (C-5$_{Sia1}$), 40.81 (C-3$_{Sia1}$, C-3$_{Sia2}$), 35.27 (-$\underline{C}$H$_2$-), 32.94 (-$\underline{C}$H$_2$-), 31.22 (-$\underline{C}$H$_2$-), 30.85 (-$\underline{C}$H$_2$-), 28.48 (-$\underline{C}$H$_2$-), 28.14 (-$\underline{C}$H$_2$-), 24.79 (-$\underline{C}$H$_2$-), 21.49 (-$\underline{C}$H$_2$-), 20.69 (NCO$\underline{C}$H$_3$), 20.45 (NCO$\underline{C}$H$_3$), 18.69 (CO$\underline{C}$H$_3$), 18.56 (CO$\underline{C}$H$_3$), 12.20(-CH$_2\underline{C}$H$_3$); HRMS (ESI): *m/z* calculated for $C_{74}H_{128}N_3O_{31}$ ([M-H]$^-$) 1554.8537, found 1554.8494.

| **1** | H1 | H2 | H3 | H4 | H5 | H6 | H7 | H8 | H9 |
|---|---|---|---|---|---|---|---|---|---|
| Glc | 4.20 (d, $J_{1,2}$ = 7.9 Hz) | 3.19 | 3.43 | 3.87-3.80 | 3.36 | 3.79 | - | - | - |
| Gal | 4.36 (d, $J_{1,2}$ = 7.8 Hz) | 3.54 | 3.95 | 3.87-3.80 | 3.45 | 3.72-3.61 | - | - | - |
| Sia1 | - | - | 2.90 (d, $J_{3a,3e}$ = 12.6 Hz), 1.54 | 3.40 | 3.69 | 3.69 | 3.49 (dd, $J_{7,8}$ = 9.2 Hz, $J_{6,7}$ = 2.3 Hz) | 3.58 | 3.72-3.61 |
| Sia2 | - | - | 2.90 (d, $J_{3a,3e}$ = 12.6 Hz), 1.54 | 3.40 | 3.71 | 3.87-3.80 | 5.00 | 4.01 | 3.93 |

| 1 | C1 | C2 | C3 | C4 | C5 | C6 | C7 | C8 | C9 |
|---|---|---|---|---|---|---|---|---|---|
| Glc | 102.2 | 72.6 | 74.0 | 71.4 | 74.3 | 59.6 | - | - | - |
| Gal | 102.9 | 68.4 | 74.7 | 71.4 | 78.5 | 60.7 | - | - | - |
| Sia1 | na | na | 40.8 | 68.5 | 50.4 | 71.3 | 74.1 | 74.3 | 60.7 |
| Sia2 | na | na | 40.8 | 68.5 | 51.7 | 71.4 | 68.6 | 67.1 | 63.7 |

**7-*O*-Acetyl GD3 (2)**

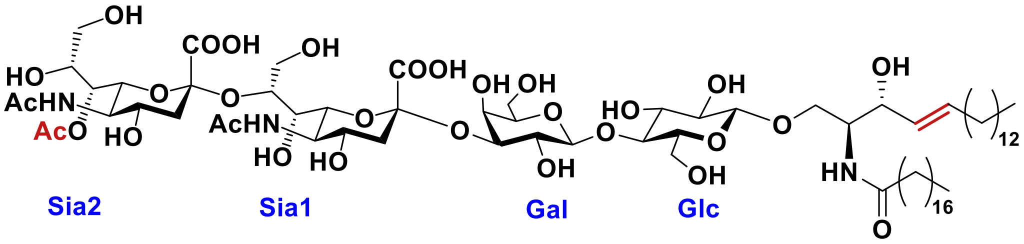


To a solution of 7-OAc-GD3βSph **17** (1.0 mg, 0.8 μmol) in DMSO (500 μL) were added CsF (0.2 mg, 1.3 μmol) and stearic acid-NHS ester (0.5 mg, 1.3 μmol). The resulting reaction mixture was stirred vigorously at RT for 2 h. The solution was concentrated, and the residue was purified by silica gel column chromatography (ethyl acetate: methanol: water= 7: 2: 1) to give compound **2** (0.96 mg, 80%). $^1$H NMR (600 MHz, $CD_3OD$) $\delta$ 5.63 (dd, $J_{trans}$ = 14.9 Hz, , $J_{vic}$ = 7.1 Hz, 1H, H-$e_{CH=CH}$), 5.39 (dd, $J_{trans}$ = 15.3 Hz, $J_{vic}$ = 7.7 Hz, 1H, H-$d_{CH=CH}$), 4.90 (d, $J_{Sia7,8}$ = 9.3 Hz, 1H, H-$7_{Sia2}$), 4.44 (d, $J_{Gal1,2}$ = 7.9 Hz, 1H, H-$1_{Gal}$), 4.24 (d, $J_{Glc1,2}$ = 7.8 Hz, 1H, H-$1_{Glc}$), 4.14 (dd, $J_{gem}$ = 10.0 Hz, $J_{vic}$ = 4.4 Hz, 1H, OC$\underline{H}_{2,a}$), 4.05 (m, 2H, H-$9a_{Sia2}$, H-$3_{Gal}$), 4.02 (m, 1H, H-c), 3.96 (m, 2H, H-$8_{Sia2}$, H-$4_{Gal}$), 3.91 (dt, $J$ = 8.1, 3.8 Hz, 1H, H-b), 3.90 (m, 1H, H-$6_{Sia2}$), 3.88 – 3.83 (m, 3H, H-$6a_{Glc}$, H-$6b_{Glc}$, H-$4_{Glc}$), 3.81 (s, 2H, H-$6_{Sia1}$, H-$9b_{Sia2}$), 3.78 – 3.59 (m, 4H, H-$5_{Sia1}$, H-$5_{Sia2}$, H-$6a_{Gal}$, H-$6b_{Gal}$), 3.58 (s, 2H, H-$8_{Sia1}$, H-$5_{Gal}$), 3.56 (m, 1H, H-$2_{Gal}$), 3.50 (m, 2H, H-$9a_{Sia1}$, OC$\underline{H}_{2,a}$), 3.47 (m, 3H, H-$4_{Sia2}$, H-$7_{Sia1}$, H-$3_{Glc}$), 3.42 (m, 1H, H-$4_{Sia1}$), 3.37 (m, 2H, H-$9b_{Sia1}$, H-$5_{Glc}$), 3.23 (m, 1H, H-$2_{Glc}$), 2.90 (d, $J_{Sia3a,3e}$= 12.0 Hz, 1H, H-$3eq_{Sia2}$), 2.61 (m, 1H, H-$3eq_{Sia1}$), 2.06 – 1.83 (m, 11H, 3xAc, C$\underline{H}_{2,f}$), 1.67 (m, 1H, H-$3ax_{Sia1}$), 1.59 (t, $J_{Sia3a,3e}$ = $J_{Sia3a,4}$ = 12.0 Hz, 1H, H-$3ax_{Sia2}$), 1.24 (s, 52H, 26x$CH_2$), 0.85 (t, $J$ = 7.0 Hz, 6H, 2xC$\underline{H}_3$); $^{13}$C NMR (150 MHz, $CD_3OD$) $\delta$ 132.94 (=$\underline{C}$H sphingosine), 129.31 (H$\underline{C}$=C sphingosine), 102.75 (C-$1_{Gal}$), 102.27 (C-$1_{Glc}$), 78.55 (C-$5_{Gal}$), 74.54 (C-$3_{Gal}$), 74.25 (C-$5_{Glc}$), 74.09 (C-$3_{Glc}$), 73.28 (C-$6_{Sia1}$), 72.84 (C-$2_{Glc}$), 72.21 (C-$7_{Sia1}$) , 71.24 (C-$4_{Glc}$, C-$6_{Sia2}$), 70.72 (-$\underline{C}$HOH sphingosine), 69.44 (C-$4_{Gal}$, C-$8_{Sia2}$), 69.16 (C-$7_{Sia2}$), 68.59 (C-$4_{Sia2}$), 68.20 (C-$2_{Gal}$, C-$4_{Sia1}$), 67.67(-O$\underline{C}H_2$- C1-sphingosine), 62.04 (C-$9_{Sia1}$), 61.18 (C-$9_{Sia2}$), 60.60 (C-$6_{Gal}$), 59.66 (C-$6_{Glc}$), 52.53 (-$\underline{C}HNH_2$ sphingosine), 52.10 (C-$5_{Sia2}$), 50.49 (C-$5_{Sia1}$), 41.01

(C-3$_{Sia2}$), 39.93 (C-3$_{Sia1}$), 35.25 (-C̲H$_2$-), 31.27 (-C̲H$_2$-), 30.78 (-C̲H$_2$-), 28.57 (-C̲H$_2$-), 28.19 (-C̲H$_2$-), 21.41 (-C̲H$_2$-), 20.79 (NCOC̲H$_3$), 20.61 (NCOC̲H$_3$), 18.89 (COC̲H$_3$), 12.26 (-CH$_2$C̲H$_3$); HRMS (ESI): *m/z* calculated for $C_{72}H_{126}N_3O_{30}$ ([M-H]$^-$) 1512.8432, found 1512.8419.

| **2** | H1 | H2 | H3 | H4 | H5 | H6 | H7 | H8 | H9 |
|---|---|---|---|---|---|---|---|---|---|
| Glc | 4.24 (d, $J_{1,2}$ = 7.8 Hz) | 3.24 | 3.48 | 3,86 | 3.37 | 3.86 | - | - | - |
| Gal | 4.44 (d, $J_{1,2}$ = 7.9 Hz) | 3.50 | 4.06 | 3.96 | 3.58 | 3.70, 3.65 | - | - | - |
| Sia1 | - | - | 2.61, 1.67 | 3.42 | 3.74 | 3.81 | 3.48 | 3.58 | 3.51, 3.37 |
| Sia2 | - | - | 2.90 (d, $J_{3a,3e}$ = 12.0 Hz), 1.59 (t, $J_{3a,3e}$ = $J_{3a,4}$ = 12.0 Hz) | 3.47 | 3.75 | 3.90 | 4.90 (d, $J_{7,8}$ = 9.3 Hz) | 3.96 | 4.05, 3.81 |

| **2** | C1 | C2 | C3 | C4 | C5 | C6 | C7 | C8 | C9 |
|---|---|---|---|---|---|---|---|---|---|
| Glc | 102.3 | 72.8 | 74.1 | 71.2 | 74.2 | 59.7 | - | - | - |
| Gal | 102.8 | 68.2 | 74.5 | 69.4 | 78.6 | 60.6 | - | - | - |
| Sia1 | na | na | 39.9 | 68.2 | 50.5 | 73.3 | 72.2 | 74.5 | 62.0 |
| Sia2 | na | na | 41.0 | 68.6 | 52.1 | 71.2 | 69.2 | 69.4 | 61.2 |

**9-*O*-Acetyl GD3 (3)**

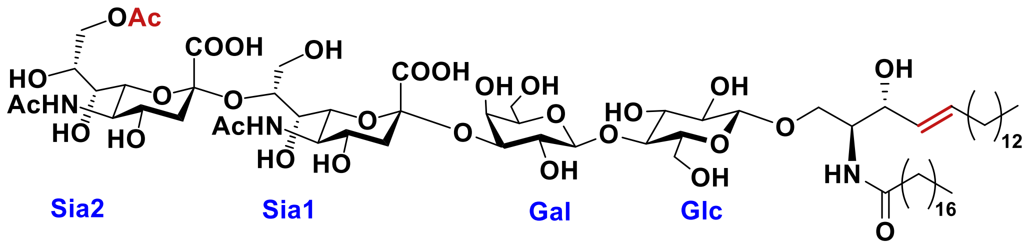


Compound **2** (0.2 mg, 0.1 μmol) was dissolved in 50 mM $NH_4HCO_3$ (pH 8.0, 500 μL), and the solution was incubated at 37 °C overnight. The solution was lyophilized to give compound **3** (0.2 mg, quant.). $^{1}$H NMR (600 MHz, $CD_3OD$) $\delta$ 5.58 (dt, $J_{trans}$ = 15.1 Hz, $J_{vic}$ = 6.7 Hz, 1H, H-$e_{CH=CH}$), 5.34 (dd, $J_{trans}$ = 15.4 Hz, $J_{vic}$ = 7.9 Hz, 1H, H-$d_{CH=CH}$), 4.40 (d, $J_{Gal1,2}$ = 8.0 Hz, 1H, H-$1_{Gal}$), 4.25 (dd, $J_{Sia9a,9b}$ = 11.5 Hz, $J_{Sia8,9a}$ = 2.2 Hz, 1H, H-$9a_{Sia2}$), 4.20 (d, $J_{Glc1,2}$ = 7.8 Hz, 1H, H-$1_{Glc}$), 4.09 (dd, $J_{gem}$ = 10.1 Hz, $J_{vic}$ = 4.4 Hz, 1H, OC$\underline{H}_{2,a}$), 4.03 (m, 2H, H-$9b_{Sia2}$, H-$3_{Gal}$), 3.99 (m, 1H, H-$4_{Glc}$), 3.98 (m, 1H, H-c), 3.95 (m, 1H, H-$8_{Sia2}$), 3.93 (m, 1H, H-$4_{Gal}$), 3.86 (m, 2H, H-b, H-$6_{Sia1}$), 3.82 (m, 2H, H-$6a_{Glc}$, H-$6b_{Glc}$), 3.75 (m, 2H, H-$5_{Sia2}$, H-$6_{Sia2}$), 3.66 (m, 1H, H-$6a_{Gal}$), 3.60 (m, 1H, H-$5_{Sia1}$), 3.58 (m, 1H, H-$6b_{Gal}$), 3.52 (m, 2H, H-$8_{Sia1}$, H-$5_{Gal}$), 3.50 (m, 1H, H-$7_{Sia2}$)3.48 (m, 2H, H-$9a_{Sia1}$, H-$2_{Gal}$), 3.45 (m, 3H, OC$\underline{H}_{2,a}$, H-$3_{Glc}$, H-$7_{Sia1}$), 3.42 (m, 1H, H-$9b_{Sia1}$) 3.36 (m, 2H, H-$4_{Sia1}$, H-$4_{Sia2}$), 3.33 (m, 1H, H-$5_{Glc}$), 3.19 (m, 1H, H-$2_{Glc}$), 2.83 (dd, $J_{Sia3a,3e}$ = 11.8 Hz, $J_{Sia3e,4}$ = 3.4 Hz, 1H, H-$3eq_{Sia2}$), 2.61 (dd, $J_{Sia3a,3e}$ = 10.4 Hz, $J_{Sia3e,4}$ = 3.4 Hz, 1H, H-$3eq_{Sia1}$), 1.98 – 1.85 (m, 11H, 3xAc, C$\underline{H}_{2,f}$), 1.65 (m, 1H, H-$3ax_{Sia1}$), 1.60 (t, $J_{Sia3a,3e}$ = $J_{Sia3a,4}$ = 12.0 Hz, 1H, H-$3ax_{Sia2}$), 1.22 – 1.17 (m, 52H, 26xC$\underline{H}_2$), 0.81 (t, $J$ = 6.8 Hz, 6H, 2xC$\underline{H}_3$); $^{13}$C NMR (150 MHz, $CD_3OD$) $\delta$ 132.93 (=$\underline{C}$H sphingosine), 128.62 (H$\underline{C}$=C sphingosine), 102.43 (C-$1_{Gal}$), 102.21 (C-$1_{Glc}$), 78.61 (C-$5_{Gal}$), 76.13 (C-$3_{Gal}$), 74.64 (C-$4_{Glc}$), 74.51 (C-$8_{Sia1}$), 74.17 (C-$5_{Glc}$), 74.03 (C-$3_{Glc}$), 72.79 (C-$6_{Sia1}$), 72.62 (C-$2_{Glc}$), 72.19 (C-$7_{Sia1}$), 70.69 (-$\underline{C}$HOH sphingosine), 68.64 (C-$2_{Gal}$), 68.21 (C-$8_{Sia2}$), 68.08 (C-$4_{Sia1}$, C-$4_{Sia2}$), 67.88 (C-$6_{Sia2}$), 67.56 (-O$\underline{C}H_2$- C1-sphingosine), 67.50 (C-$7_{Sia2}$), 66.29 (C-$4_{Gal}$), 64.84 (C-$9_{Sia2}$), 62.27 (C-$9_{Sia1}$), 60.54 (C-$6_{Gal}$), 59.49 (C-$6_{Glc}$), 52.47 (-$\underline{C}HNH_2$ sphingosine), 52.00 (C-$5_{Sia2}$), 51.72 (C-$5_{Sia1}$), 40.19 (C-$3_{Sia2}$), 40.03 (C-$3_{Sia1}$), 35.11 (-$\underline{C}H_2$-), 31.21 (-$\underline{C}H_2$-), 30.81 (-$\underline{C}H_2$-), 28.52 (-$\underline{C}H_2$-), 28.21 (-$\underline{C}H_2$-), 21.37 (-$\underline{C}H_2$-), 20.39 (NCO$\underline{C}H_3$), 18.60 (CO$\underline{C}H_3$), 12.11 (-$CH_2\underline{C}H_3$); HRMS (ESI): *m/z* calculated for $C_{72}H_{126}N_3O_{30}$ ($[M-H]^-$) 1512.8432, found 1512.8439.

| **3** | H1 | H2 | H3 | H4 | H5 | H6 | H7 | H8 | H9 |
|---|---|---|---|---|---|---|---|---|---|
| Glc | 4.20 (d, $J_{1,2}$ = 7.8 Hz) | 3.19 | 3.45 | 3,99 | 3.33 | 3.82 | - | - | - |
| Gal | 4.40 (d, $J_{1,2}$ = 8.0 Hz) | 3.48 | 4.03 | 3.92 | 3.52 | 3.66, 3.58 | - | - | - |
| Sia1 | - | - | 2.61 (dd, $J_{3a,3e}$ = 10.4 Hz, $J_{3e,4}$ = 3.4 Hz), 1.65 | 3.36 | 3.60 | 3.86 | 3.45 | 3.52 | 3.48, 3.42 |
| Sia2 | - | - | 2.83 (dd, $J_{3a,3e}$ = 11.8 Hz, $J_{3e,4}$ = 3.4 Hz), 1.60 (t, $J_{3a,3e}$ = $J_{3a,4}$ = 12.0 Hz) | 3.36 | 3.75 | 3.75 | 3.50 | 3.95 | 4.25 (dd, $J_{9a,9b}$ = 11.5 Hz, $J_{8,9a}$ = 2.2 Hz), 4.03 |

| **3** | C1 | C2 | C3 | C4 | C5 | C6 | C7 | C8 | C9 |
|---|---|---|---|---|---|---|---|---|---|
| Glc | 102.2 | 72.6 | 74.0 | 74.6 | 74.2 | 59.5 | - | - | - |
| Gal | 102.4 | 68.6 | 76.1 | 66.3 | 78.6 | 60.5 | - | - | - |
| Sia1 | na | na | 40.0 | 68.1 | 51.7 | 72.8 | 72.2 | 74.5 | 62.3 |
| Sia2 | na | na | 40.2 | 68.1 | 52.0 | 67.9 | 67.5 | 68.2 | 64.8 |

**GD3 (18)**

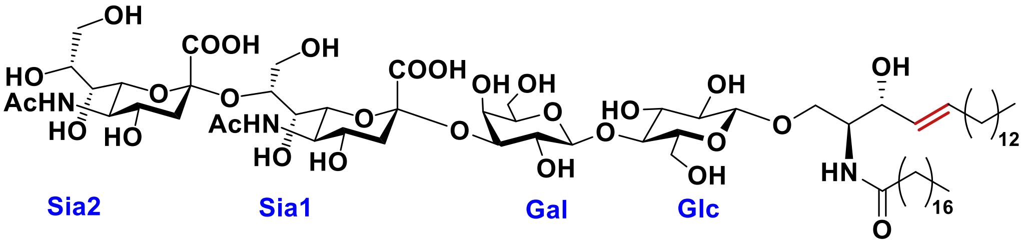


To a solution of **1** (0.2 mg, 0.12 μmol) in $H_2O$ (1 mL) was added NaOH (0.1 mg, 2.5 mmol) at RT. After being stirred for 2 h, the reaction mixture was concentrated *in vacuo*. The residue was purified by size exclusion chromatography (LH-20, MeOH: $H_2O$ = 1:1) to afford compound **18** (0.1 mg, 53%). HRMS (ESI): *m/z* calculated for $C_{70}H_{124}N_3O_{29}$ ($[M-H]^-$) 1470.8326, found 1470.8348. The NMR data agree with reported data.[7]

$^1H$ NMR (600 MHz, $CD_3OD$) $\delta$ (non-carbohydrate): 5.58 (dt, $J_{trans}$ = 14.5 Hz, $J_{vic}$ = 6.7 Hz, 1H, H-e$_{CH=CH}$), 5.34 (dd, $J_{trans}$ = 15.3 Hz, $J_{vic}$ = 7.7 Hz, 1H, H-d$_{CH=CH}$), 4.09 (dd, $J_{gem}$ = 10.1 Hz, $J_{vic}$ = 4.3 Hz, 1H, OC$\underline{H}_{2,a}$), 3.97 (m, 1H, H-c), 3.87 (m, 1H, H-b), 3.46 (m, 1H, OC$\underline{H}_{2,a}$), 2.00 – 1.84 (m, 8H, 2xAc, C$\underline{H}_{2,f}$), 1.63 – 1.15 (m, 52H, 26xC$\underline{H}_2$), 0.80 (t, $J$ = 7.1 Hz, 6H, 2xC$\underline{H}_3$);

| **18** | H1 | H2 | H3 | H4 | H5 | H6 | H7 | H8 | H9 |
|---|---|---|---|---|---|---|---|---|---|
| Glc | 4.20 (d, $J_{1,2}$ = 7.8 Hz) | 3.19 | 3.44 | 3,99 | 3.32 | 3.81 | - | - | - |
| Gal | 4.40 | 3.52 | 4.00 | 3.89 | 3.51 | 3.67, 3.61 | - | - | - |
| Sia1 | - | - | 2.53, 1.65 | 3.46 | 3.59 | na | 3.45 | 3.54 | 3.73, 3.51 |
| Sia2 | - | - | 2.85, 1.58 | 3.46 | 3.59 | 3.76 | 3.59 | na | 3.68-3.64 |

#### 4. References

[1] Z. Li, Y. Lang, L. Liu, M. I. Bunyatov, A. I. Sarmiento, R. J. de Groot, G. J. Boons. Synthetic *O*-acetylated sialosides facilitate functional receptor identification for human respiratory viruses. *Nat. Chem.* **2021**, *13*, 496-503.

[2] S. M. Hancock, J. R. Rich, M. E. Caines, N. C. Strynadka, S. G. Withers. Designer enzymes for glycosphingolipid synthesis by directed evolution. *Nat. Chem. Biol.* **2009**, *5*, 508-514.

[3] G. M. Vos, K. C. Hooijschuur, Z. Li, J. Fjeldsted, C. Klein, R. P. de Vries, J. Toraño Sastre, G. J. Boons. Sialic acid *O*-acetylation patterns and glycosidic linkage type determination by ion mobility-mass spectrometry. *Nat. Commun.* **2023**, *14*, 6795.

[4] M. L. Killian, in *Animal Influenza Virus. Methods in Molecular Biology,, Vol. 1161* (Ed.: E. Spackman), Humana Press, New York, NY, **2014**, pp. 3-9.

[5] S. S. Weng, Y. D. Lin, C. T. Chen. Highly diastereoselective thioglycosylation of functionalized peracetylated glycosides catalyzed by MoO2Cl2. *Org. Lett.* **2006**, *8*, 5633-5636.

[6] X. Dai, W. Liu, Q. Zhou, C. Cheng, C. Yang, S. Wang, M. Zhang, P. Tang, H. Song, D. Zhang, Y. Qin. Formal synthesis of anticoagulant drug fondaparinux sodium. *J. Org. Chem.* **2016**, *81*, 162-184.

[7] a) J. C. Castro-Palomino, B. Simon, O. Speer, M. Leist, R. R. Schmidt. Synthesis of ganglioside GD3 and its comparison with bovine GD3 with regard to oligodendrocyte apoptosis mitochondrial damage. *Chem. Eur. J.* **2001**, *7*, 2178-2184; b) Q. Li, M. Jaiswal, R. S. Rohokale, Z. Guo. A diversity-oriented strategy for chemoenzymatic synthesis of glycosphingolipids and related derivatives. *Org. Lett.* **2020**, *22*, 8245-8249.

#### 5. NMR Specta

[1]H NMR of **S2**. 600 MHz, $CDCl_3$

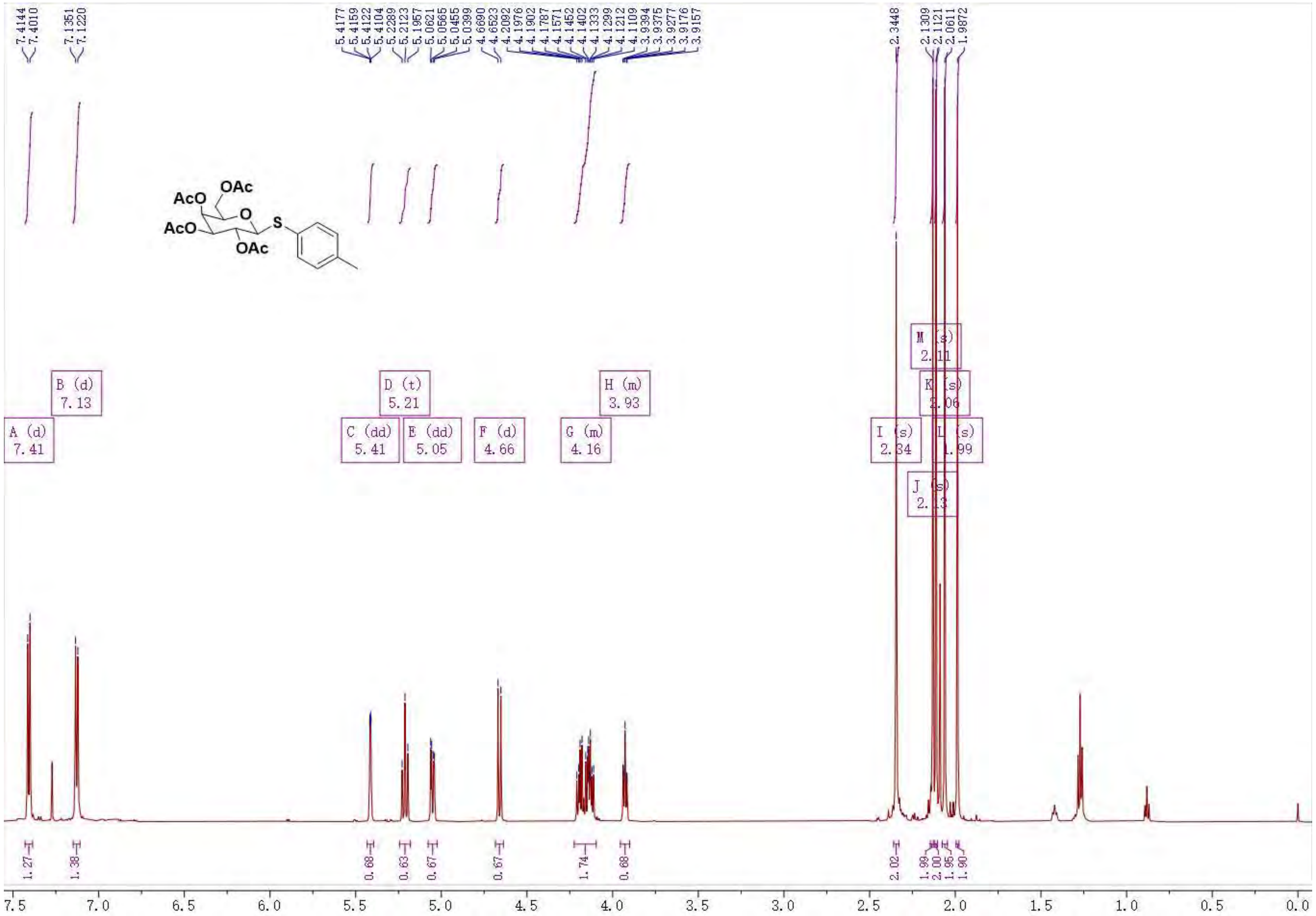

$^{13}C$ NMR of **S2**. 125 MHz, $CDCl_3$.

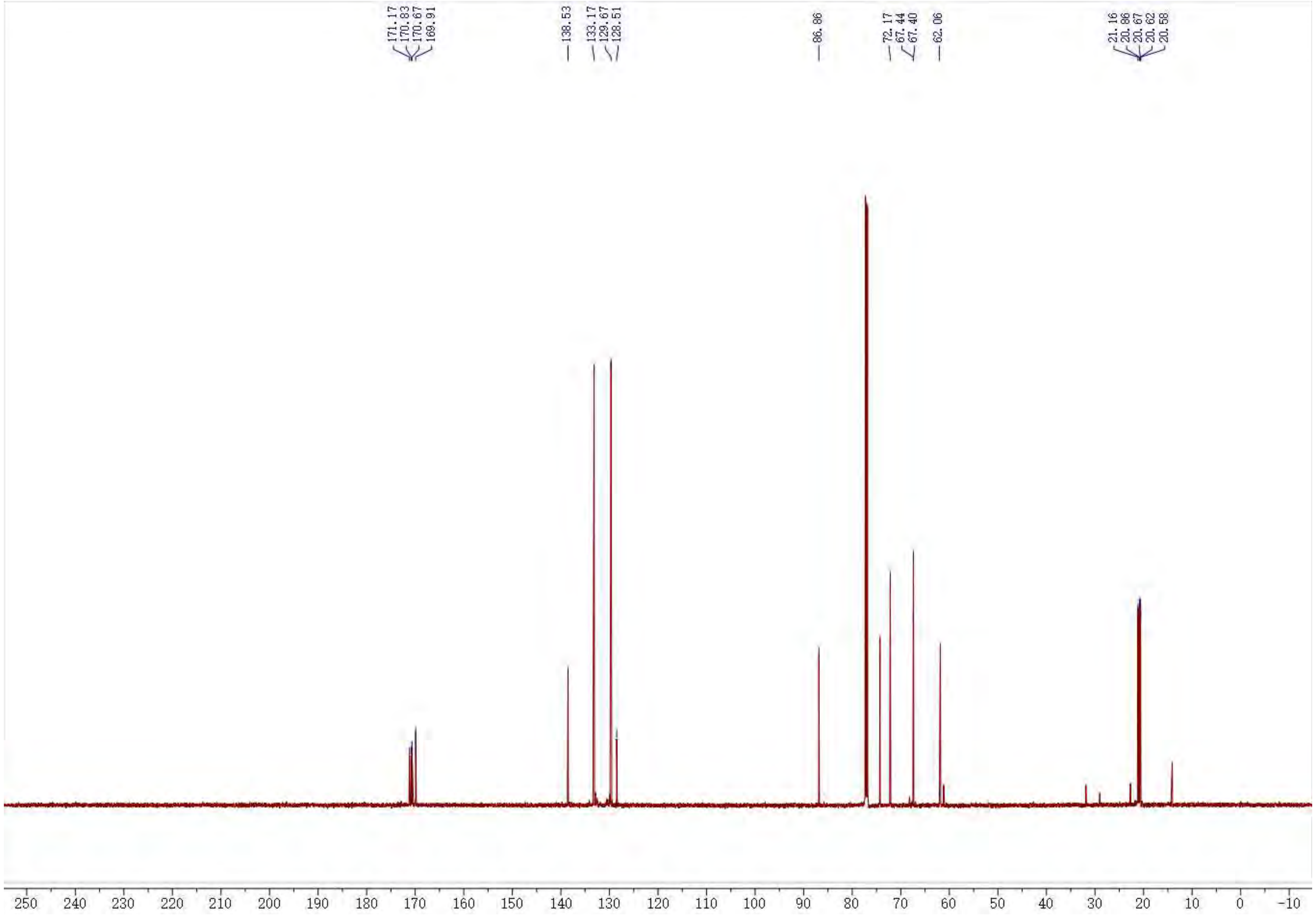

[1]H NMR of **S3**. 600 MHz, $CDCl_3$

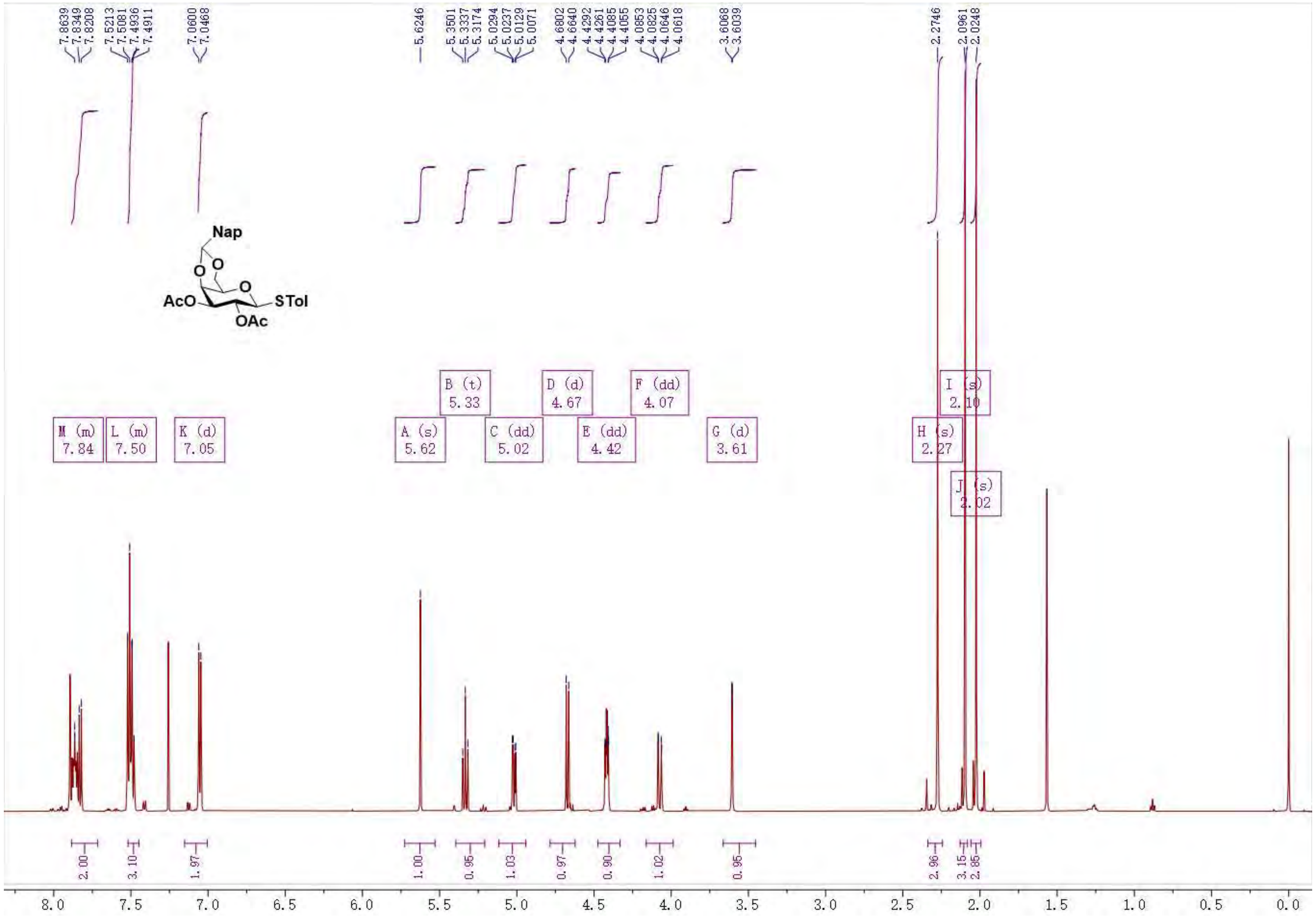

$^{13}C$ NMR of **S3**. 125 MHz, $CDCl_3$

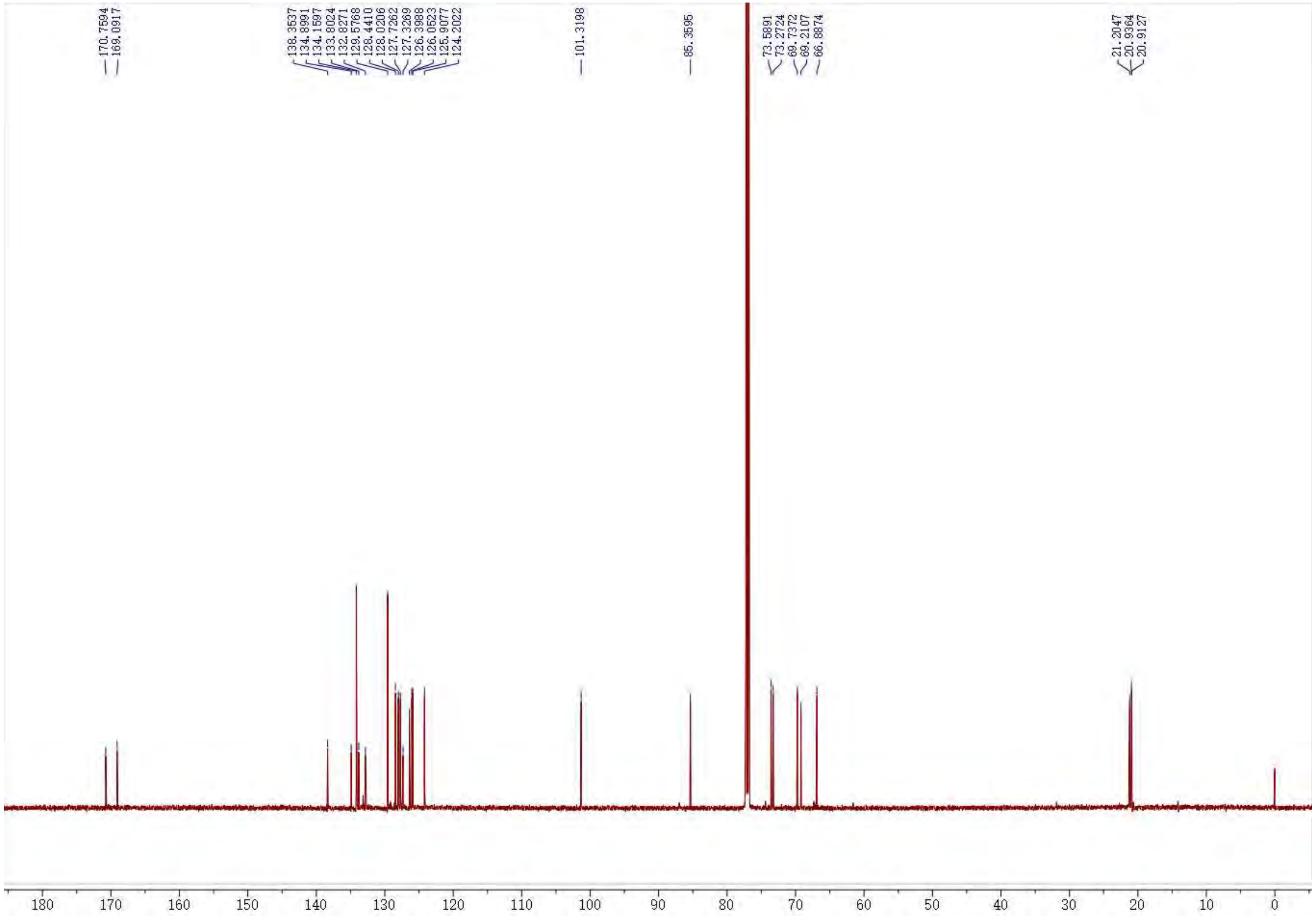

[1]H NMR of **S4**. 600 MHz, $CDCl_3$

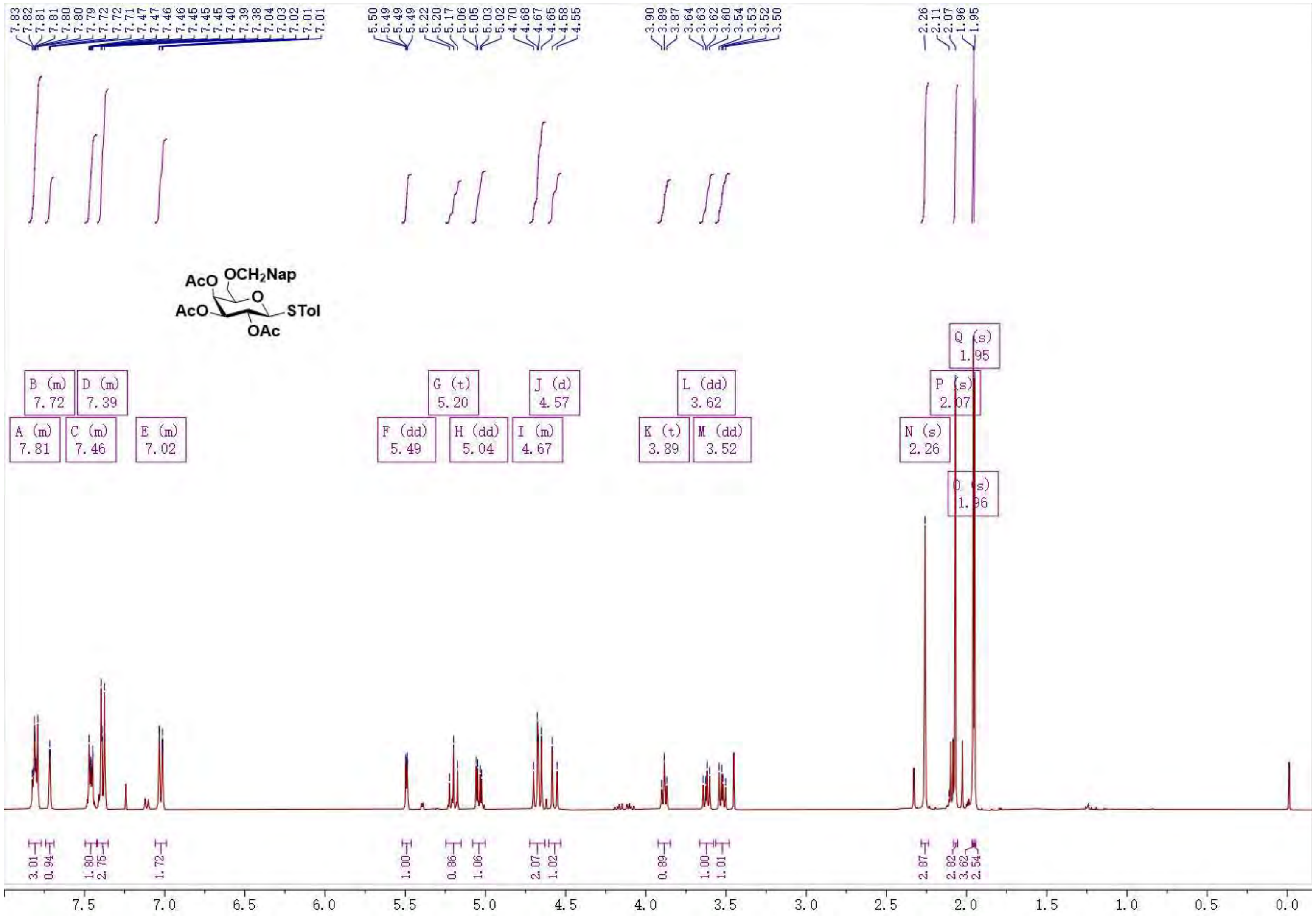

$^{13}$C NMR of **S4**. 125 MHz, $CDCl_3$

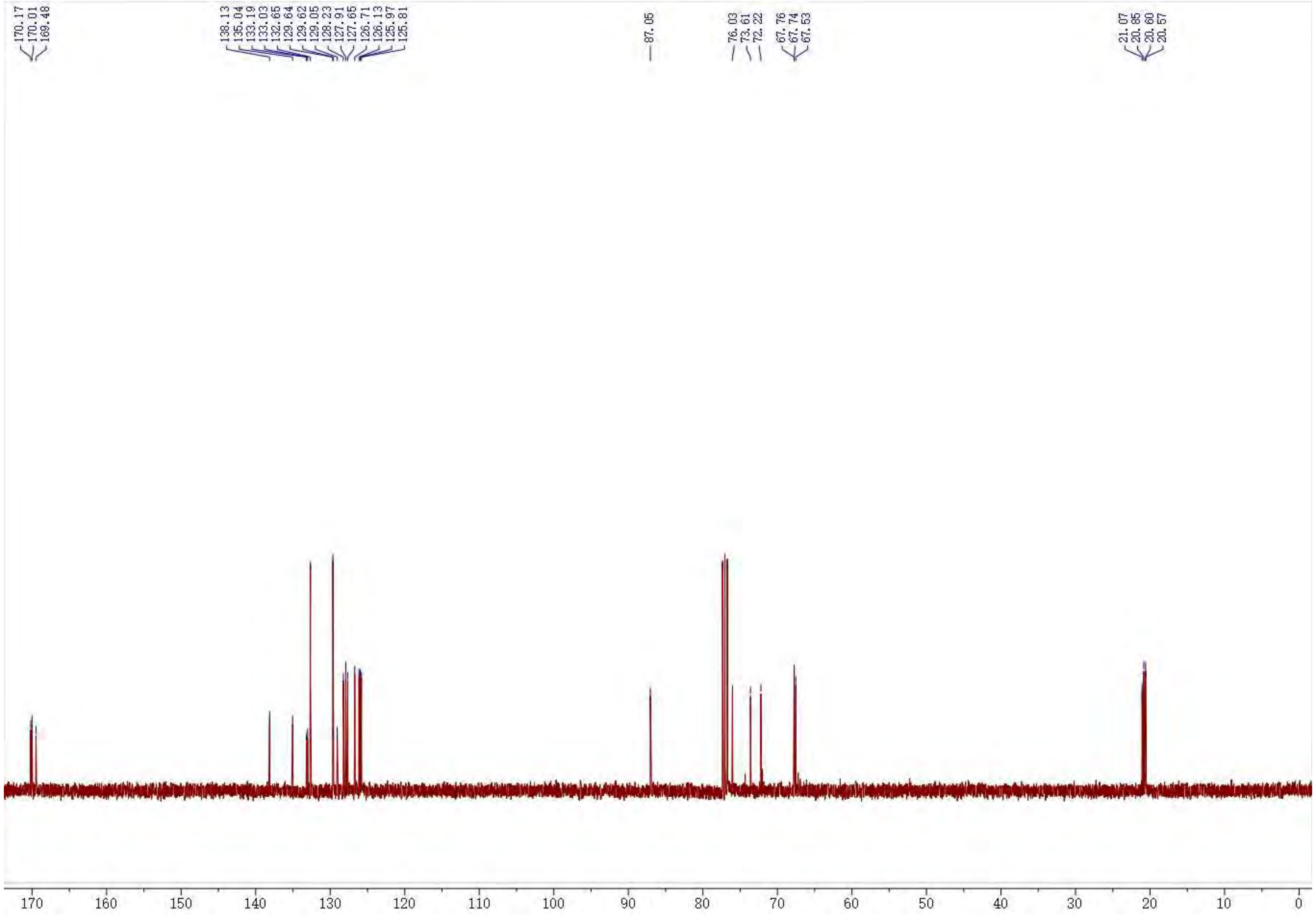

$^1$H NMR of **S5**. 600 MHz, $CDCl_3$

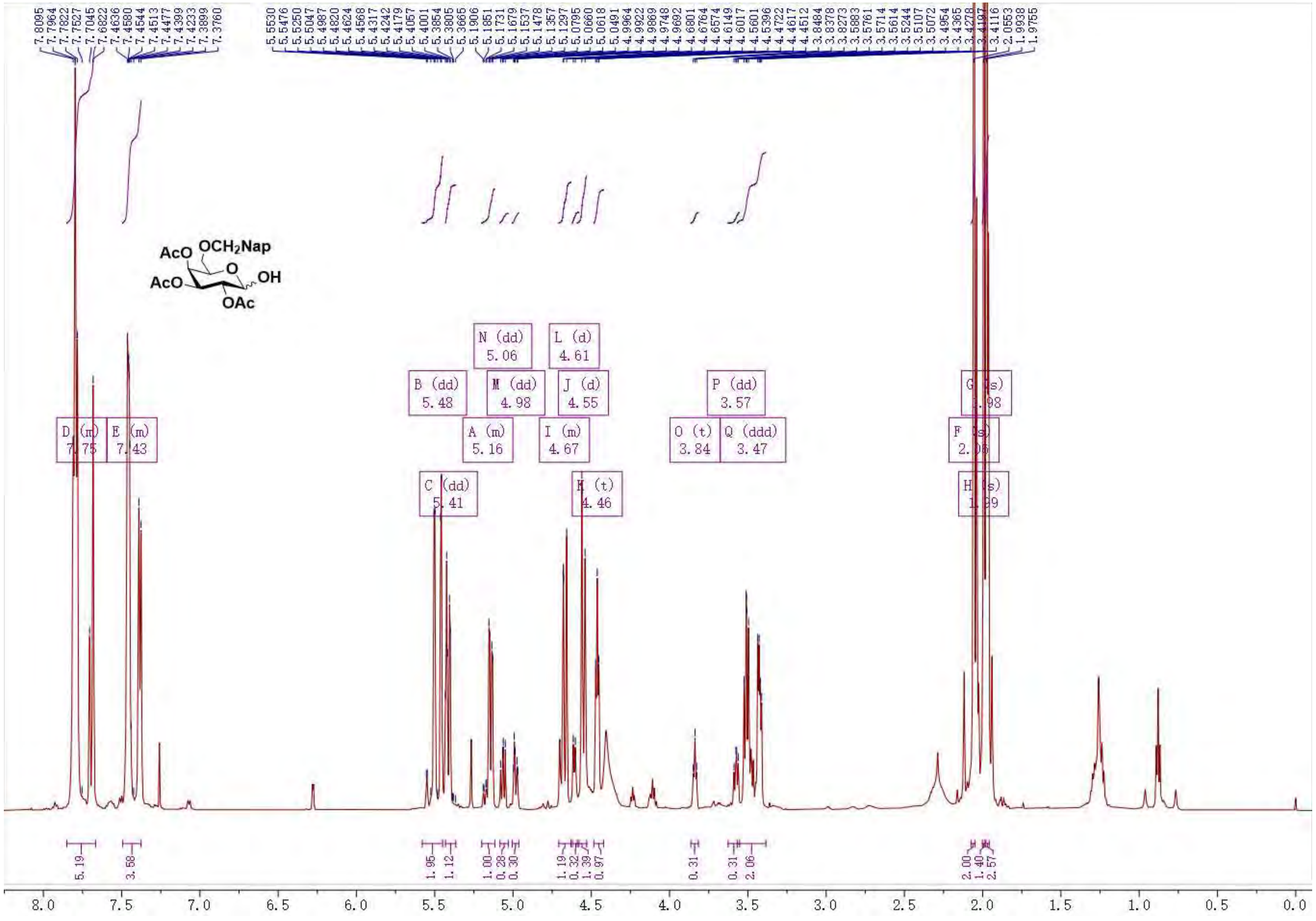

$^{13}C$ NMR of **S5**. 125 MHz, $CDCl_3$

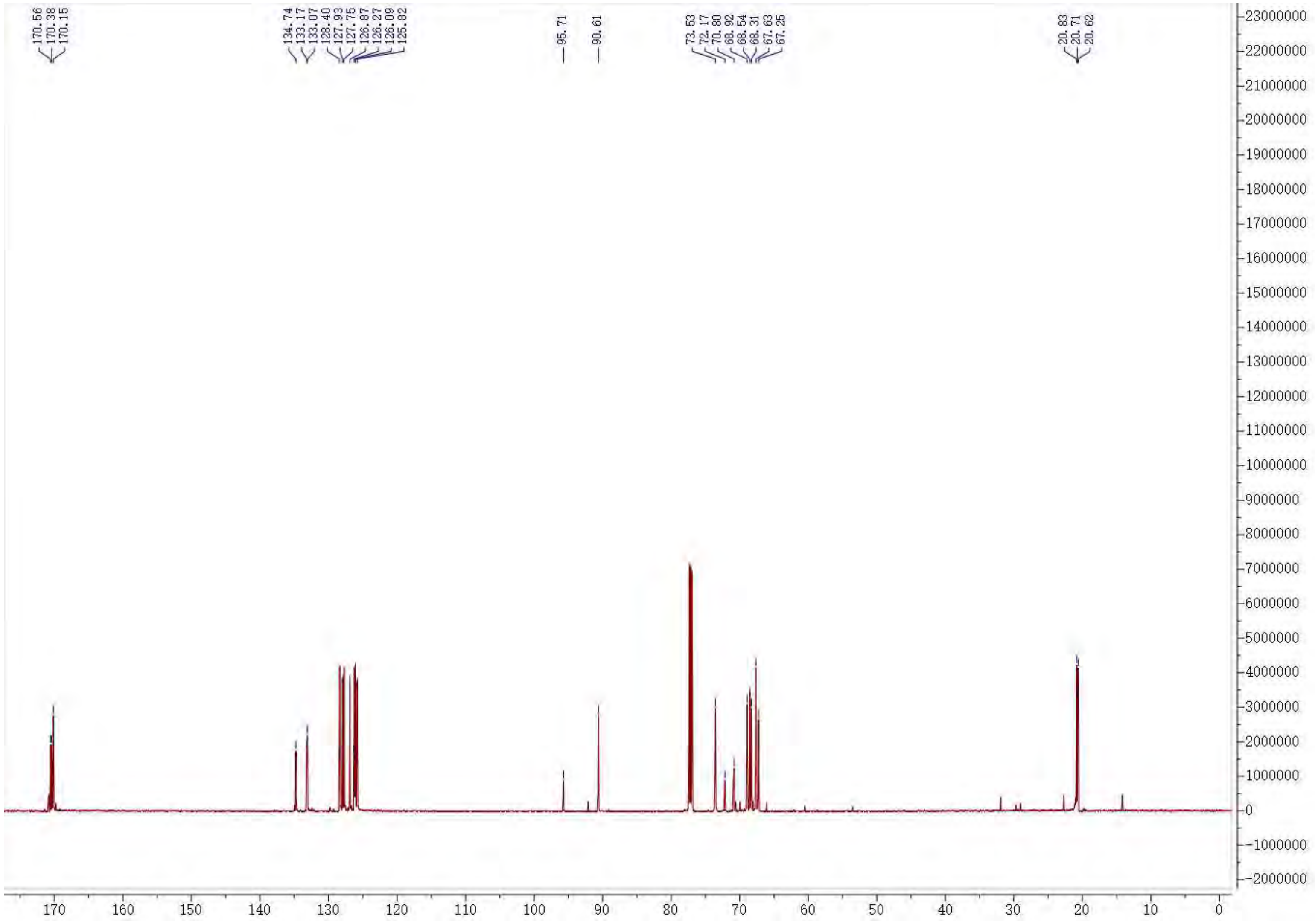

[1]H NMR of **S6**. 400 MHz, $CDCl_3$

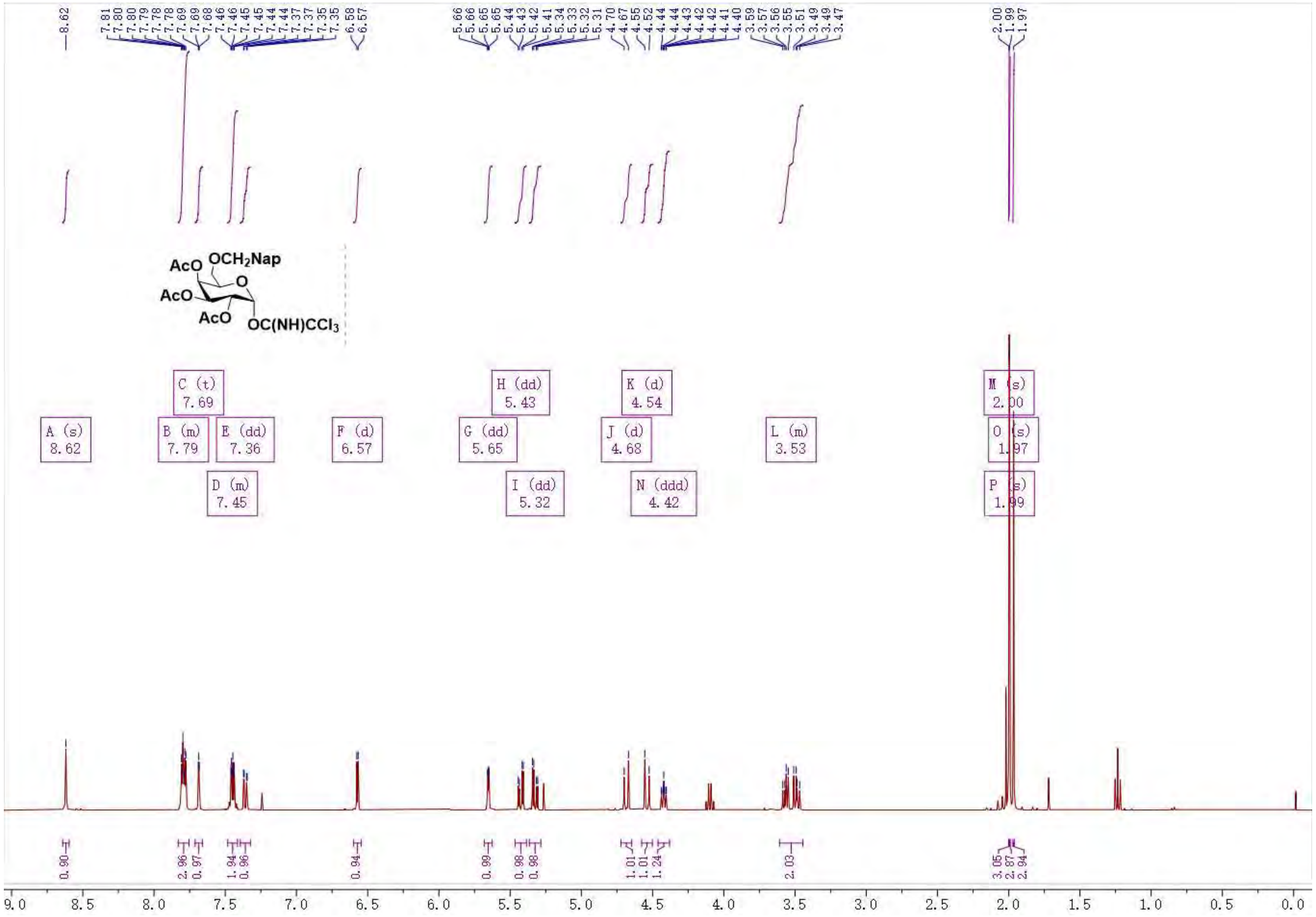

$^{13}$C NMR of **S6**. 100 MHz, $CDCl_3$

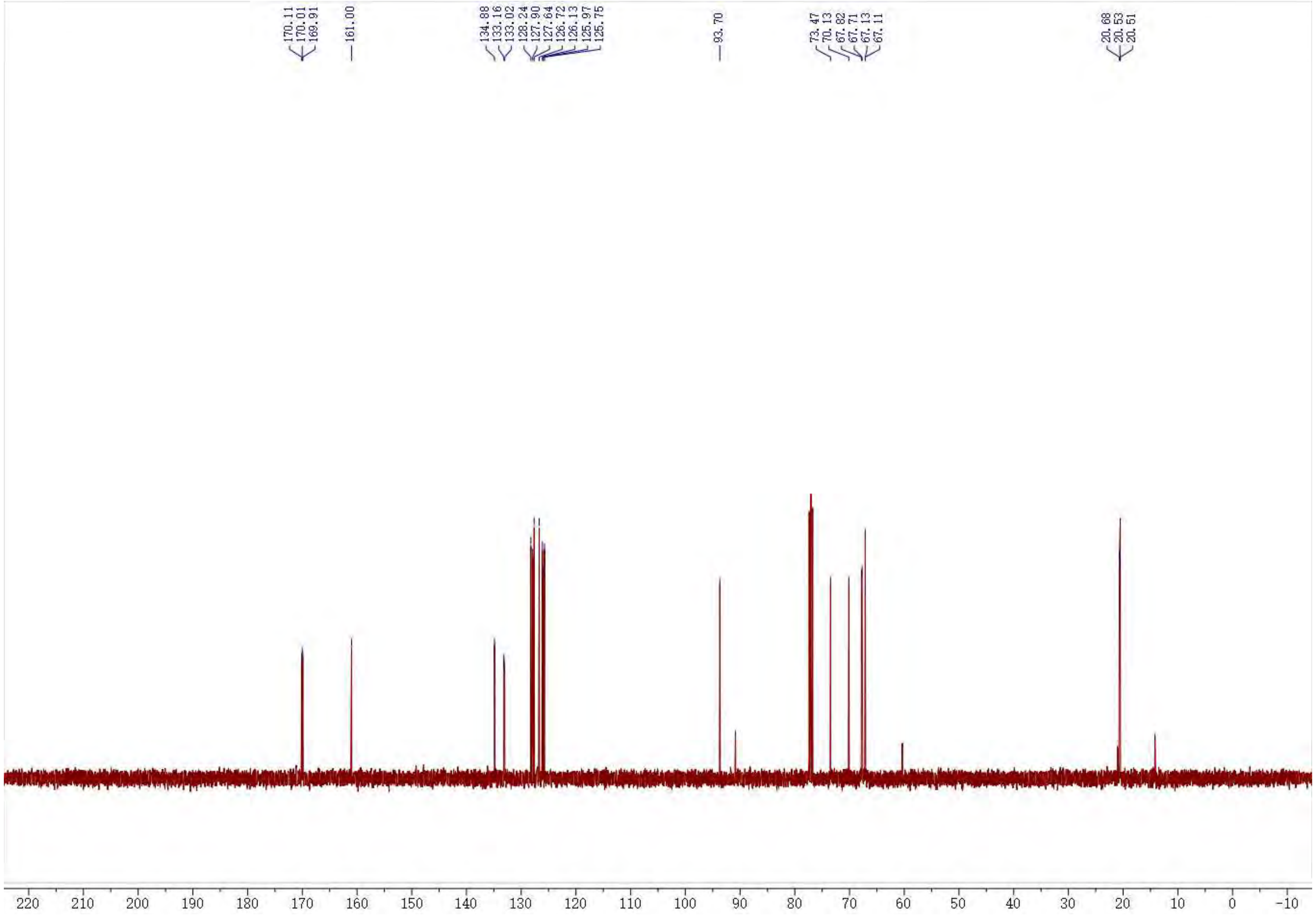

[1]H NMR of **S10**. 400 MHz, $CDCl_3$

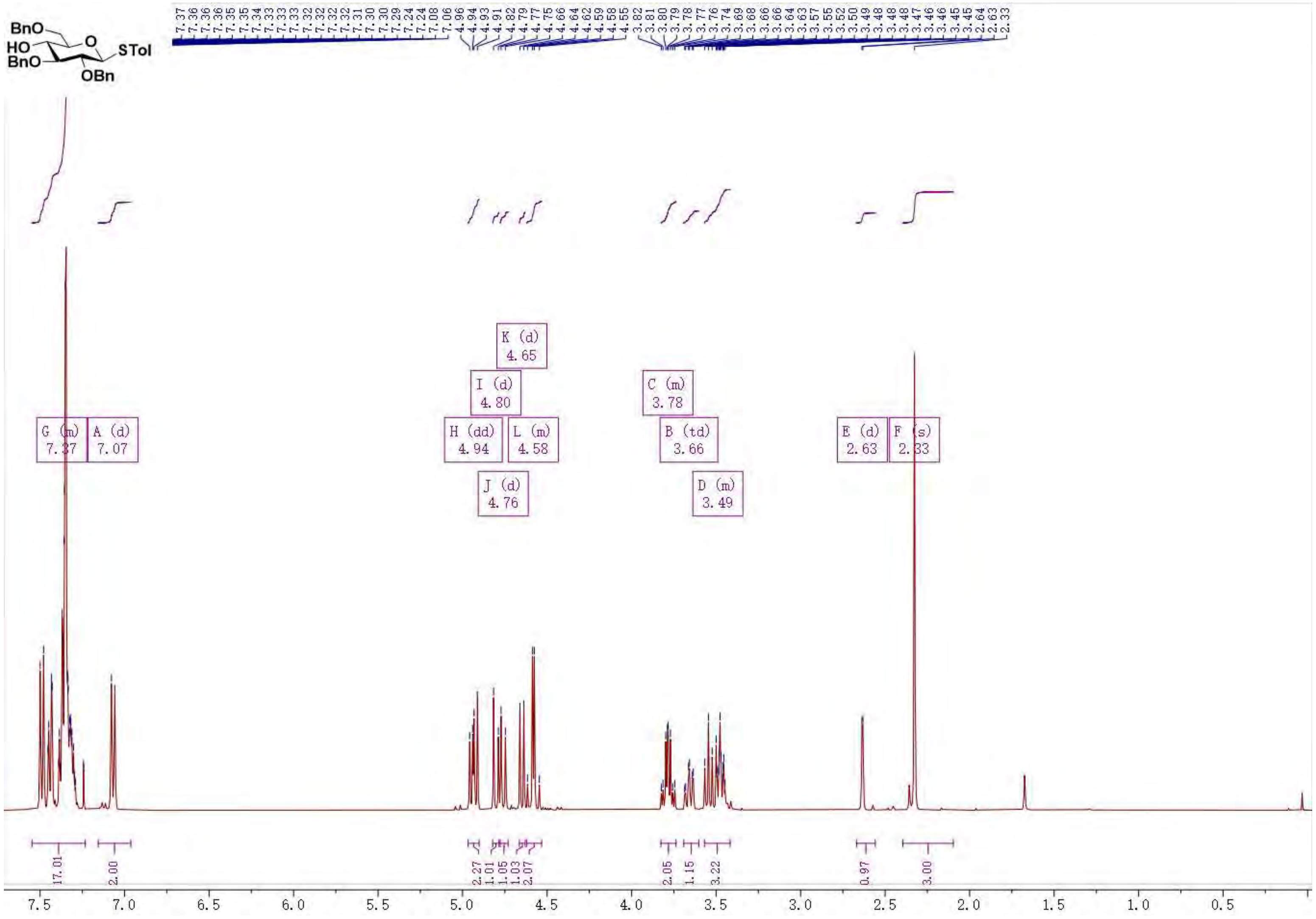

$^{13}$C NMR of **S10**. 100 MHz, $CDCl_3$

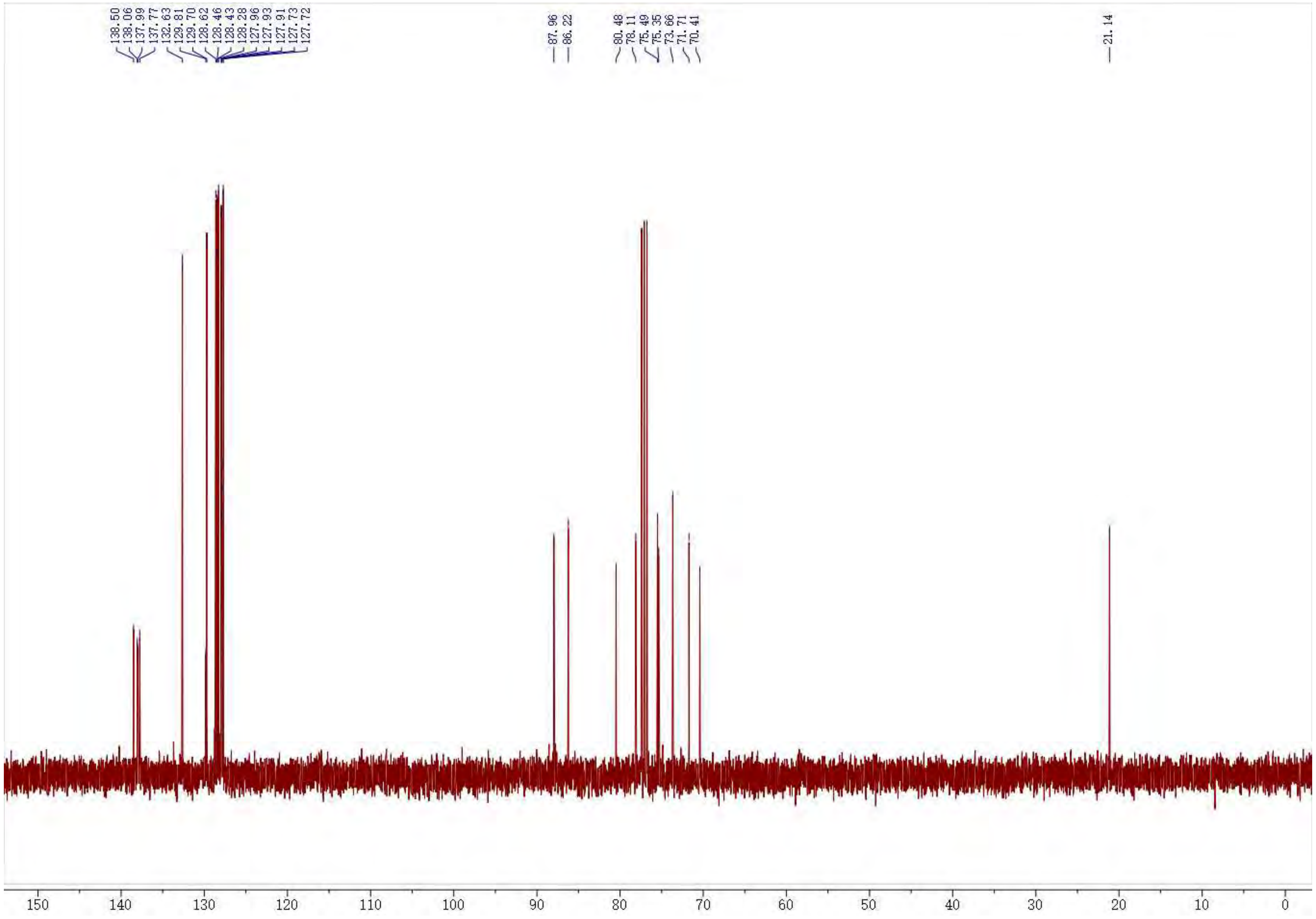

$^1$H NMR of **S12**. 400 MHz, $CDCl_3$

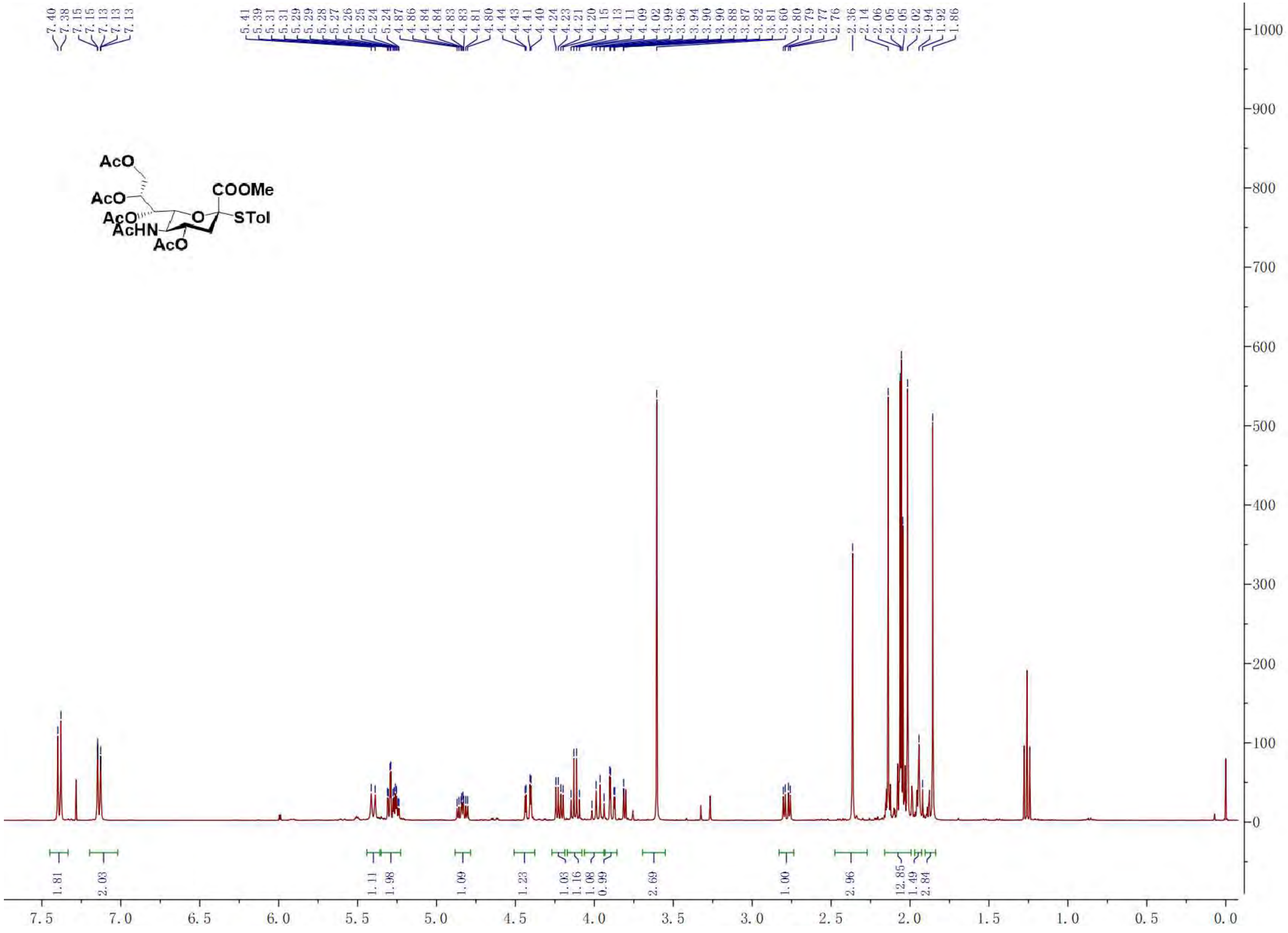

$^{13}$C NMR of **S12**. 100 MHz, $CDCl_3$

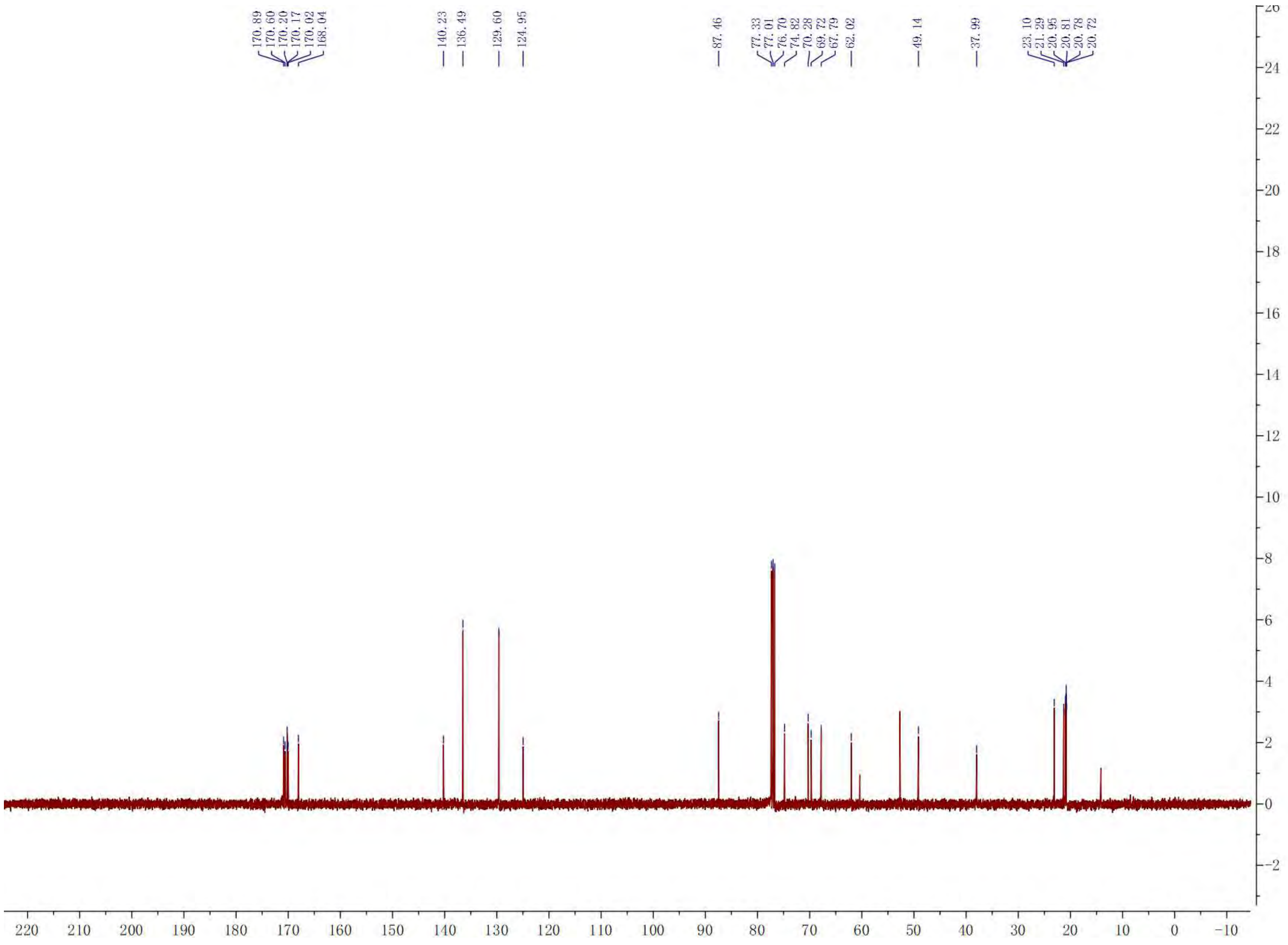

[1]H NMR of **S13**. 400 MHz, $CDCl_3$

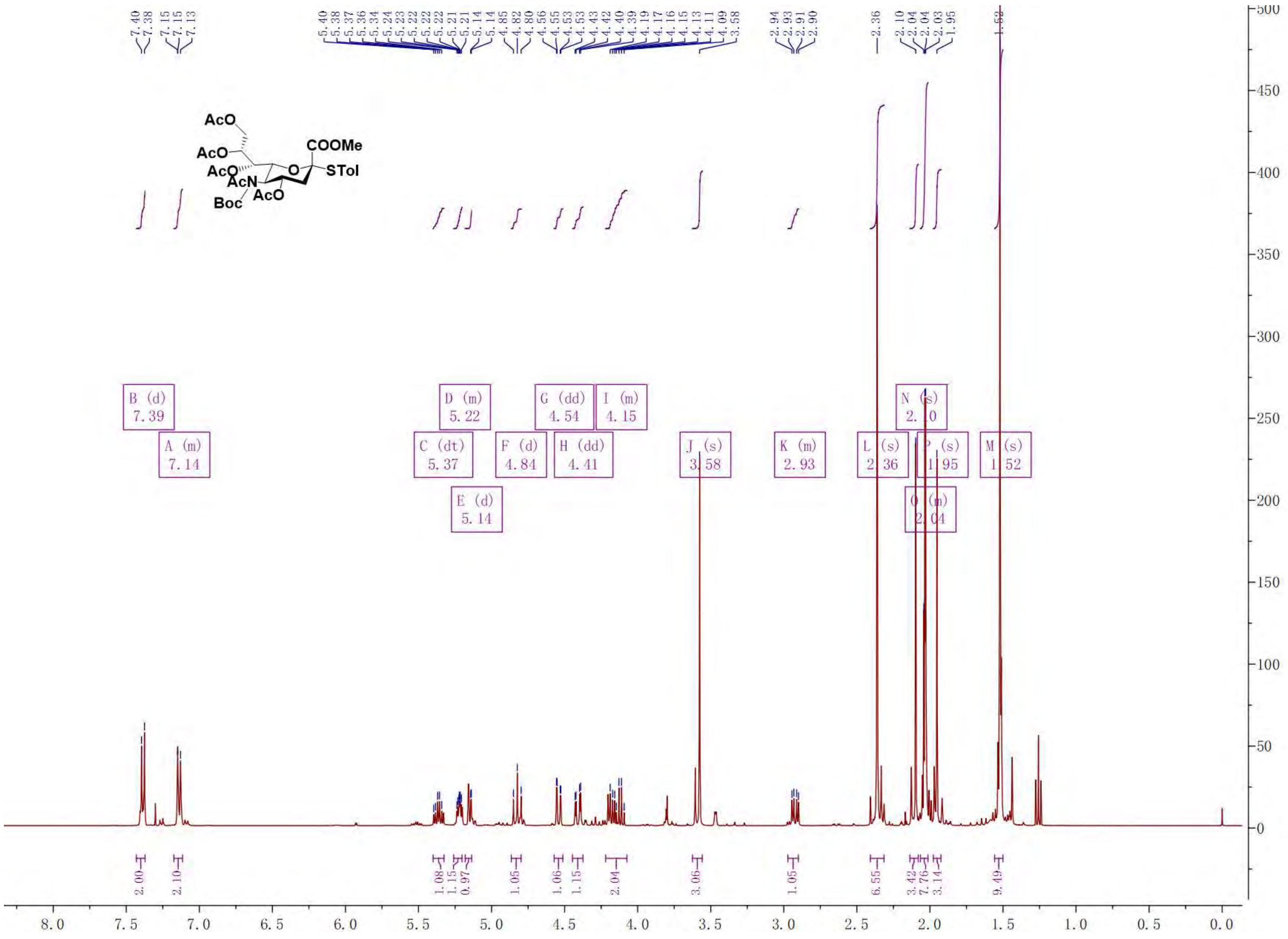

$^{13}$C NMR of **S13**. 100 MHz, $CDCl_3$

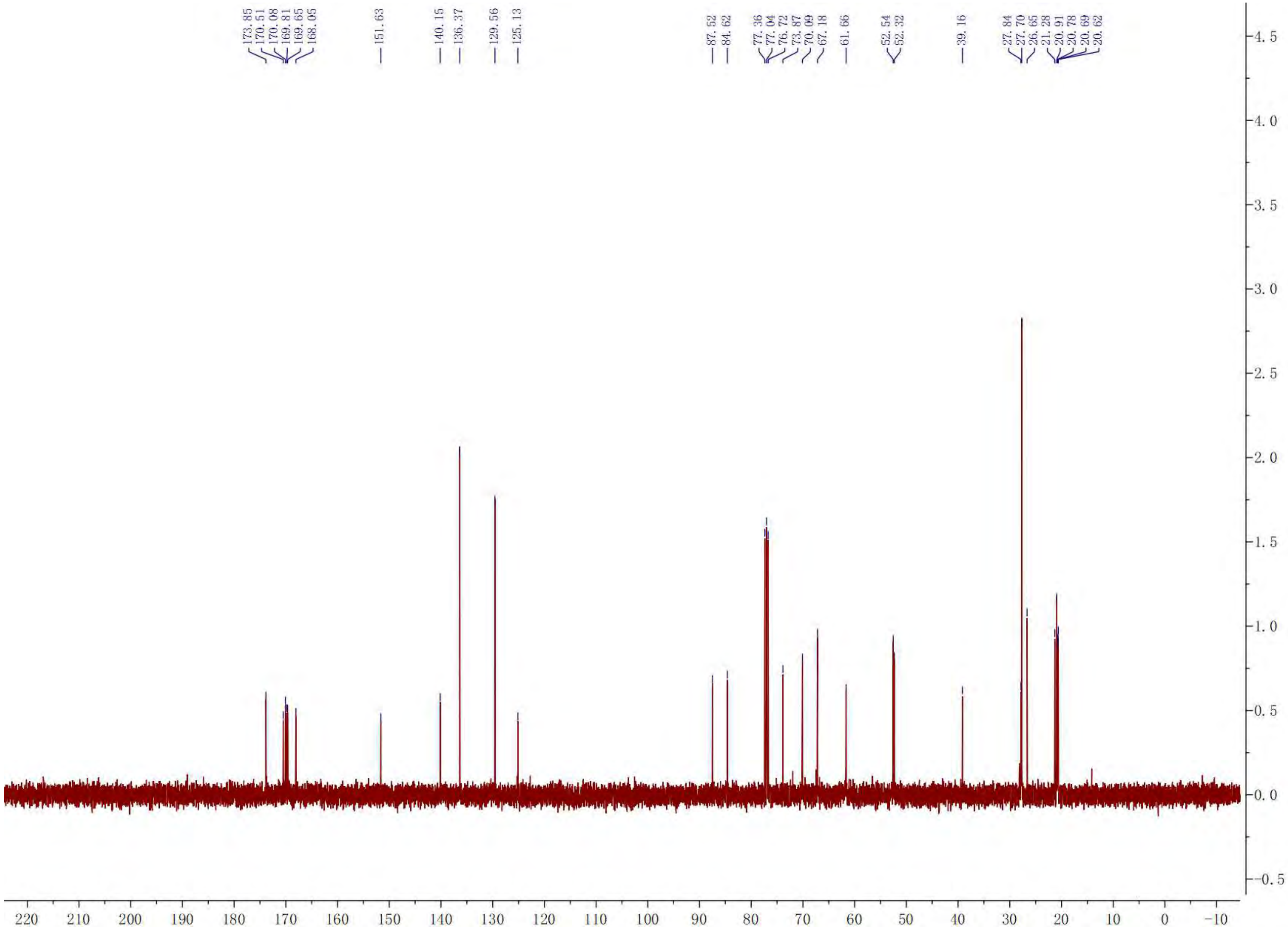

$^{1}$H NMR of **S14**. 400 MHz, $CDCl_3$

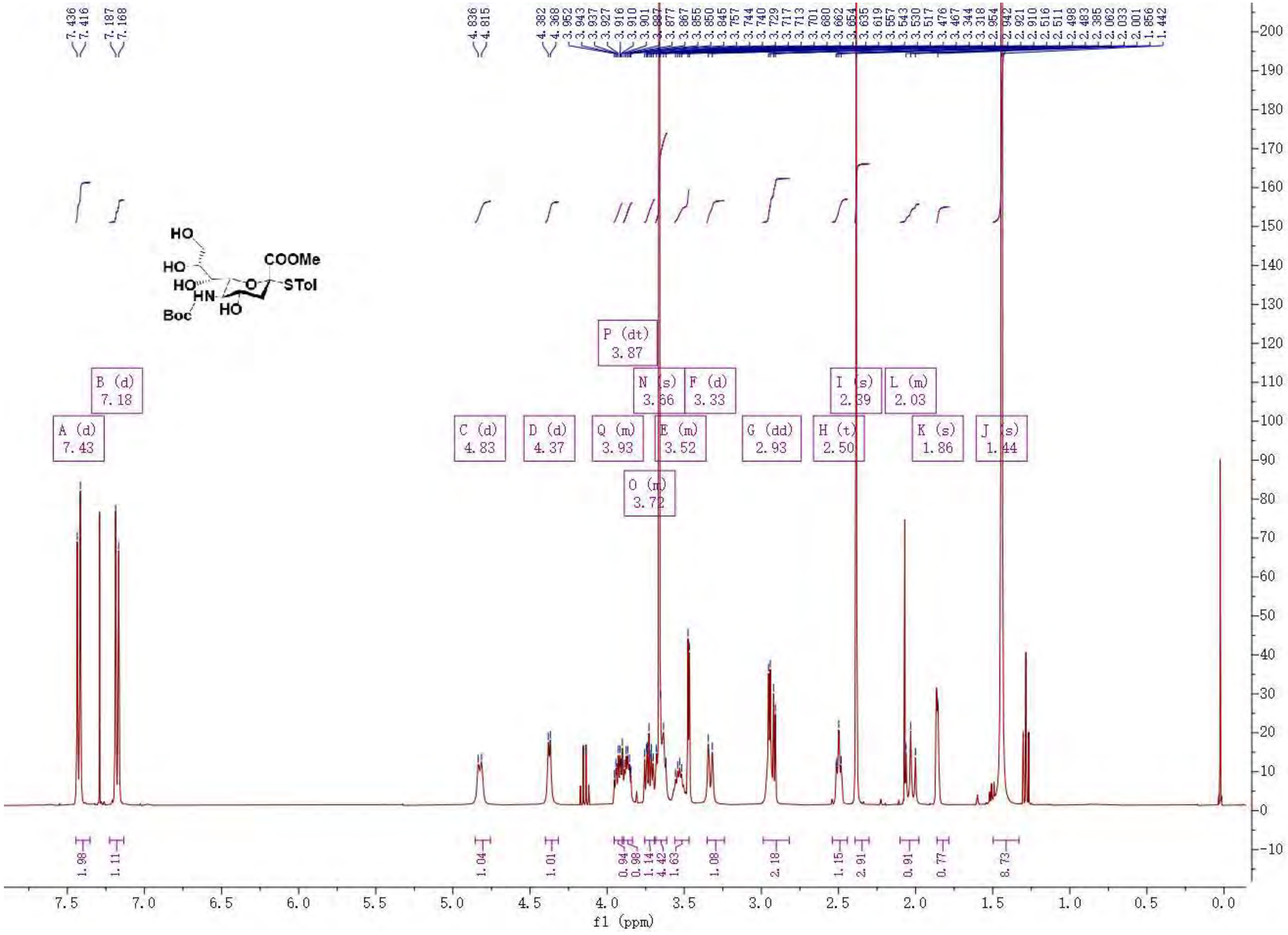

$^{13}C$ NMR of **S14**. 100 MHz, $CDCl_3$

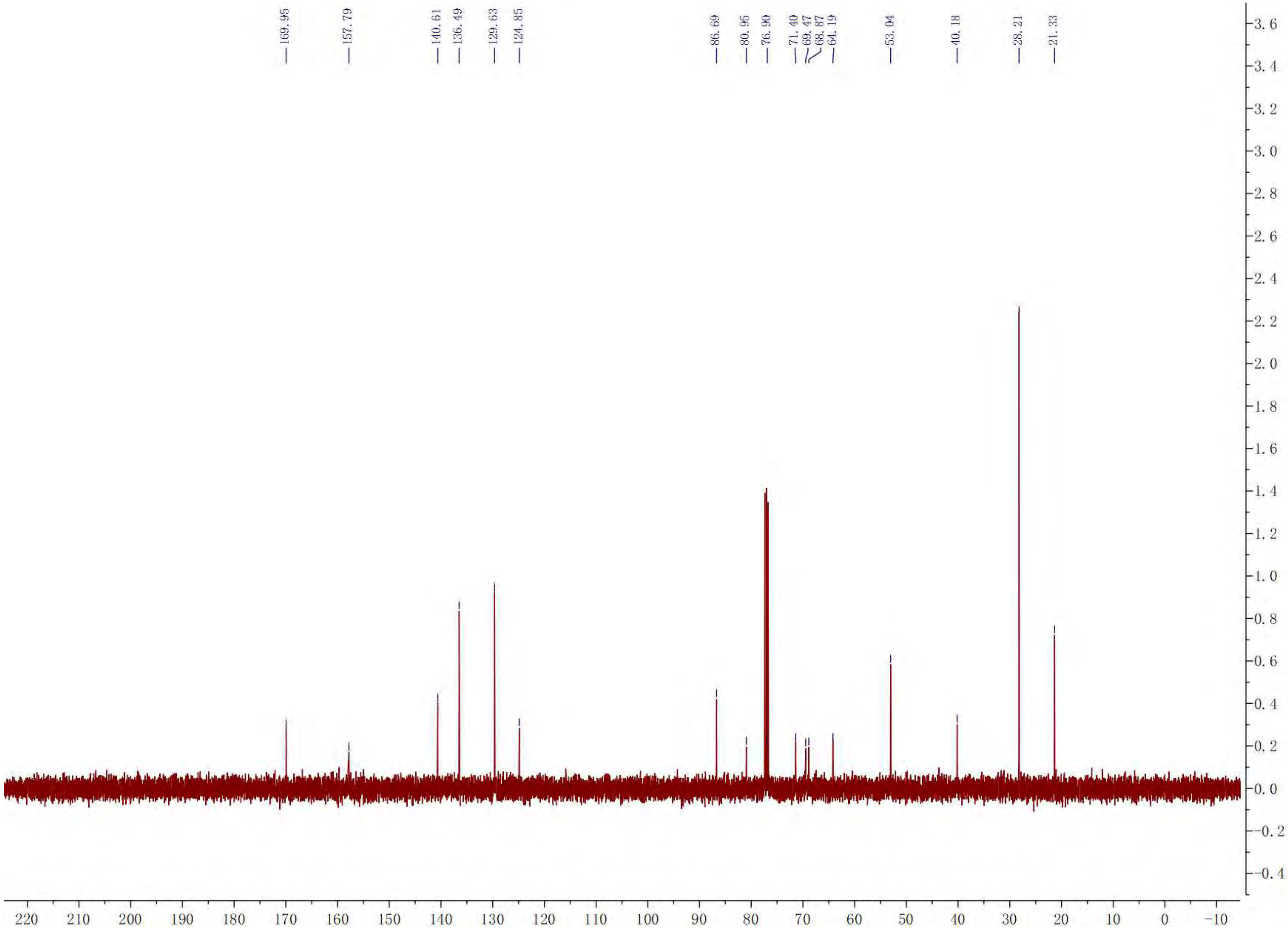

[1]H NMR of **S15**. 400 MHz, $CDCl_3$

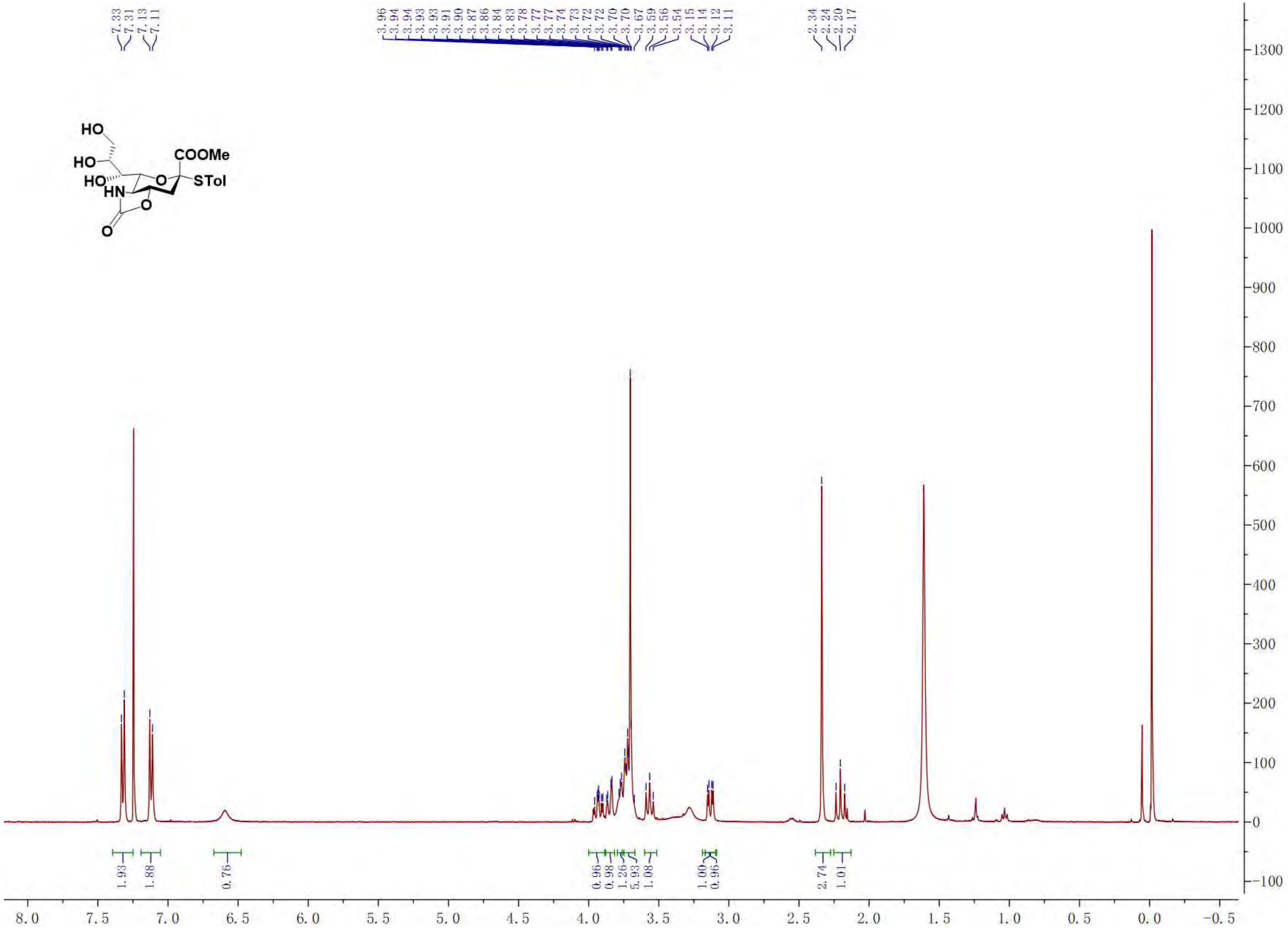

[13]C NMR of **S15**. 100 MHz, $CDCl_3$

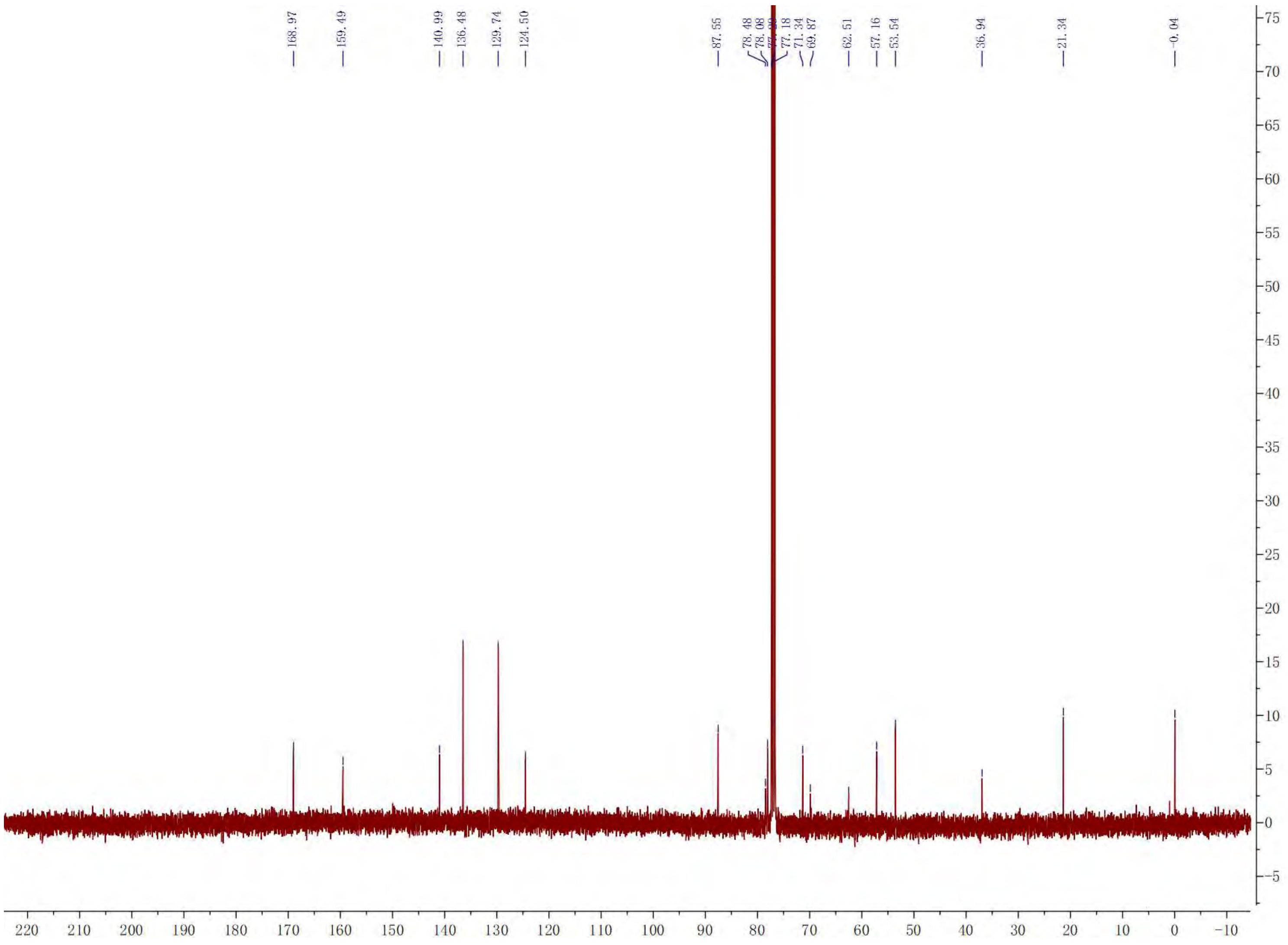

$^{1}$H NMR of **S16**. 400 MHz, $CDCl_3$

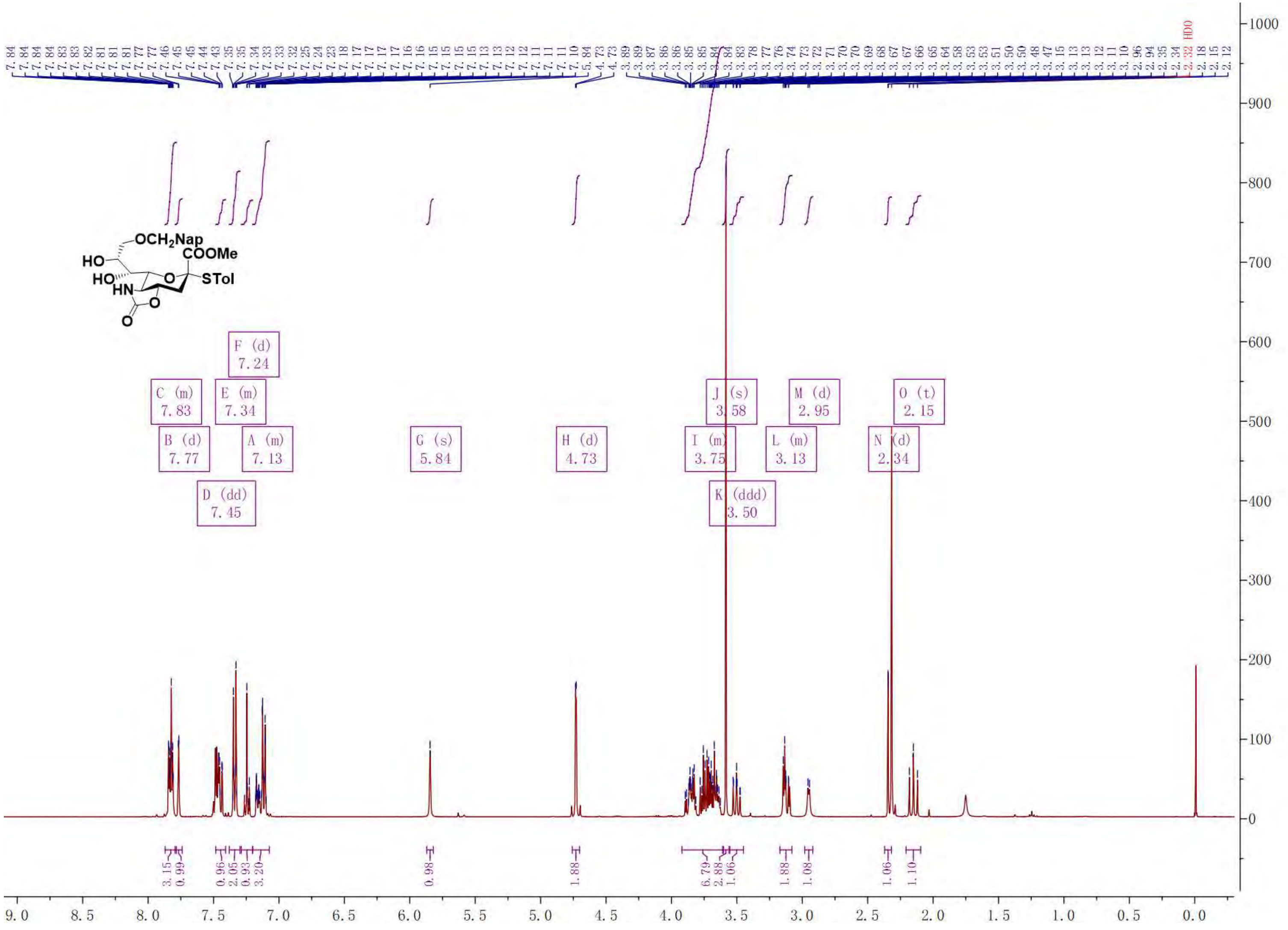

$^{13}C$ NMR of **S16**. 100 MHz, $CDCl_3$

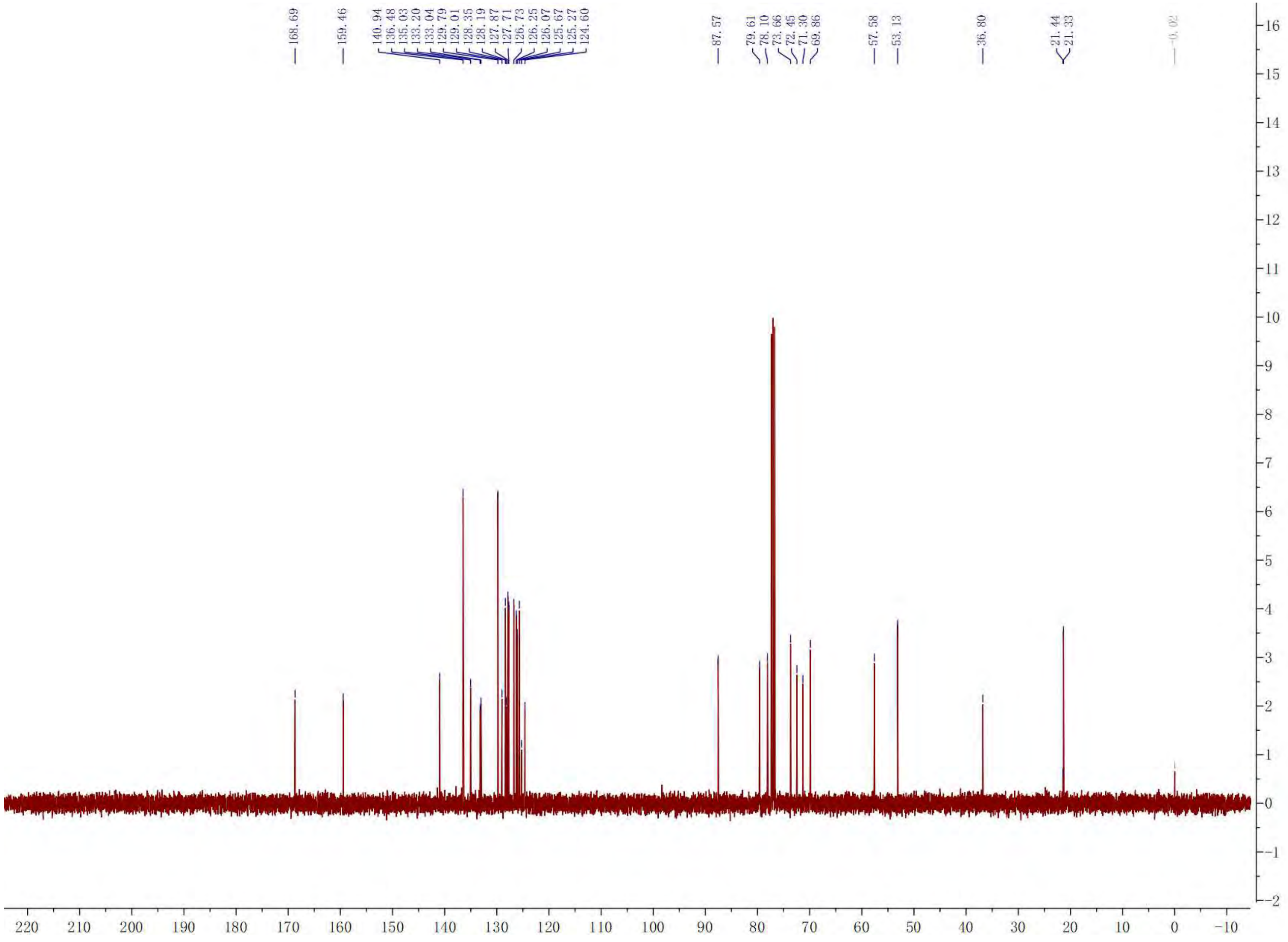

[1]H NMR of **S17**. 400 MHz, $CDCl_3$

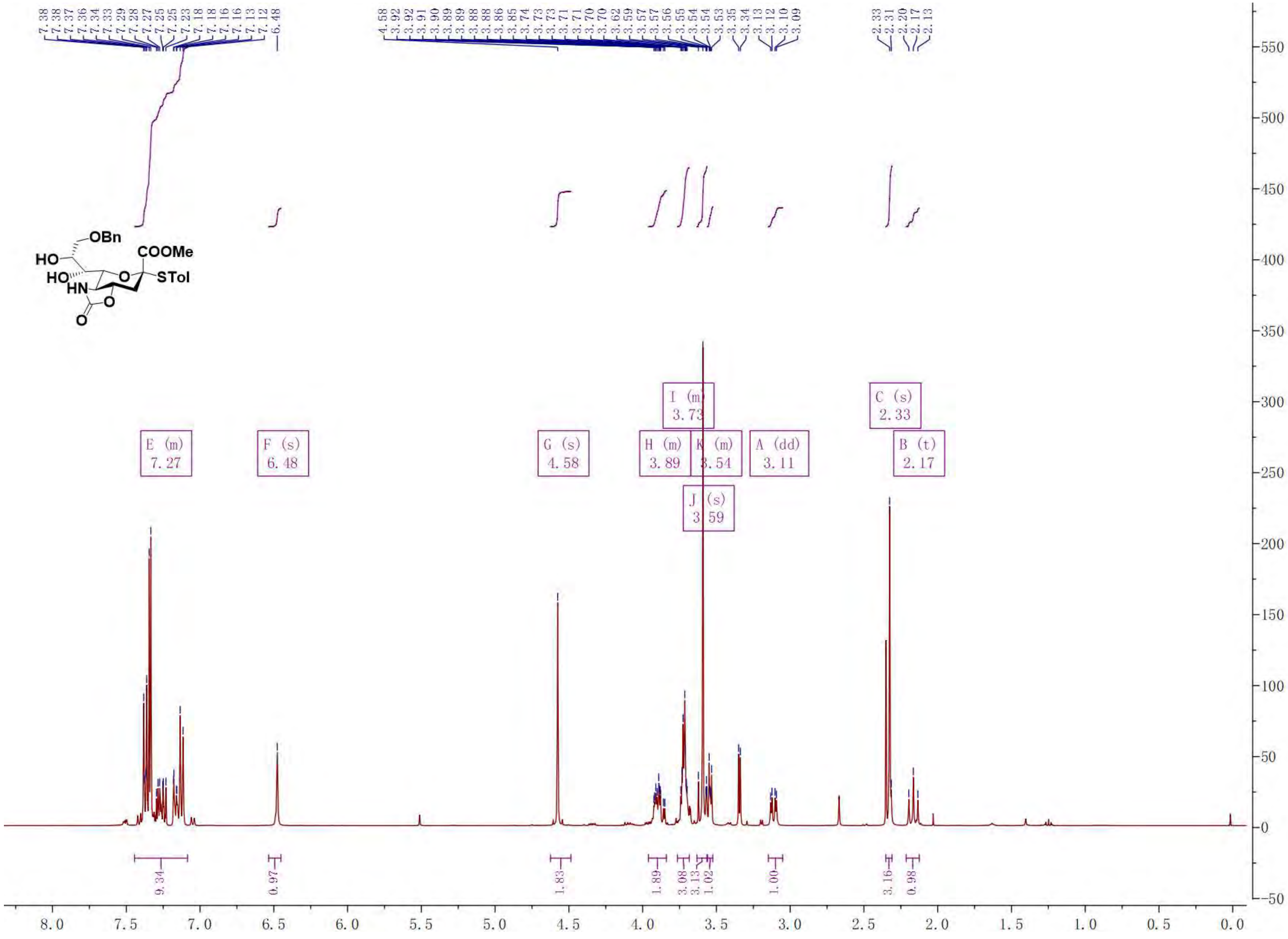

[13]C NMR of **S17**. 100 MHz, $CDCl_3$

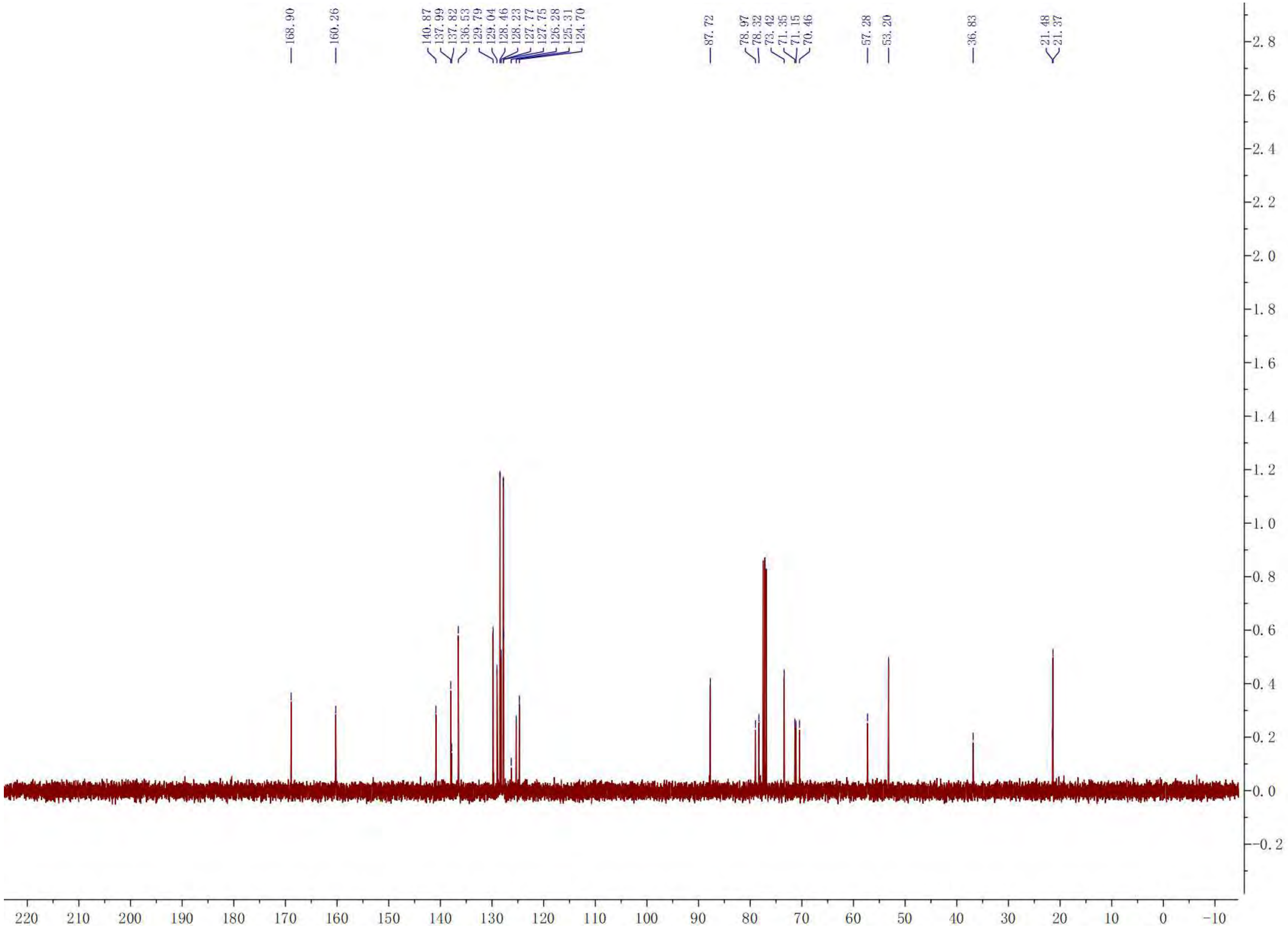

[1]H NMR of **S18**. 400 MHz, $CDCl_3$

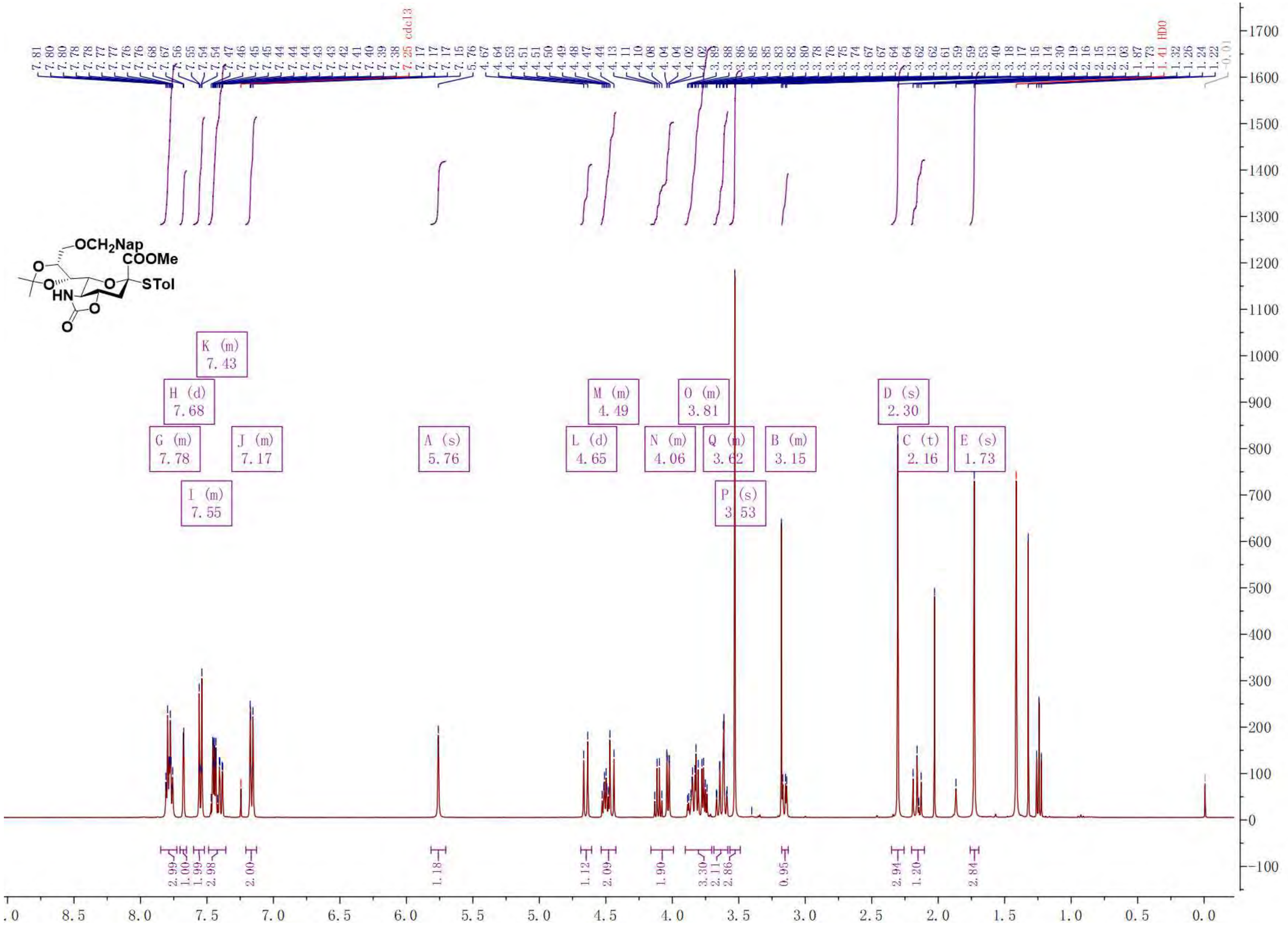

$^{13}$C NMR of **S18**. 100 MHz, $CDCl_3$

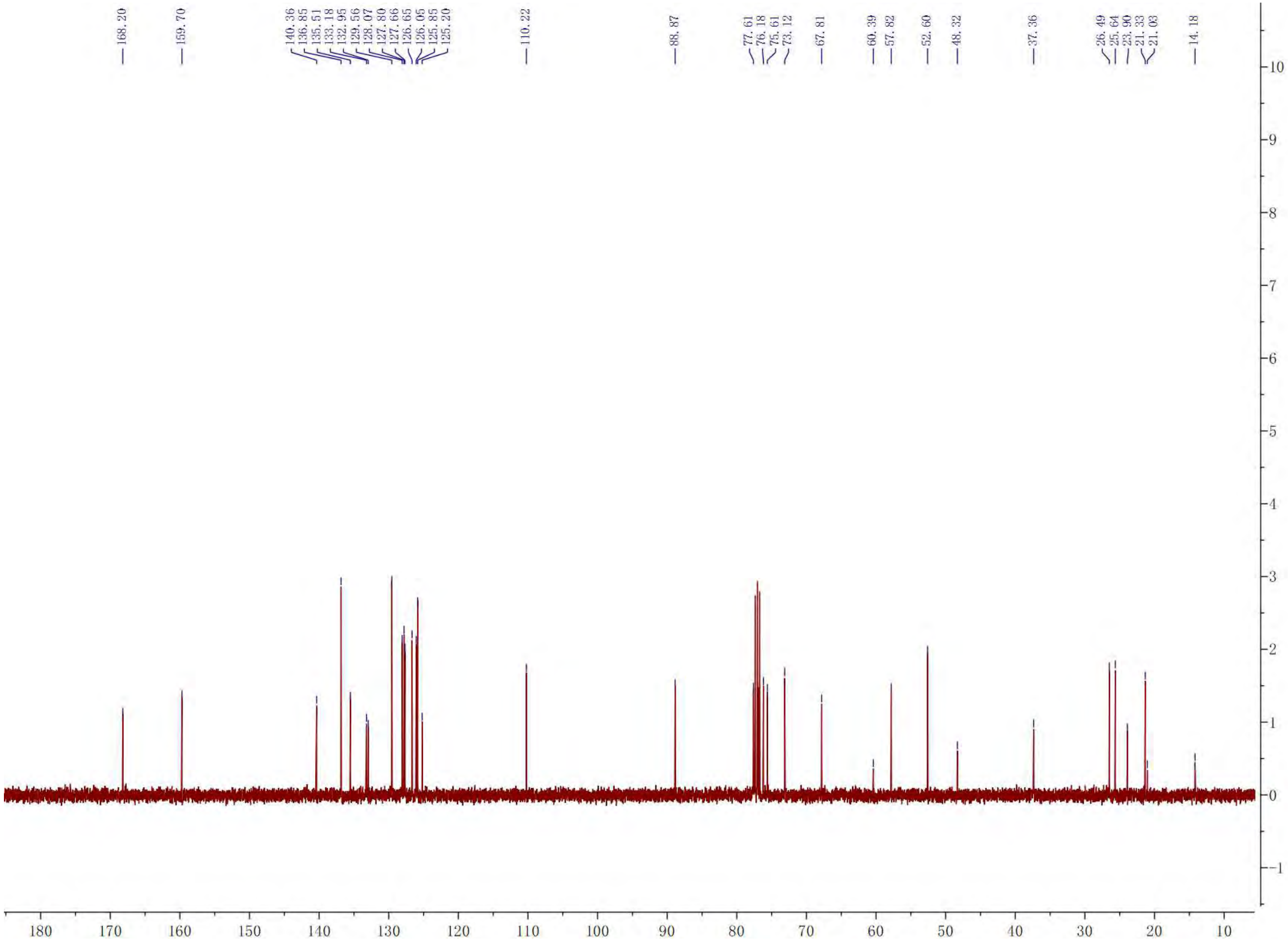

[1]H NMR of **S19**. 400 MHz, $CDCl_3$

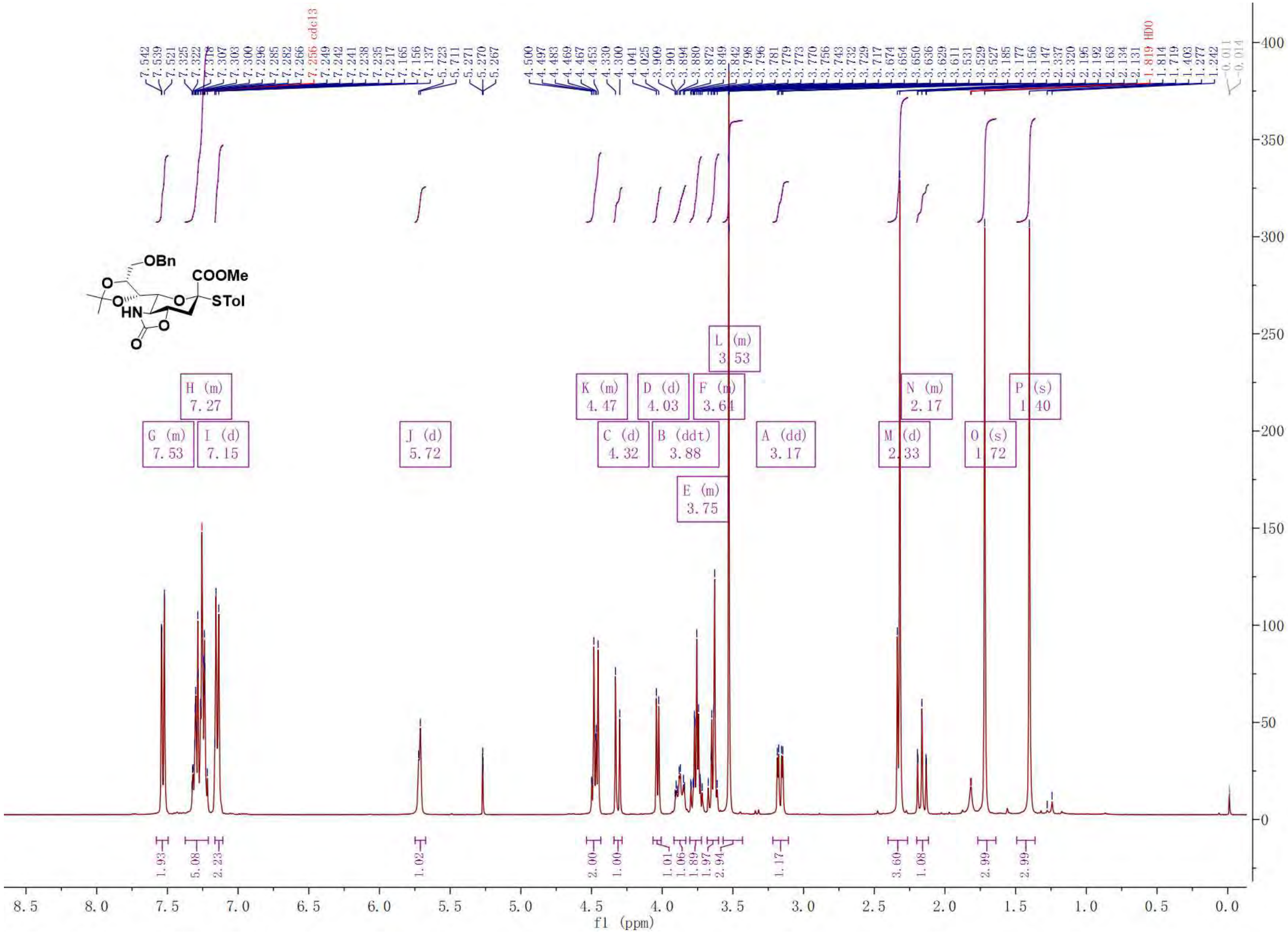

$^{13}C$ NMR of **S19**. 100 MHz, $CDCl_3$

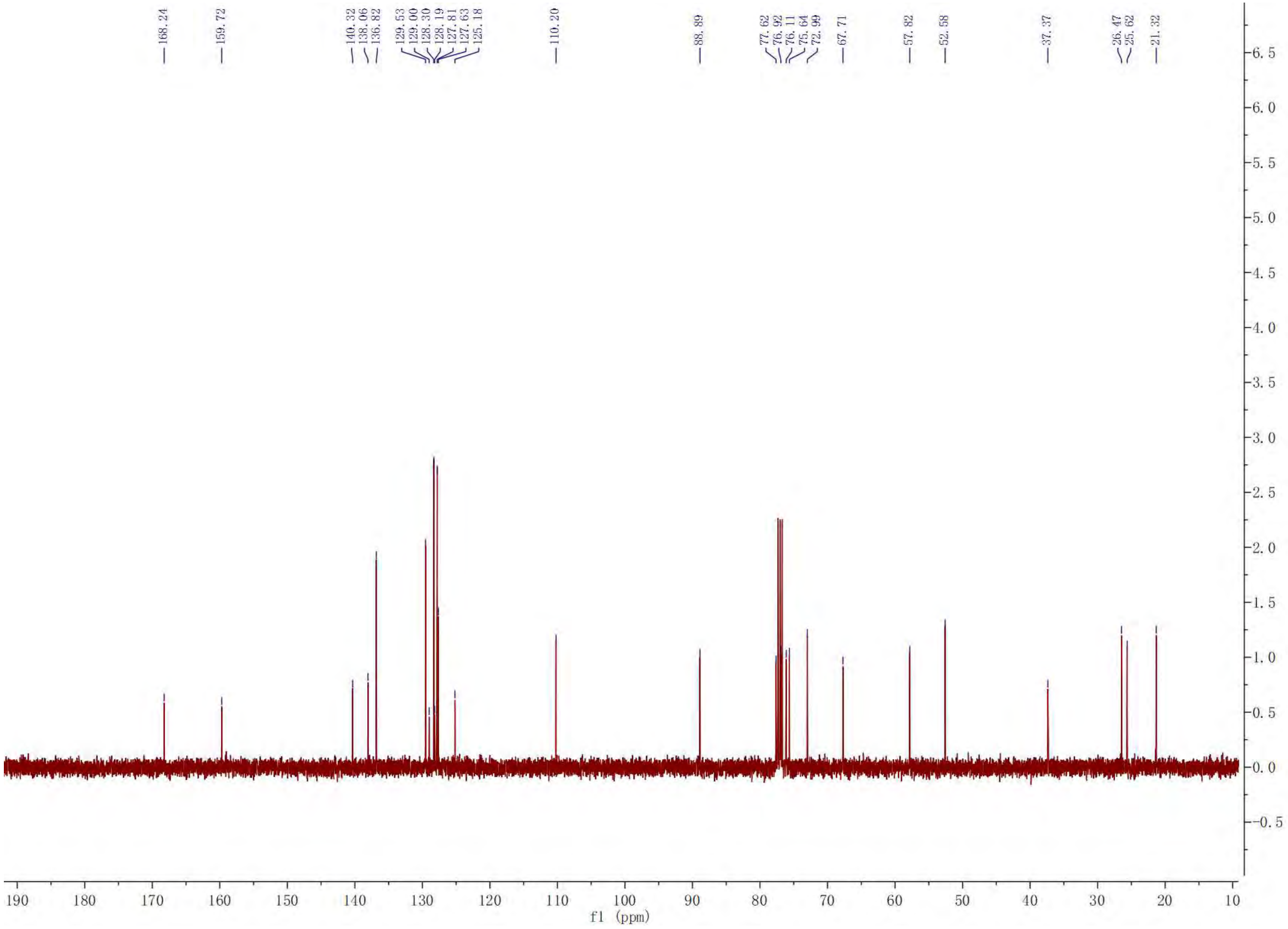

[1]H NMR of **9**. 400 MHz, $CDCl_3$

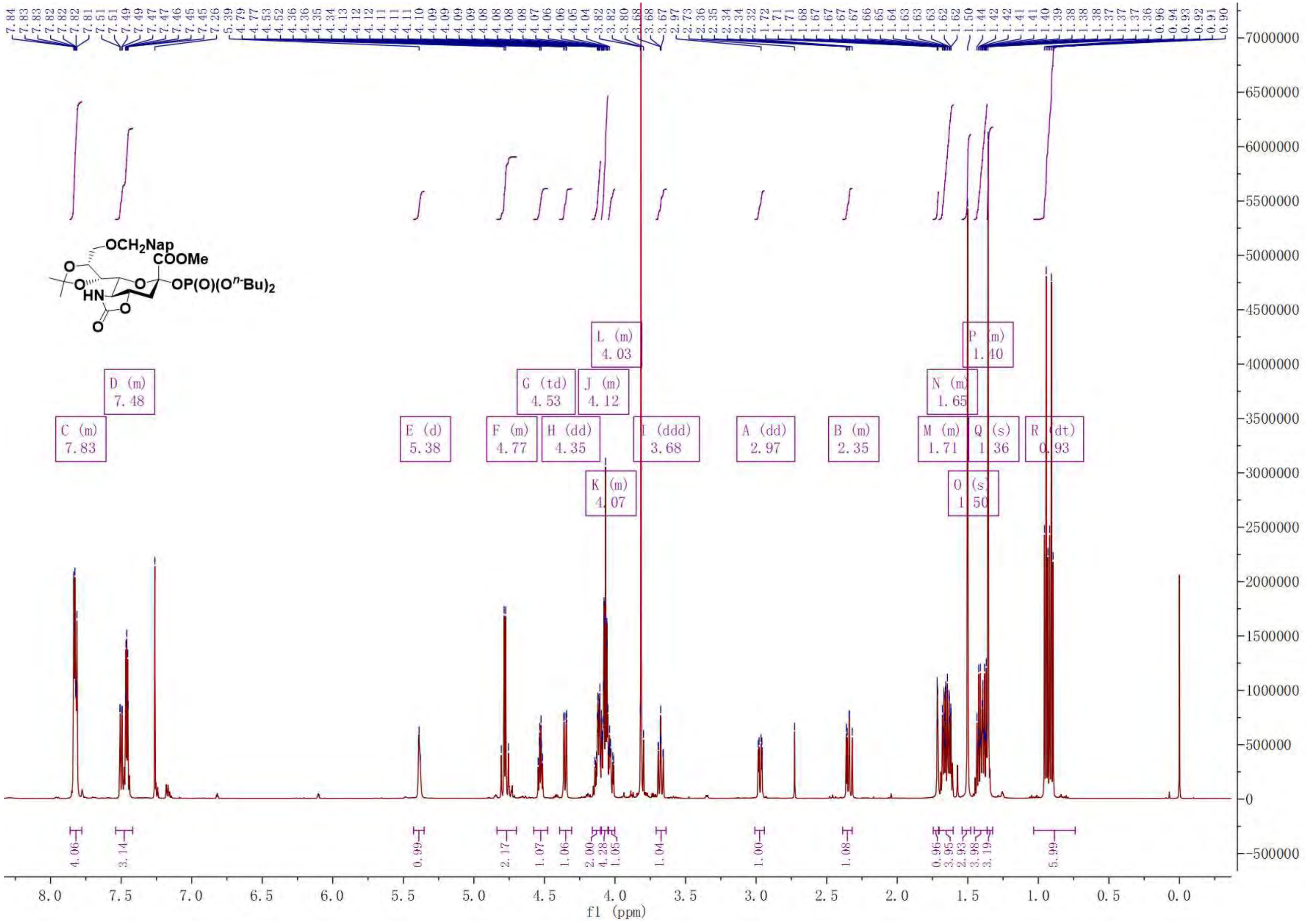

[13]C NMR of **9**. 100 MHz, $CDCl_3$

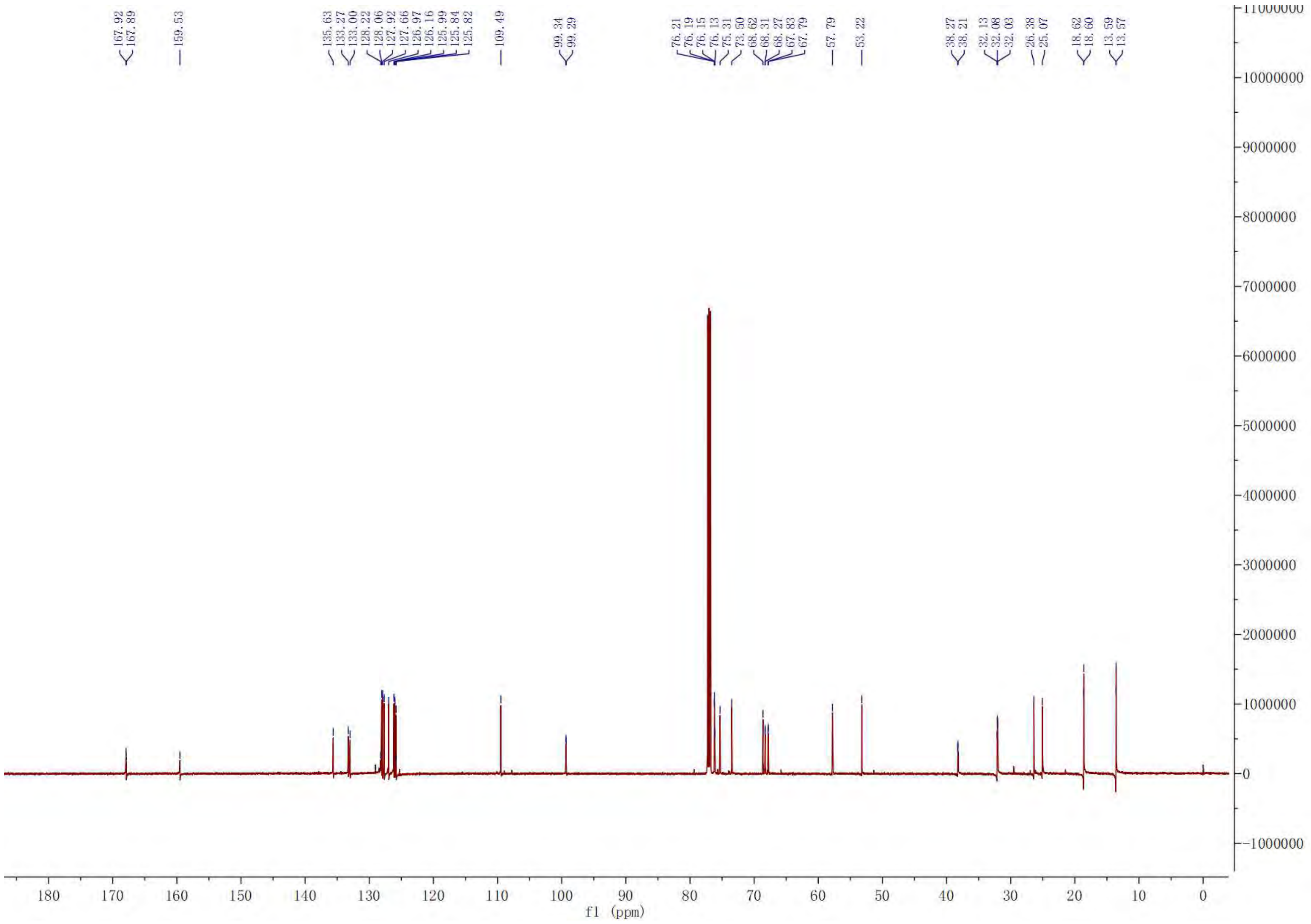

[1]H NMR of **10**. 400 MHz, $CDCl_3$

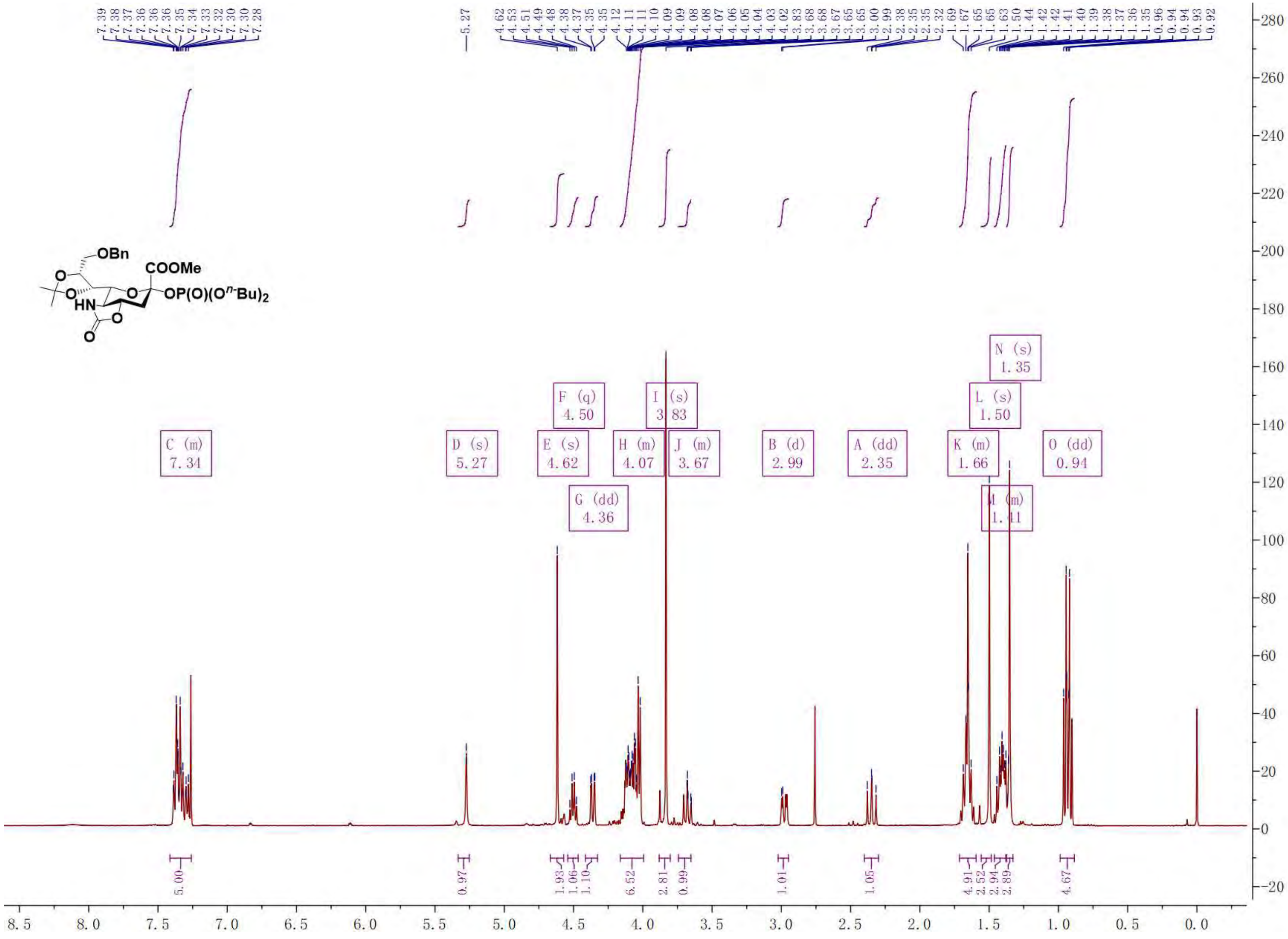

$^{13}$C NMR of **10**. 100 MHz, $CDCl_3$.

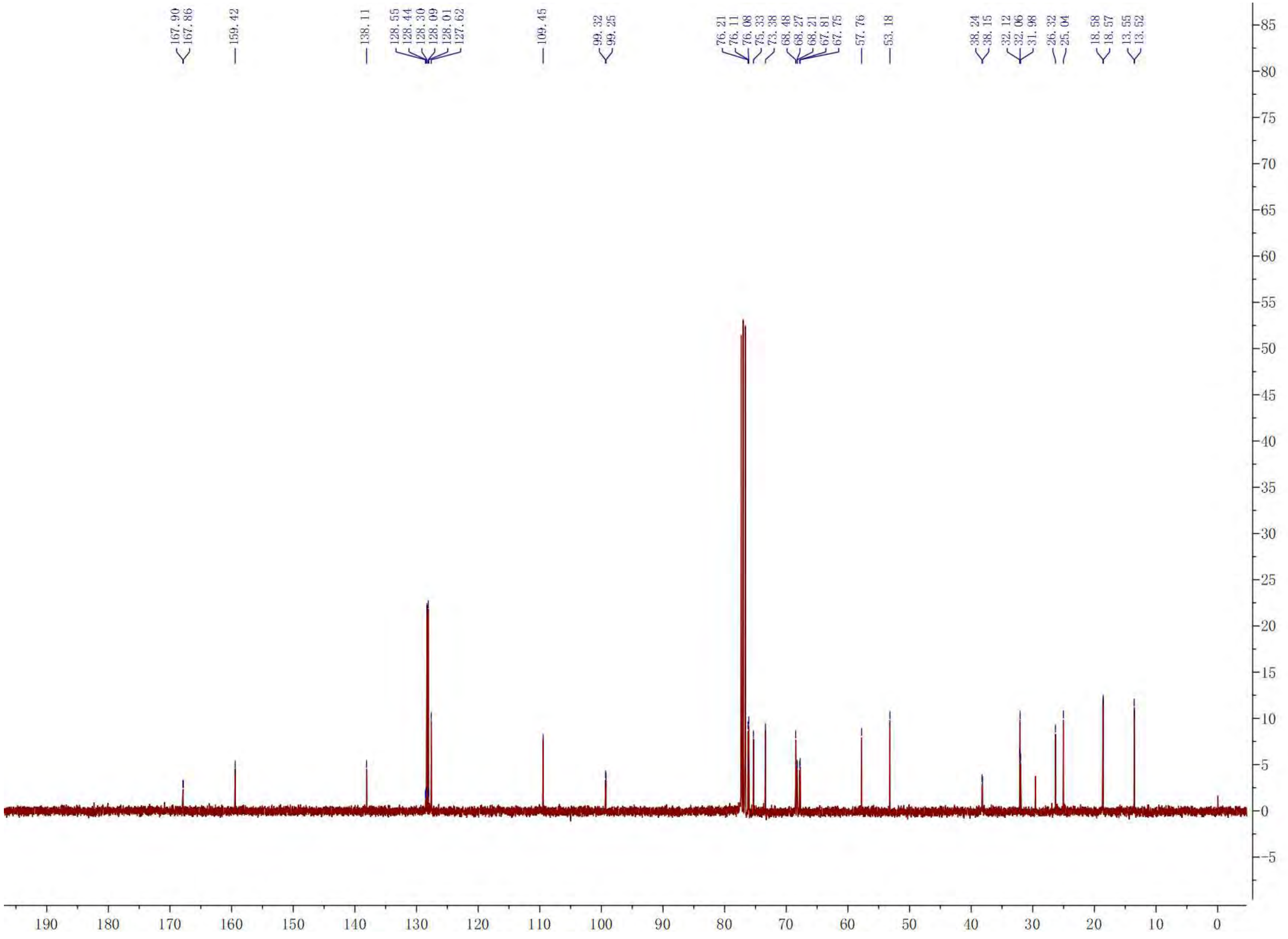

<sup></sup>[1]H NMR of **11**. 400 MHz, $CDCl_3$.

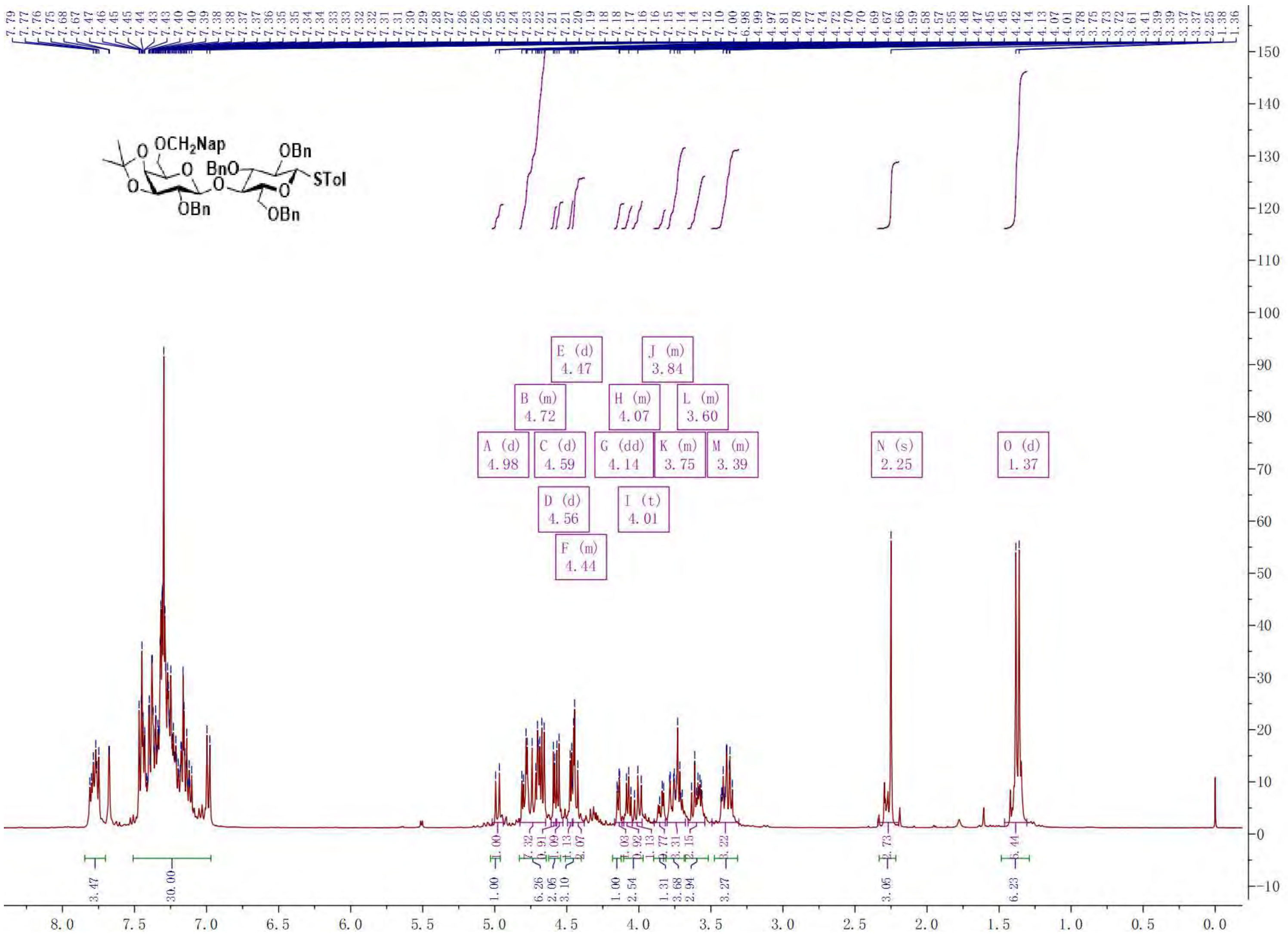

COSY of **11**

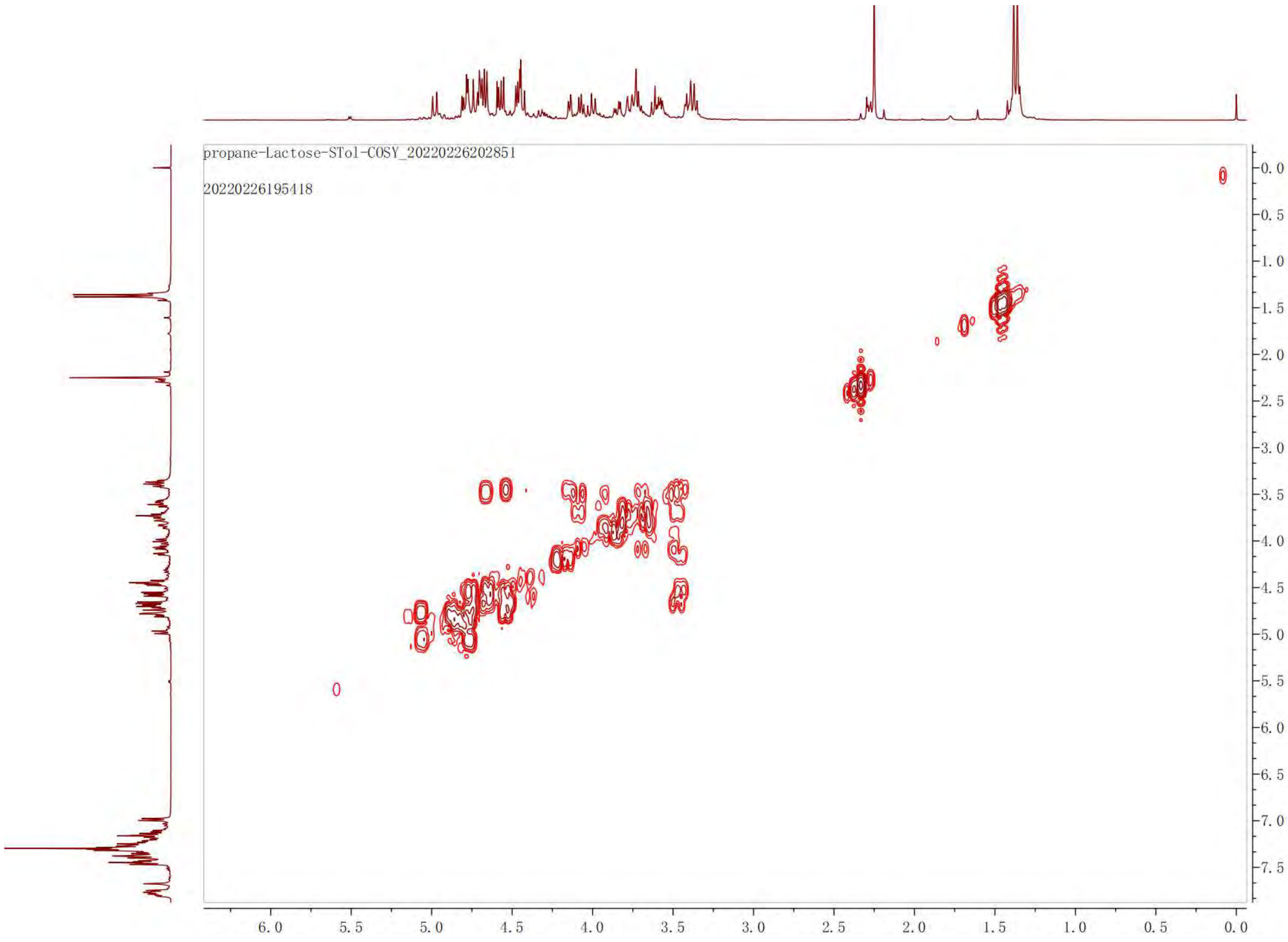
propane-Lactose-STol-COSY_20220226202851
20220226195418
6.0
5.5
5.0
4.5
4.0
3.5
3.0
2.5
2.0
1.5
1.0
0.5
0.0
-0.0
-0.5
-1.0
-1.5
-2.0
-2.5
-3.0
-3.5
-4.0
-4.5
-5.0
-5.5
-6.0
-6.5
-7.0
-7.5

TOCSY of **11**

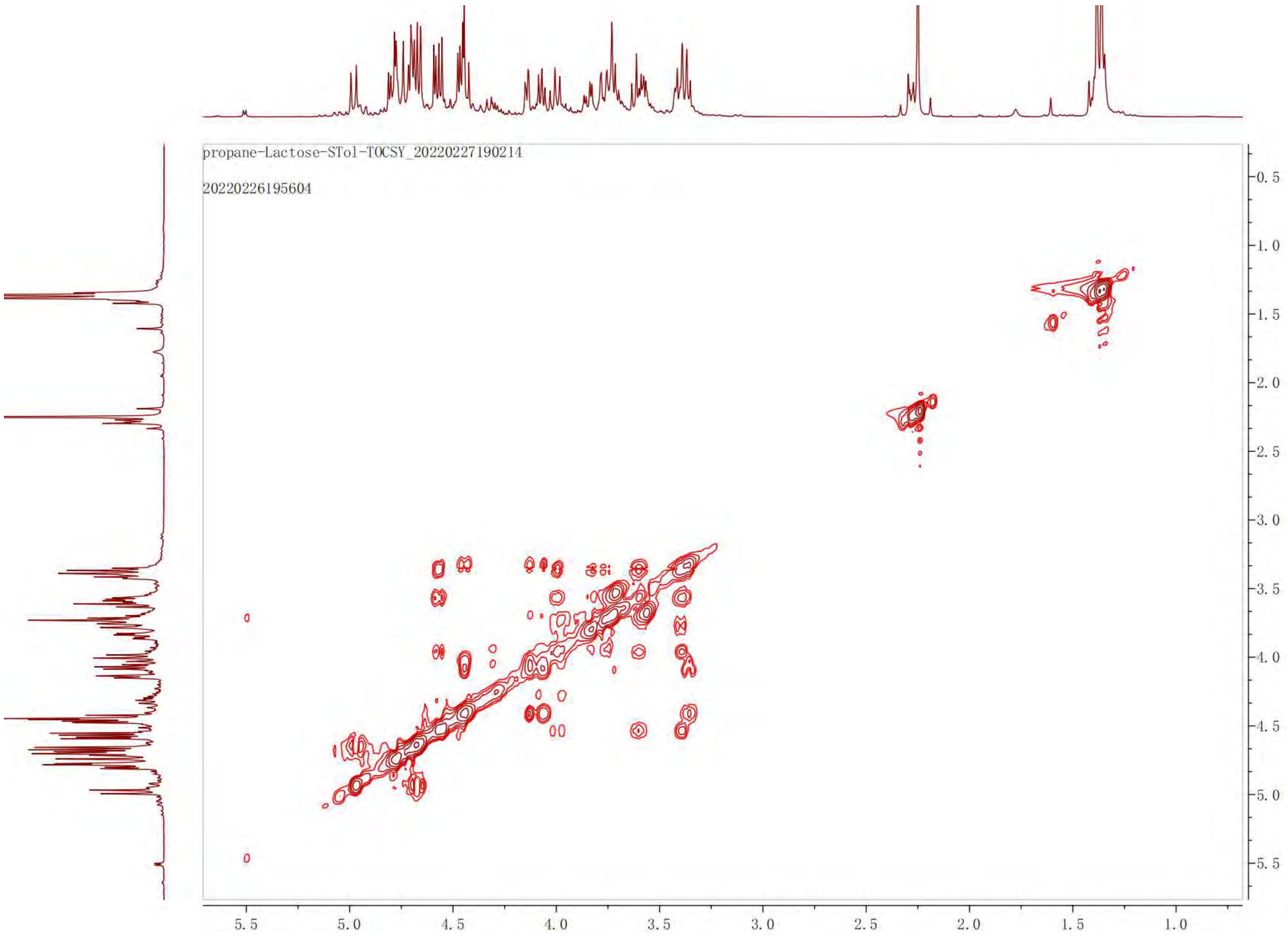

propane-Lactose-STol-TOCSY_20220227190214
20220226195604
5.5
5.0
4.5
4.0
3.5
3.0
2.5
2.0
1.5
1.0
0.5
1.0
1.5
2.0
2.5
3.0
3.5
4.0
4.5
5.0
5.5

HSQC of **11**

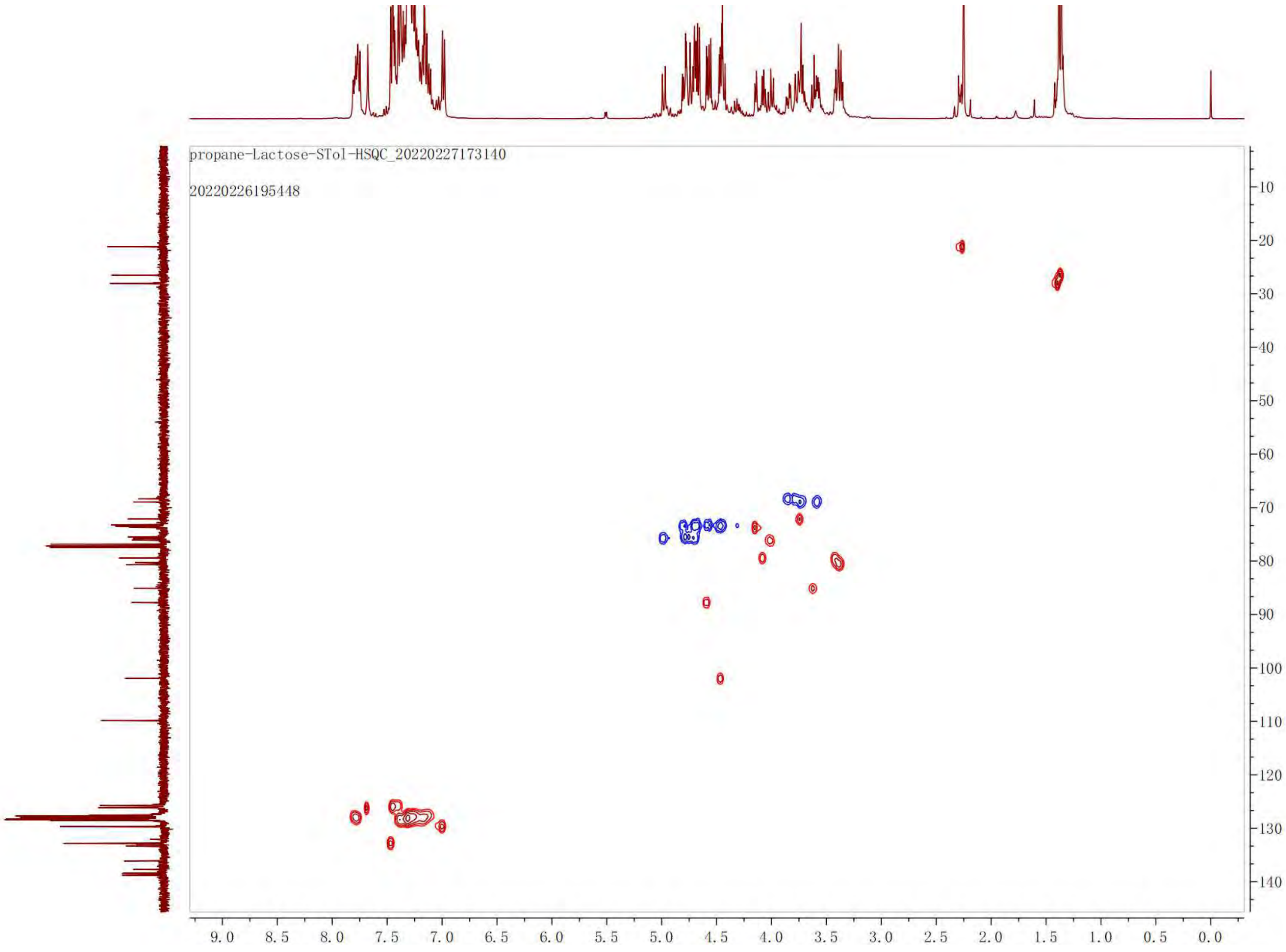

propane-Lactose-STol-HSQC_20220227173140
20220226195448
9.0 8.5 8.0 7.5 7.0 6.5 6.0 5.5 5.0 4.5 4.0 3.5 3.0 2.5 2.0 1.5 1.0 0.5 0.0
10 20 30 40 50 60 70 80 90 100 110 120 130 140

$^{13}$C NMR of **11**. 100 MHz, $CDCl_3$.

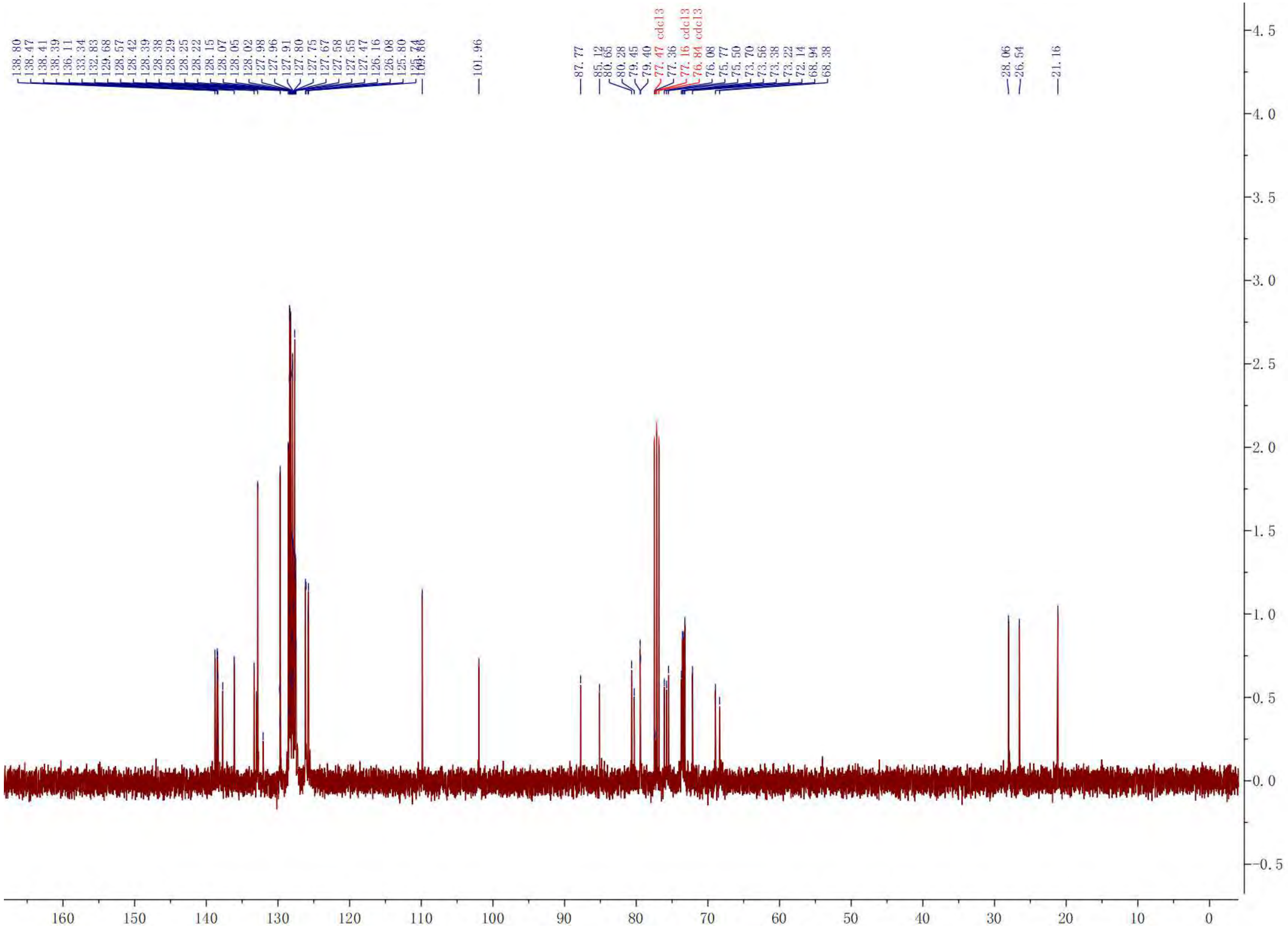

[1]H NMR of **12**. 400 MHz, $CDCl_3$.

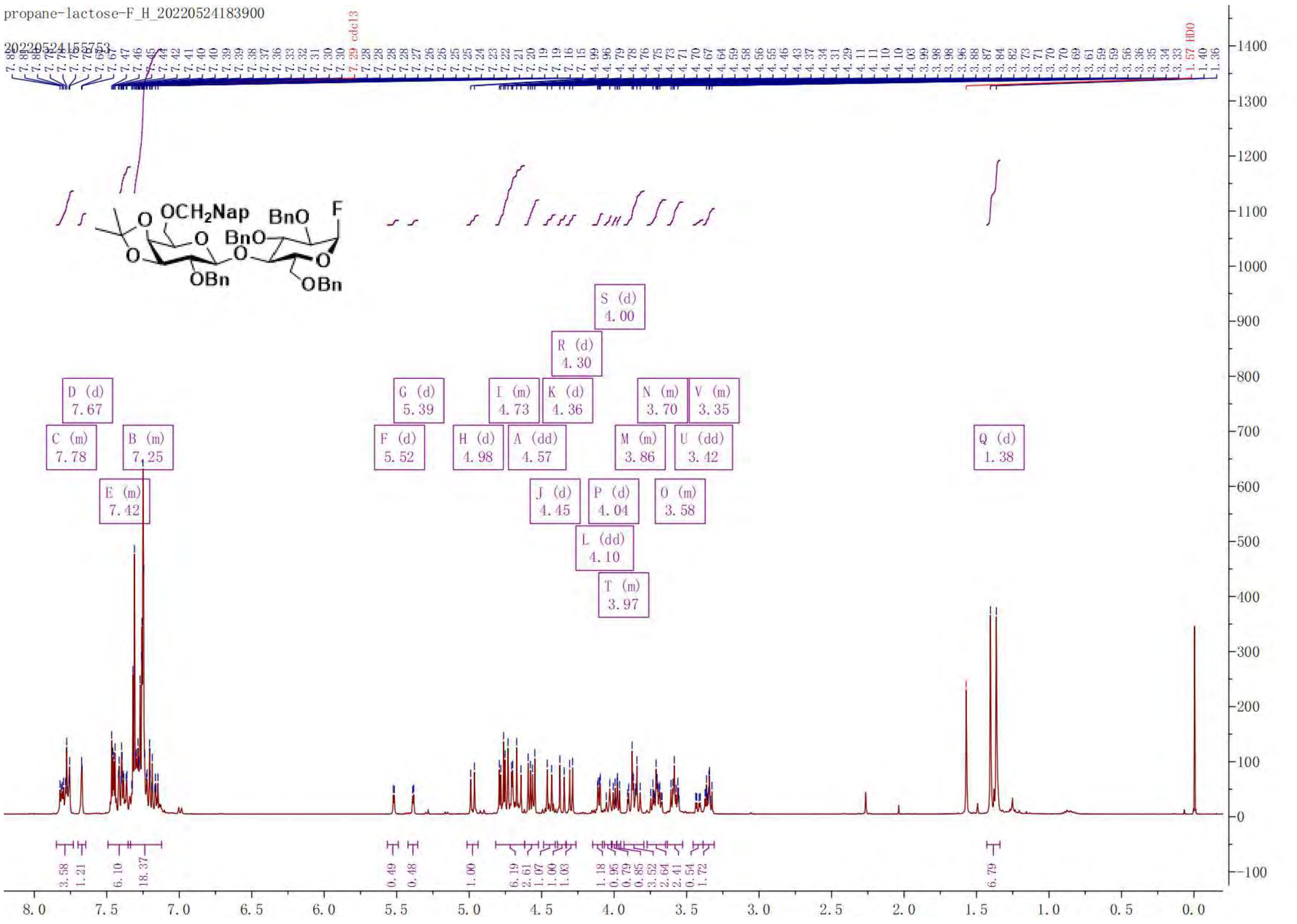

COSY of **12**

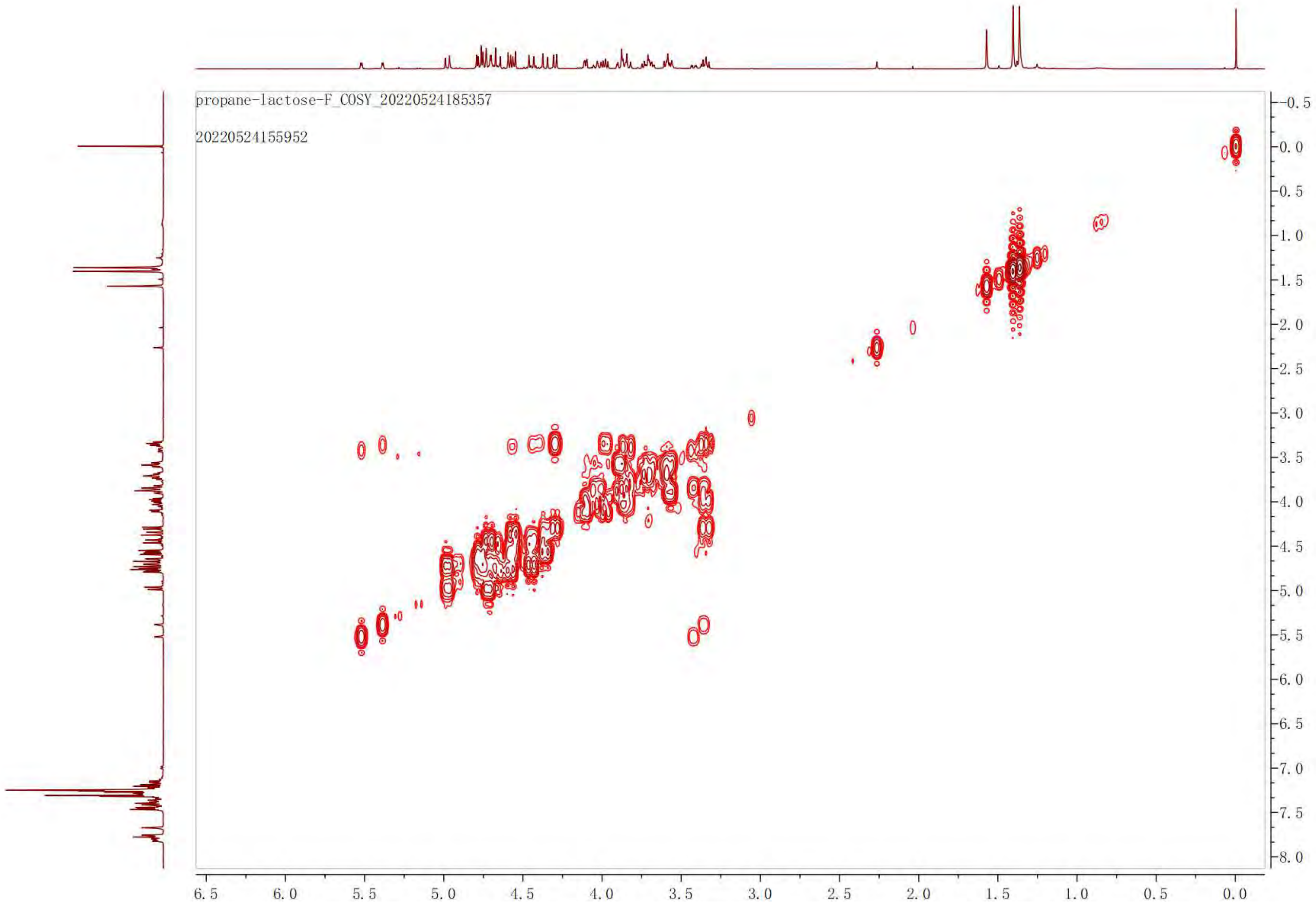

propane-lactose-F_COSY_20220524185357
20220524155952
-0.5
0.0
0.5
1.0
1.5
2.0
2.5
3.0
3.5
4.0
4.5
5.0
5.5
6.0
6.5
7.0
7.5
8.0
6.5
6.0
5.5
5.0
4.5
4.0
3.5
3.0
2.5
2.0
1.5
1.0
0.5
0.0

[19]F NMR of **12**

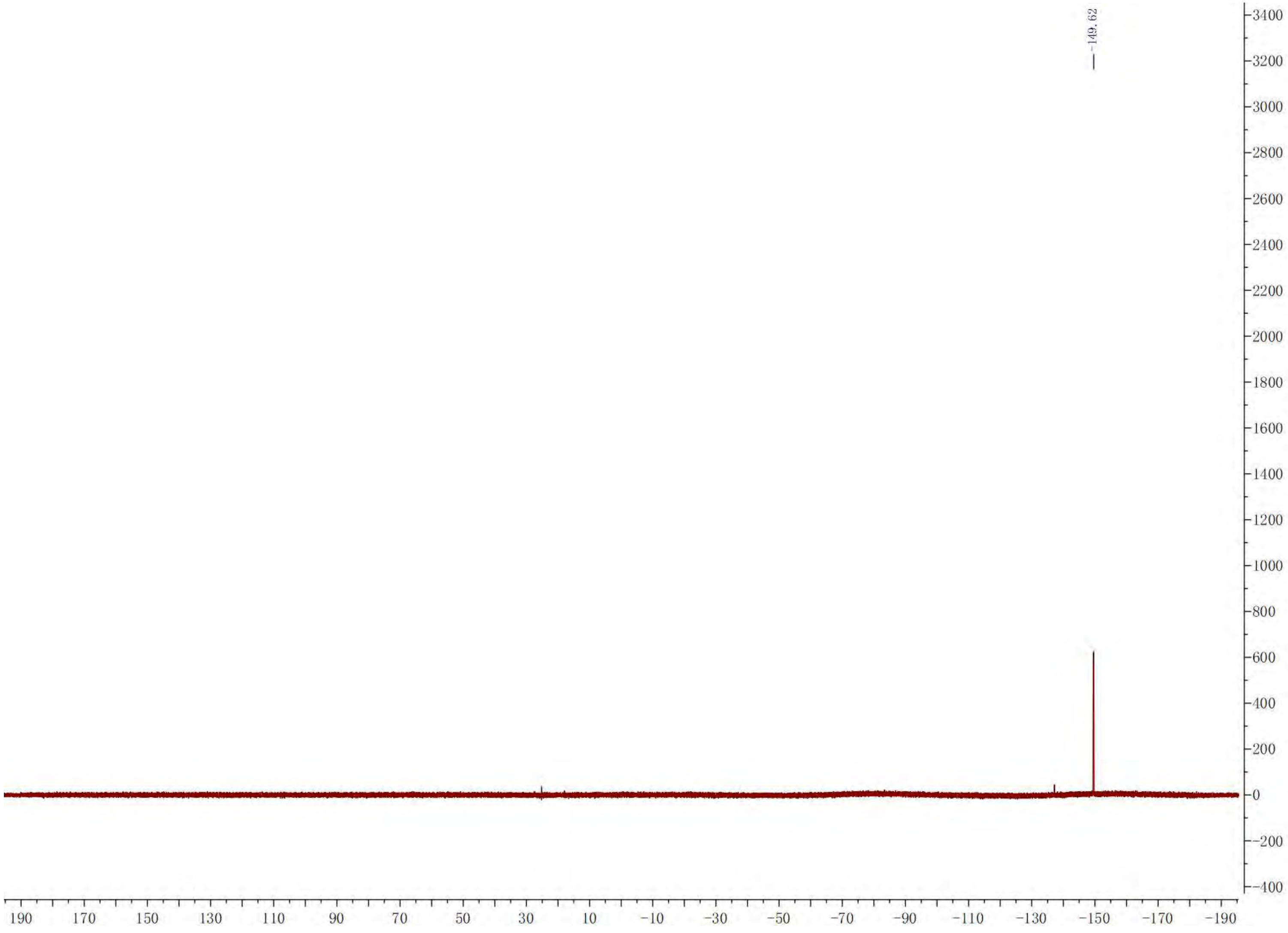

-149.62
3400
3200
3000
2800
2600
2400
2200
2000
1800
1600
1400
1200
1000
800
600
400
200
0
-200
-400
190 170 150 130 110 90 70 50 30 10 -10 -30 -50 -70 -90 -110 -130 -150 -170 -190

HSQC of **12**

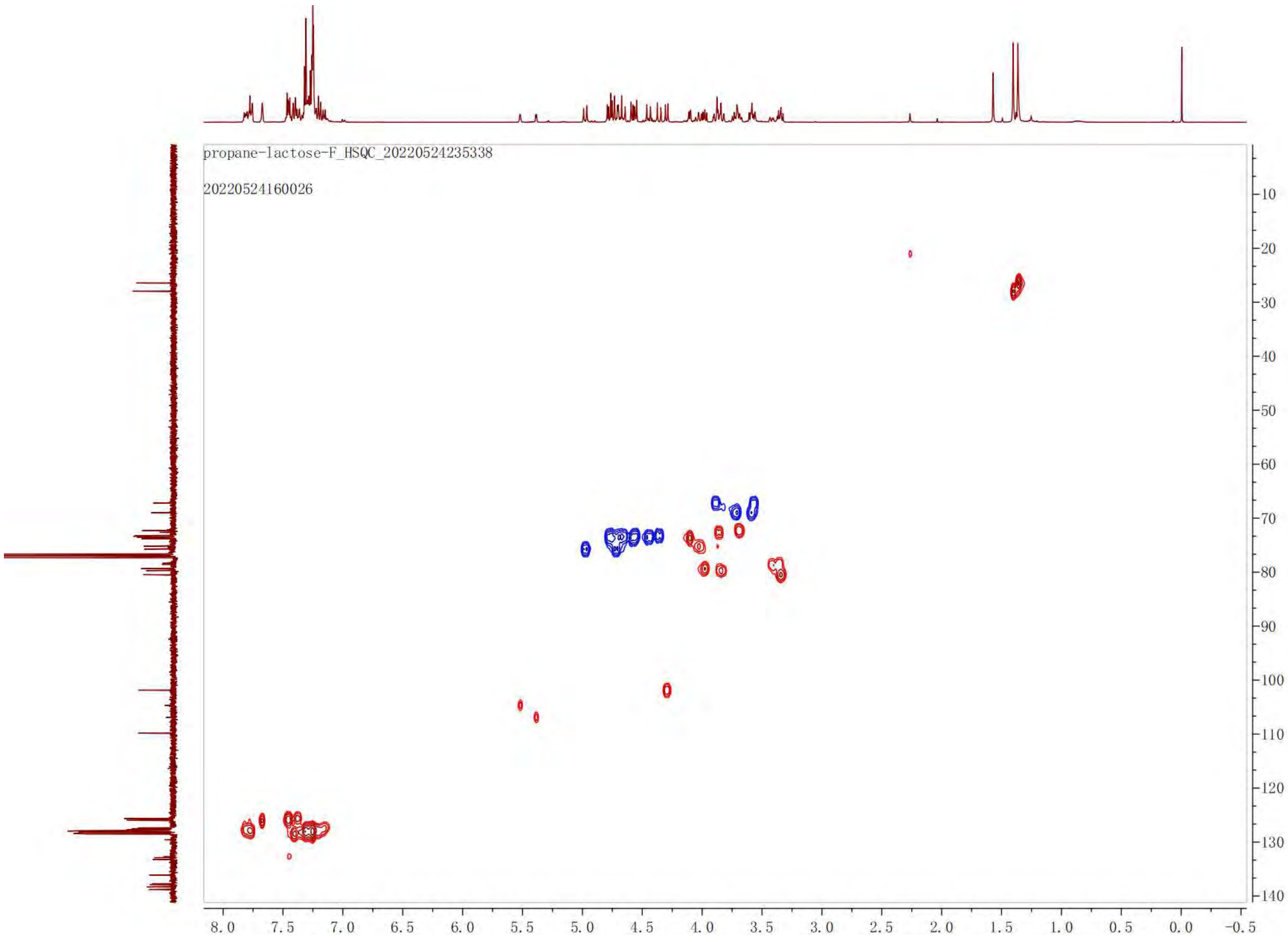
propane-lactose-F_HSQC_20220524235338
20220524160026
10
20
30
40
50
60
70
80
90
100
110
120
130
140
8.0
7.5
7.0
6.5
6.0
5.5
5.0
4.5
4.0
3.5
3.0
2.5
2.0
1.5
1.0
0.5
0.0
-0.5

$^{13}$C NMR of **12**. 100 MHz, $CDCl_3$.

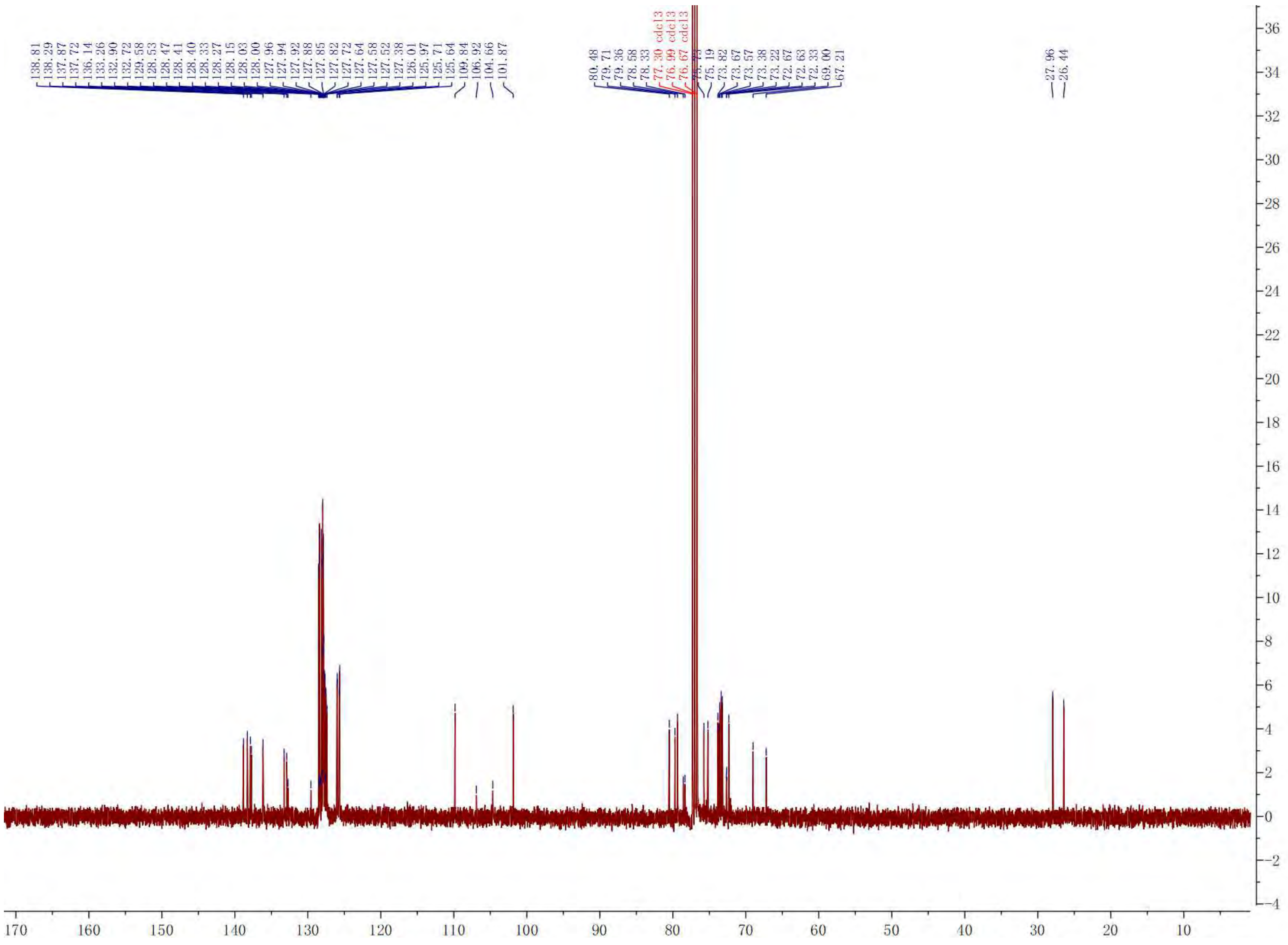

$^{1}$H NMR of **8**. 400 MHz, $CDCl_3$.

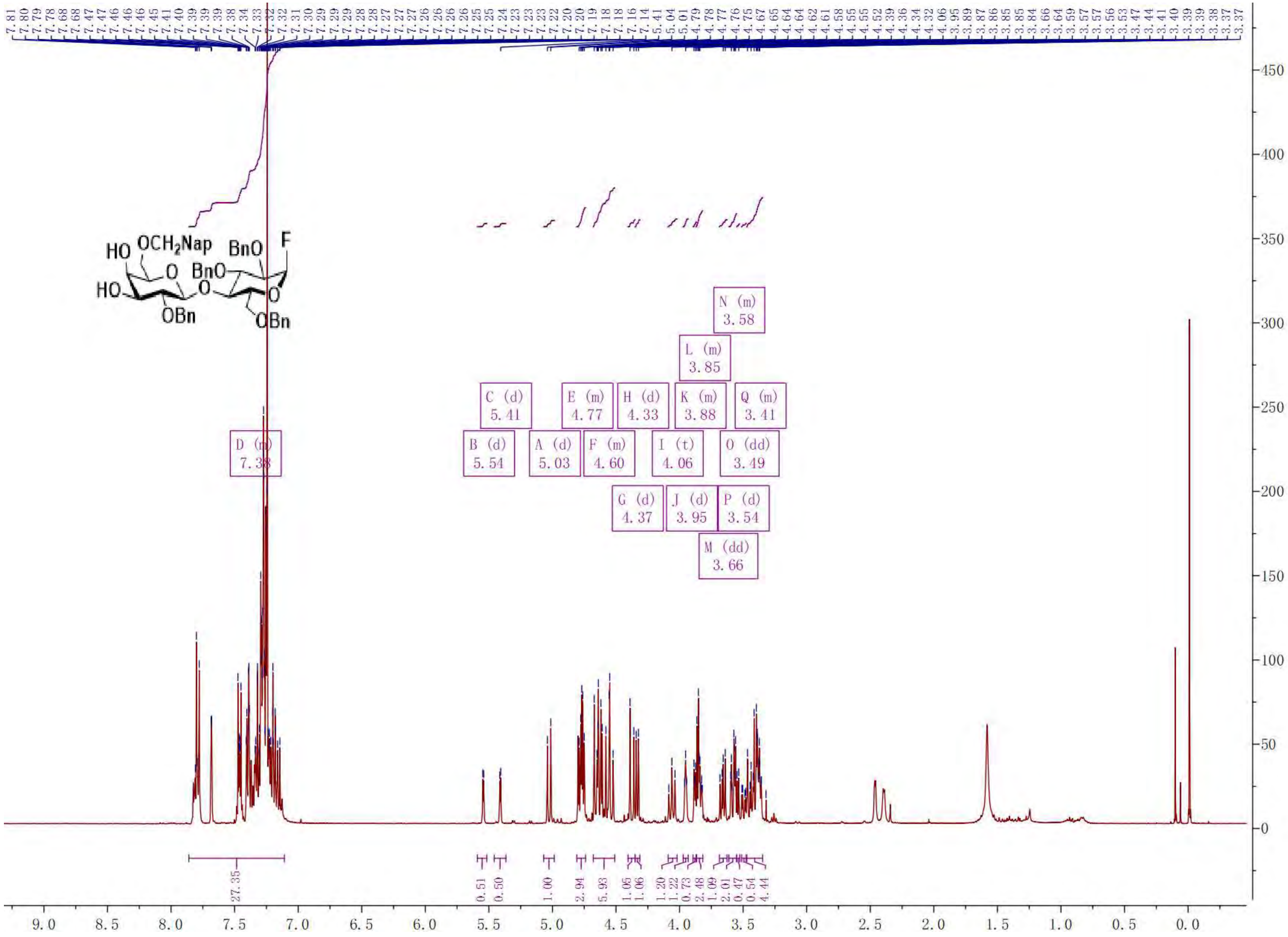

COSY of **8**

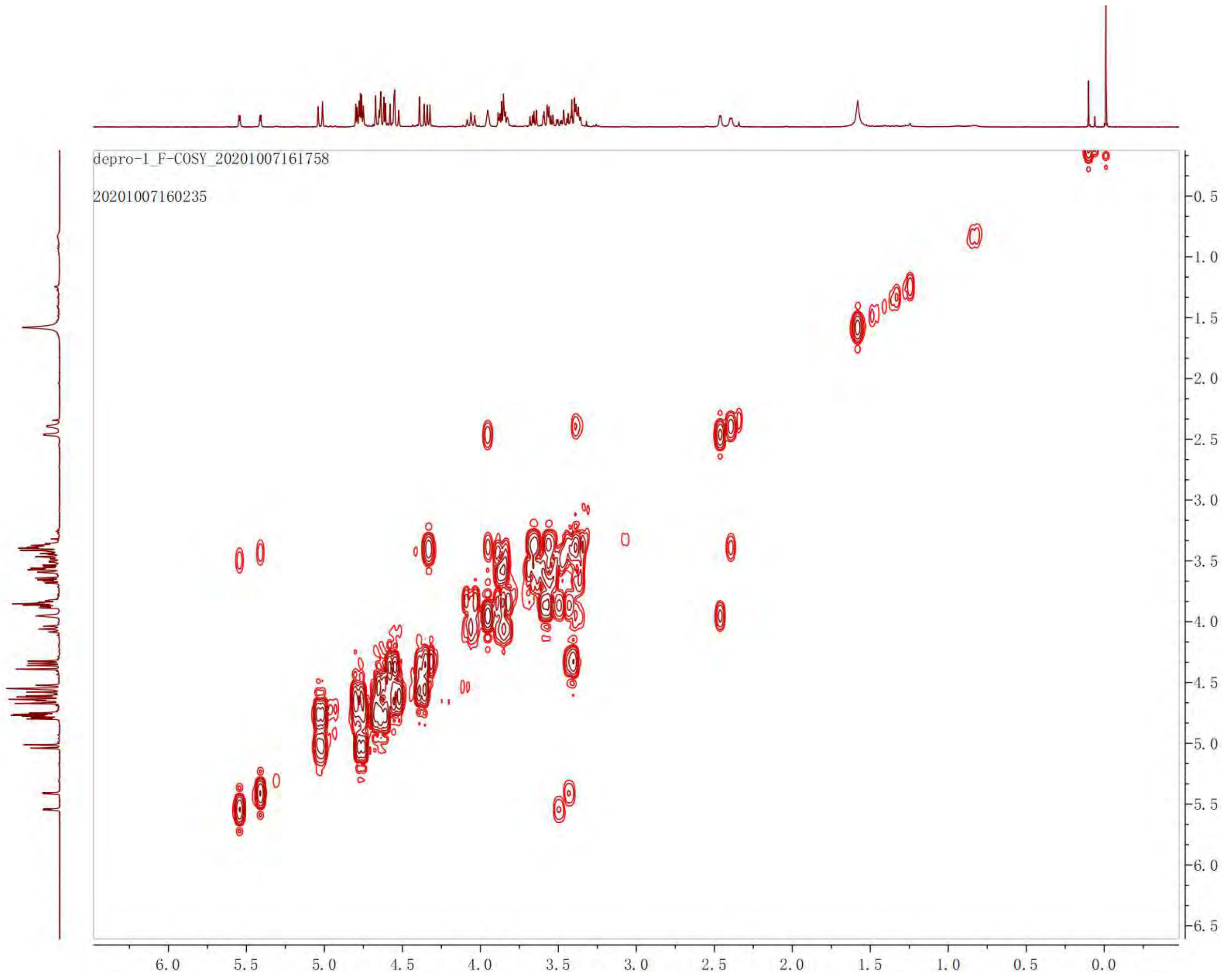

depro-1_F-COSY_20201007161758
20201007160235
6.0
5.5
5.0
4.5
4.0
3.5
3.0
2.5
2.0
1.5
1.0
0.5
0.0
-0.5
-1.0
-1.5
-2.0
-2.5
-3.0
-3.5
-4.0
-4.5
-5.0
-5.5
-6.0
-6.5

$^{19}$F NMR of **8**

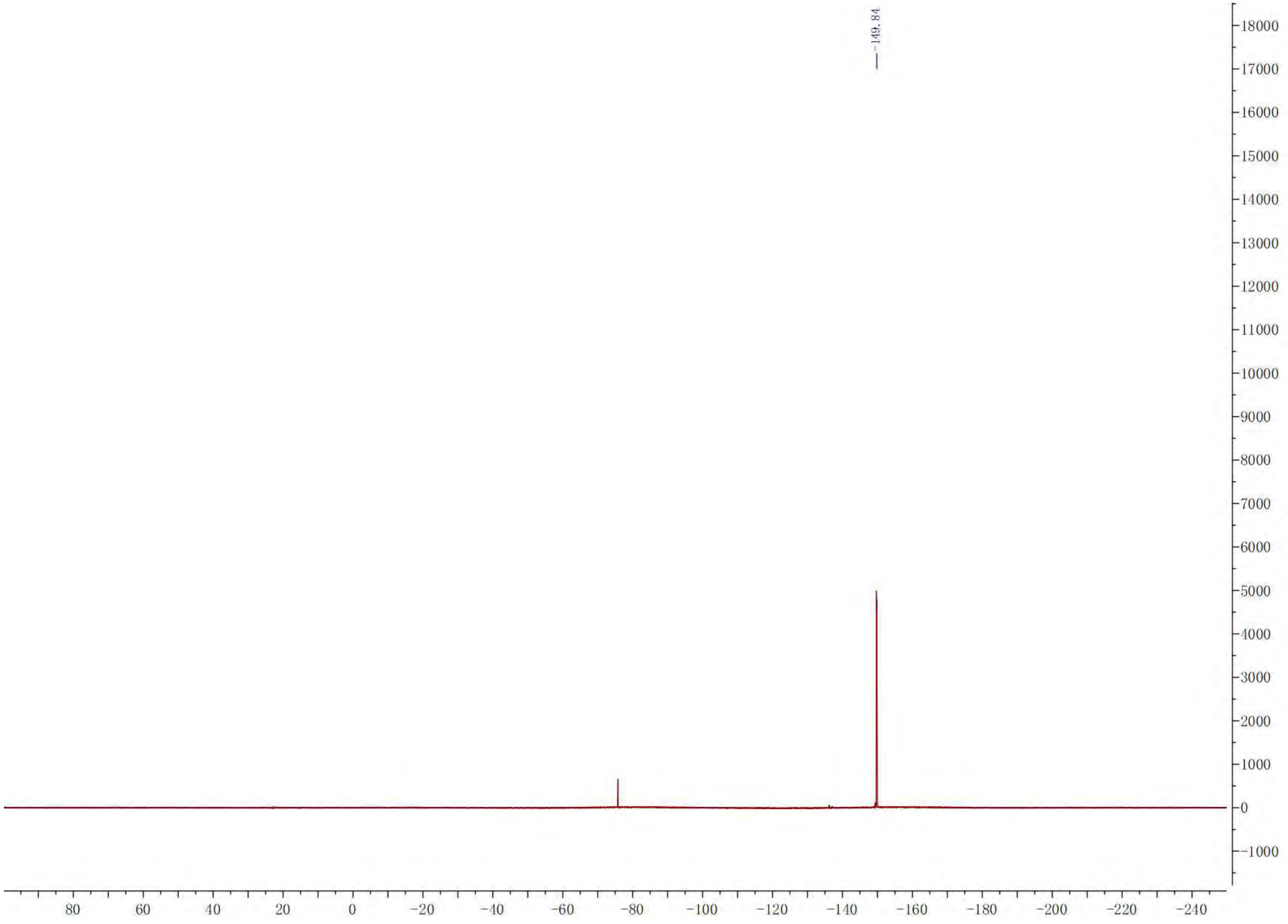
-149.84
18000
17000
16000
15000
14000
13000
12000
11000
10000
9000
8000
7000
6000
5000
4000
3000
2000
1000
0
-1000
80 60 40 20 0 -20 -40 -60 -80 -100 -120 -140 -160 -180 -200 -220 -240

HSQC of **8**

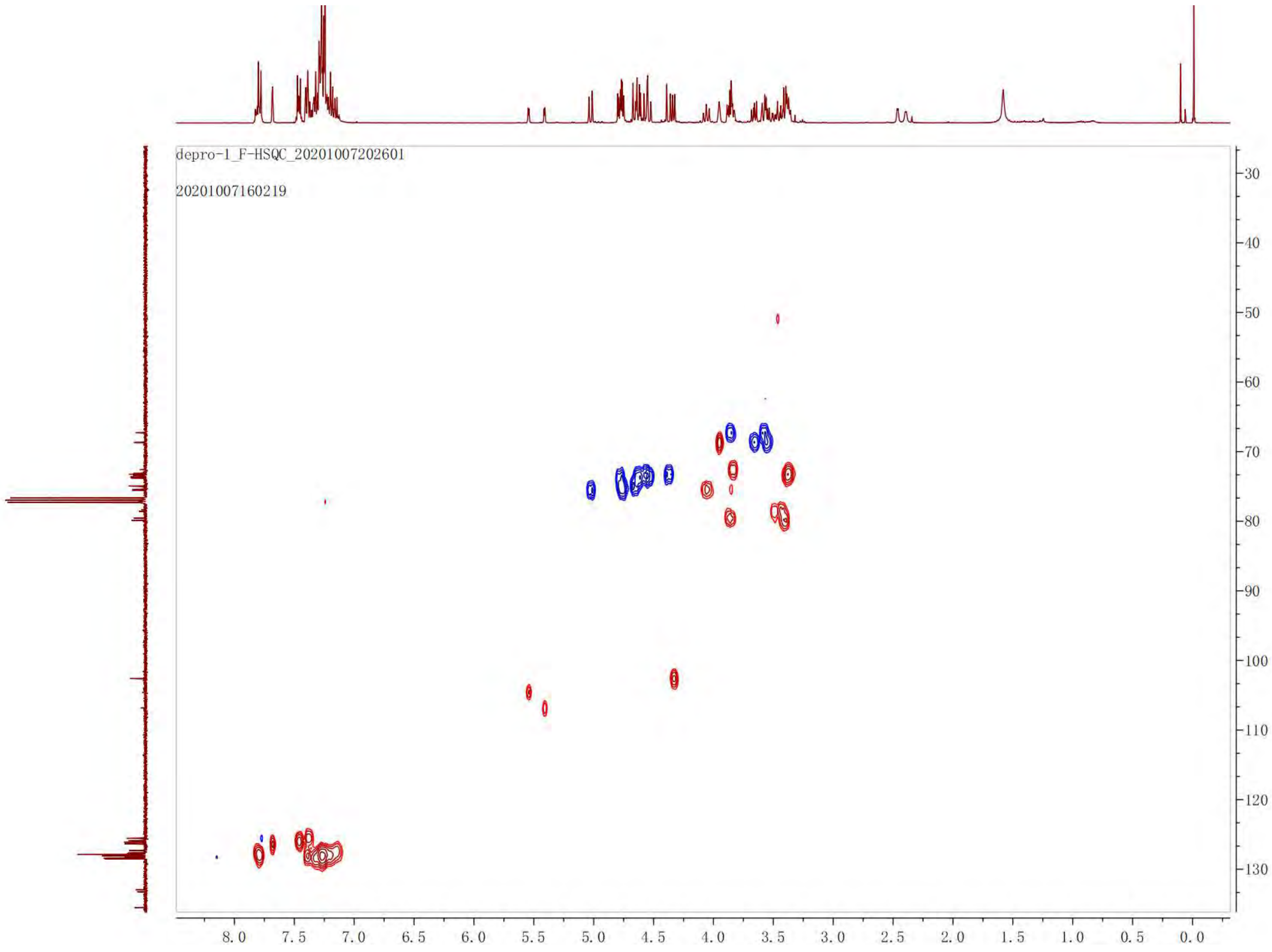
depro-1_F-HSQC_20201007202601
20201007160219
30
40
50
60
70
80
90
100
110
120
130
8.0
7.5
7.0
6.5
6.0
5.5
5.0
4.5
4.0
3.5
3.0
2.5
2.0
1.5
1.0
0.5
0.0

$^{13}$C NMR of **8**. 100 MHz, $CDCl_3$.

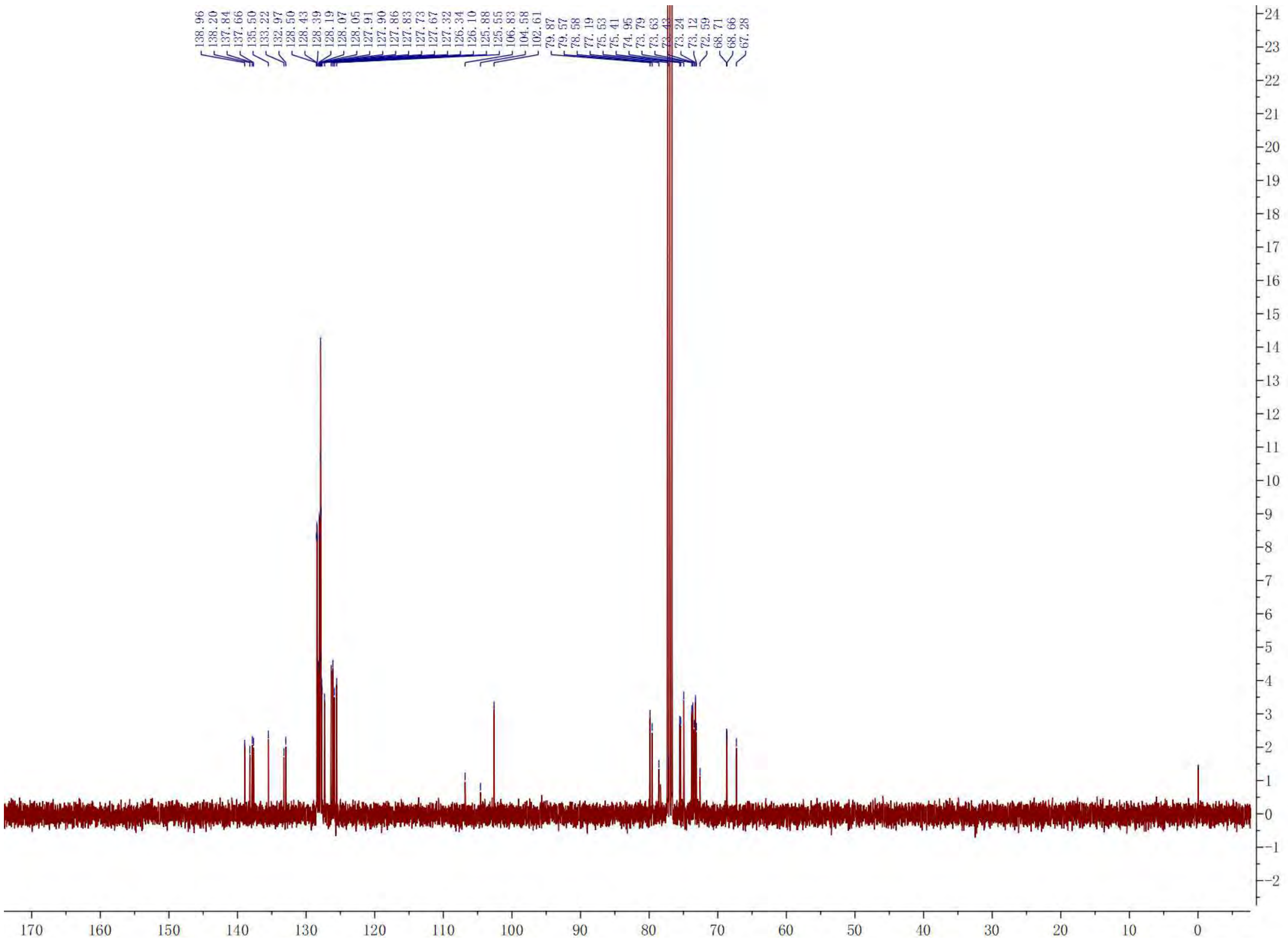

[1]H NMR of **13**. 600 MHz, $CDCl_3$.

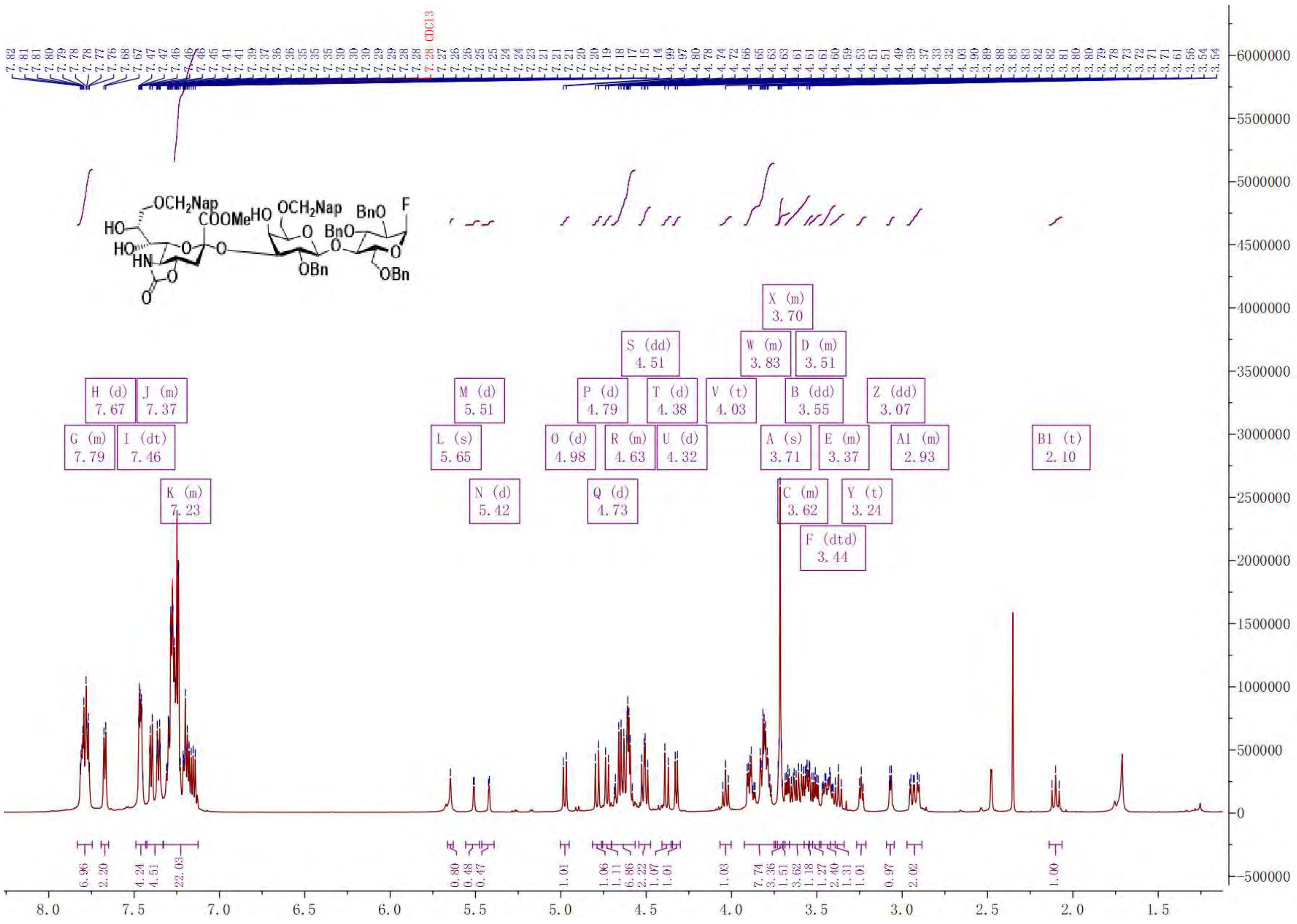

COSY of **13**

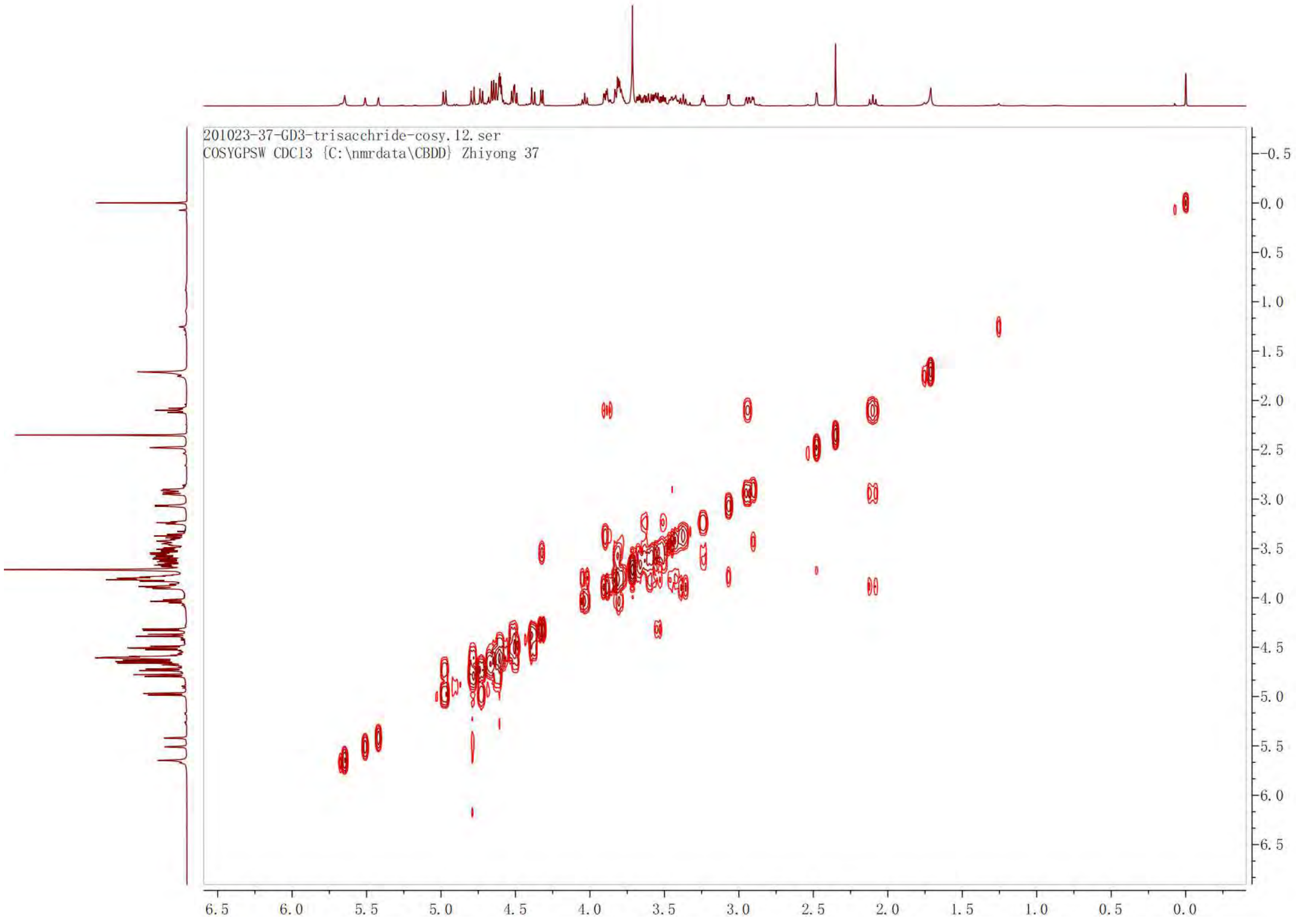

201023-37-GD3-trisacchride-cosy.12.ser
COSYGPSW CDCl3 {C:\nmrdata\CBDD} Zhiyong 37

$^{19}$F NMR of **13**

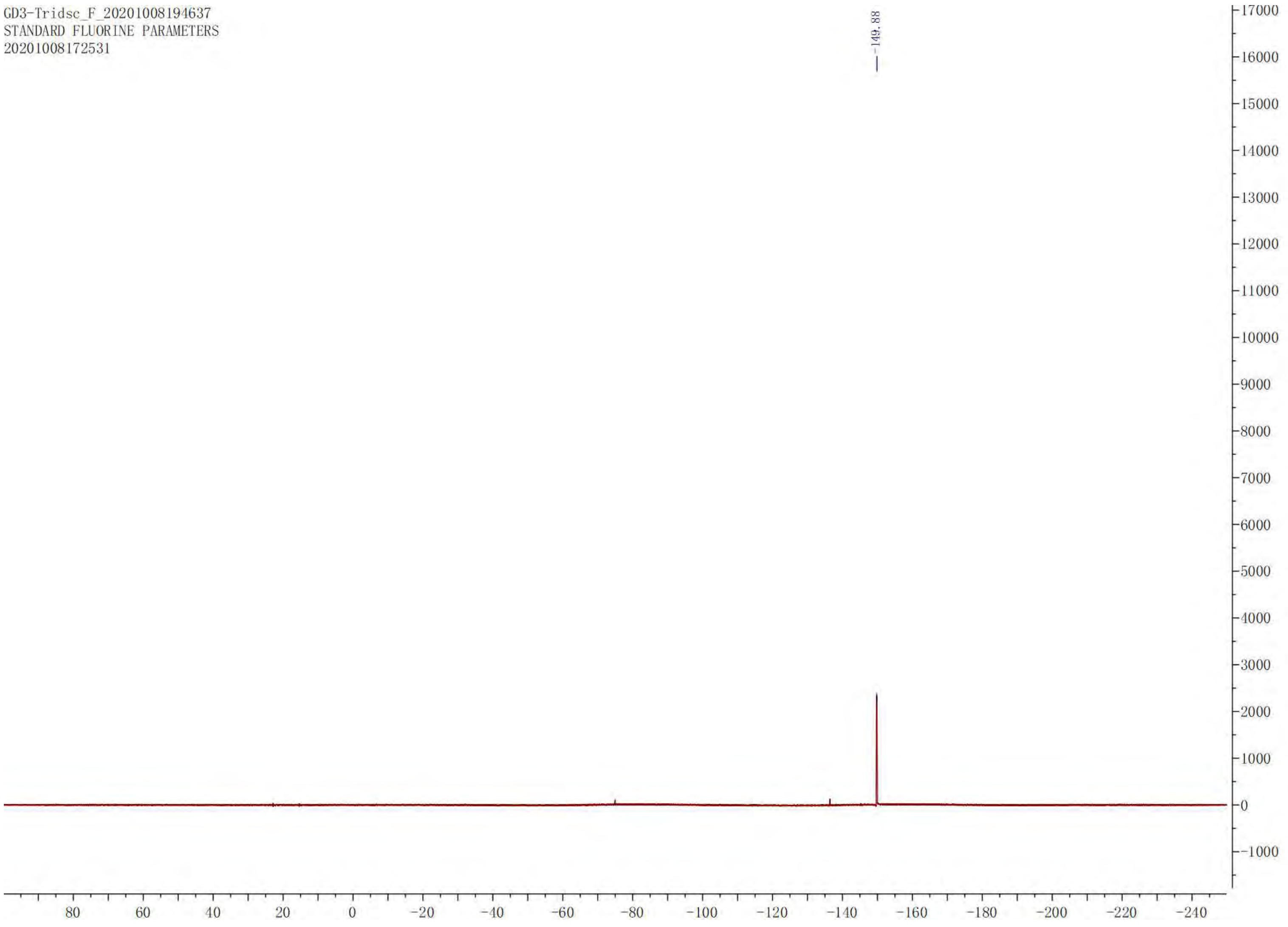

GD3-Tridsc_F_20201008194637
STANDARD FLUORINE PARAMETERS
20201008172531
-149.88
17000
16000
15000
14000
13000
12000
11000
10000
9000
8000
7000
6000
5000
4000
3000
2000
1000
0
-1000
80
60
40
20
0
-20
-40
-60
-80
-100
-120
-140
-160
-180
-200
-220
-240

HSQC of **13**

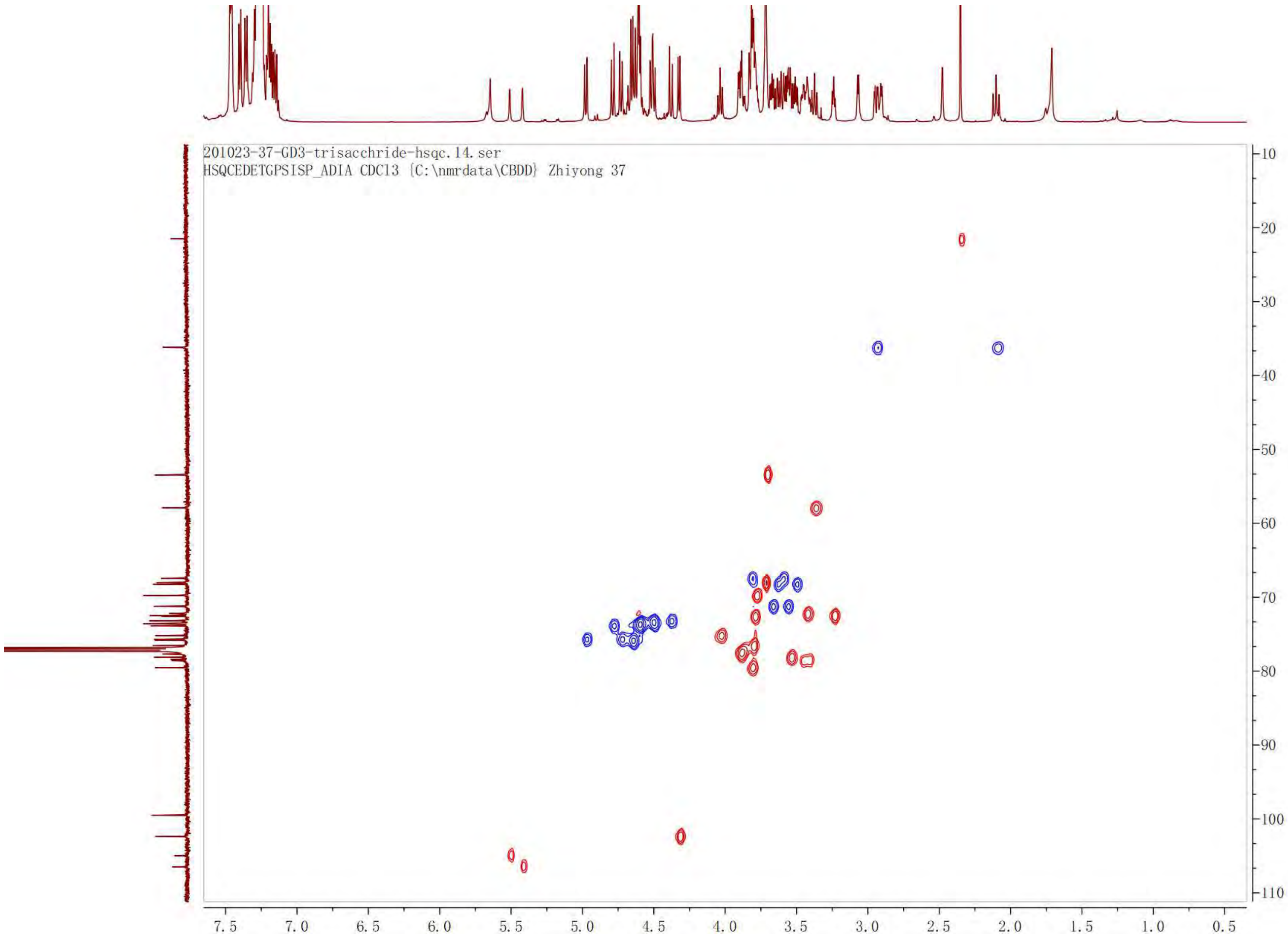

201023-37-GD3-trisacchride-hsqc.14.ser
HSQCEDETGPSISP_ADIA CDCl3 {C:\nmrdata\CBDD} Zhiyong 37
7.5
7.0
6.5
6.0
5.5
5.0
4.5
4.0
3.5
3.0
2.5
2.0
1.5
1.0
0.5
10
20
30
40
50
60
70
80
90
100
110

$^{13}$C NMR of **13**. 125 MHz, $CDCl_3$.

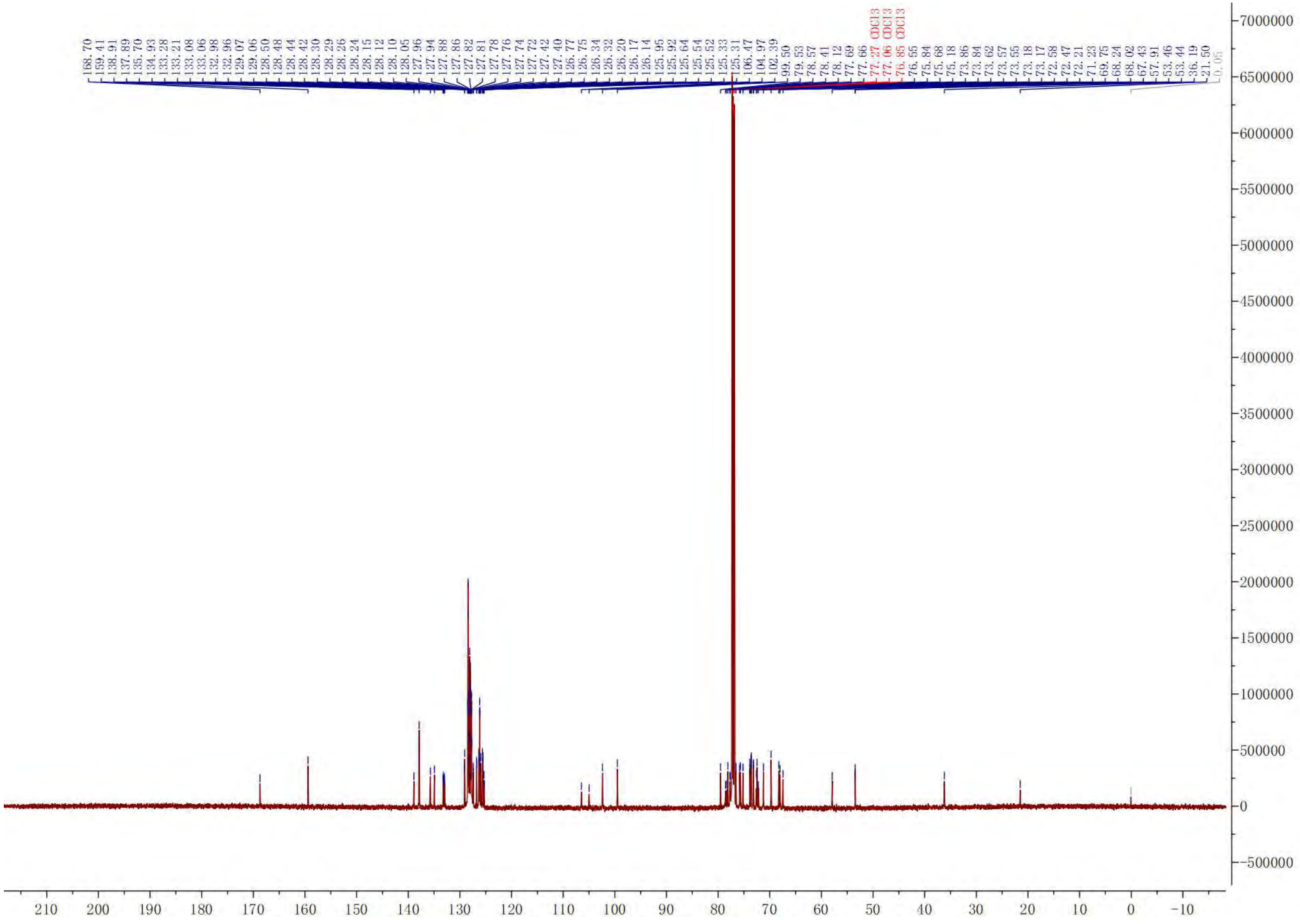

[1]H NMR of **14**. 600 MHz, $CDCl_3$.

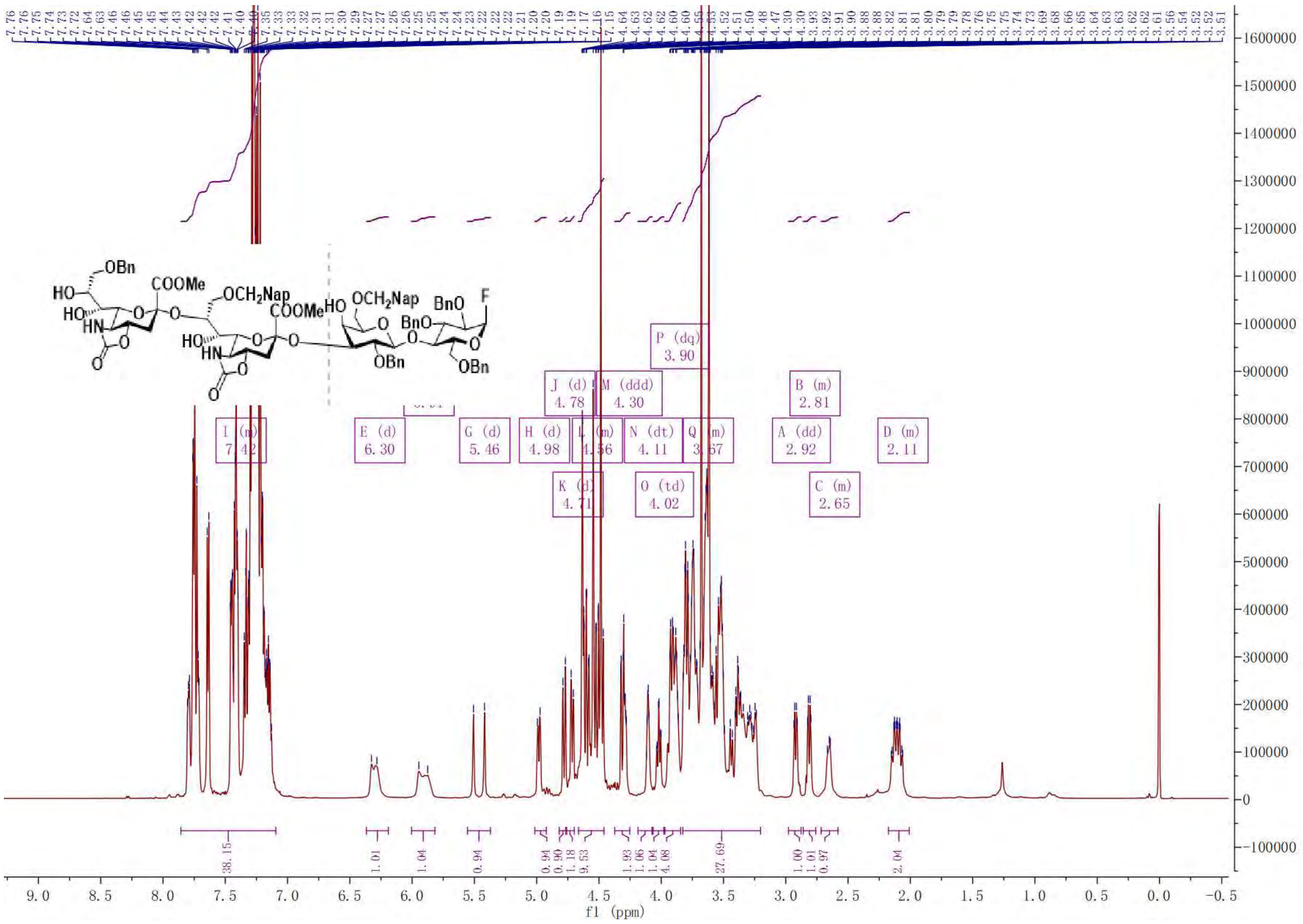

COSY of **14**

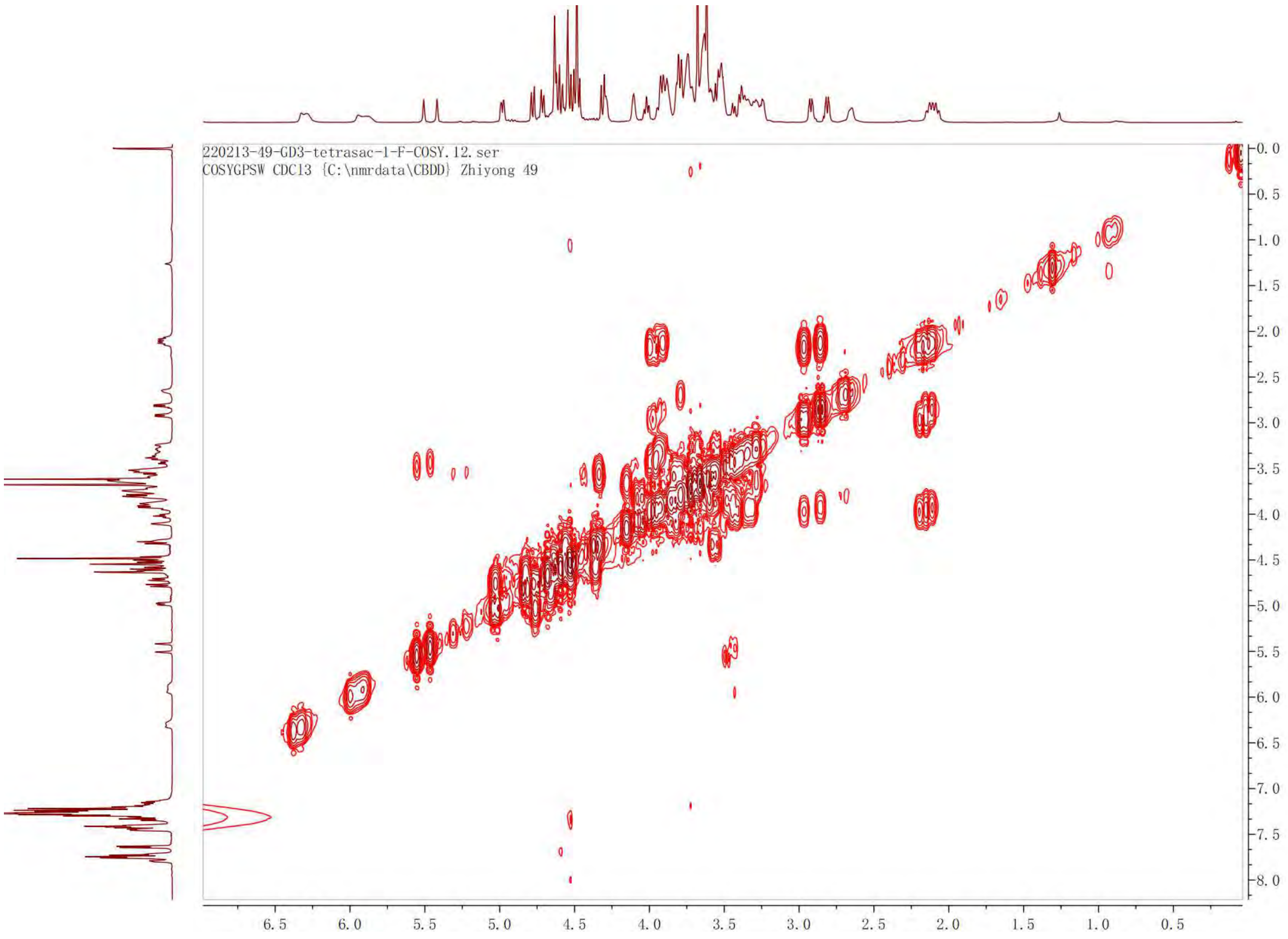
220213-49-GD3-tetrasac-1-F-COSY.12.ser
COSYGPSW CDCl3 {C:\nmrdata\CBDD} Zhiyong 49
6.5
6.0
5.5
5.0
4.5
4.0
3.5
3.0
2.5
2.0
1.5
1.0
0.5
0.0
0.5
1.0
1.5
2.0
2.5
3.0
3.5
4.0
4.5
5.0
5.5
6.0
6.5
7.0
7.5
8.0

TOCSY of **14**

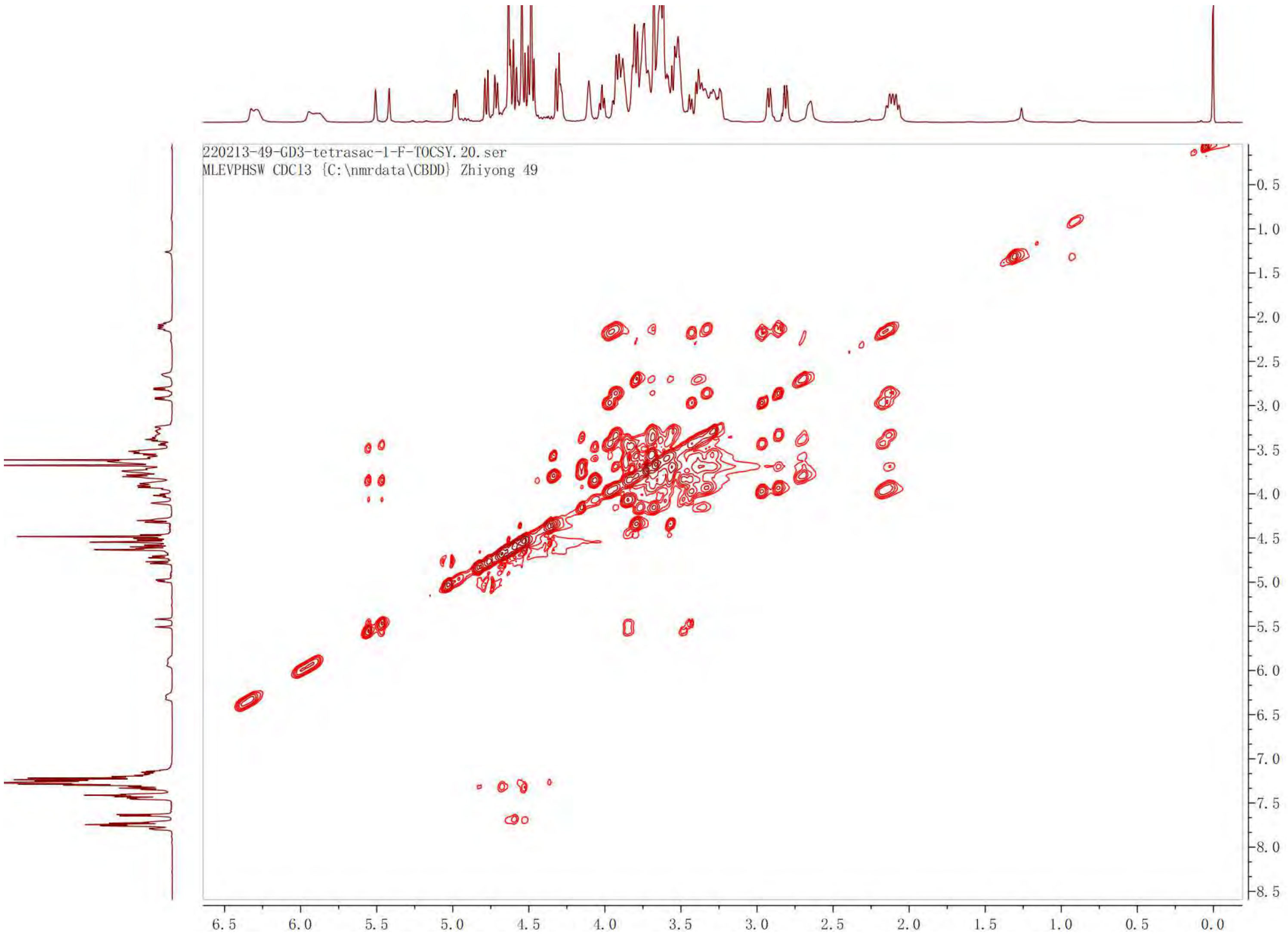

220213-49-GD3-tetrasac-1-F-TOCSY.20.ser
MLEVPHSW CDCl3 {C:\nmrdata\CBDD} Zhiyong 49
0.5
1.0
1.5
2.0
2.5
3.0
3.5
4.0
4.5
5.0
5.5
6.0
6.5
7.0
7.5
8.0
8.5
6.5
6.0
5.5
5.0
4.5
4.0
3.5
3.0
2.5
2.0
1.5
1.0
0.5
0.0

[19]F NMR of **14**

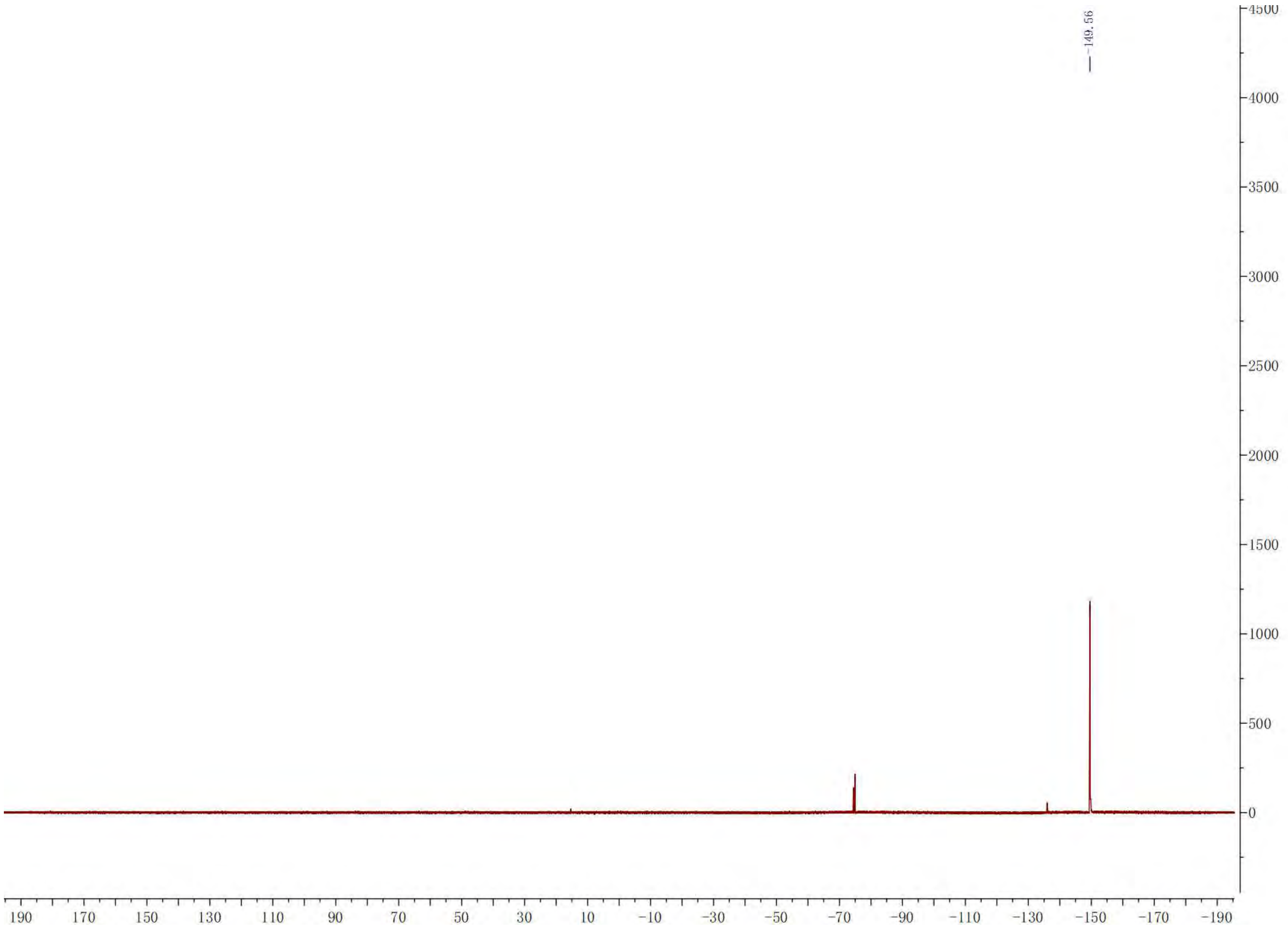

-149.56
4500
4000
3500
3000
2500
2000
1500
1000
500
0
190
170
150
130
110
90
70
50
30
10
-10
-30
-50
-70
-90
-110
-130
-150
-170
-190

HSQC of **14**

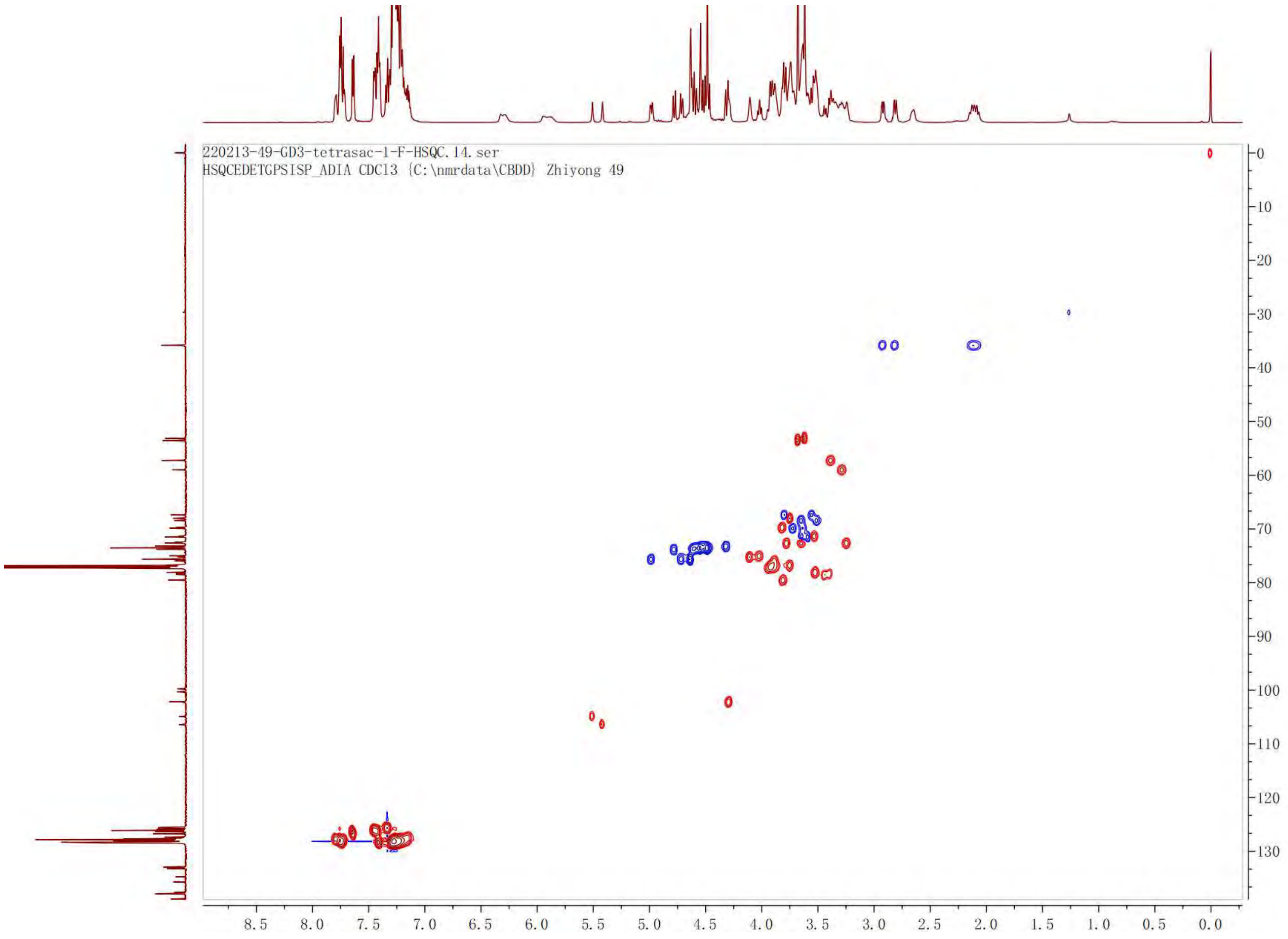

220213-49-GD3-tetrasac-1-F-HSQC.14.ser
HSQCEDETGPSISP_ADIA CDCl3 {C:\nmrdata\CBDD} Zhiyong 49
8.5
8.0
7.5
7.0
6.5
6.0
5.5
5.0
4.5
4.0
3.5
3.0
2.5
2.0
1.5
1.0
0.5
0.0
0
10
20
30
40
50
60
70
80
90
100
110
120
130

[13]C NMR of **14**. 125 MHz, $CDCl_3$.

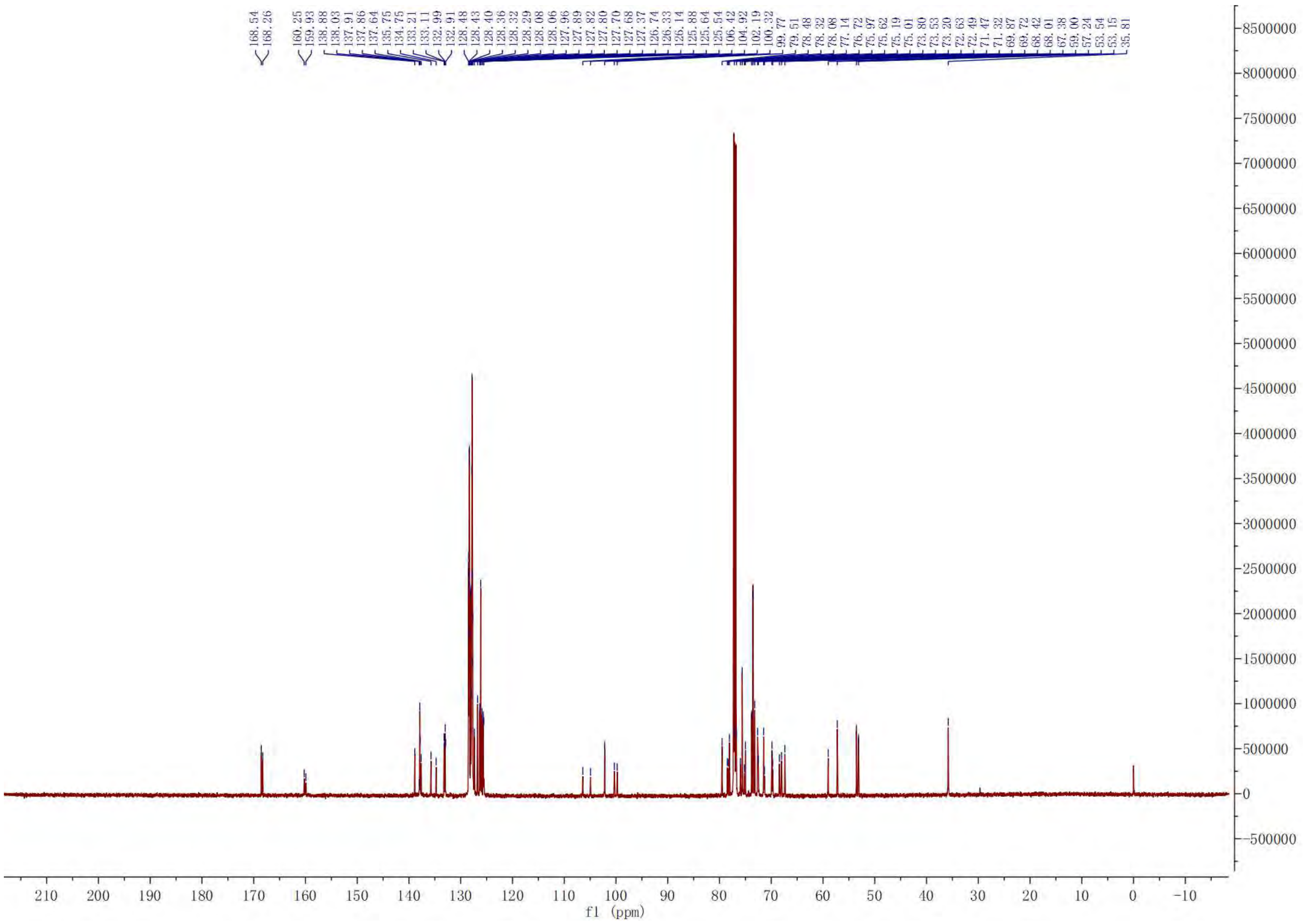

[1]H NMR of **7’**. 600 MHz, $CDCl_3$.

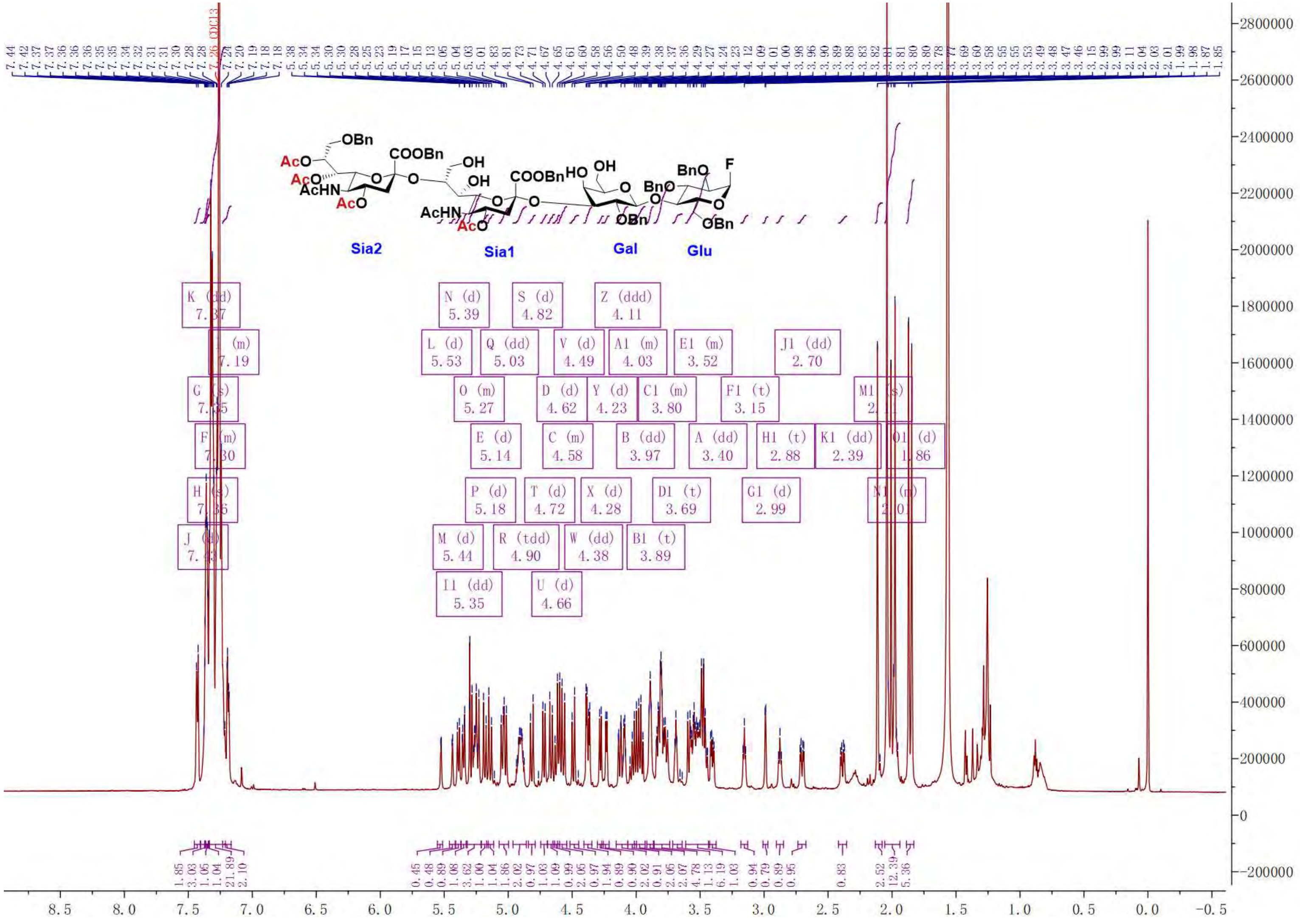

COSY of **7'**

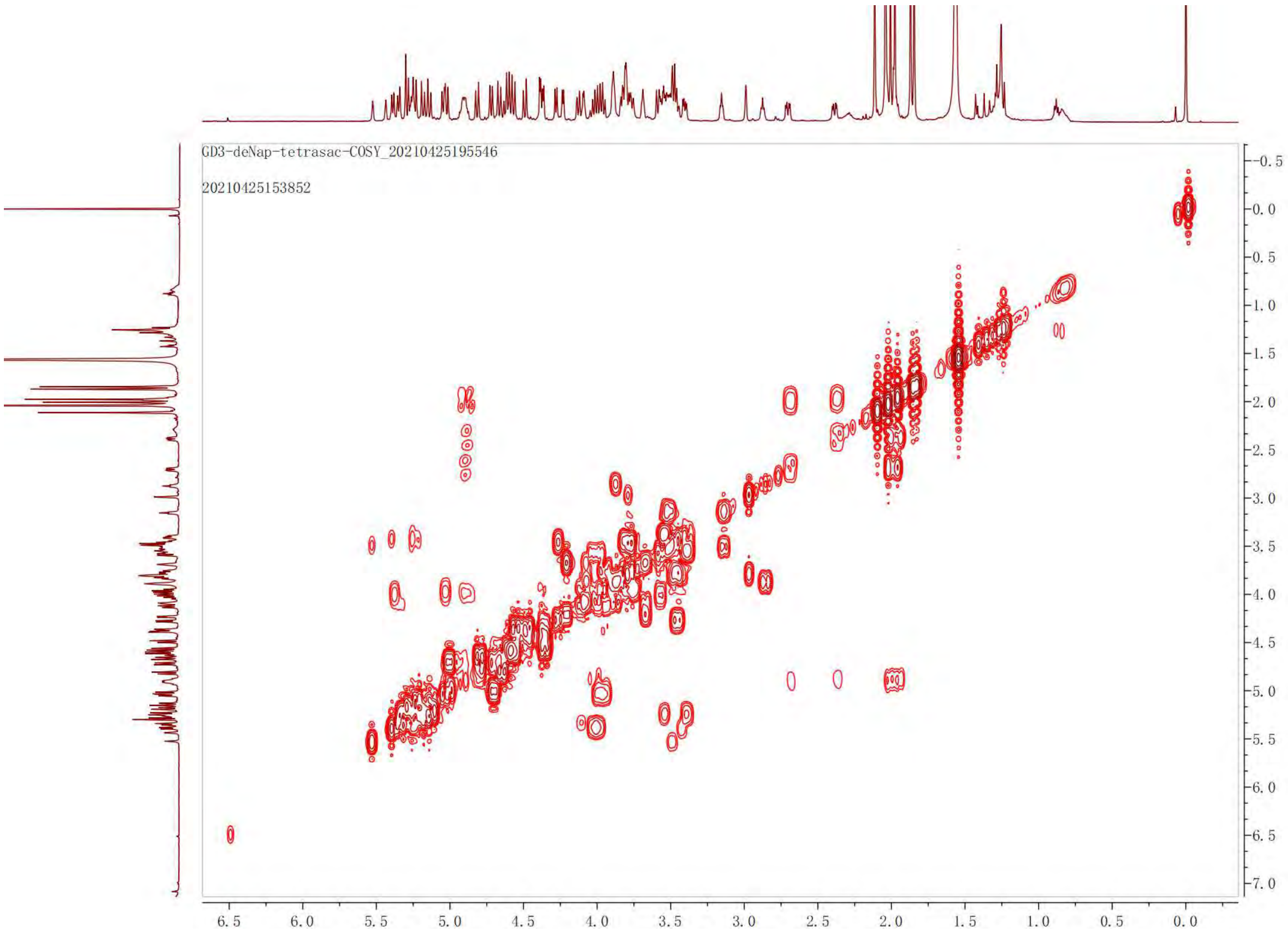

GD3-deNap-tetrasac-COSY_20210425195546
20210425153852
6.5
6.0
5.5
5.0
4.5
4.0
3.5
3.0
2.5
2.0
1.5
1.0
0.5
0.0
-0.5
0.0
0.5
1.0
1.5
2.0
2.5
3.0
3.5
4.0
4.5
5.0
5.5
6.0
6.5
7.0

HSQC of **7’**

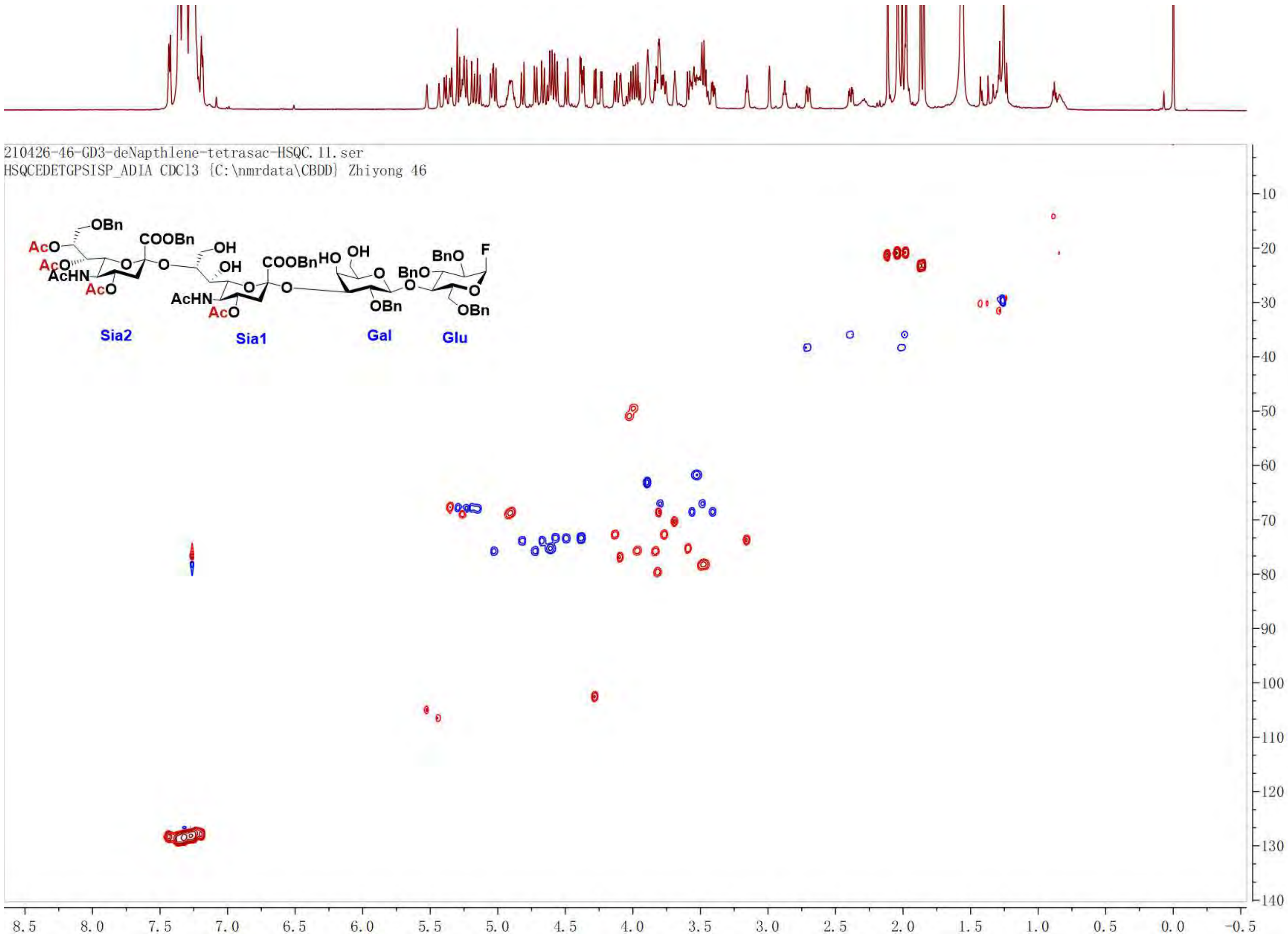

210426-46-GD3-deNapthlene-tetrasac-HSQC.11.ser
HSQCEDETGPSISP_ADIA CDCl3 {C:\nmrdata\CBDD} Zhiyong 46
OBn
COOBn
AcO
AcO
AcHN
AcO
OH
OH
COOBn
HO
OH
BnO
F
AcHN
AcO
OBn
OBn
Sia2
Sia1
Gal
Glu

[1]H NMR of **6**. 600 MHz, $D_2O$.

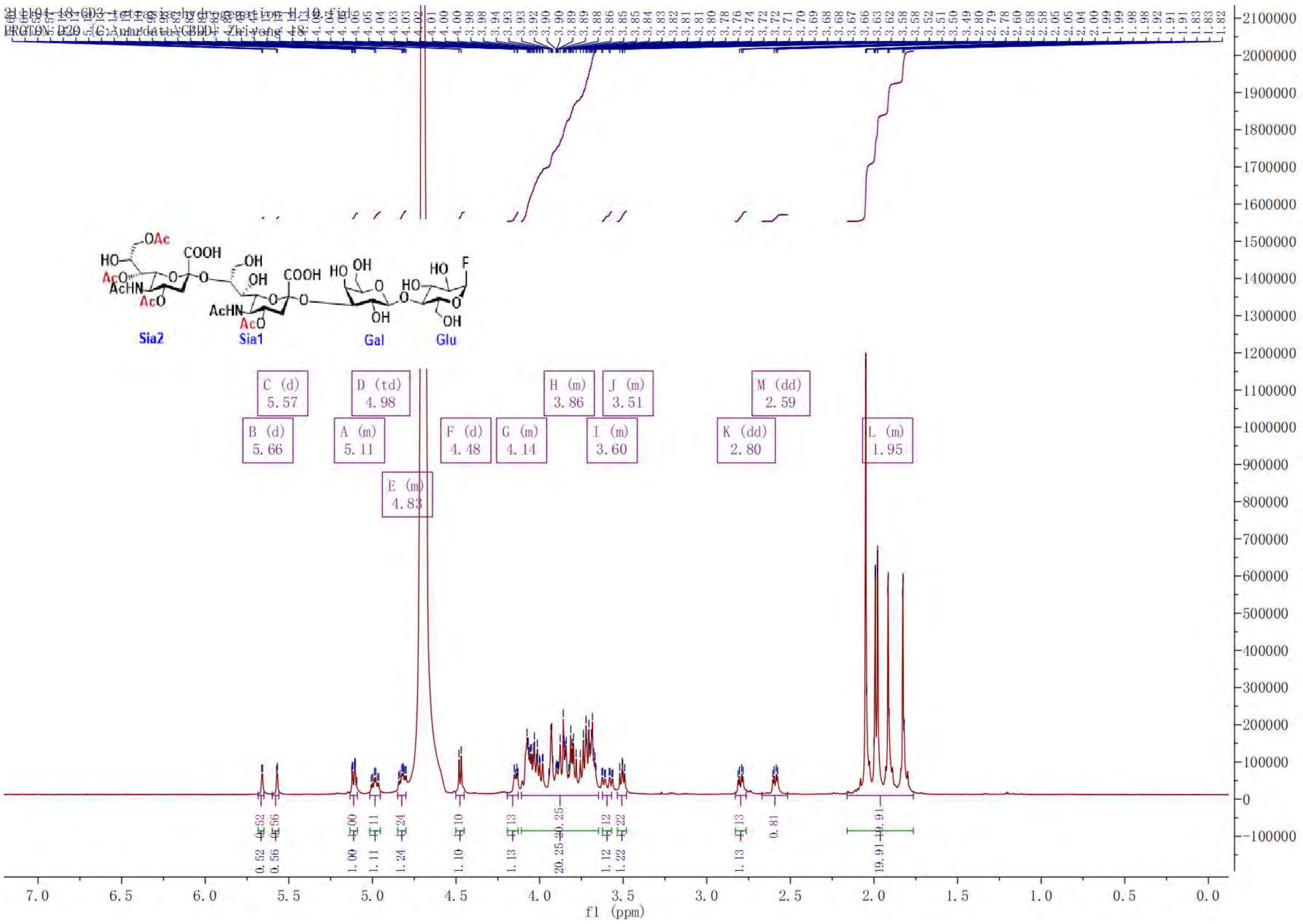

COSY of **6**

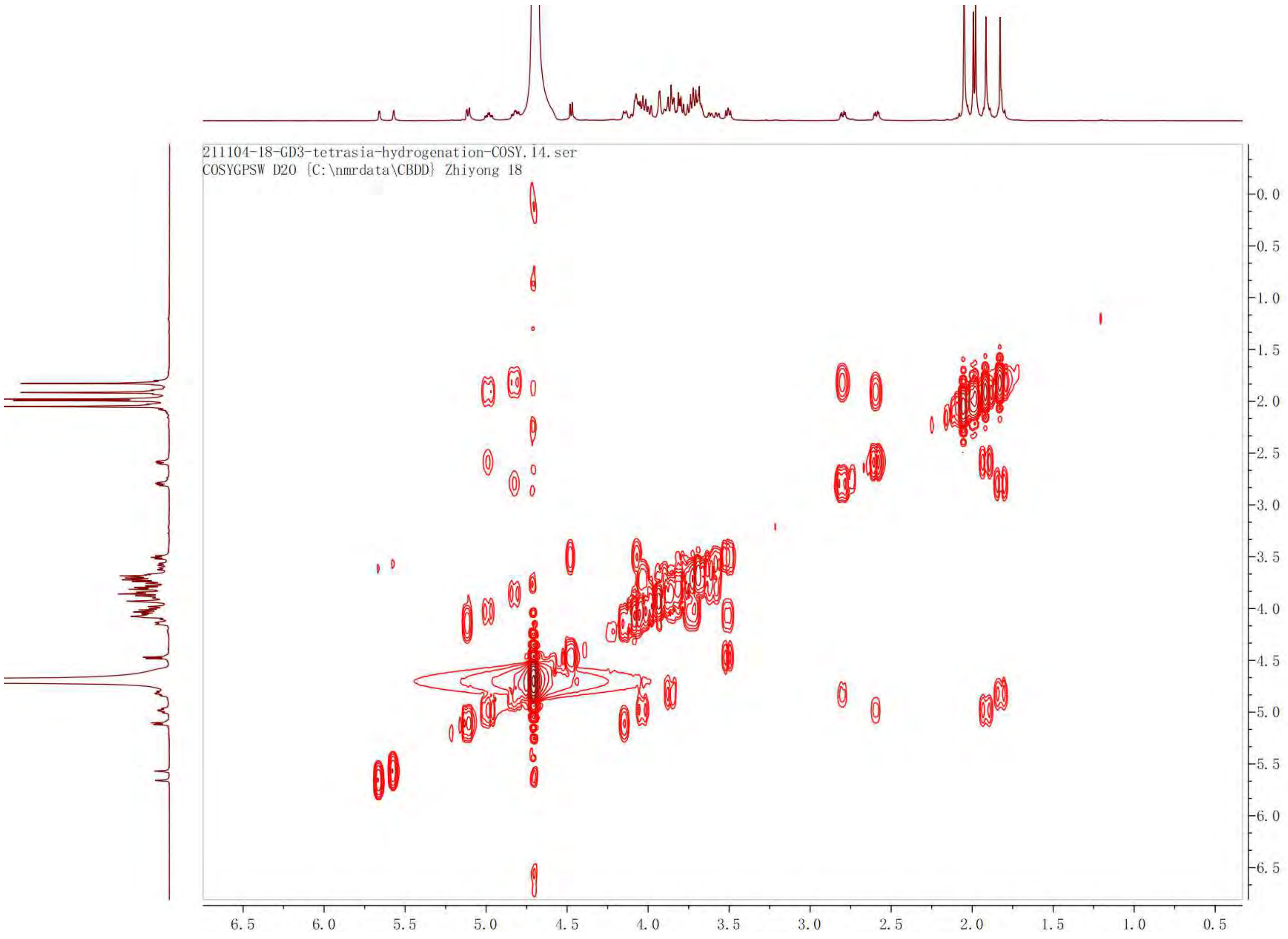

211104-18-GD3-tetrasia-hydrogenation-COSY.14.ser
COSYGPSW D2O {C:\nmrdata\CBDD} Zhiyong 18

HSQC of **6**

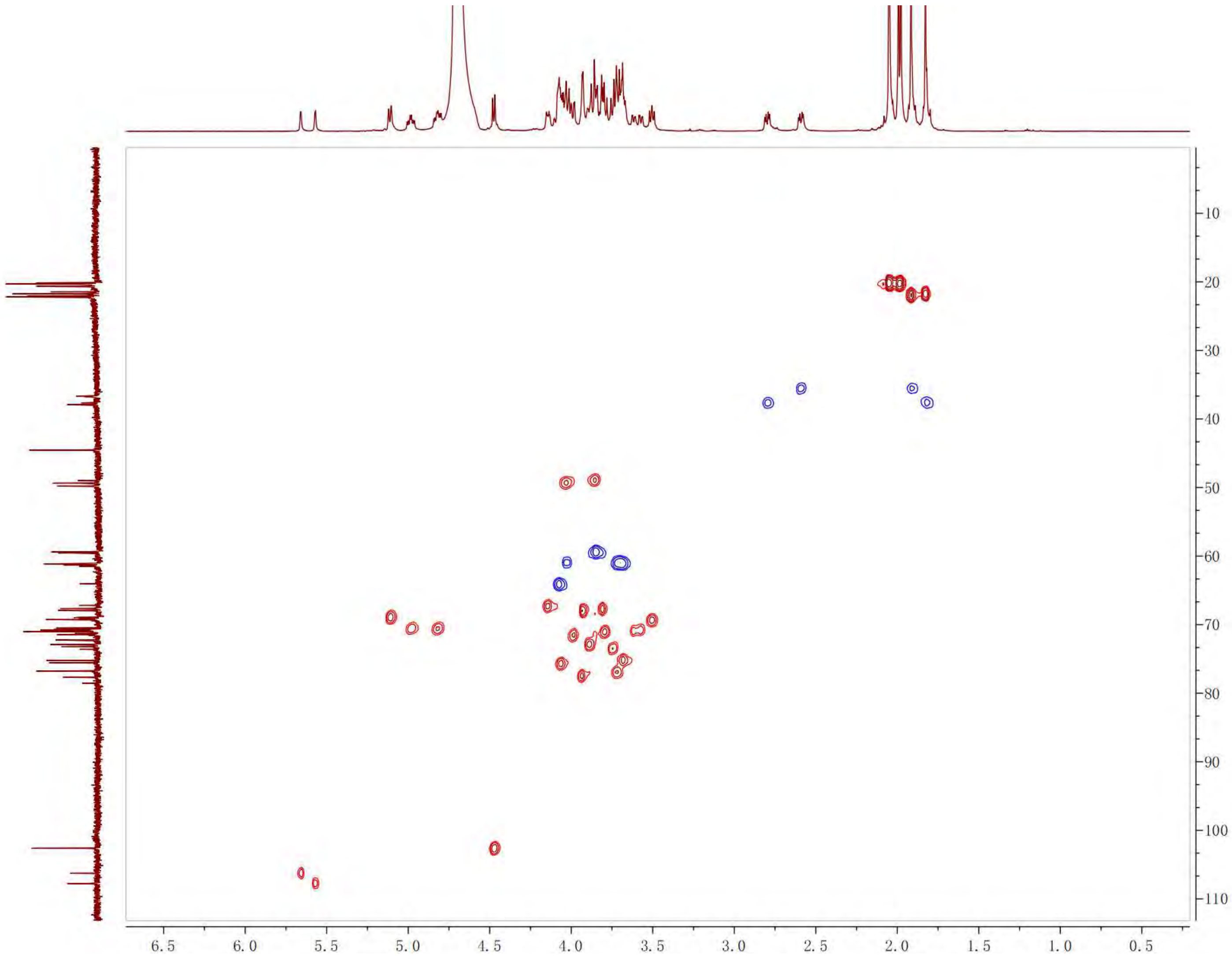
6.5 6.0 5.5 5.0 4.5 4.0 3.5 3.0 2.5 2.0 1.5 1.0 0.5
10 20 30 40 50 60 70 80 90 100 110

HMBC of **6**

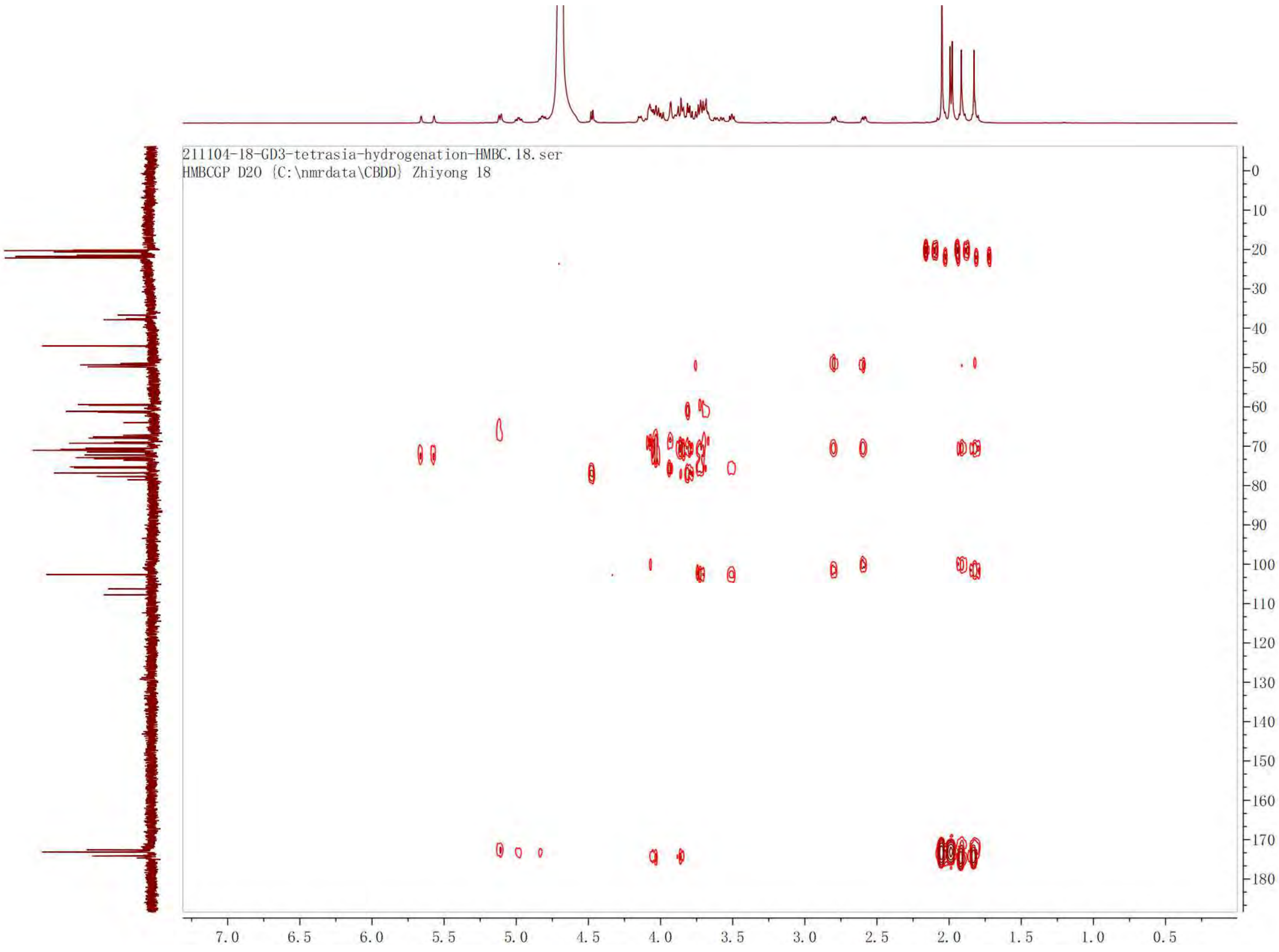
211104-18-GD3-tetrasia-hydrogenation-HMBC.18.ser
HMBCGP D2O {C:\nmrdata\CBDD} Zhiyong 18
0
10
20
30
40
50
60
70
80
90
100
110
120
130
140
150
160
170
180
7.0
6.5
6.0
5.5
5.0
4.5
4.0
3.5
3.0
2.5
2.0
1.5
1.0
0.5

$^{13}C$ NMR of **6**. 125 MHz, $D_2O$.

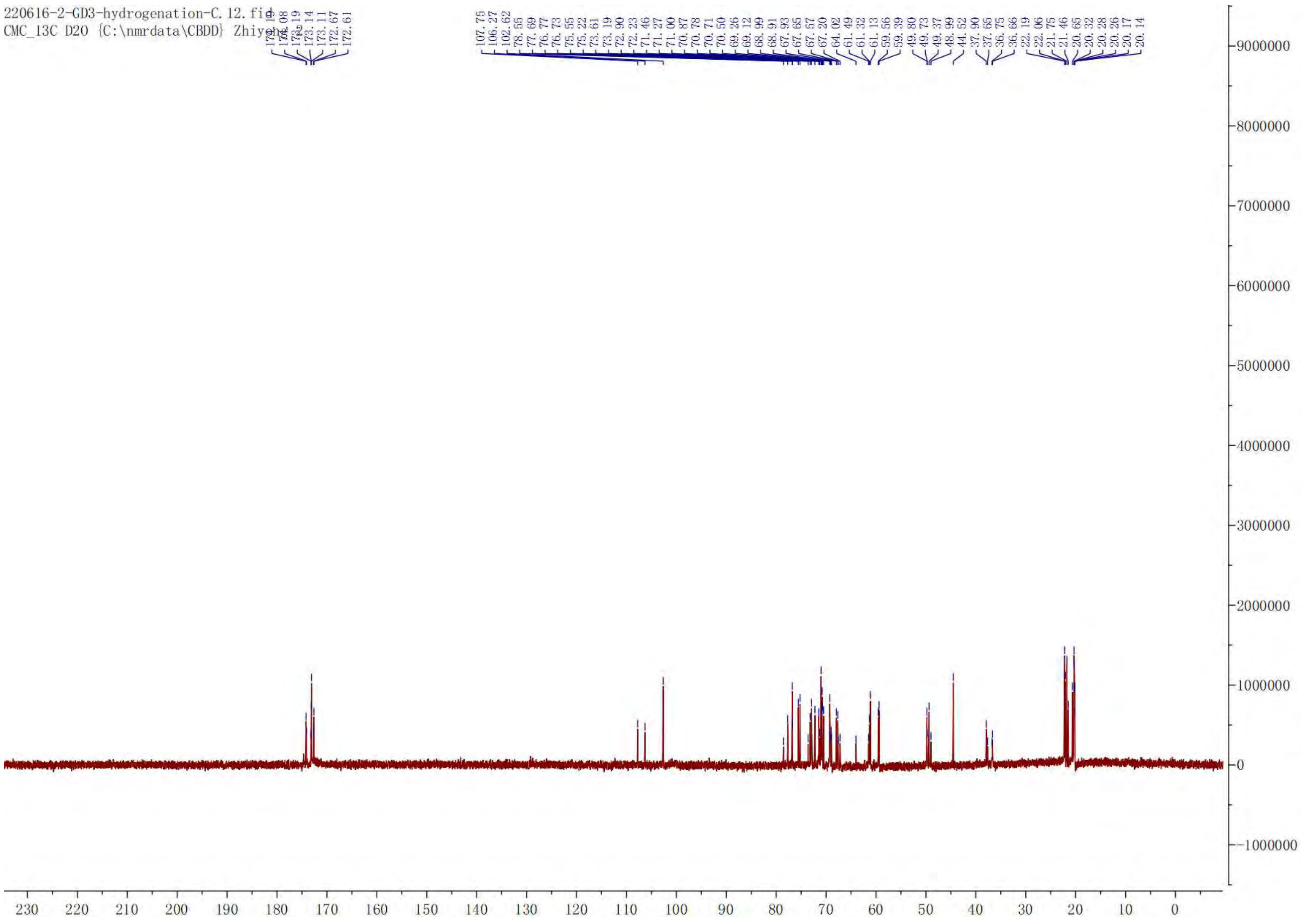

[1]H NMR of **4**. 600 MHz, $D_2O$.

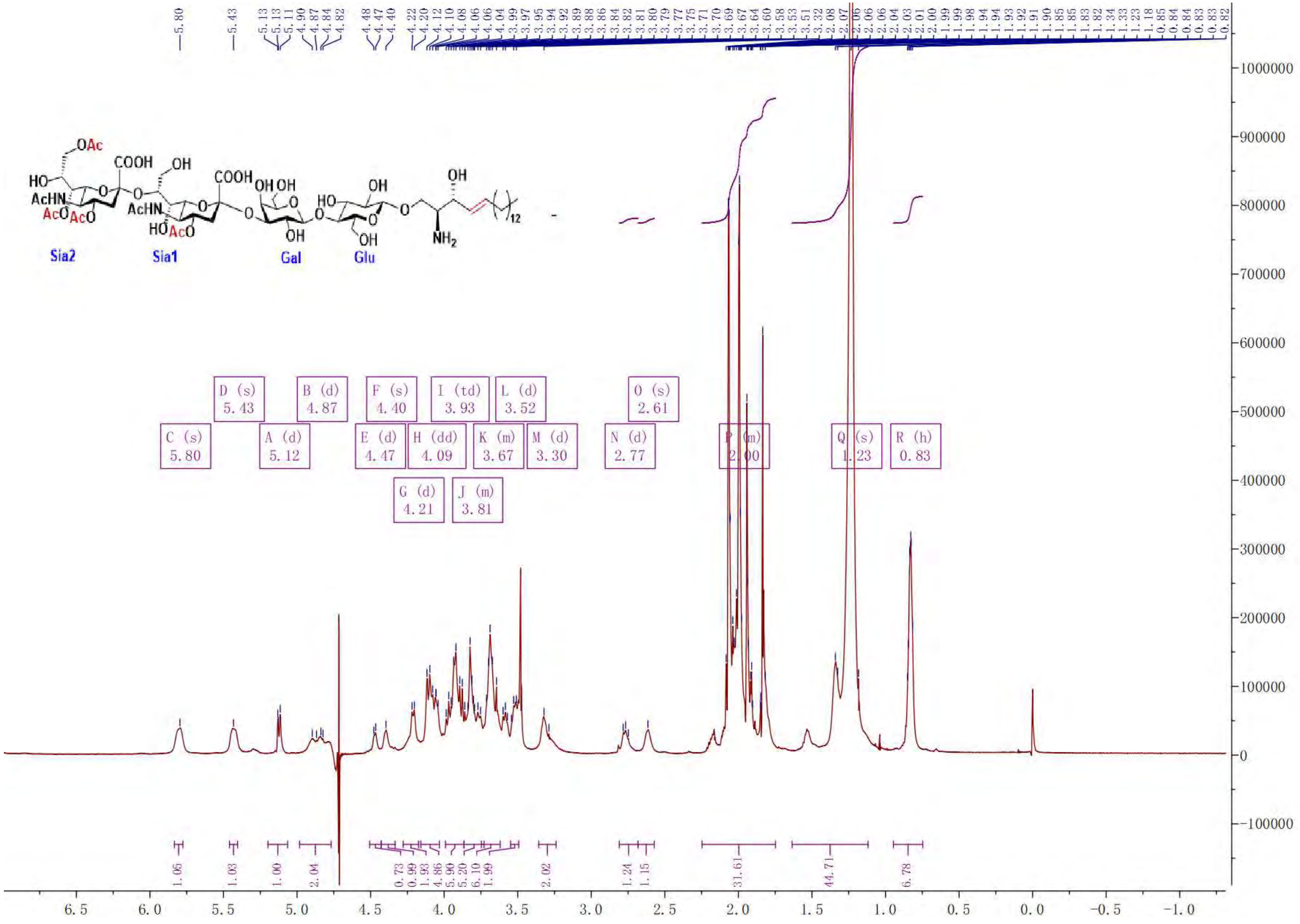

COSY of **4**

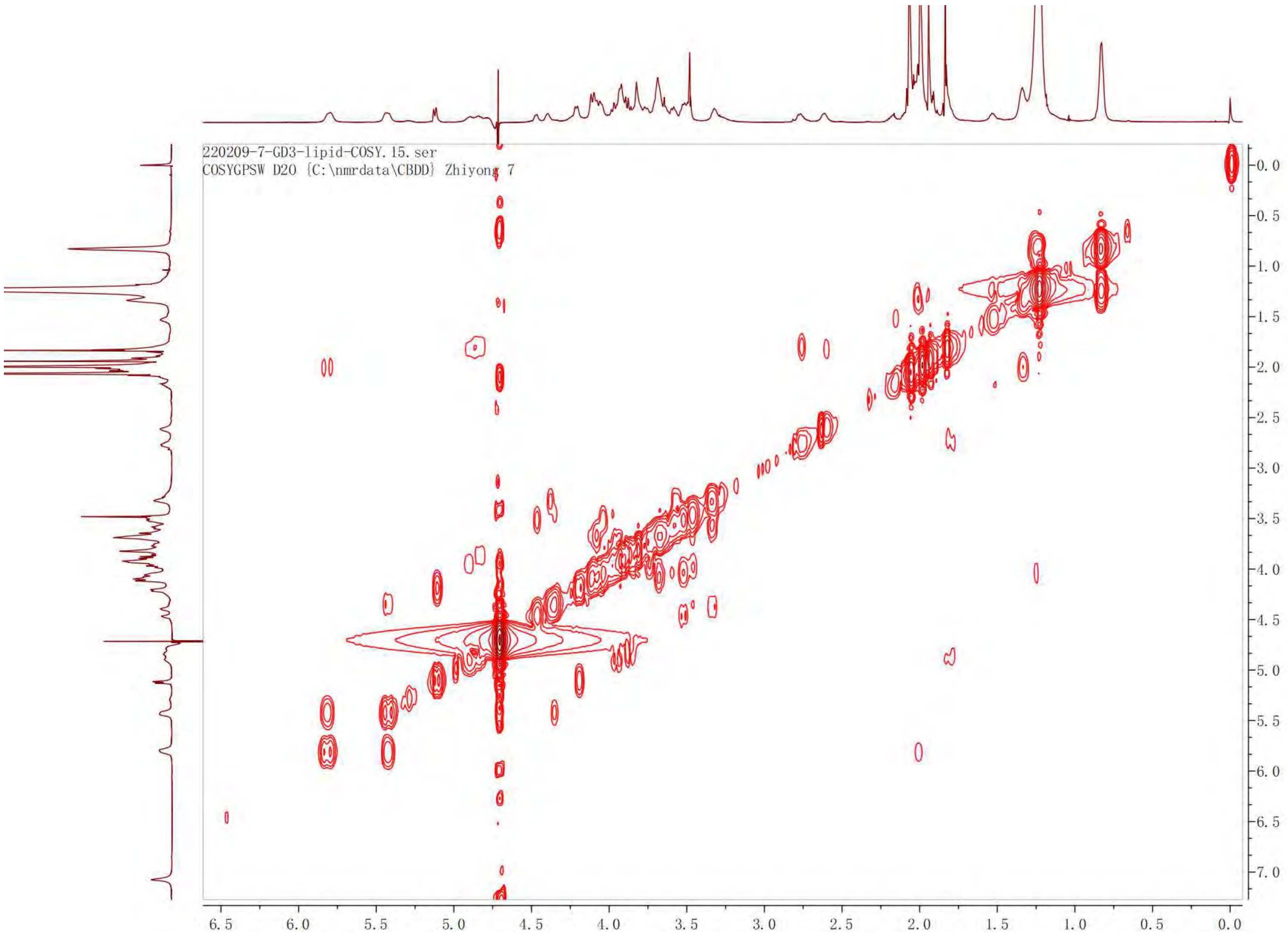

220209-7-GD3-lipid-COSY.15.ser
COSYGPSW D2O {C:\nmrdata\CBDD} Zhiyong 7

TOCSY of **4**

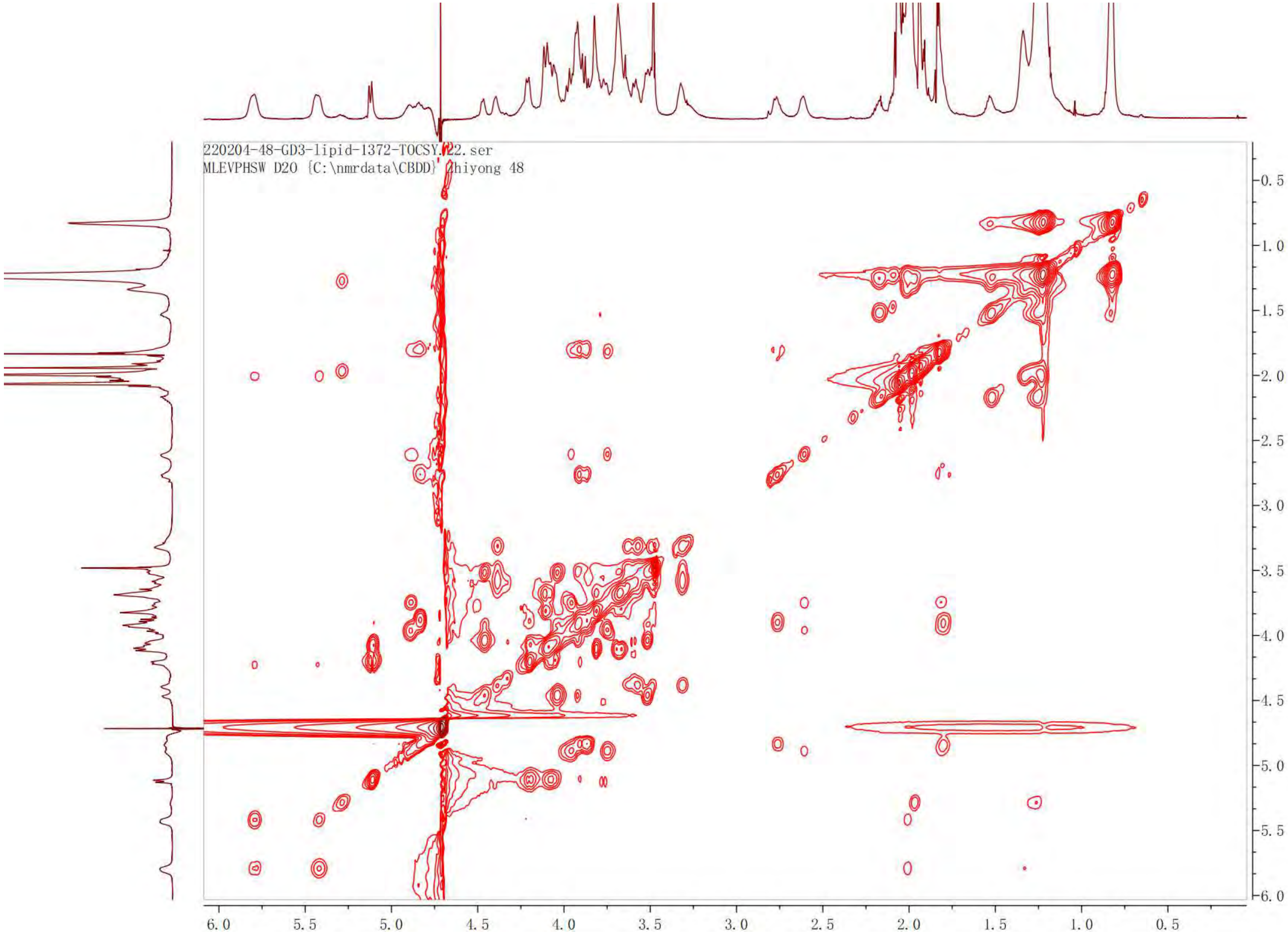

220204-48-GD3-lipid-1372-TOCSY.22.ser
MLEVPHSW D2O {C:\nmrdata\CBDD} zhiyong 48
0.5
1.0
1.5
2.0
2.5
3.0
3.5
4.0
4.5
5.0
5.5
6.0
6.0
5.5
5.0
4.5
4.0
3.5
3.0
2.5
2.0
1.5
1.0
0.5

HSQC of **4**

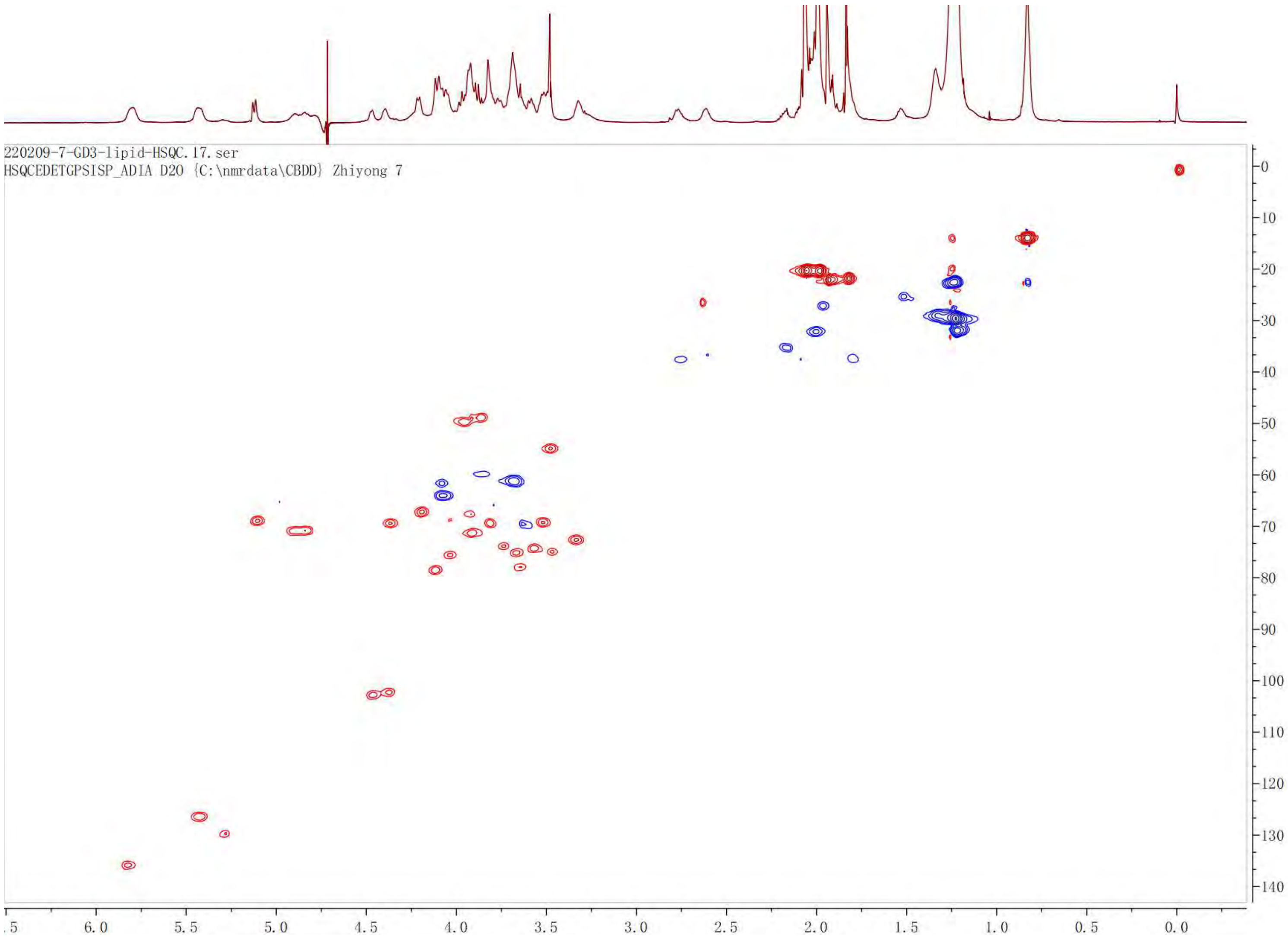
220209-7-GD3-lipid-HSQC.17.ser
HSQCEDETGPSISP_ADIA D2O {C:\nmrdata\CBDD} Zhiyong 7
.5
6.0
5.5
5.0
4.5
4.0
3.5
3.0
2.5
2.0
1.5
1.0
0.5
0.0
0
10
20
30
40
50
60
70
80
90
100
110
120
130
140

[1]H NMR of **16**. 600 MHz, $CD_3OD$.

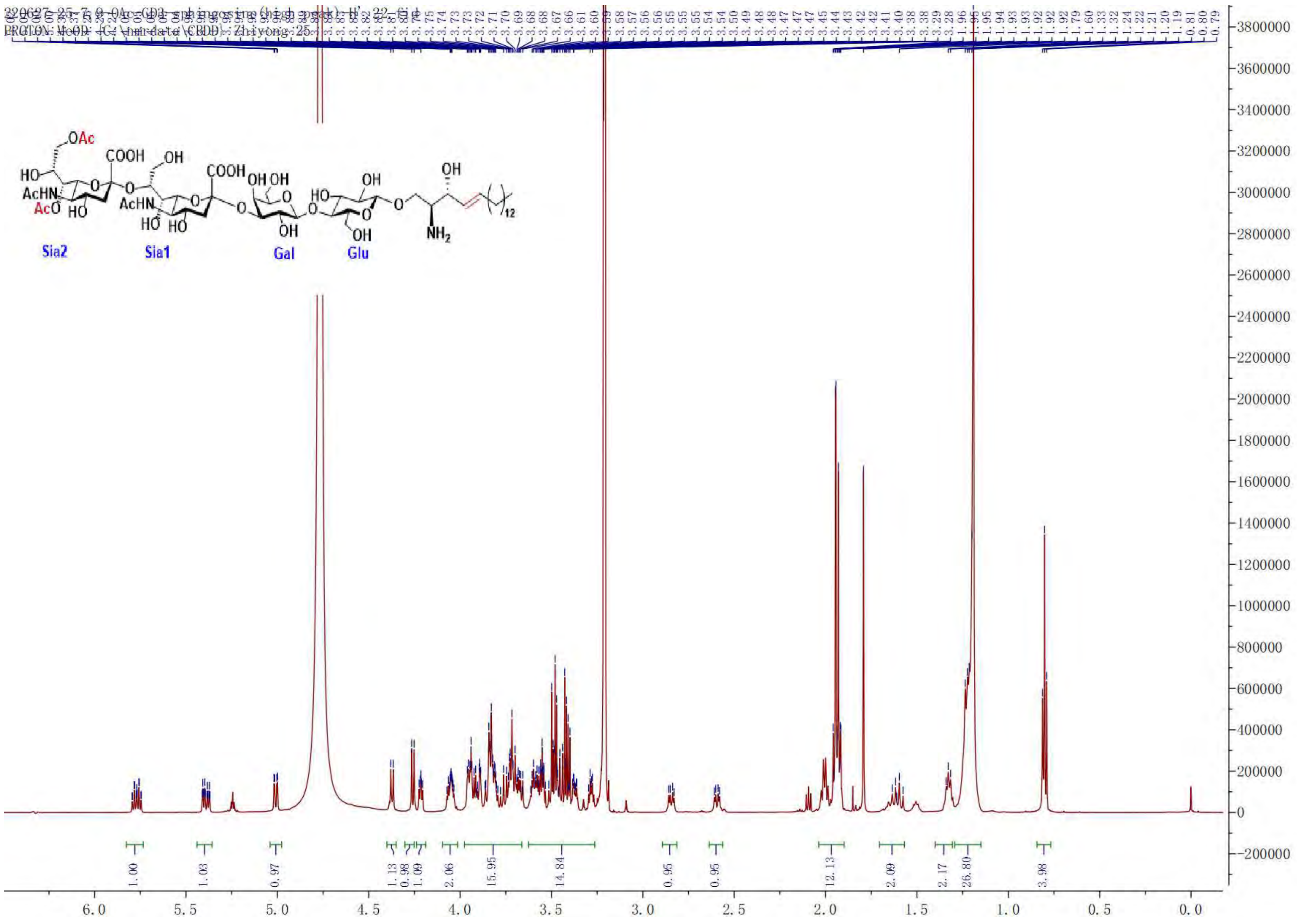

COSY of **16**

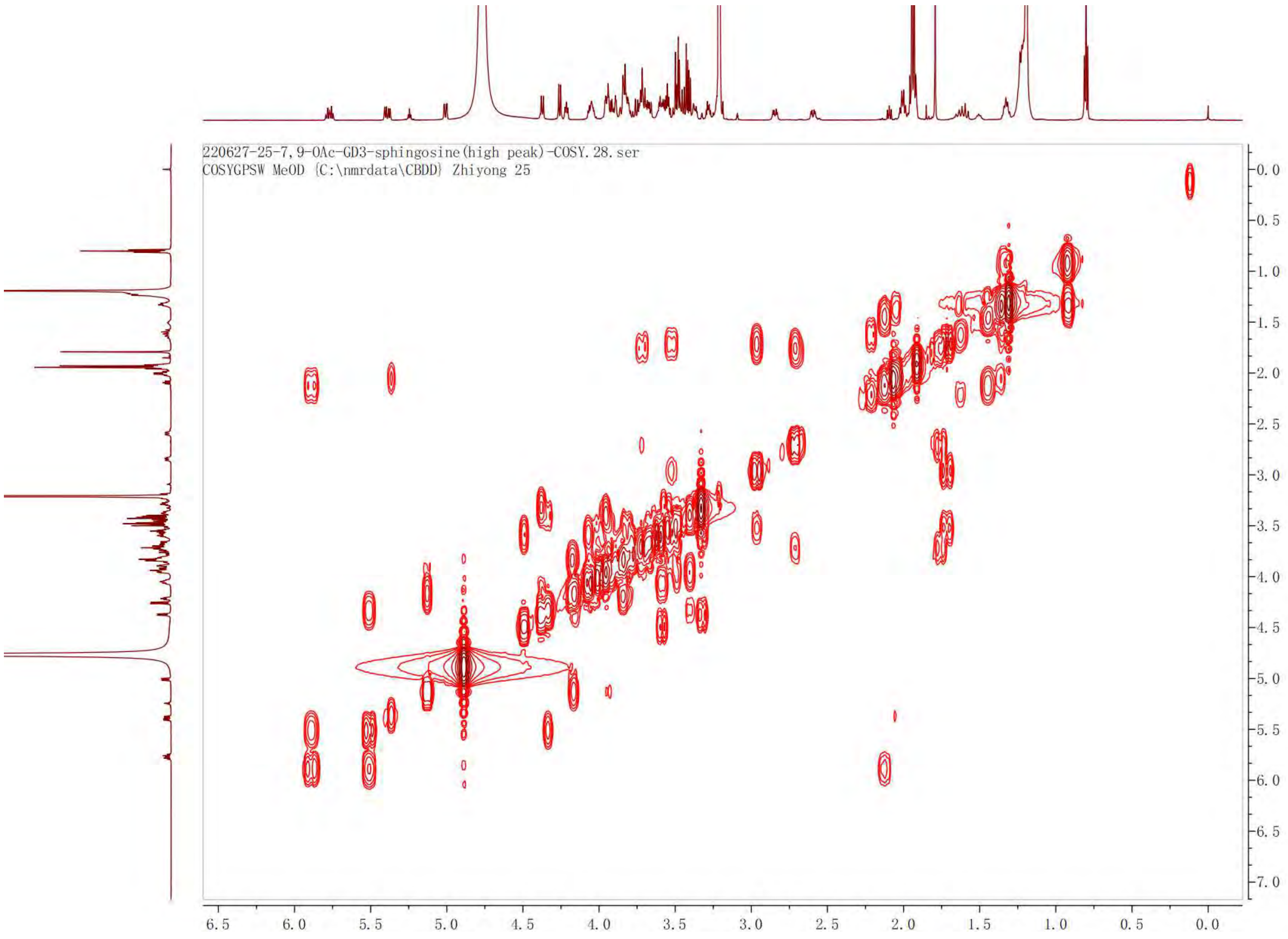

220627-25-7,9-OAc-GD3-sphingosine(high peak)-COSY.28.ser
COSYGPSW MeOD {C:\nmrdata\CBDD} Zhiyong 25

TOCSY of **16**

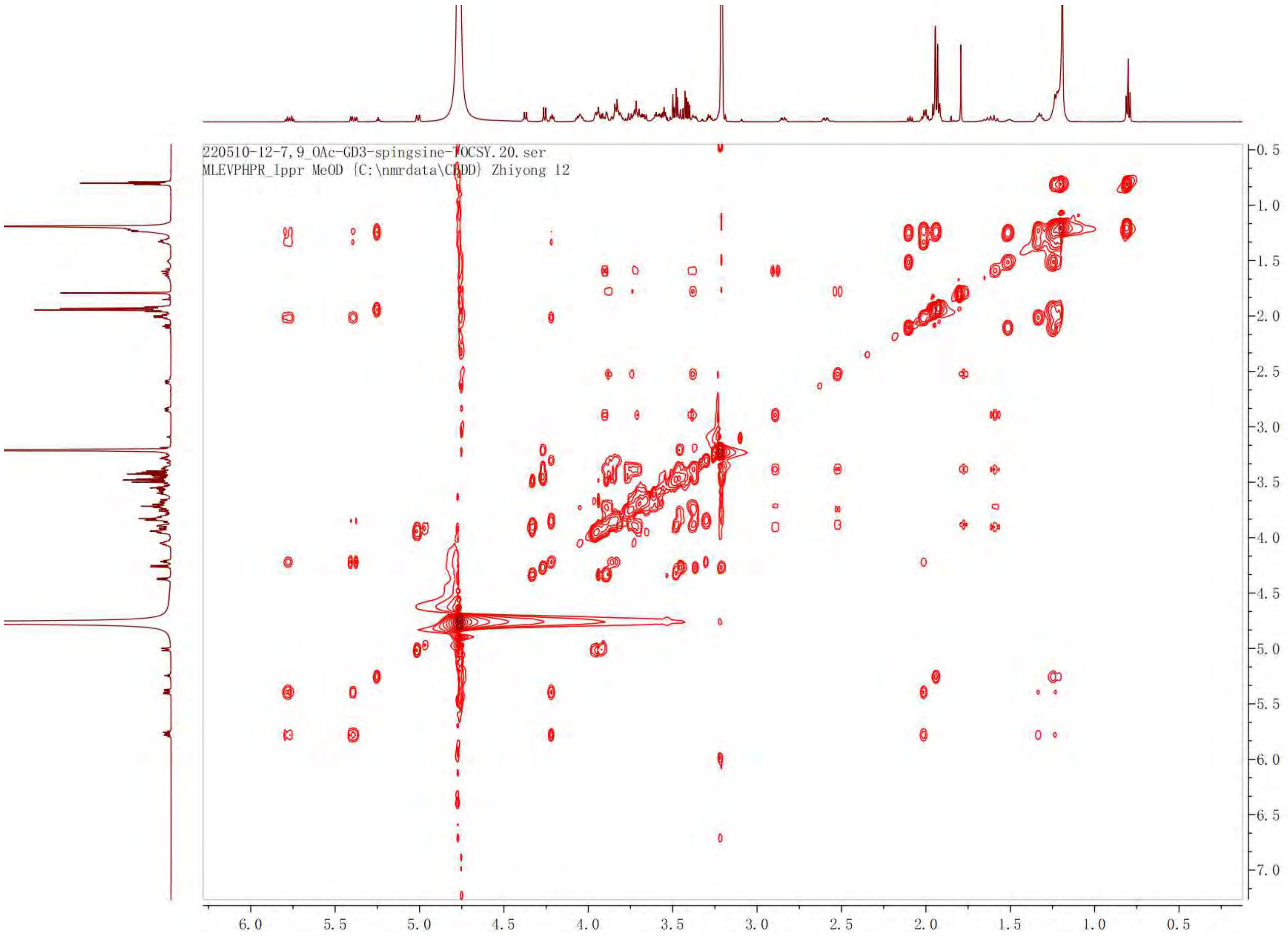
220510-12-7,9_OAc-GD3-spingsine-TOCSY.20.ser
MLEVPHPR_1ppr MeOD {C:\nmrdata\CDDD} Zhiyong 12
6.0 5.5 5.0 4.5 4.0 3.5 3.0 2.5 2.0 1.5 1.0 0.5
0.5 1.0 1.5 2.0 2.5 3.0 3.5 4.0 4.5 5.0 5.5 6.0 6.5 7.0

HSQC of **16**

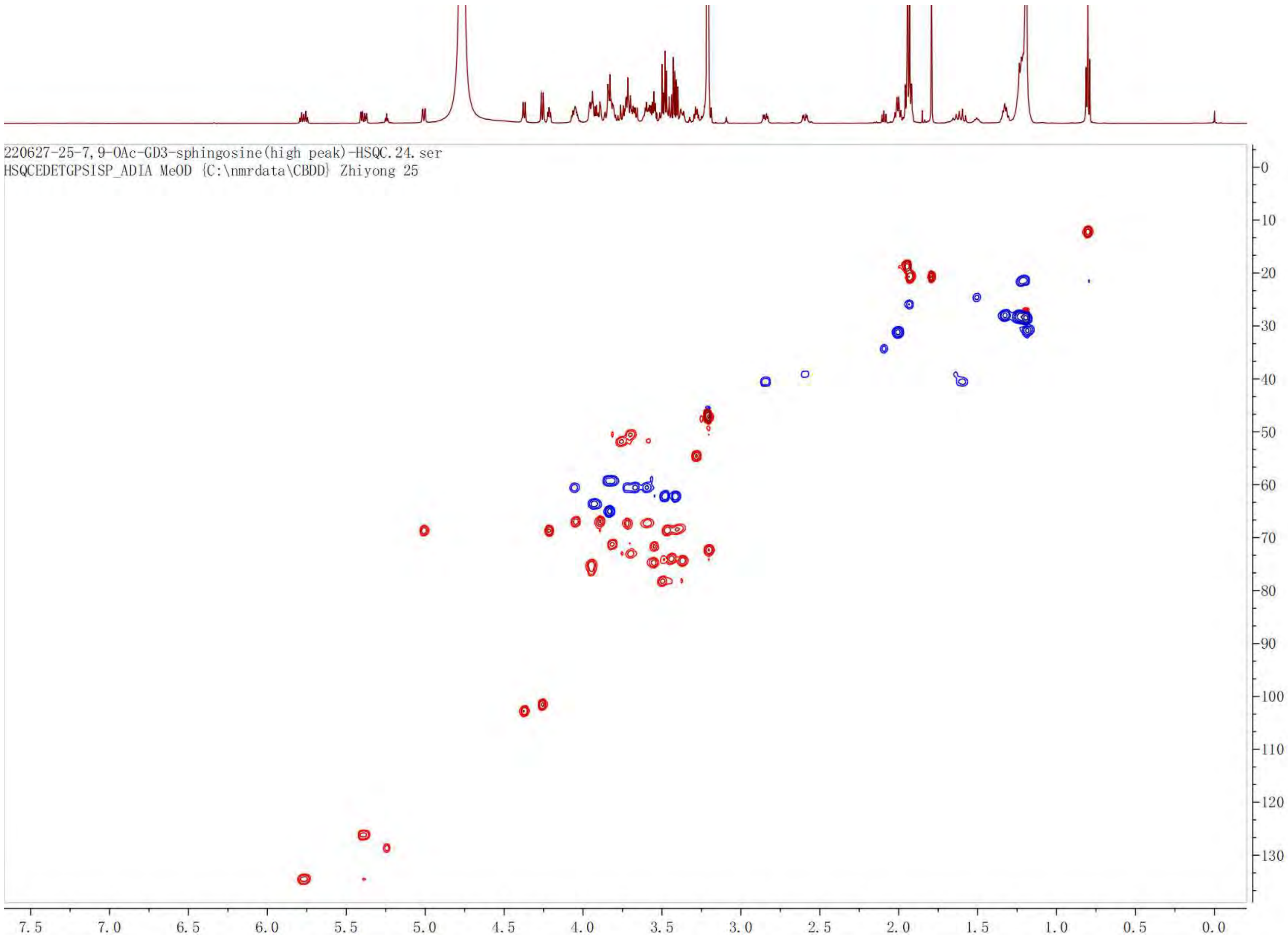

220627-25-7,9-OAc-GD3-sphingosine(high peak)-HSQC.24.ser
HSQCEDETGPSISP_ADIA MeOD {C:\nmrdata\CBDD} Zhiyong 25

HMBC of **16**

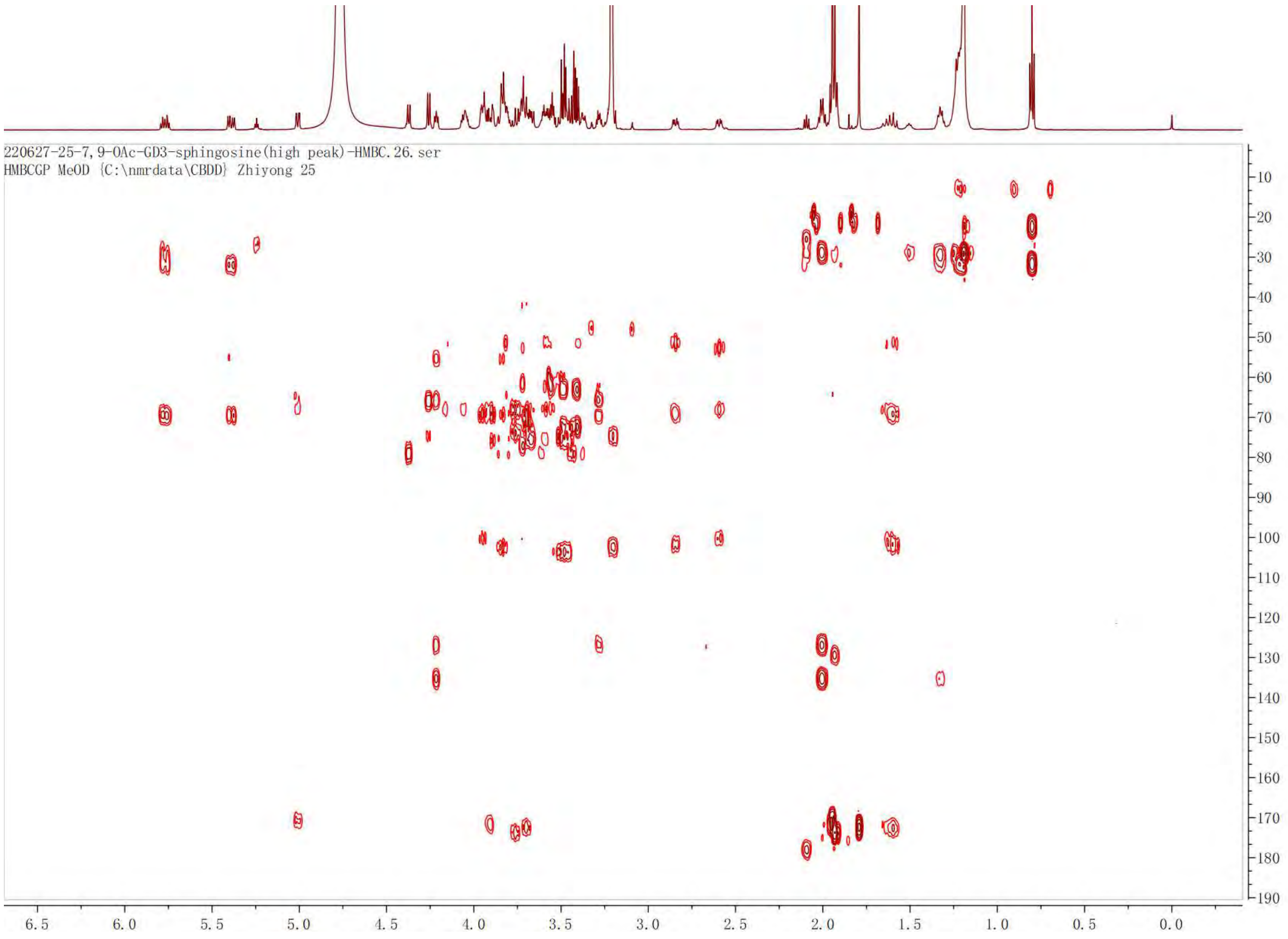
220627-25-7,9-OAc-GD3-sphingosine(high peak)-HMBC.26.ser
HMBCGP MeOD {C:\nmrdata\CBDD} Zhiyong 25
6.5 6.0 5.5 5.0 4.5 4.0 3.5 3.0 2.5 2.0 1.5 1.0 0.5 0.0
10 20 30 40 50 60 70 80 90 100 110 120 130 140 150 160 170 180 190

[1]H NMR of **17**. 600 MHz, $CD_3OD$.

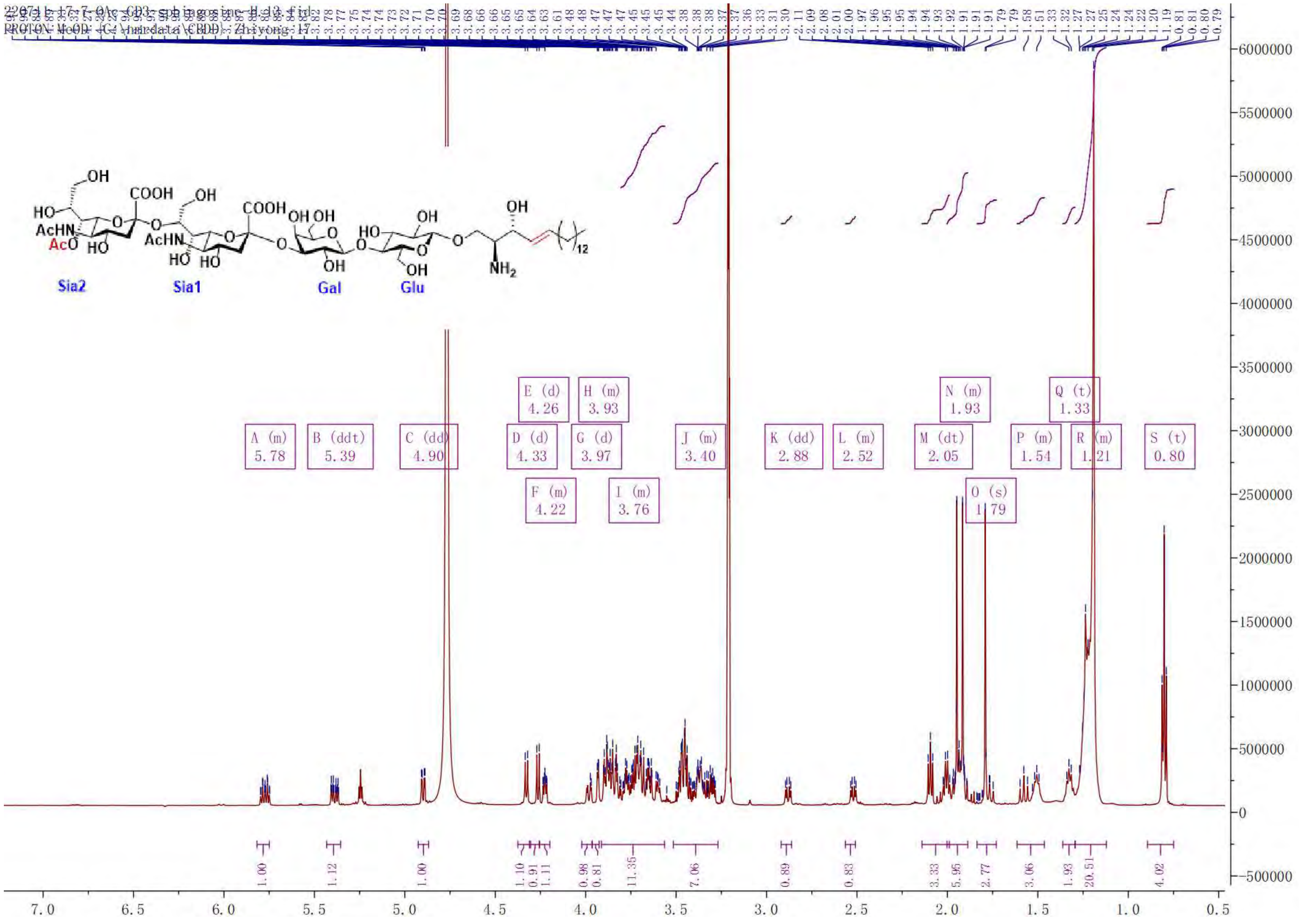

COSY of **17**

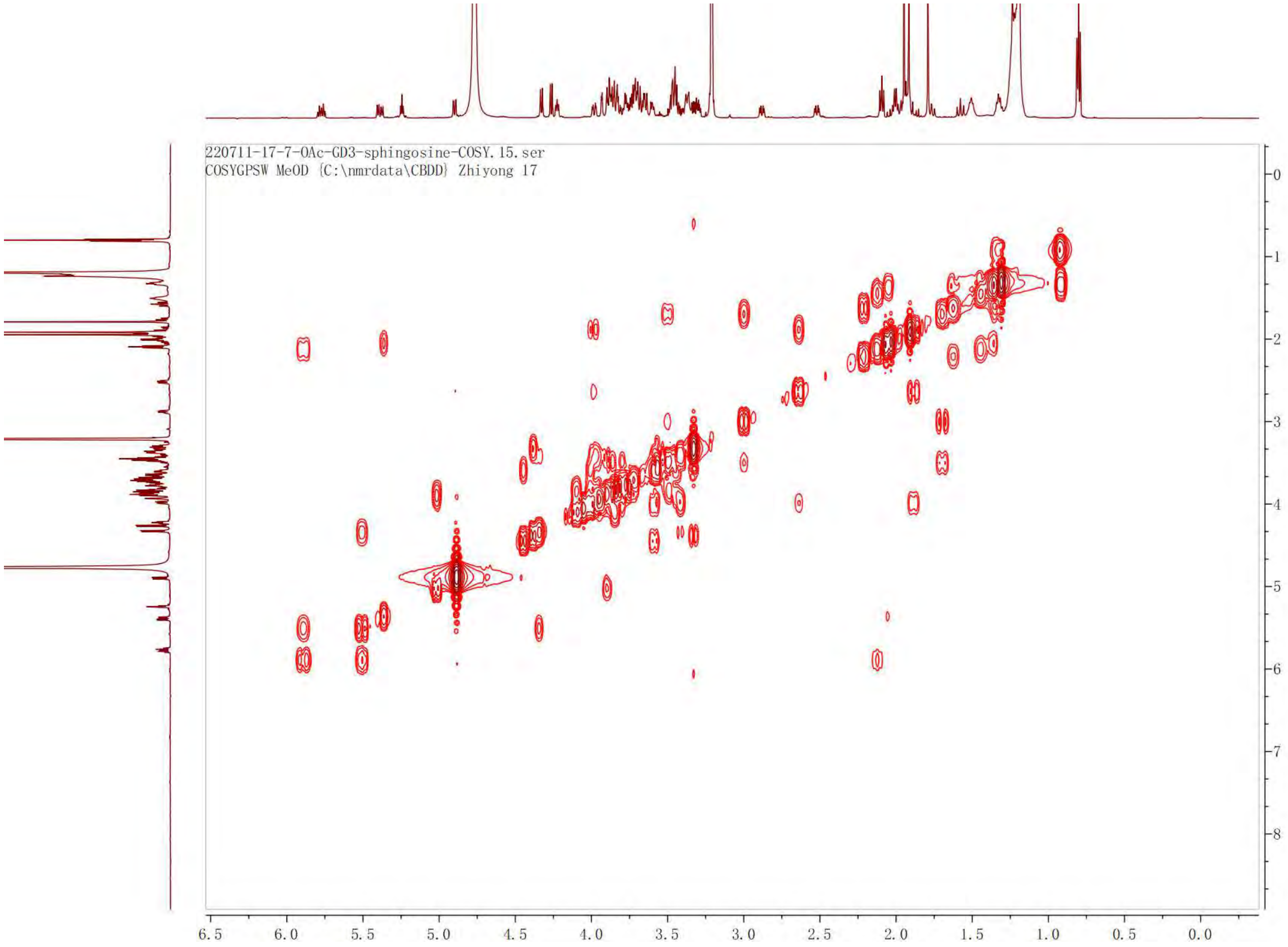

220711-17-7-OAc-GD3-sphingosine-COSY.15.ser
COSYGPSW MeOD {C:\nmrdata\CBDD} Zhiyong 17
6.5
6.0
5.5
5.0
4.5
4.0
3.5
3.0
2.5
2.0
1.5
1.0
0.5
0.0
0
1
2
3
4
5
6
7
8

TOCSY of **17**

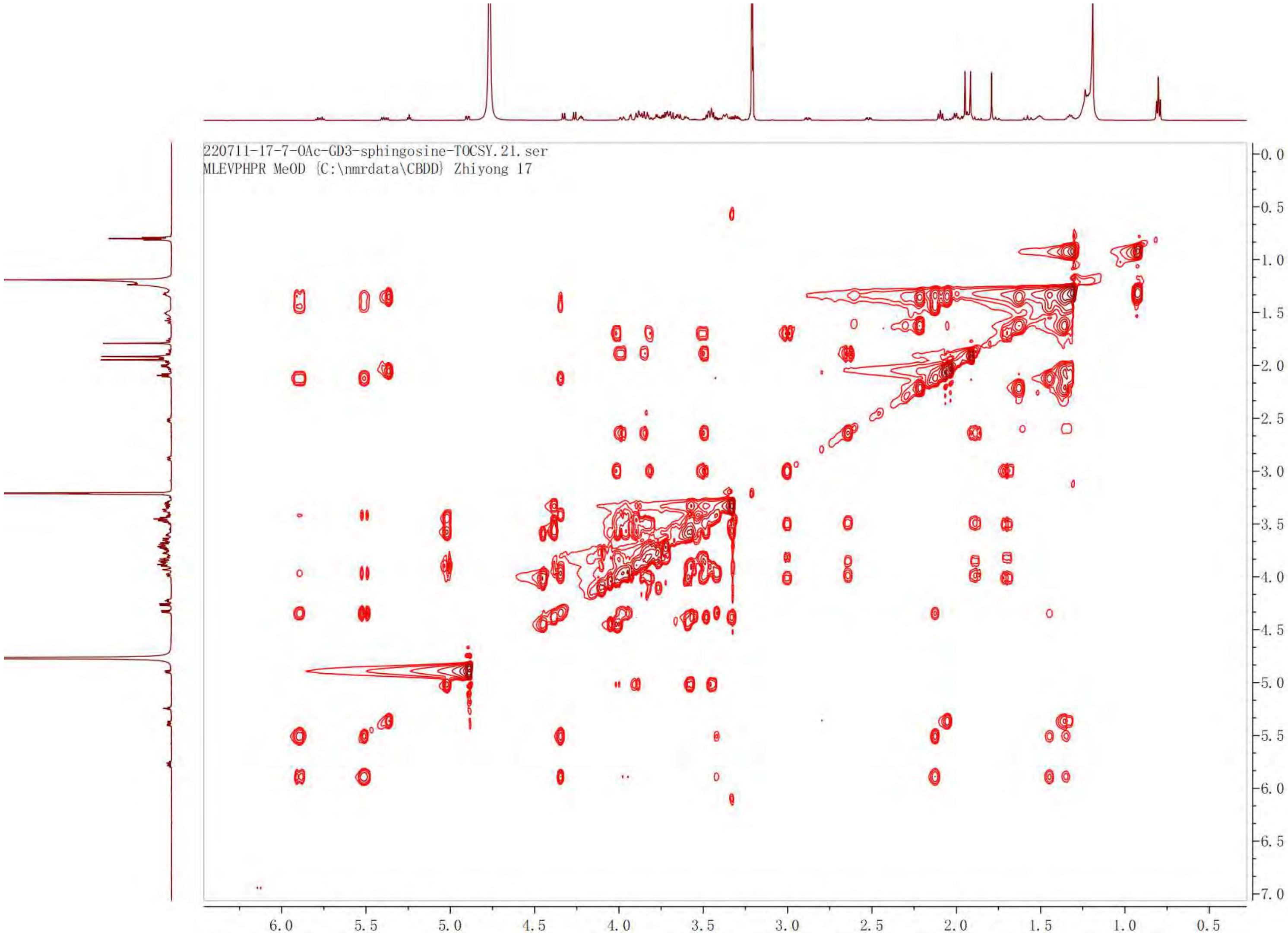

220711-17-7-OAc-GD3-sphingosine-TOCSY.21.ser
MLEVPHPR MeOD {C:\nmrdata\CBDD} Zhiyong 17
0.0
0.5
1.0
1.5
2.0
2.5
3.0
3.5
4.0
4.5
5.0
5.5
6.0
6.5
7.0
6.0
5.5
5.0
4.5
4.0
3.5
3.0
2.5
2.0
1.5
1.0
0.5

HSQC of **17**

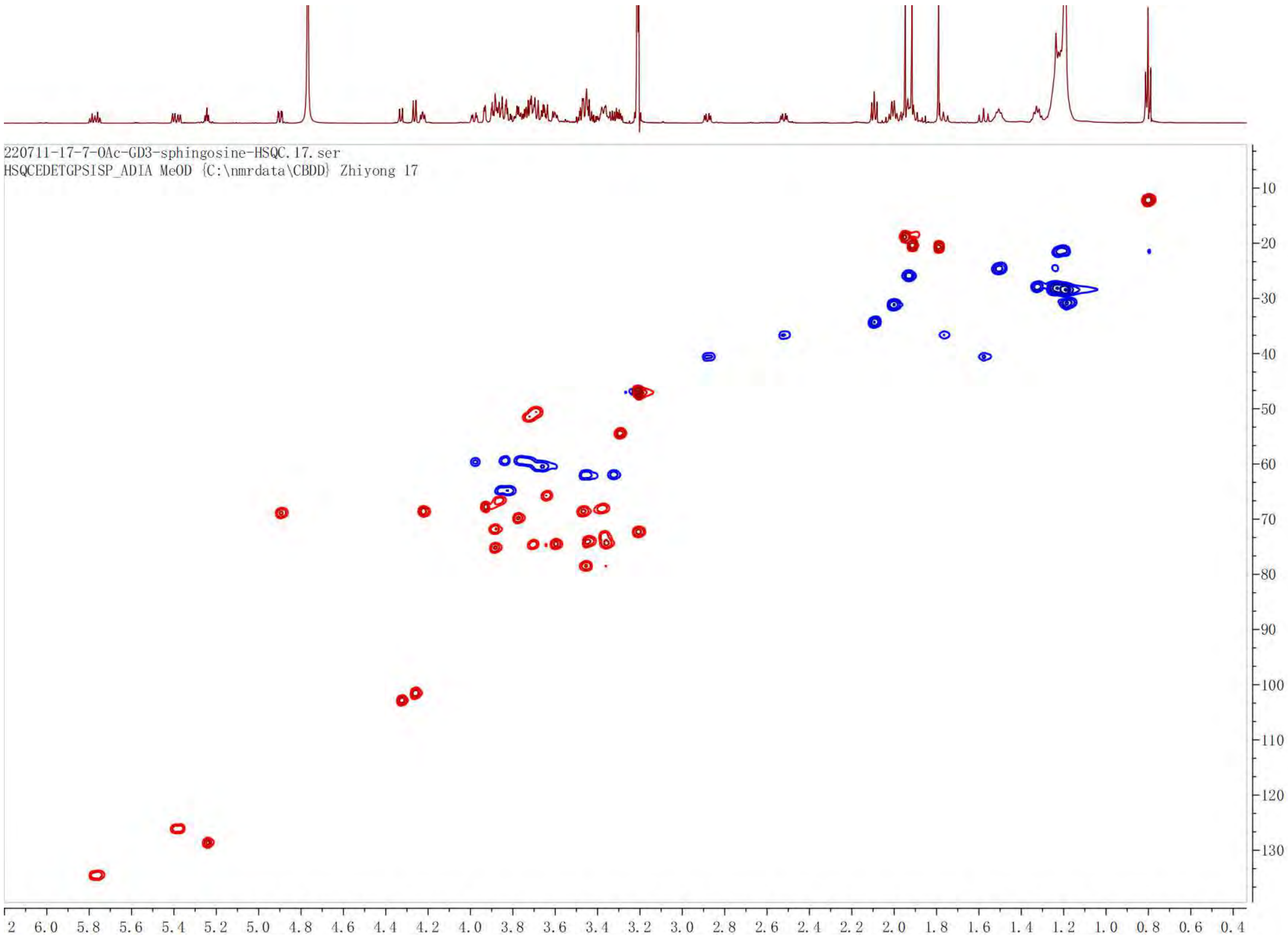

220711-17-7-OAc-GD3-sphingosine-HSQC.17.ser
HSQCEDETGPSISP_ADIA MeOD {C:\nmrdata\CBDD} Zhiyong 17

HMBC of **17**

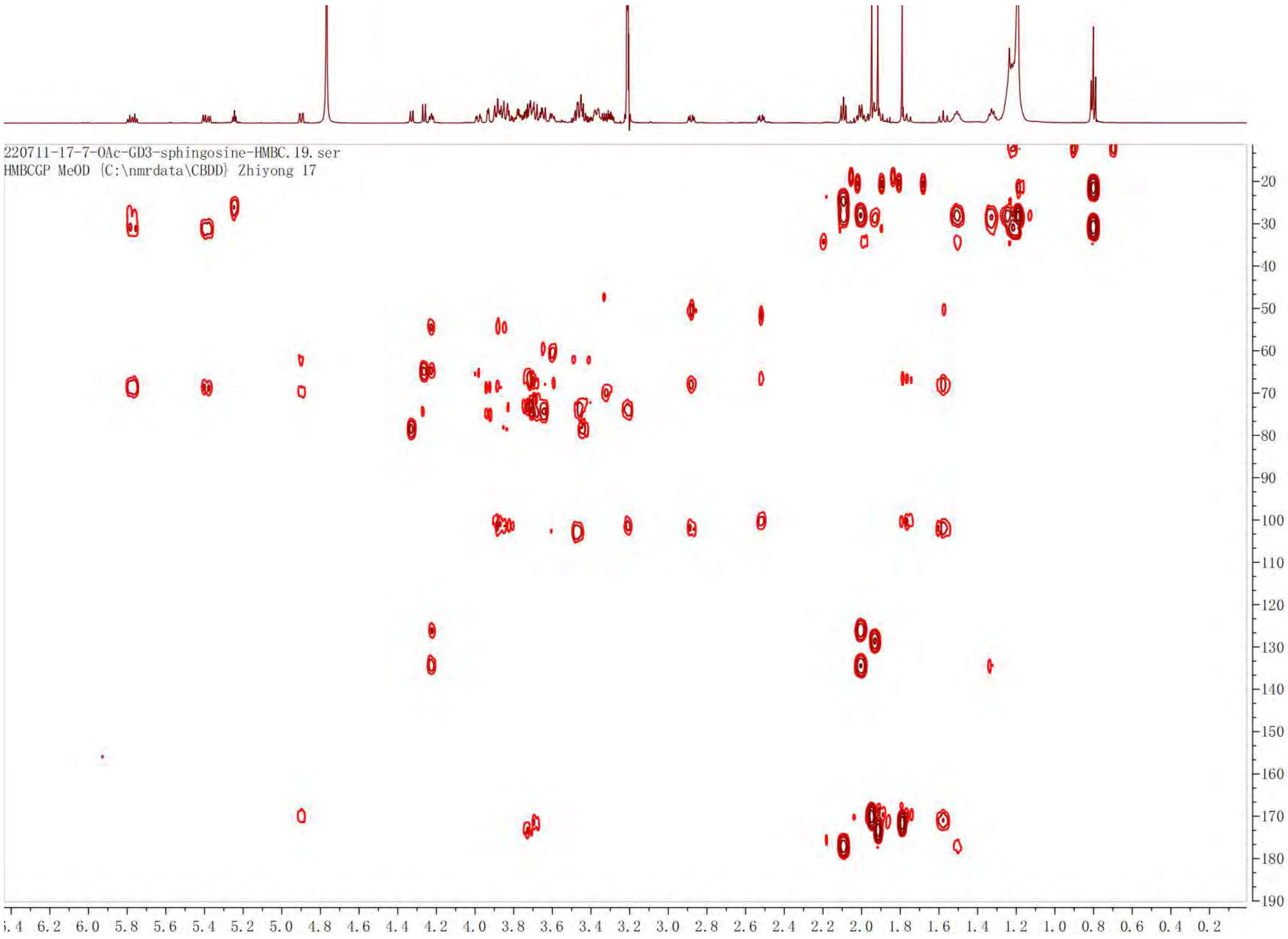

220711-17-7-OAc-GD3-sphingosine-HMBC.19.ser
HMBCGP MeOD {C:\nmrdata\CBDD} Zhiyong 17

[1]H NMR of **1**. 600 MHz, $CD_3OD$.

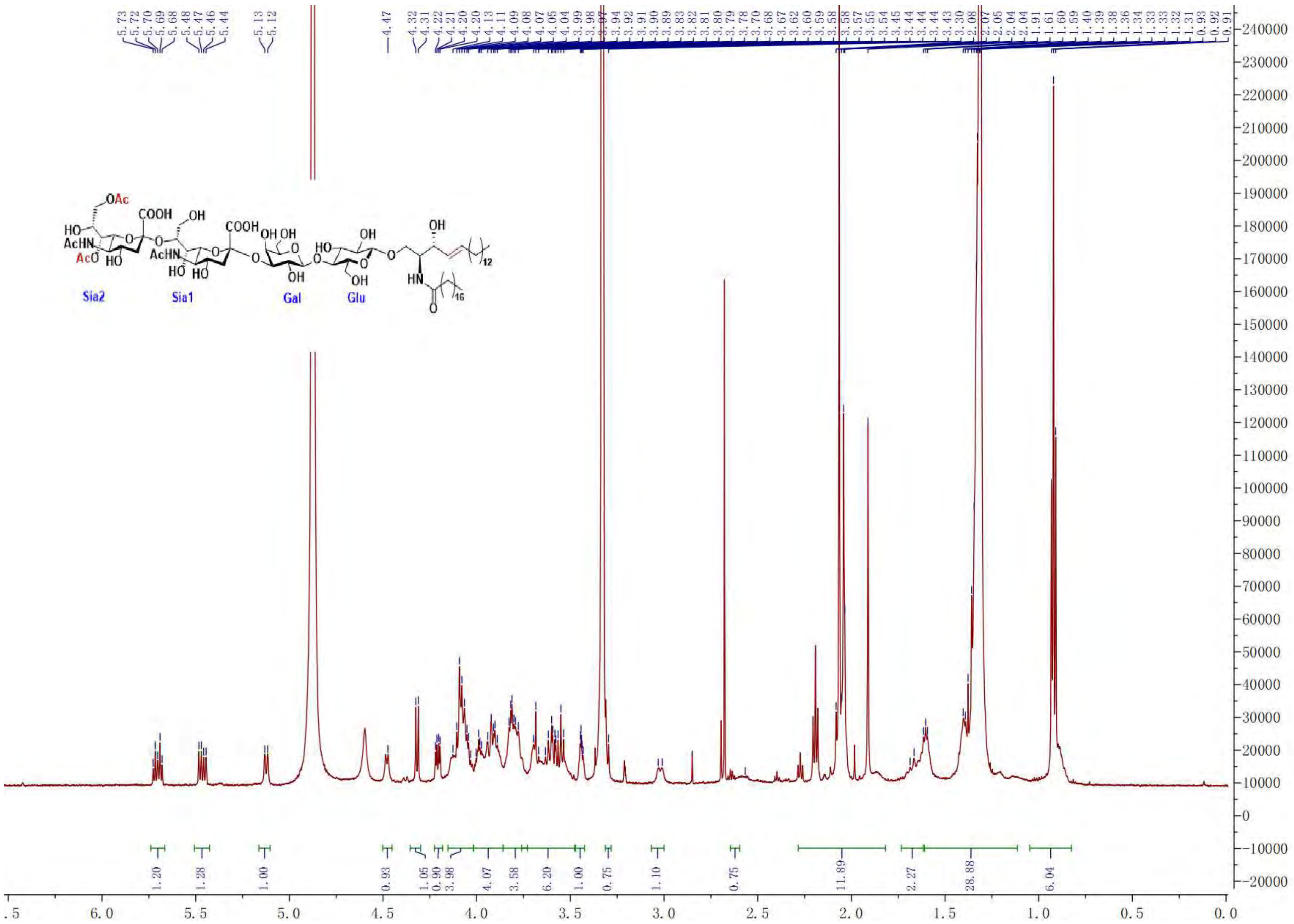

COSY of **1**

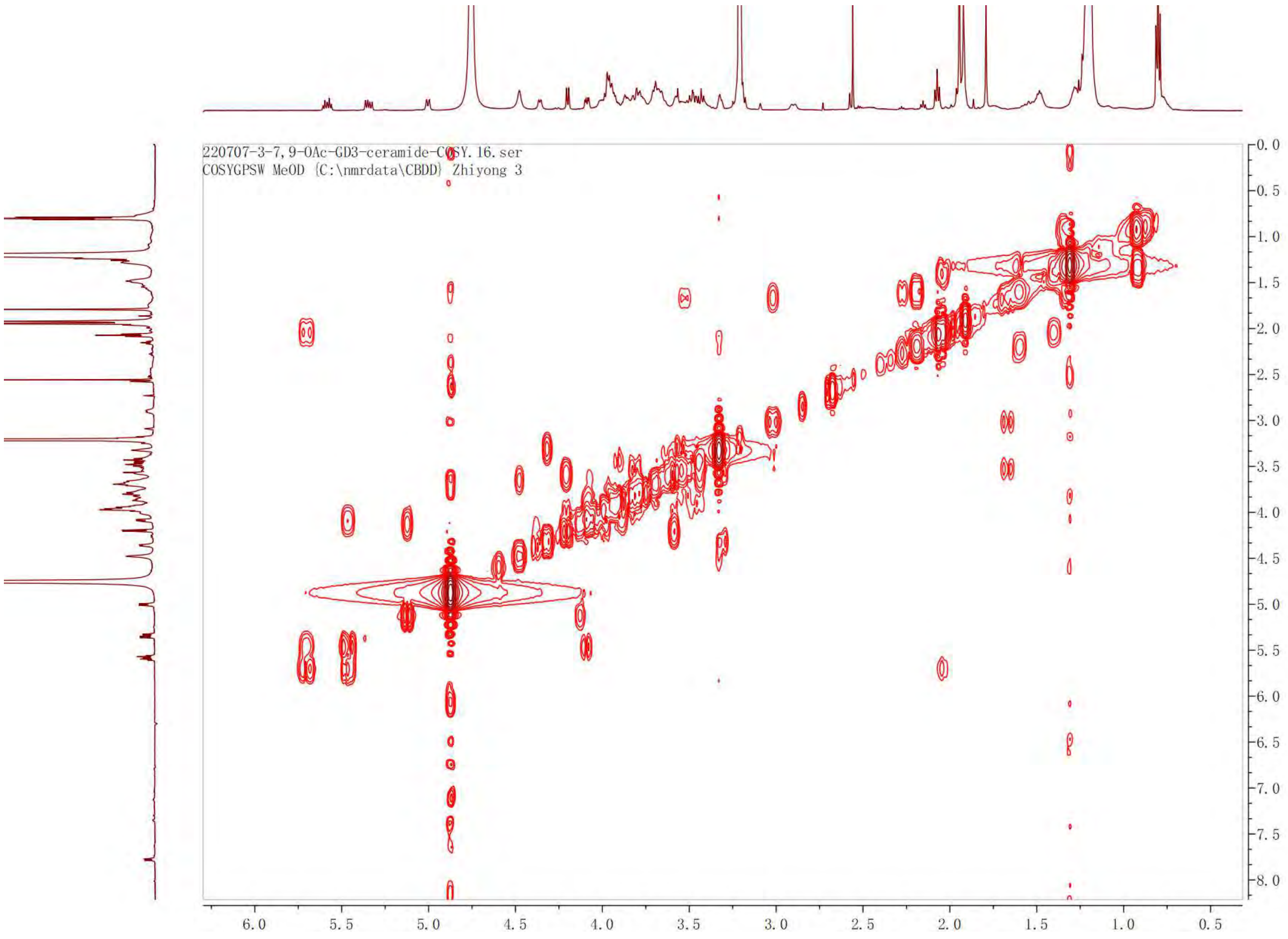

220707-3-7,9-OAc-GD3-ceramide-COSY.16.ser
COSYGPSW MeOD {C:\nmrdata\CBDD} Zhiyong 3
6.0
5.5
5.0
4.5
4.0
3.5
3.0
2.5
2.0
1.5
1.0
0.5
0.0
-0.5
-1.0
-1.5
-2.0
-2.5
-3.0
-3.5
-4.0
-4.5
-5.0
-5.5
-6.0
-6.5
-7.0
-7.5
-8.0

HSQC of **1**

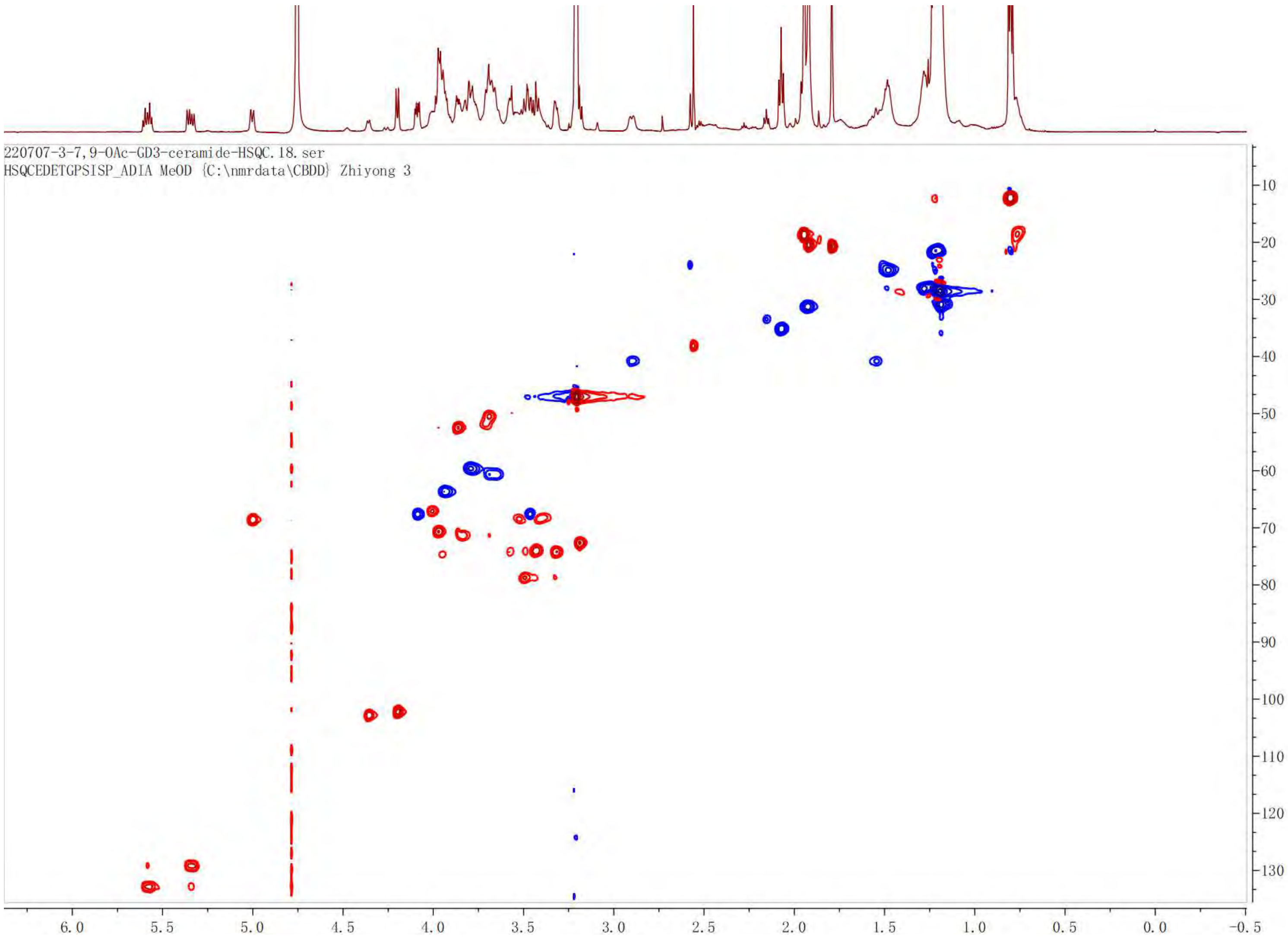
220707-3-7,9-OAc-GD3-ceramide-HSQC.18.ser
HSQCEDETGPSISP_ADIA MeOD {C:\nmrdata\CBDD} Zhiyong 3
6.0
5.5
5.0
4.5
4.0
3.5
3.0
2.5
2.0
1.5
1.0
0.5
0.0
-0.5
10
20
30
40
50
60
70
80
90
100
110
120
130

[1]H NMR of **2**. 600 MHz, $CD_3OD$.

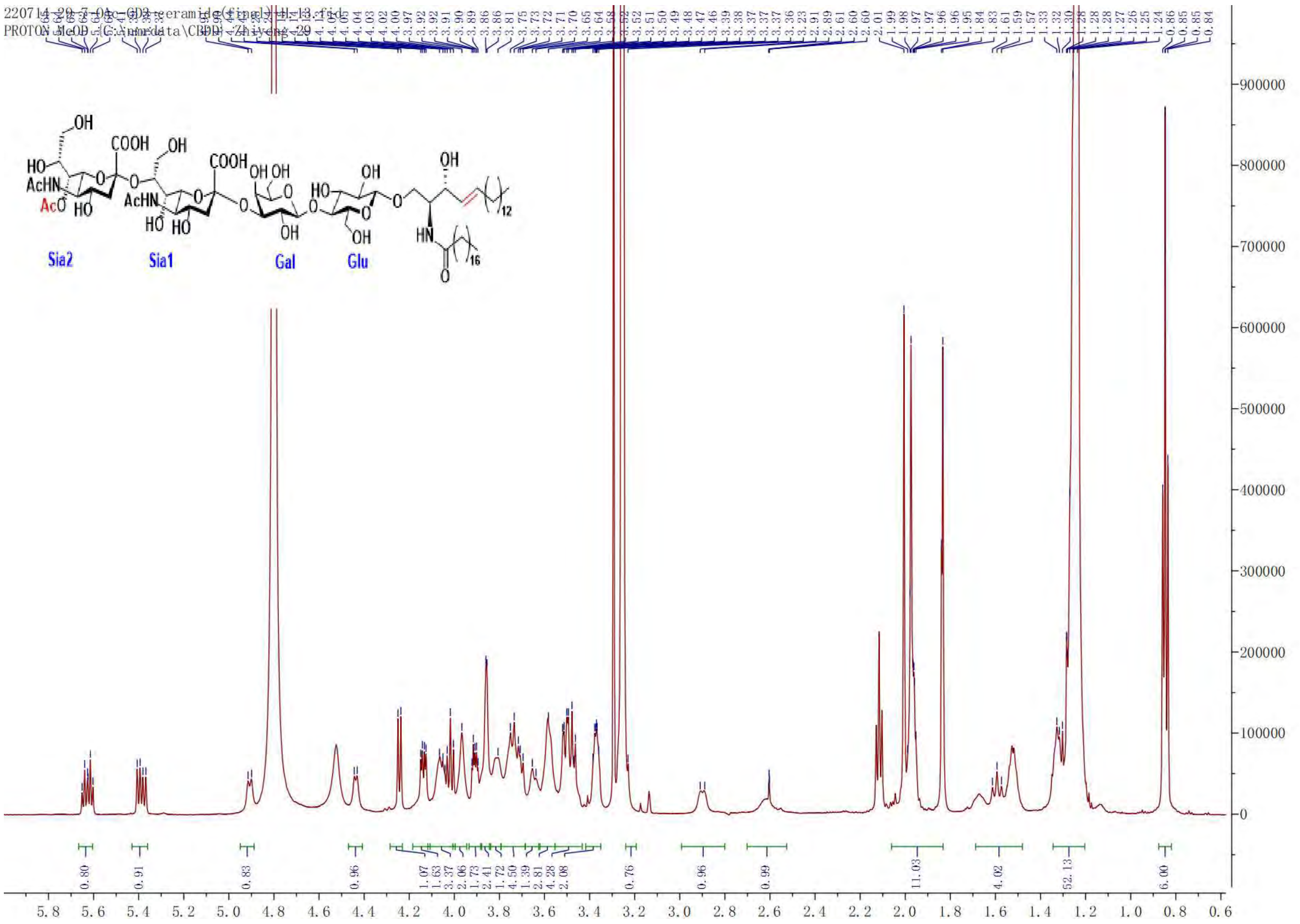

COSY of **2**

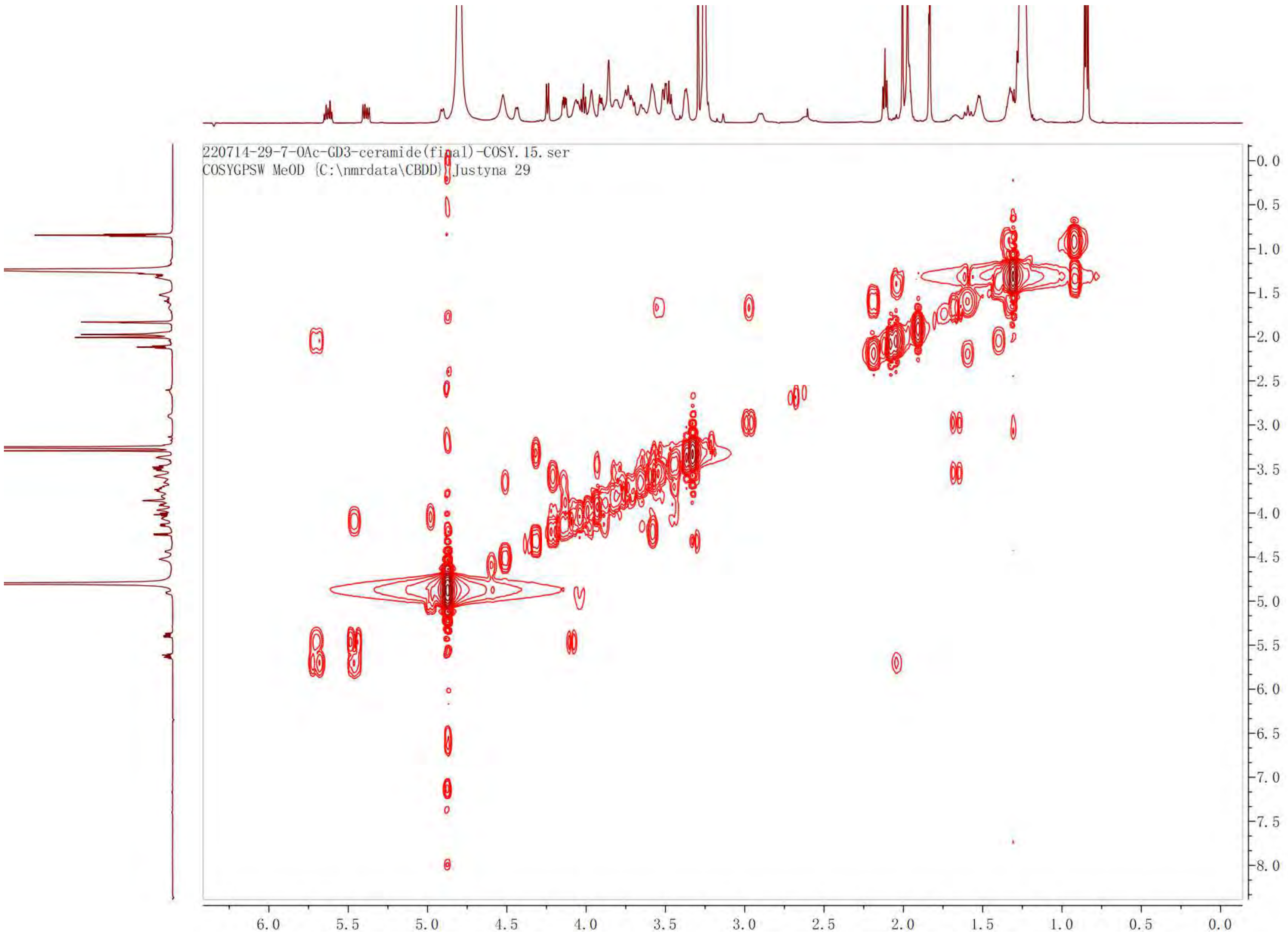

220714-29-7-OAc-GD3-ceramide(final)-COSY.15.ser
COSYGPSW MeOD {C:\nmrdata\CBDD} Justyna 29

TOCSY of **2**

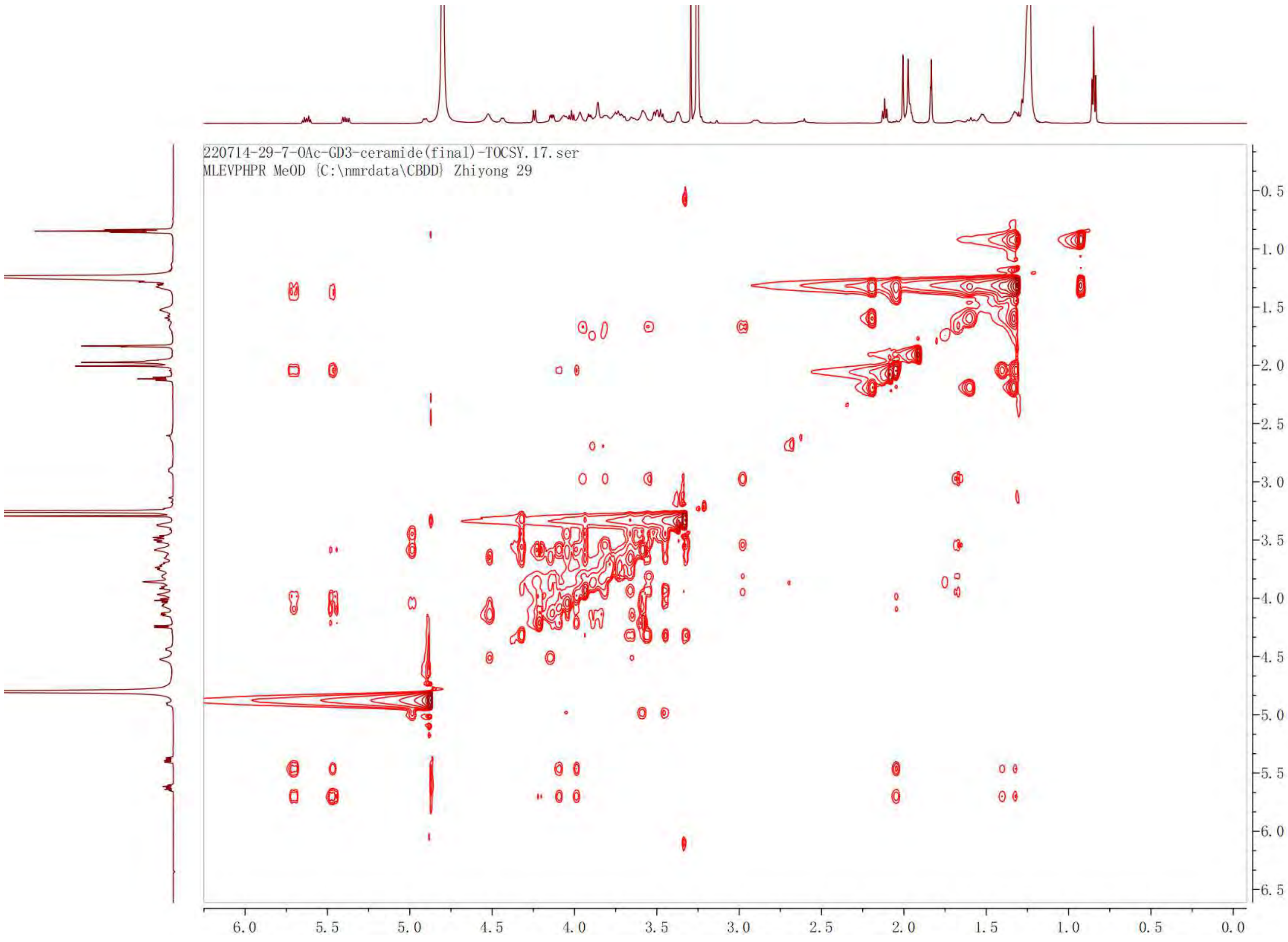
220714-29-7-OAc-GD3-ceramide(final)-TOCSY.17.ser
MLEVPHPR MeOD {C:\nmrdata\CBDD} Zhiyong 29
0.5
1.0
1.5
2.0
2.5
3.0
3.5
4.0
4.5
5.0
5.5
6.0
6.5
6.0
5.5
5.0
4.5
4.0
3.5
3.0
2.5
2.0
1.5
1.0
0.5
0.0

HSQC of **2**

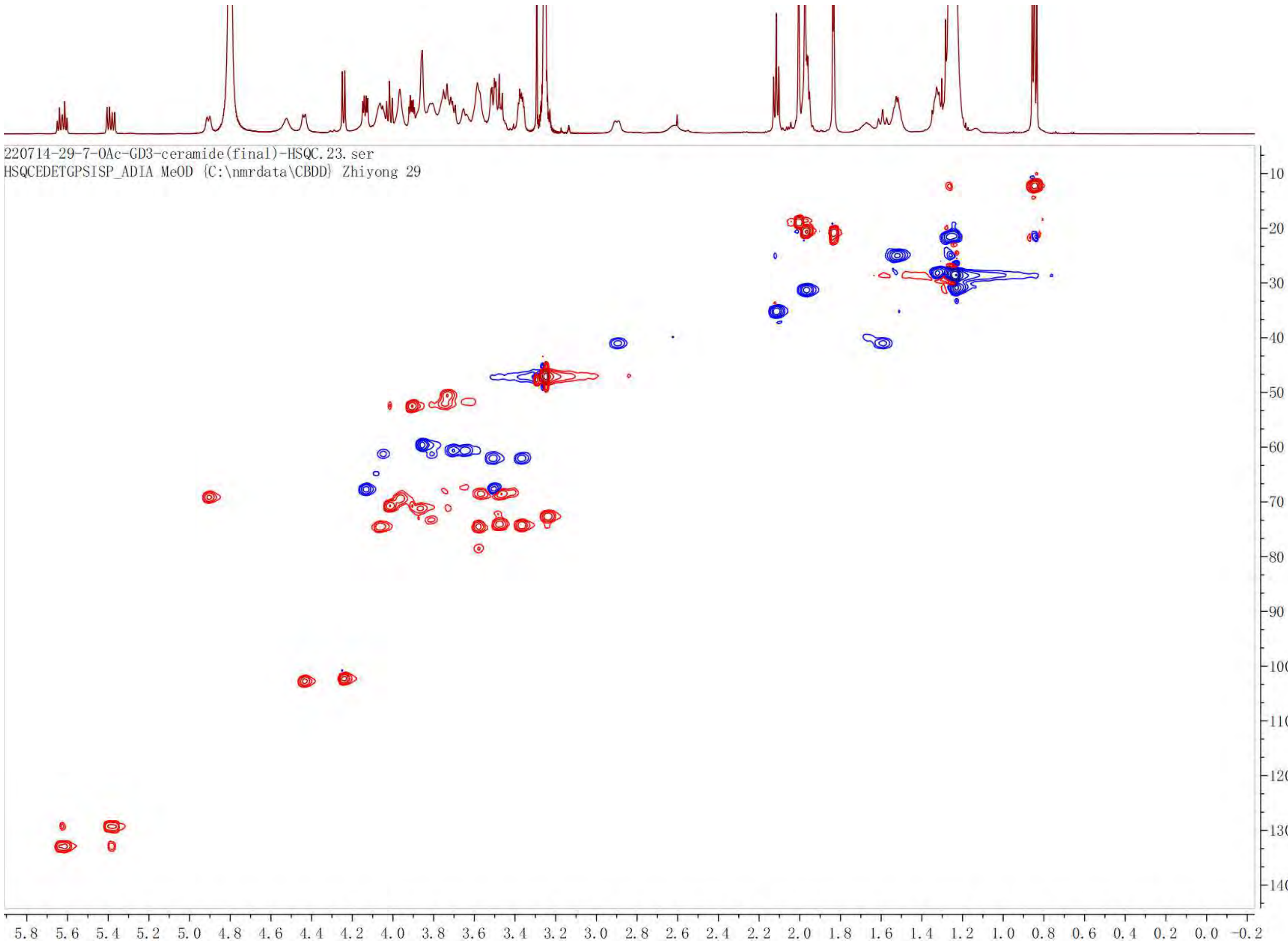

220714-29-7-OAc-GD3-ceramide(final)-HSQC.23.ser
HSQCEDETGPSISP_ADIA MeOD {C:\nmrdata\CBDD} Zhiyong 29

$^{1}$H NMR of **3**. 600 MHz, $CD_3OD$.

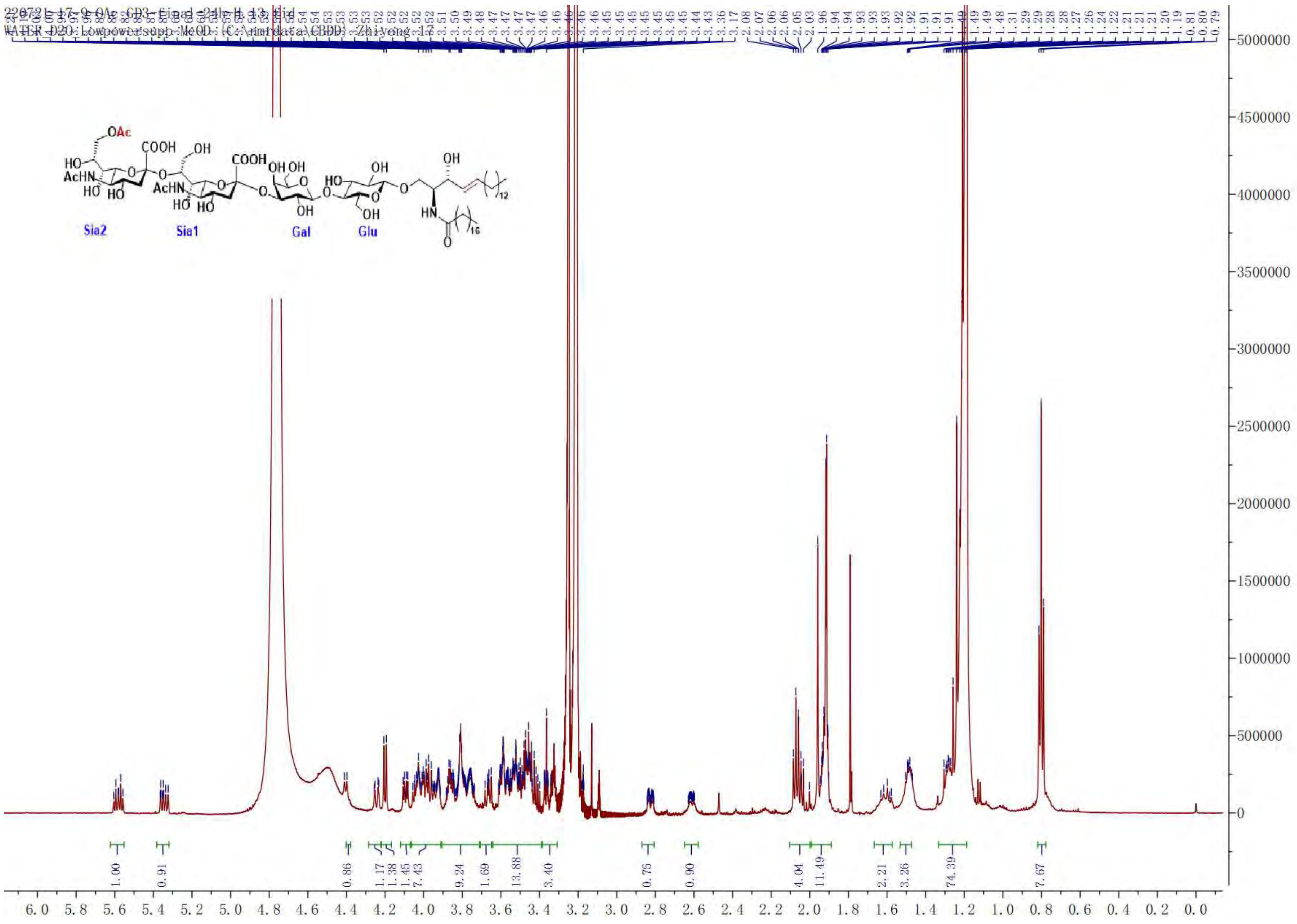

COSY of **3**

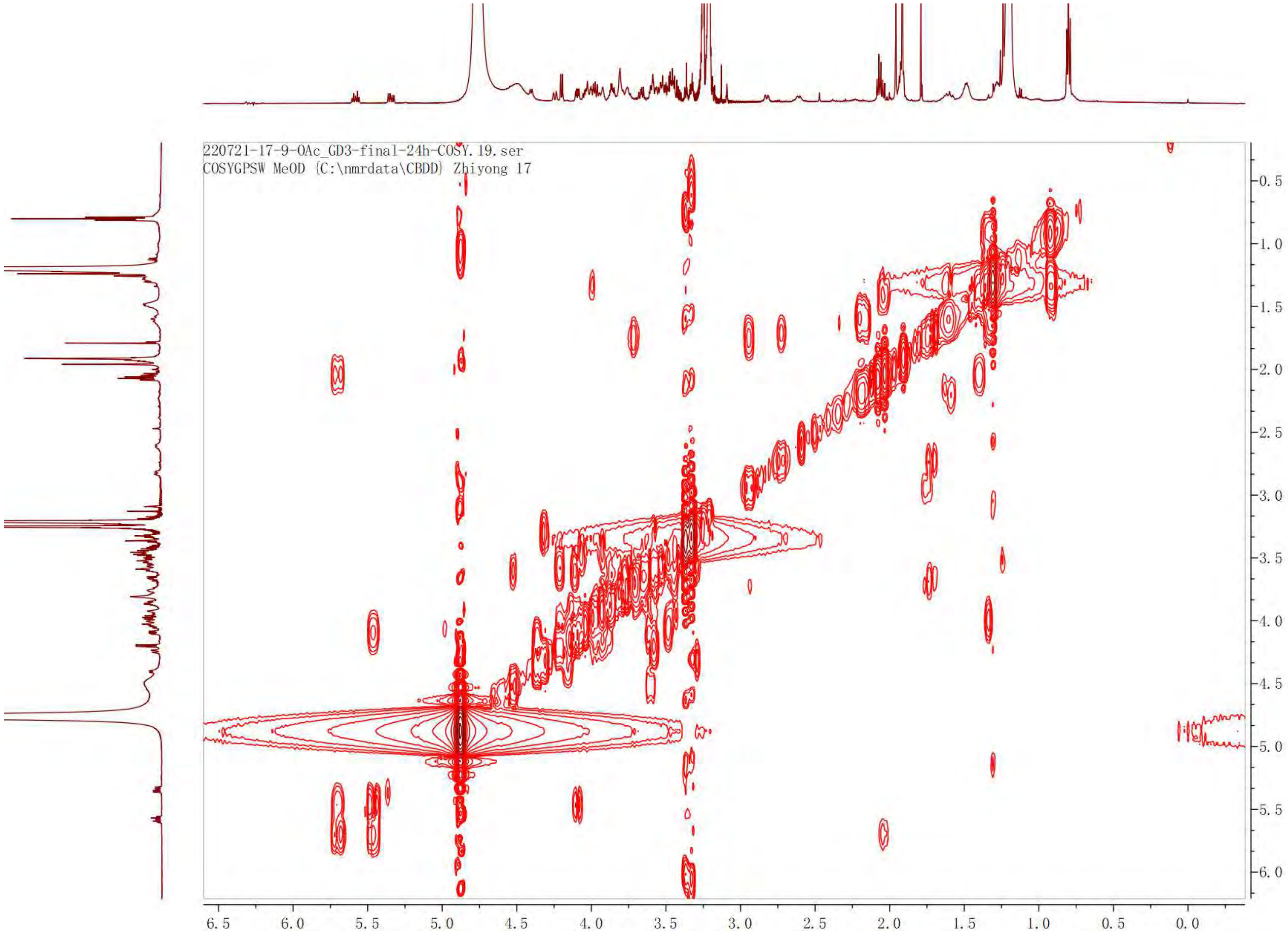

220721-17-9-OAc_GD3-final-24h-COSY.19.ser
COSYGPSW MeOD {C:\nmrdata\CBDD} Zhiyong 17
-0.5
-1.0
-1.5
-2.0
-2.5
-3.0
-3.5
-4.0
-4.5
-5.0
-5.5
-6.0
6.5
6.0
5.5
5.0
4.5
4.0
3.5
3.0
2.5
2.0
1.5
1.0
0.5
0.0

TOCSY of **3**

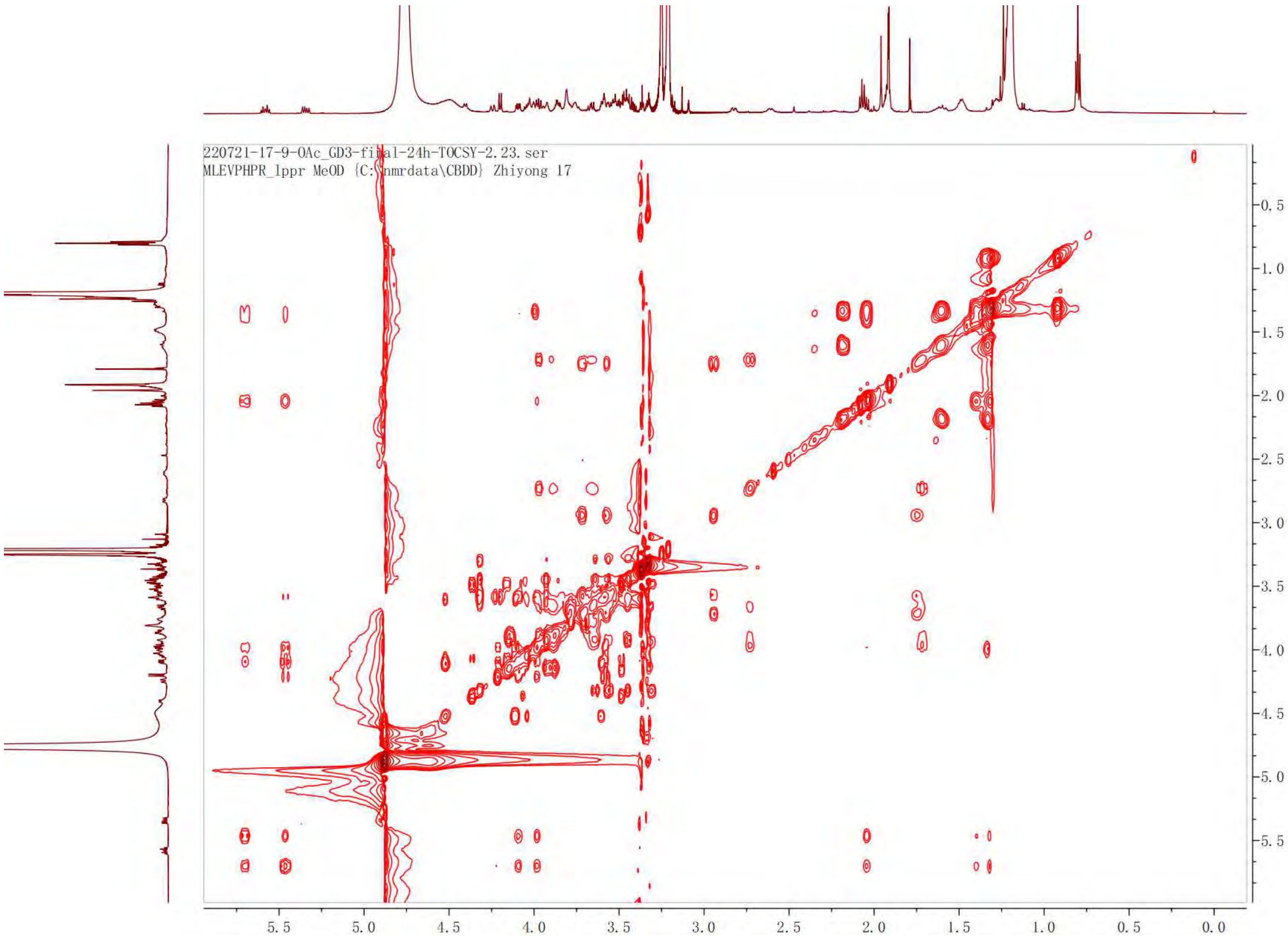

220721-17-9-OAc_GD3-final-24h-TOCSY-2.23.ser
MLEVPHPR_1ppr MeOD {C:\nmrdata\CBDD} Zhiyong 17
0.5
1.0
1.5
2.0
2.5
3.0
3.5
4.0
4.5
5.0
5.5
5.5
5.0
4.5
4.0
3.5
3.0
2.5
2.0
1.5
1.0
0.5
0.0

HSQC of **3**

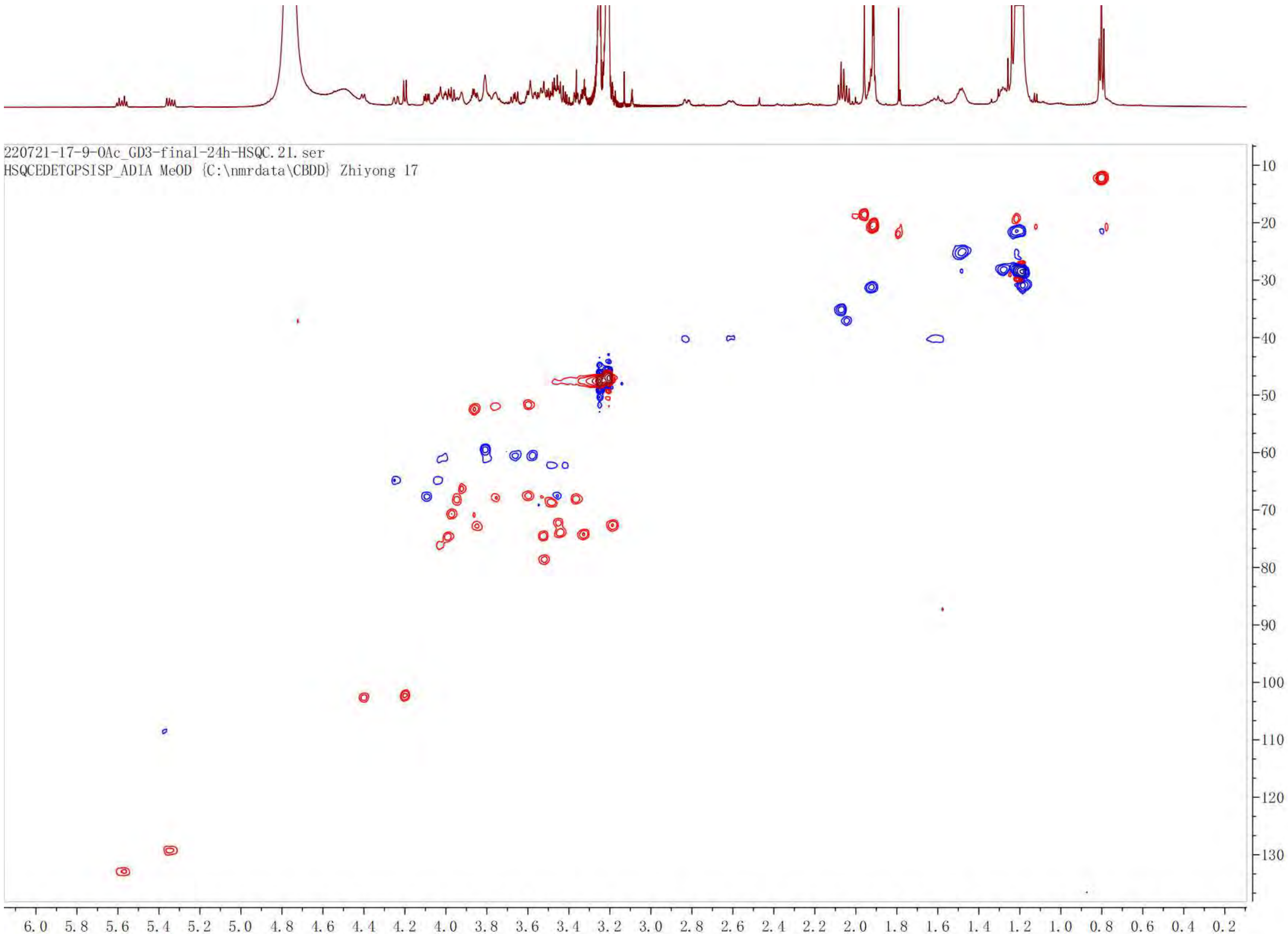

220721-17-9-OAc_GD3-final-24h-HSQC.21.ser
HSQCEDETGPSISP_ADIA MeOD {C:\nmrdata\CBDD} Zhiyong 17
10
20
30
40
50
60
70
80
90
100
110
120
130
6.0 5.8 5.6 5.4 5.2 5.0 4.8 4.6 4.4 4.2 4.0 3.8 3.6 3.4 3.2 3.0 2.8 2.6 2.4 2.2 2.0 1.8 1.6 1.4 1.2 1.0 0.8 0.6 0.4 0.2

$^{1}$H NMR of **18**. 600 MHz, $CD_3OD$.

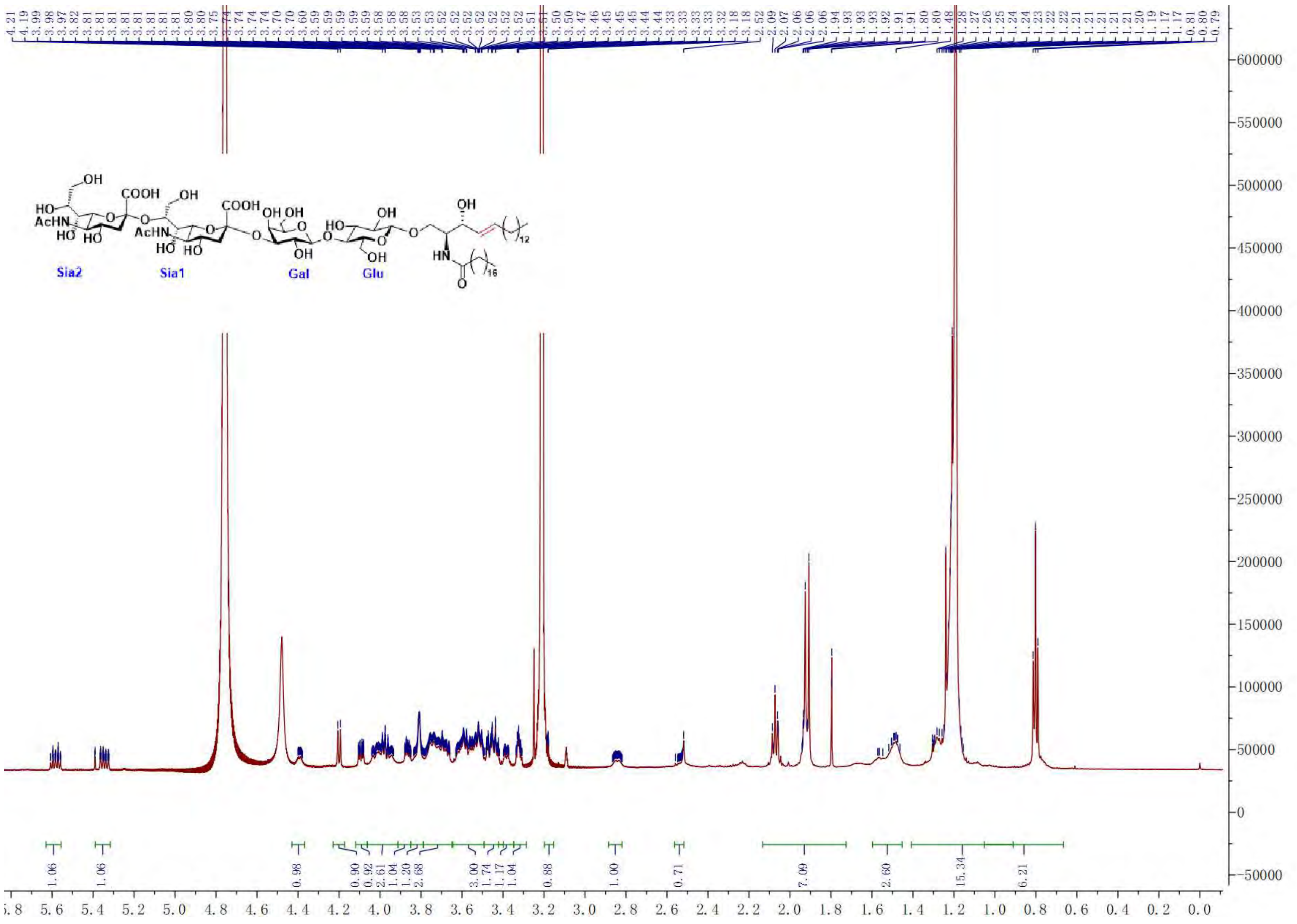

COSY of **18**

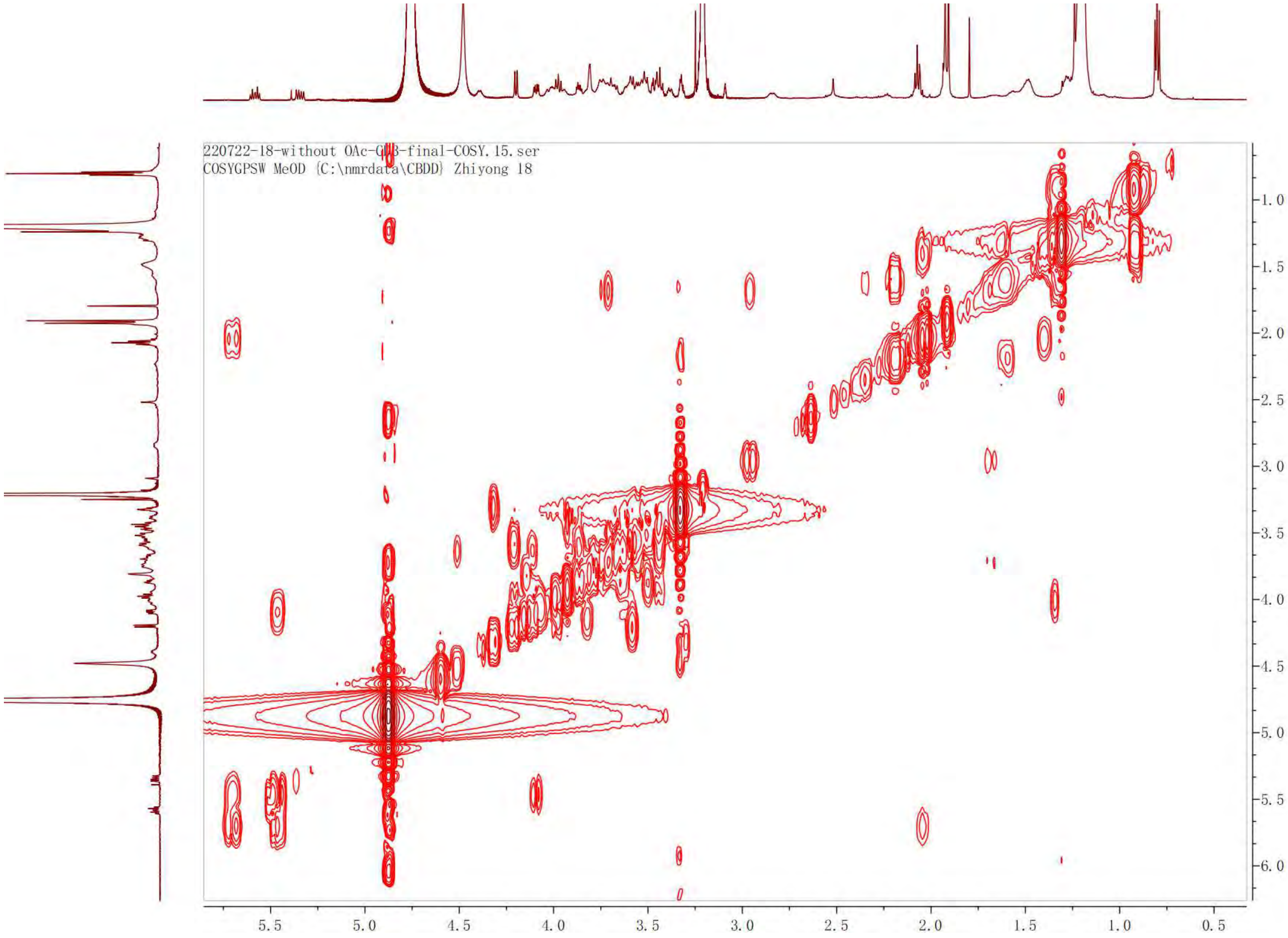

220722-18-without OAc-G[illegible]B-final-COSY.15.ser
COSYGPSW MeOD {C:\nmrdata\CBDD} Zhiyong 18
5.5
5.0
4.5
4.0
3.5
3.0
2.5
2.0
1.5
1.0
0.5
1.0
1.5
2.0
2.5
3.0
3.5
4.0
4.5
5.0
5.5
6.0

TOCSY of **18**

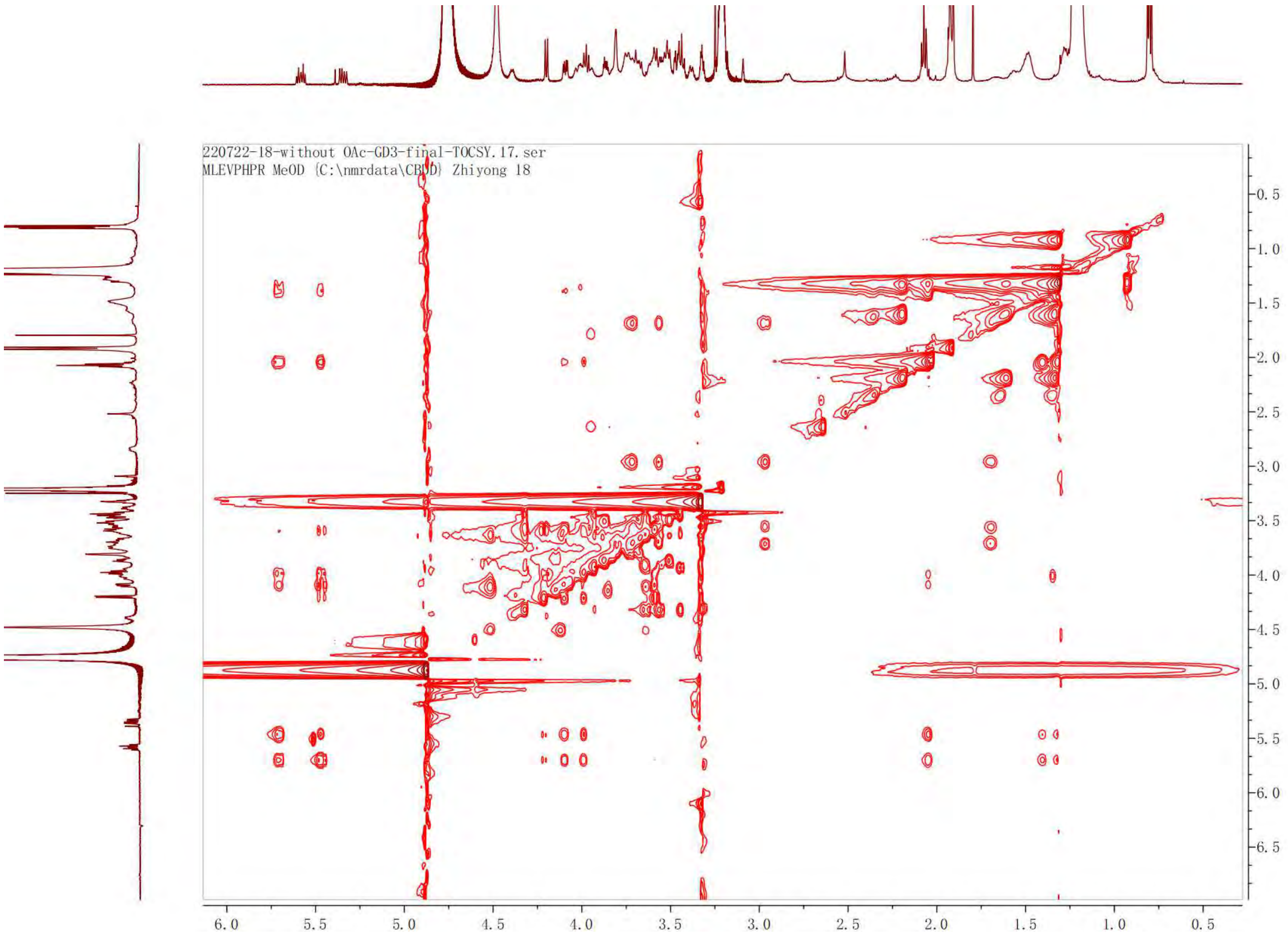

220722-18-without OAc-GD3-final-TOCSY.17.ser
MLEVPHPR MeOD {C:\nmrdata\CBDD} Zhiyong 18
0.5
1.0
1.5
2.0
2.5
3.0
3.5
4.0
4.5
5.0
5.5
6.0
6.5
6.0
5.5
5.0
4.5
4.0
3.5
3.0
2.5
2.0
1.5
1.0
0.5

HSQC of **18**

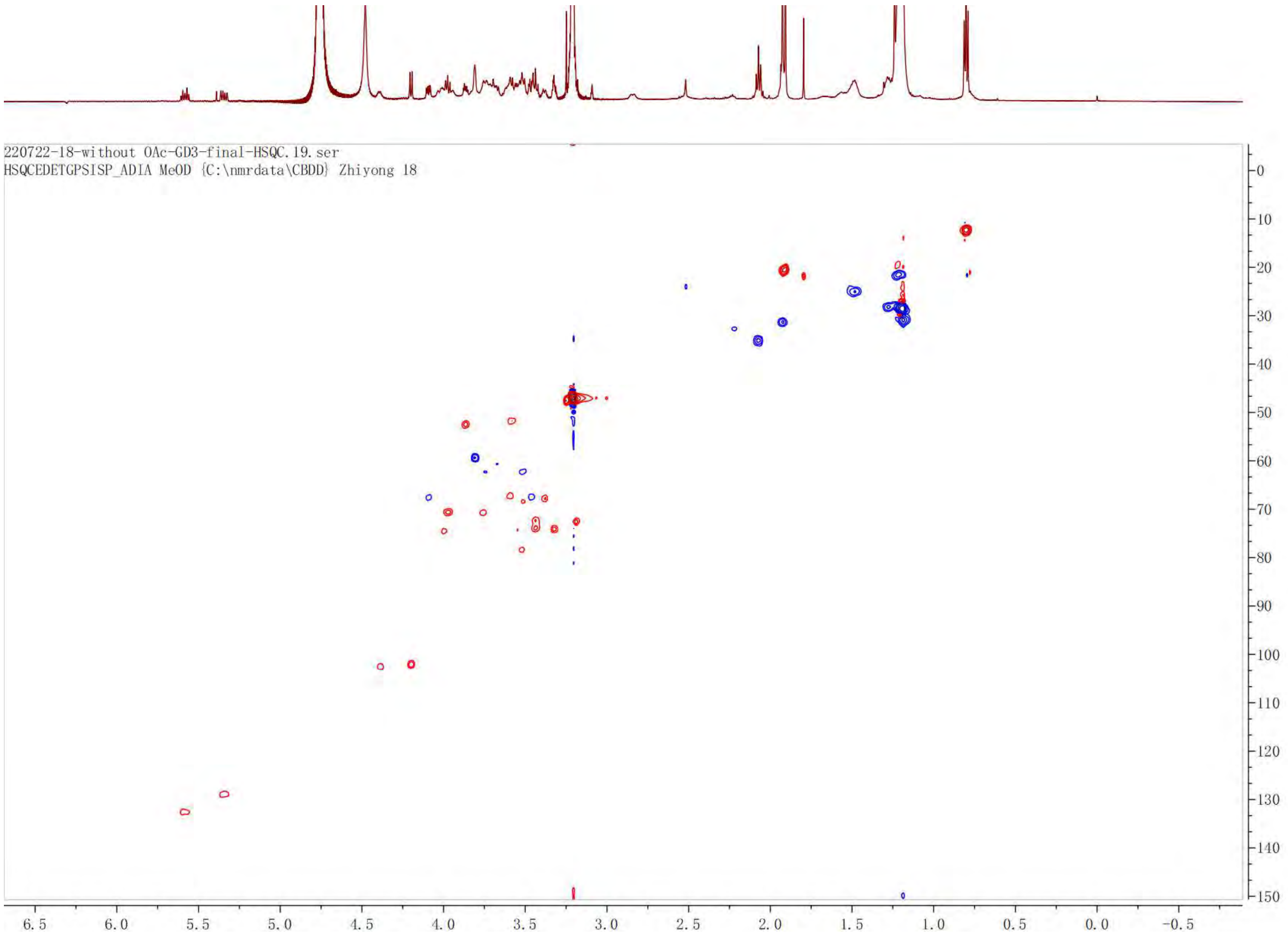
220722-18-without OAc-GD3-final-HSQC.19.ser
HSQCEDETGPSISP_ADIA MeOD {C:\nmrdata\CBDD} Zhiyong 18
0
10
20
30
40
50
60
70
80
90
100
110
120
130
140
150
6.5
6.0
5.5
5.0
4.5
4.0
3.5
3.0
2.5
2.0
1.5
1.0
0.5
0.0
-0.5
